# Models of Throughput for Multi-Cell, Multi-Type Droplet Microfluidics

**DOI:** 10.1101/2022.09.23.509249

**Authors:** William Krinsman

**Author notes:** Corresponding author: William Krinsman.

## Abstract

New experimental platforms encapsulate multiple cells per microfluidic droplet, with each cell belonging to one of multiple possible types. The motivating example comes from microbial ecology, where we want to observe the interactions of microbial strains. Because droplets are formed randomly, we want to accurately predict the data throughput, the numbers of droplets containing desired combinations of cell types.

Herein I identify the default statistical model for predicting the data throughput of multi-cell, multi-type droplet microfluidics experiments, which fits to cell type count data. I explain the assumptions behind this model and issues that in practice may cause these assumptions to fail. One such issue, “compositional heterogeneity”, is unique to multi-type experiments. I show how to modify the default statistical model to describe the consequences of these issues, without needing to mechanistically model their causes.

In practice, only two of these issues may substantially change the data throughput predictions. The changes depend on both (1) which combination of these issues are present, and (2) the precise definition of data throughput. Finally, I show that for a given experimental platform one can estimate the severity of these two issues, enabling more accurate data throughput predictions that account for these two issues.

## INTRODUCTION

This paper highlights the work presented in Part II of the PhD Thesis (Krinsman, 2022).

Droplet microfluidics has proven useful for single-cell, single-type experiments. See, for example, (Lagus and Edd, 2013) or (Matula et al., 2020), for a review. New experimental platforms, such as kChip (Kehe et al., 2019) or MINI-Drop (Hsu et al., 2019), use droplet microfluidics to probe the interactions of multiple types of cells. For kChip and MINI-Drop specifically the motivation is to enable more systematic explorations of microbial interactions. Microbial ecology would benefit from experimental, rather than merely observational, data characterizing the interactions of large numbers of microbial strains. The current state of the art for such experiments, by manually culturing and plating combinations of cell types, probes the interactions of fewer than 20 microbial strains simultaneously (Venturelli et al., 2018). Because droplet microfluidics chips produce large numbers of microfluidic droplets, multi-cell, multi-type droplet microfluidics experimental platforms potentially could drastically increase the number of microbial strains whose interactions can be probed simultaneously.

A major tradeoff associated with this approach, compared to manually culturing and plating combinations of cell types, is that droplet microfluidics chips form microfluidic droplets randomly. Cf. Supplementary Figure S4. Therefore, even starting with known microbial communities, it is a priori unclear approximately how many droplets will be produced containing any given combination of cell types. This paper aims to introduce a framework that fills this gap. Doing so will increase the relative advantage of multi-cell, multi-type droplet microfluidics experimental platforms over manually culturing and plating combinations of cell types. Hopefully this can abet future dramatic increases in the state of the art for the number of microbial strains whose interactions can be probed simultaneously.

### Notation

Let *S* denote the total number of cell types in the sampling pool. In the motivating microbial ecology example, “*S*” for “**s**trains” is the total number of microbial strains in the microbial community sample.

Given a positive integer 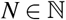, define [*N*]:= {1,…, *n,…,N*}. Unless specified otherwise, given a (non-negative) vector 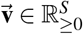, its entries are denoted *v^(s)^*, i.e.

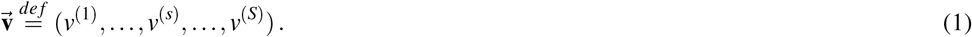

The shorthand 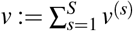 is used for the sum of the entries of 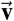.

For a given droplet, the random variable *N*(0) is the number of cells from the population encapsulated in the droplet during its formation (“*N*” for “**n**umber”). Equivalently, *N*(0) is the number of cells at “time 0”. This is the sum of the entries of the random vector:

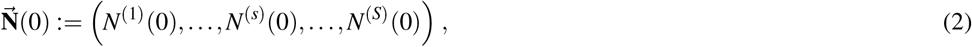

where for each type *s* (“*s*” for “**s**train”) of the S total types, *N^(s)^*(0) is the random variable equalling the number of cells from type *s* at time 0.

For each cell type *s*, let *f*^(*s*)^ (“*f*“ for “**f**requency”) denote the relative abundance of cell type *s* in the sampling pool. In particular, we necessarily have that 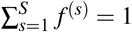.

### Precise Definition of Throughput

The below sections give two different possible definitions of throughput. Cf. supplementary section S4.2.1 for mathematically rigorous versions of these definitions.

#### Gluttonous Definition of Data Throughput

The treatment group 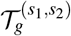 for inferring the effect of type *s*_1_ on the growth of type *s*_2_ corresponds to droplets where *s*_1_ and *s*_2_ co-occur. This is called the “**gluttonous**” **treatment** group because *all* droplets where *s*_1_ and *s*_2_ co-occur are included. The control group 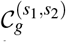 for inferring the effect of type *s*_1_ on the growth of type *s*_2_ corresponds to droplets where *s*_2_ occurs but *s*_1_ does not. This is called the “**gluttonous**” **control** group because *all* droplets where *s*_2_ occurs but *s*_1_ does not are included. Cf. figure 1.

**Figure 1.**
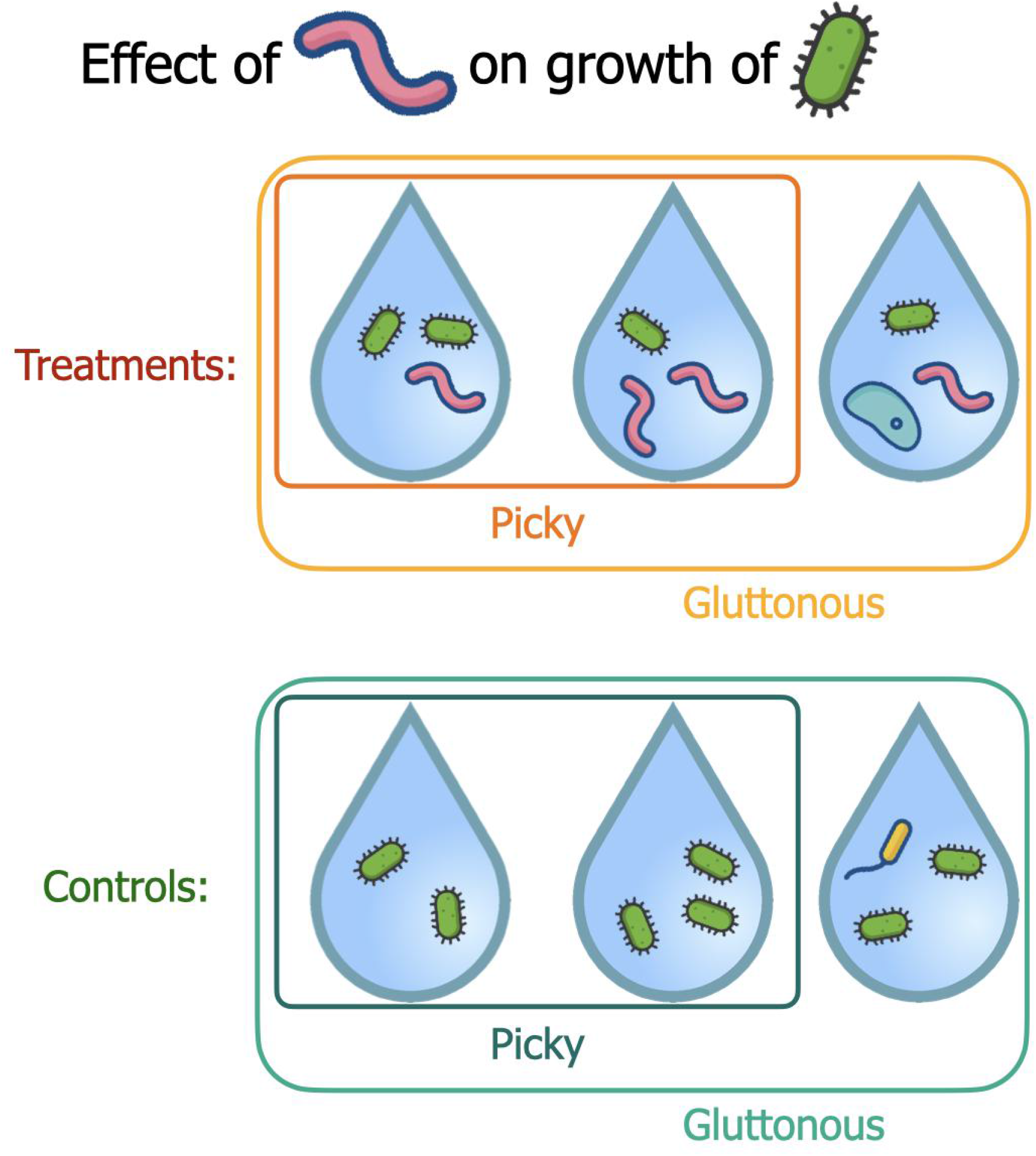
Gluttonous vs. Picky Definitions of Data Throughput. Cf. text.

These definitions place no constraints on the presence or absence of types *s* ∈ [*S*] besides *s*_1_ and *s*_2_. This can be favorable for example when studying very rare types, for which there may be no or very few droplets containing *s*_1_ and *s*_2_ only, but some containing *s*_1_ and *s*_2_ along with other types. These definitions give us the best possible chance of avoiding “data starvation”.

#### Picky Definition of Data Throughput

The above “gluttonous” definitions correspond to more possible combinations of cell types than we probably would have included when studying the effect of type *s*_1_ on the growth of type *s*_2_ by manually plating combinations of cell types. This motivates the following definitions.

The set of droplets where *s*_1_ and *s*_2_ co-occur, and all other types are absent, is called the “**picky**” **treatment** group 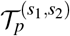. The set of droplets where *only s*_2_ occurs, and all other types are absent, is called the “**picky**” **control** group 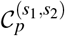. Cf. figure 1.

Unlike the gluttonous groups, the picky groups *do* place constraints on the presence or absence of types *s* ∈ [*S*] besides *s*_1_ and *s*_2_. For both the treatments and the controls, for “picky” groups all types besides *s*_1_ or *s*_2_ must have zero counts. Every picky group is a subset of its gluttonous counterpart.

The picky definitions can be favorable when we have plenty of droplets from which to make estimates. They reduce the possibility of confounding effects on the growth of type *s*_2_ that could be caused by types that are not type *s*_1_. On the other hand, in instances where there are only very few or no droplets without “extra” types, the picky definitions could lead to “data starvation”.

### Statistical Motivation

The treatment group 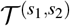 for the effect of type *s*_1_ on type *s*_2_ consists of droplets containing both types. Likewise, the control group 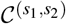 for the effect of type *s*_1_ on type *s*_2_ consists of droplets containing type *s*_2_ but not type *s*_1_. With these groups, we can quantify the effect of type *s*_1_ on type *s*_2_ by targeting an average treatment effect (ATE) or other similar causal estimand.

Even when we can implement estimators for such causal estimands, sometimes the results may be unreliable. Consider e.g. a situation where our treatment group 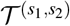 consists of only **T**^(*s*_1_,*s*_2_)^ = 5 droplets and our control group 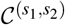 consists of **C**^(*s*_1_,*s*_2_)^ ≈ 50,000 droplets. This example may seem unrealistic, but it is actually optimistic compared to some of the disparities between the treatment and control group sizes **T**^(*s*_1_,*s*_2_)^, **C**^(*s*_1_,*s*_2_)^ that one may expect to see in practice. Cf. the discussion in (Krinsman, 2022, Appendix 2.C). Regardless of how we choose to quantify interaction effects, we need to estimate the reliability of our estimates in order for them to be scientifically useful. We need to estimate their statistical power 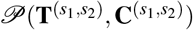.

Of course (even for a fixed causal estimand) the statistical power 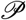 function will depend on the particular estimator, so at this level of generality we cannot complete such a calculation. However, a priori we do also know that the statistical power 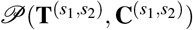 will always depend on the sizes **T**^(*s*_1_,*s*_2_)^, **C**^(*s*_1_,*s*_2_)^ of the treatment group and control group, respectively. Thus, to estimate statistical power 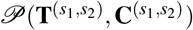, we always need to know the treatment group and control group sizes **T**^(*s*_1_,*s*_2_)^, **C**^(*s*_1_,*s*_2_)^, regardless of the particular estimator.

Unfortunately, due to the random nature of droplet formation in droplet-based microfluidics experiments, the scientist can never directly control the treatment group and control group sizes **T**^(*s*_1_,*s*_2_)^, **C**^(*s*_1_,*s*_2_)^. The treatment group and control group sizes **T**^(*s*_1_,*s*_2_)^, **C**^(*s*_1_,*s*_2_)^ are random variables. Therefore, for any estimator of any causal estimand, the statistical power 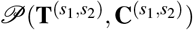 will itself be a random variable. This is not inherently a “deal-breaker”, but we do still need to constrain the possibilities for the distribution of the statistical power 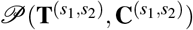. At the very least, we would like to be able to compute the expected statistical power 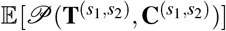.

Specifically, for a given estimator of a given causal estimand, the statistical power 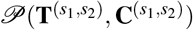 will be a deterministic function 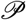 of the treatment and control group sizes **T**^(*s*_1_,*s*_2_)^, **C**^(*s*_1_,*s*_2_)^ (and possibly other factors which for simplicity we will assume to be known and fixed). If we know specific values **T**^(*s*_1_,*s*_2_)^ = *T*^(*s*_1_,*s*_2_)^, **C**^(*s*_1_,*s*_2_)^ = *C*^(*s*_1_,*s*_2_)^ for the treatment and control group sizes **T**^(*s*_1_,*s*_2_)^, **C**^(*s*_1_,*s*_2_)^, then we can evaluate this statistical power function 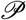 and get a deterministic value 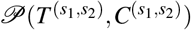. If instead the treatment and control group sizes **T**^(*s*_1_,*s*_2_)^, **C**^(*s*_1_,*s*_2_)^ are random variables, then when evaluating this statistical power function we get a new random variable 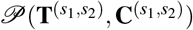 whose distribution is determined by the pushforward measure.

Obviously, without specifying the particular causal estimand and its particular estimator, we still cannot complete the calculation of the expectation 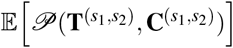 of a random statistical power 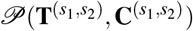. Nevertheless, for any given estimator of any given causal estimand, the distribution of the statistical power 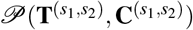 will be determined by the distribution of the treatment group and control group sizes **T**^(*s*_1_,*s*_2_)^, **C**^(*s*_1_,*s*_2_)^. Thus, for any estimator of any causal estimand, as a prerequisite for being able to estimate expected statistical power 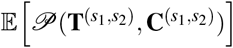, we must first understand the distribution of the treatment group and control group sizes **T**^(*s*_1_,*s*_2_)^, **C**^(*s*_1_,*s*_2_)^.

Note that the treatment group and control group sizes **T**^(*s*_1_,*s*_2_)^, **C**^(*s*_1_,*s*_2_)^ can be understood as sums of indicator random variables:

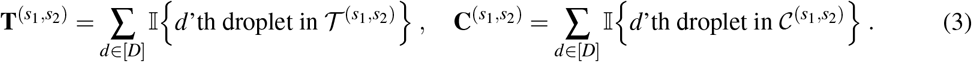

To completely characterize the distribution of indicator random variables it suffices to know their expectations, i.e. the probabilities:

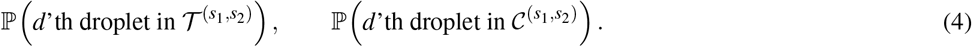

If we assume the droplets are identically distributed, then linearity of expectation allows us to compute the expected treatment and control group sizes:

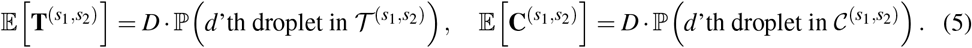

This is already enough to compute 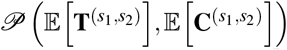, a quantity that we can use as a (potentially suboptimal) stand-in for the expected statistical power 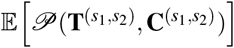 in situations where we don’t want to make additional assumptions about the distributions of the droplets.

However, if we additionally assume that the droplets are mutually independent, then we can compute the distributions of the treatment and control group sizes as marginals of a Multinomial distribution^1^. Given the statistical power function 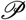, i.e. choice of a particular causal estimand and estimator thereof, this would then in principle^2^ be enough information to compute the distribution of the statistical power 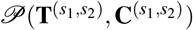 or any summary statistic thereof.

One might object to treating the statistical power 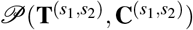 as a random variable. Certainly, after the experiment has been run, the treatment group and control group sizes **T**^(*s*_1_,*s*_2_)^, **C**^(*s*_1_,*S*_2_)^ will be known, allowing us to condition on their observed values *T*^(*s*_1_,*s*_2_)^, *C*^(*s*_1_,*s*_2_)^ and “restore determinism” by evaluating 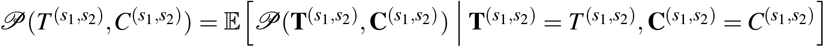. Yet this is insufficient to help the scientist before running the experiment. The scientist has *no* direct control in advance over what exactly the treatment group and control group sizes **T**^(*s*_1_,*s*_2_)^, **C**^(*s*_1_,*s*_2_)^ will turn out to be.

However, the distribution of the treatment group and control group sizes **T**^(*s*_1_,*s*_2_)^, **C**^(*s*_1_,*s*_2_)^ should depend on deterministic factors that the scientist can directly control, like the total number of droplets *D*, the expected number of cells per droplet *λ*, and the relative abundances *f^(s)^* of the various types *s* in the sampling population. The scientist would like to plan the experiment (*before* running it) in a way that allows them to (most likely) achieve certain thresholds of statistical power 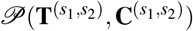 for certain interactions of types. Therefore the best way to help the scientist is to predict which values of statistical power 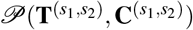 they will most likely be able to achieve as a function of the deterministic factors that the scientist can directly control. From that perspective it is inescapable that

1. the statistical power 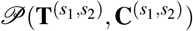 must be treated as random,
2. the most basic prerequisite for constraining the distribution of the statistical power 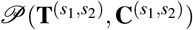 is to characterize the distributions of the treatment group and control group sizes **T**^(*s*_1_,*s*_2_)^, **C**^(*s*_1_,*s*_2_)^, and
3. (if possible) the distributions of the treatment group and control group sizes **T**^(*s*_1_,*s*_2_)^, **C**^(*s*_1_,*s*_2_)^ should be described as functions of the deterministic factors that the scientist can directly control.

There is no need to incubate the droplets in order to estimate the probabilities (4). Indeed, it should be possible to estimate the probabilities using a much smaller number of total droplets *D* than what might be required in the final experiment to achieve certain (expected) statistical power thresholds. Therefore I propose that scientists, to be able to design the experiment in a way that allows them achieve certain (expected) statistical power thresholds, first run a “*t* = 0” version of the experiment that skips incubating the droplets and uses a smaller number of total droplets *D*. This both (i) maximizes the usefulness of the data for estimating the distributions of the treatment group and control group sizes (because it should mitigate artifacts resulting from microbial growth and competition, e.g. censoring) and (ii) decreases the cost (in money and time) of this “pre-experiment”.

### A Default Working Model: hPoMu (hierarchical Poisson Multinomial)

The distributions of the “hierarchical Poisson Multinomial” (hPoMu) working model are hierarchical distributions. The total number of cells in the droplet has a Poisson distribution, while *(conditional upon the total number of cells)* the numbers of cells belonging to the multiple types have a joint Multinomial distribution. Cf. figure S5. Therefore the likelihood for this family of probability distributions equals

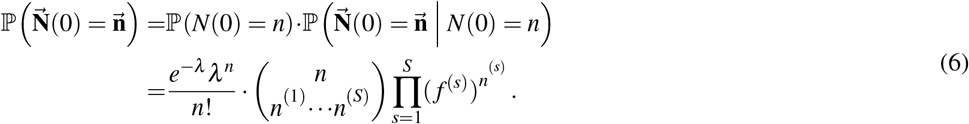

An alternative form of expressing the likelihood for hPoMu, which reveals that its marginal distributions correspond to mutually independent Poisson-distributed random variables, is

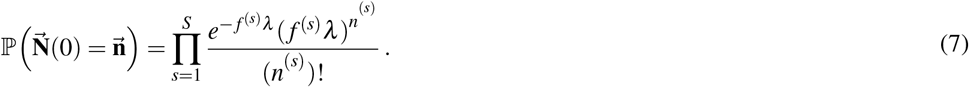

Heuristically speaking, equation (7) says that during the formation of each droplet the sampling rate *λ* is evenly spread out among the multiple types according to their frequencies in the population.

### Assumptions Justifying Default Working Model

Both of the two derivations justifying hPoMu as the default working model, cf. supplementary section S2.3.2, make the following implicit assumptions (among others):

- For *every* type s, the number of cells in the population which belong to type s is very large.
- Each cell (regardless of its type) has the same small probability of ending up in the droplet as any other cell.
- Whether a given cell (regardless of its type) ends up in the droplet is completely independent of what happens to any other cell.

The first can be thought of as stating that the population size for each type is effectively infinite. The second and third can together be thought of as stating that the sampling pool is perfectly homogeneous.

### Failures of Default Working Model Assumptions

While hPoMu is definitely a sensible working model to start with, its implicit assumptions failing to be realistic could undermine its usefulness in practice.

#### Populations are Finite

We assumed that *for each individual type s* the number of cells is “effectively infinite”. For “rare” types it is a priori unclear whether this assumption is reasonable. However in practice, even for types with relative abundance as low as 0.01%, taking the finiteness of the populations into account does not lead to substantially different predictions than the default hPoMu model. (Cf. sections S3.4.1 and S4.4.1)

#### Density Heterogeneity

In practice different sections of the sampling pool could conceivably have average numbers of cells per unit volume that differ from the average number of cells per unit volume for the entire sampling pool (i.e. the total number of cells divided by the volume of the sampling pool). Herein I call this phenomenon “density heterogeneity”. Higher density heterogeneity implies more variance in the numbers of cells per unit volume throughout the sampling pool. Cf. figures 2 and 3.

**Figure 2.**
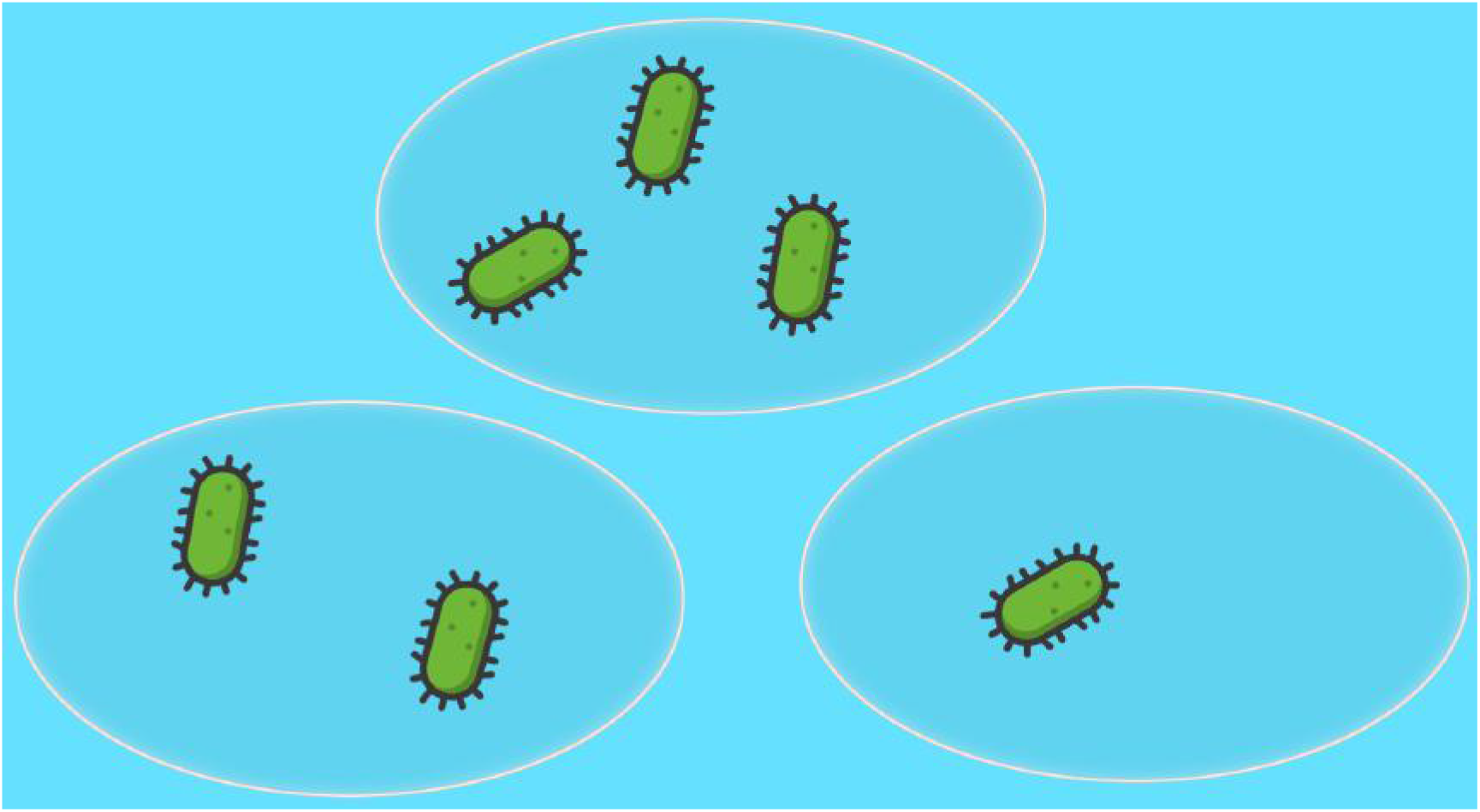
Lower Density Heterogeneity. The average number of cells per unit volume is 2.

**Figure 3.**
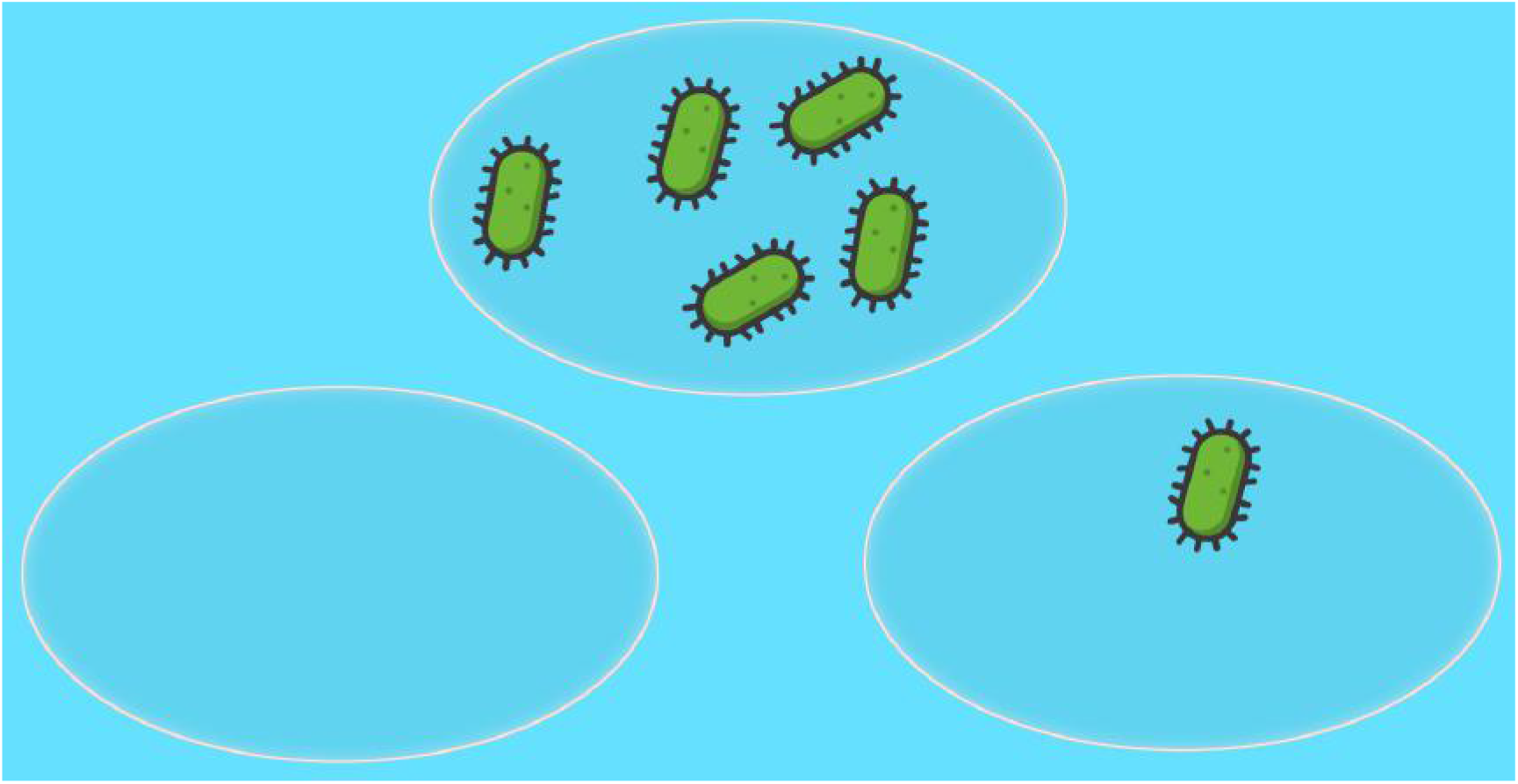
Higher Density Heterogeneity. The average number of cells per unit volume is still 2.

The default Poisson model for the distribution of the total number of cells in each droplet does not account for additional variance which might be introduced in practice by density heterogeneity.

Because this describes extra variance in the total number of cells, regardless of their types, density heterogeneity is relevant even when there is only one type. Thus density heterogeneity is relevant even for the single-cell^3^, single type experiments which have already been intensively developed.

#### Compositional Heterogeneity

Although the relative abundance of any given cell type *s* across the entire sampling pool is a fixed value *f^(s)^*, conceivably different sections of the sampling pool could have relative abundances for the cell types that differ from the relative abundances for the entire sampling pool. Herein I call this phenomenon “compositional heterogeneity”. Higher compositional heterogeneity implies more variance in the relative abundances for the cell types throughout the sampling pool. Cf. figures 4 and 5.

**Figure 4.**
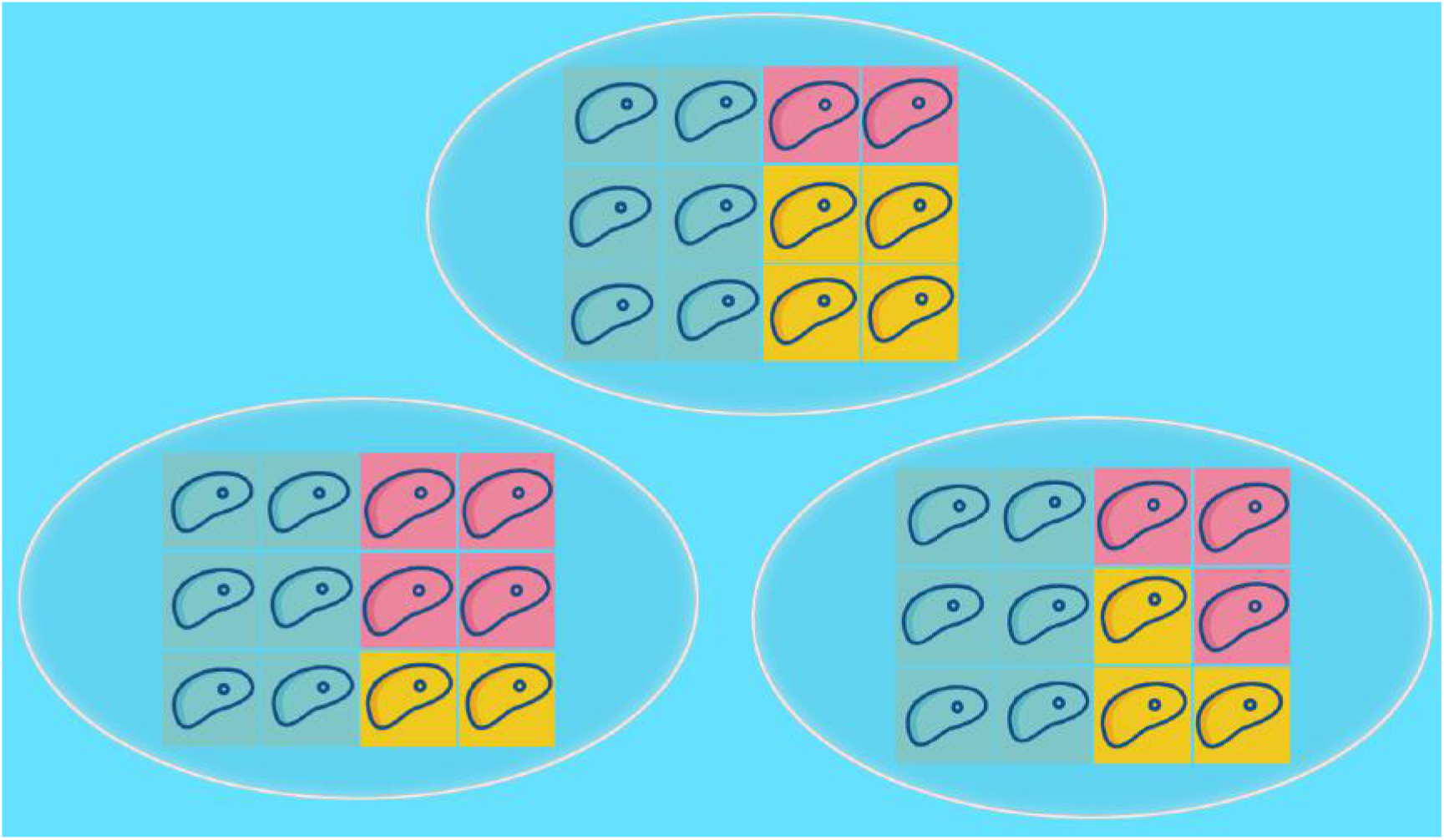
Lower Compositional Heterogeneity. The average relative abundances of each cell type for the entire sampling pool are 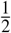 blue, 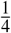 pink, 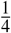 yellow.

**Figure 5.**
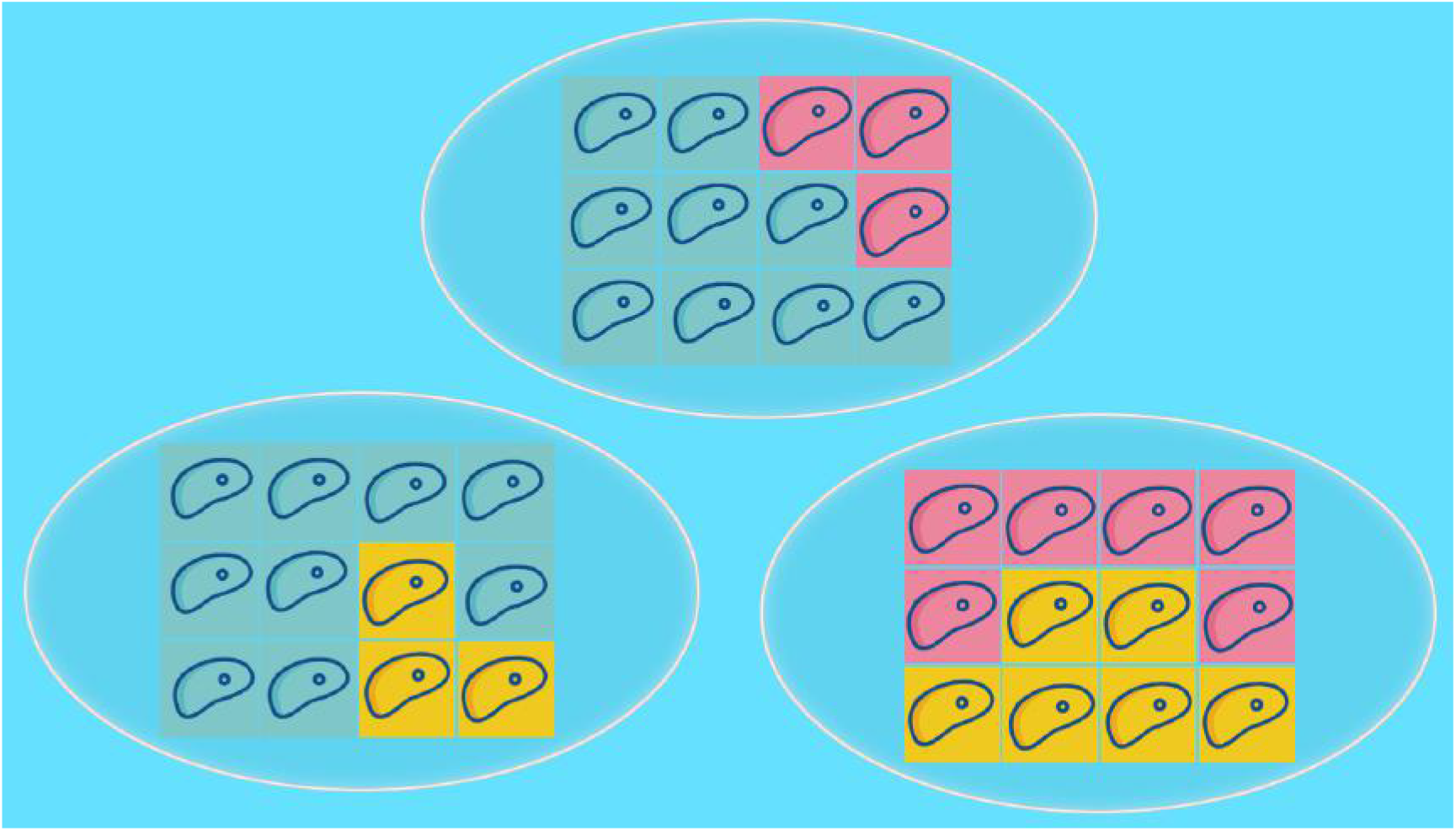
Higher Compositional Heterogeneity. The average relative abundances of each cell type for the entire sampling pool are still 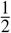 blue, 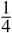 pink, 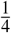 yellow.

The default multinomial model for the relative abundances of the cell types in the droplet (*conditional upon the total number of cells*) does not account for additional variance that might be introduced in practice by compositional heterogeneity.

Because compositional heterogeneity describes extra variance in the relative abundances of the cell types, compositional heterogeneity is relevant only for multi-cell, multi-type experiments. Because such experiments are newer and still less common than single-cell, single type experiments, it seems that compositional heterogeneity may have never before been identified as a potential issue affecting the data throughput of droplet microfluidics experiments.

### Working Model Accounting for Assumption Failures: ghNBDM

This working model describes how excess variances relative to the completely homogeneous situation (modelled by hPoMu) could affect data throughput. This working model is not a mechanistic model and it does not predict how physical processes create those excess variances. (Cf. supplementary section S2.4.4.) The density concentration *ζ_D_* and compositional concentration *ζ_C_* parameters specify this description of the excess variances. These parameters are separate from and not functions of the relative abundances of the cell types *f^(s)^* nor the mean number of cells per droplet *λ*. This working model can be fit to real data of cell type counts using the procedures from supplementary section S5 for estimating the density concentration *ζ_D_* and compositional concentration *ζ_C_* parameters. That alone is sufficient to access the working model’s predictions for how excess variances relative to the completely homogeneous situation could affect data throughput.

The “generalized hierarchical Negative Binomial Dirichlet-Multinomial” (ghNBDM) family is a family of hierarchical distributions. The number of cells in the droplet is Negative Binomial distributed with “density concentration” parameter *ζ_D_S*, while *(conditional upon the total number of cells)* the numbers of cells belonging to the multiple types have a joint Dirichlet-Multinomial distribution with “compositional concentration” parameter *ζ_C_S*. Thus the likelihood equals

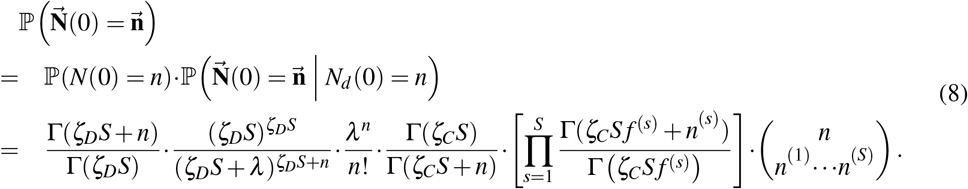

In general the marginal distributions of members of the ghNBDM family have non-zero cross-covariance. Thus the entries *N^(s)^*(0) of the random vector 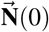 will in general *not* be mutually independent. This is unlike the default working model (hPoMu), cf. again equation (7).

Notice how this working model has two additional parameters compared to the hPoMu working model, namely the density concentration *ζ_D_* parameter and the compositional concentration *ζ_C_* parameter. Higher values of the density concentration *ζ_D_* parameter correspond to lower density heterogeneity (cf. supplementary section S2.7). As *ζ_D_* → ∞ the total number of cells in the droplet approaches a Poisson distribution. Cf. section S2.7.2. Similarly, higher values of the compositional concentration *ζ_C_* parameter correspond to lower compositional heterogeneity (cf. supplementary section S2.6). As *ζ_C_* → ∞ the joint distribution of the numbers of cells belonging to the multiple types approaches a Multinomial distribution. Cf. supplementary section S2.6.2. The hPoMu model is approached in the limit as both *ζ_D_* → ∞ and *ζ_C_* → ∞ jointly. Cf. supplementary figure S11 and supplementary section S2.8.1.

For details about the calculations demonstrating how the somewhat complicated expressions involving the Γfunction do indeed generalize the likelihoods of the Poisson and Multinomial distributions, cf. sections S2.7.2, S2.6.2, and S2.10. At a high level, the idea is to use Stirling’s approximation, in particular the formulations in terms of explicit bounds given by (Robbins, 1955) and (Gordon, 1994).

## METHODS

I computed and stored results using NumPy (Harris et al., 2020) version 1.20.2. I plotted results using Matplotlib (Hunter, 2007) version 3.4.1 and Seaborn (Waskom, 2021) version 0.11.1. Kernel density estimates were made using the default Seaborn (Waskom, 2021) settings^4^. The bar plot (Figure 6) was made using the *microbiome* (Lahti and Shetty, 2019), *phyloseq* (McMurdie and Holmes, 2013), and *ggplot2* (Wickham, 2016) packages for the *R* programming language. Complete implementation details can be found in the code at the relevant GitLab repository. See https://gitlab.com/krinsman/droplets.

**Figure 6.**
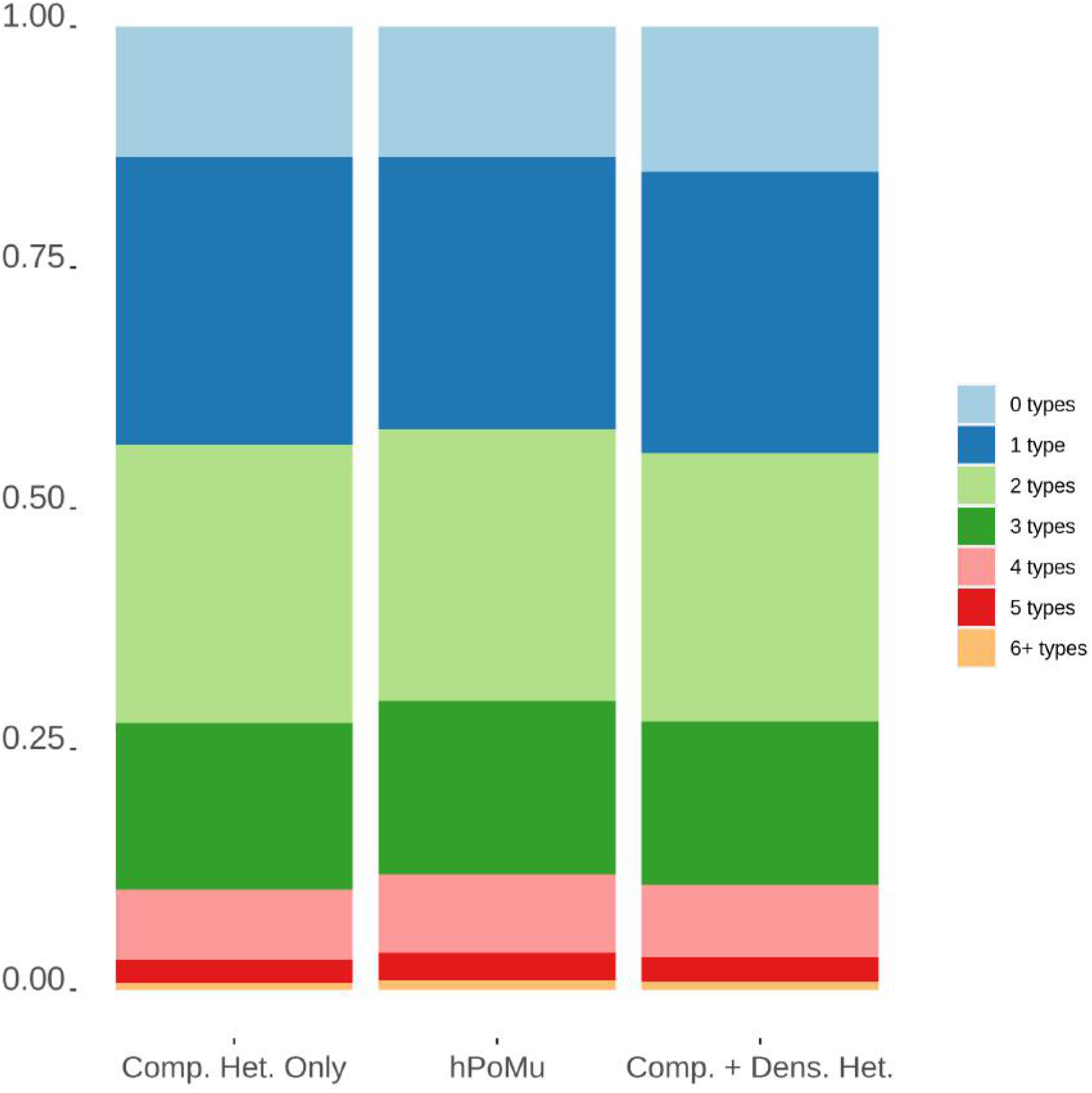
“Comp. Het. Only” = Compositional Heterogeneity Only. “Comp. + Dens. Het.” = Compositional and Density Heterogeneities. Cf. Supplementary Figure S36.

### Simulation Definitions

For all simulations and distributions, the same “simulated community” of 91 types was used. These correspond to 90 types distributed across 9 distinct relative abundances: 10 types for each of .01%, .02%, .05%, .1%, .2%, .5%, 1%, 2%, and 5%. The 91st type was a “remainder” type with abundance ≈12%. All distributions used the same value of the rate parameter, *λ* = 2, the expected total number of cells for each droplet. Each distribution was statistically independently simulated 500 times, with each simulation having 15 million statistically independently simulated droplets. See section S3.3 for details.

### Numbers of Droplets with *n* Types

For details about how the expected values under hPoMu were computed, see section S4.3.1. For each simulated distribution, I stratified and then counted the droplets from each of the 500 simulations according to how many types were present. I then computed the average values for each distribution as the arithmetic mean over the 500 simulations. I then computed the average percent change for each distribution relative to hPoMu using the computed expected values under hPoMu.

### Gluttonous Groups

For details about how the expected values under hPoMu were computed, see sections S4.3.2 and S4.2.1. For each simulated distribution, for each of the 500 simulations I computed via brute force the number of occurrences of each of the gluttonous groups, and then reported the arithmetic means over all 500 simulations. I again computed the average percent change as the observed value minus the value expected under hPoMu divided by the value expected under hPoMu.

### Picky Groups

Using the formulae (S64) and (S65), I computed the expected count for each picky group under hPoMu. The procedure was then otherwise exactly the same as for the gluttonous groups.

### Computing Estimator Distributions

For each simulated distribution, for each of the 500 simulations, for each of the three batch sizes (small = 10, 000 droplets, medium = 500, 000 droplets, large = 15, 000, 000 droplets), I partitioned the simulation results into equal parts of the given batch size. Each simulation contains 15, 000, 000 droplets, so this corresponds to 1, 500 small batches per simulation, 30 medium batches per simulation, and 1 large batch per simulation. Then for each of the batches, I computed the maximum likelihood density concentration estimate and the maximum likelihood compositional concentration estimate and stored the results. For details about how maximum likelihood estimators were computed, see sections S5.3.2 and S5.2.

## RESULTS

Compositional heterogeneity, which is unique to multi-type experiments, differently affects gluttonous and picky data throughput, depending on the presence or absence of density heterogeneity. These differences can be explained by breaking down the changes according to the number of types in each droplet.

### Compositional Heterogeneity Decreases Number of Droplets with Three or More Types

From figures 6 and 7 we see that compositional heterogeneity decreases the number of droplets with three or more types in both absolute and relative terms. Since droplets with three or more types are a component of the gluttonous groups, their decrease with increased compositional heterogeneity explains the decreases in gluttonous throughput visible in figures 8, and 9. We can also see from figures 7, 8, and 9 that the decrease compositional heterogeneity causes in the number of droplets with three or more types is more pronounced in the absence of density heterogeneity.

**Figure 7.**
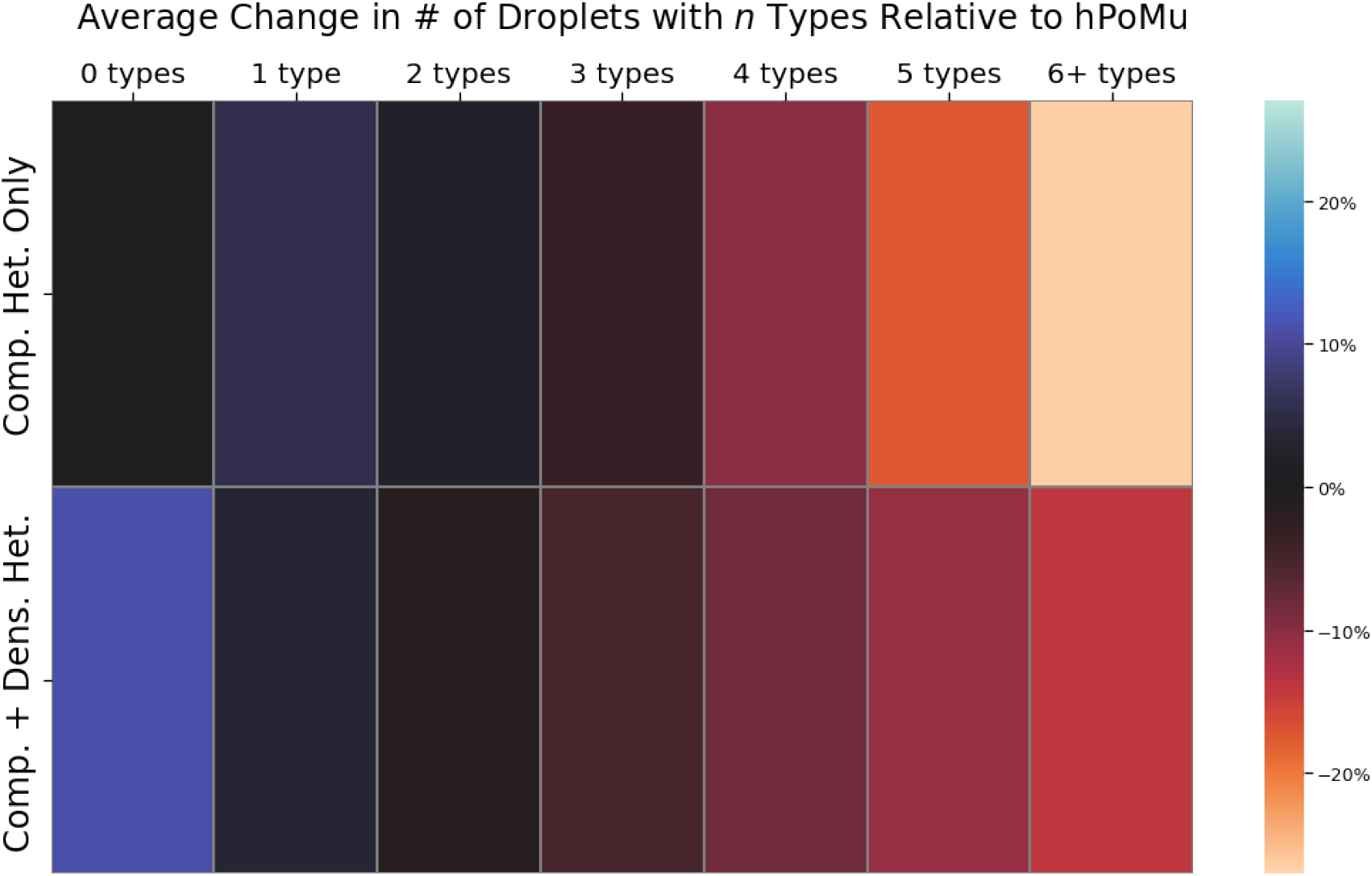
“Comp. Het. Only” = Compositional Heterogeneity Only. “Comp. + Dens. Het.” = Compositional and Density Heterogeneities. Cf. Supplementary Figure S37.

**Figure 8.**
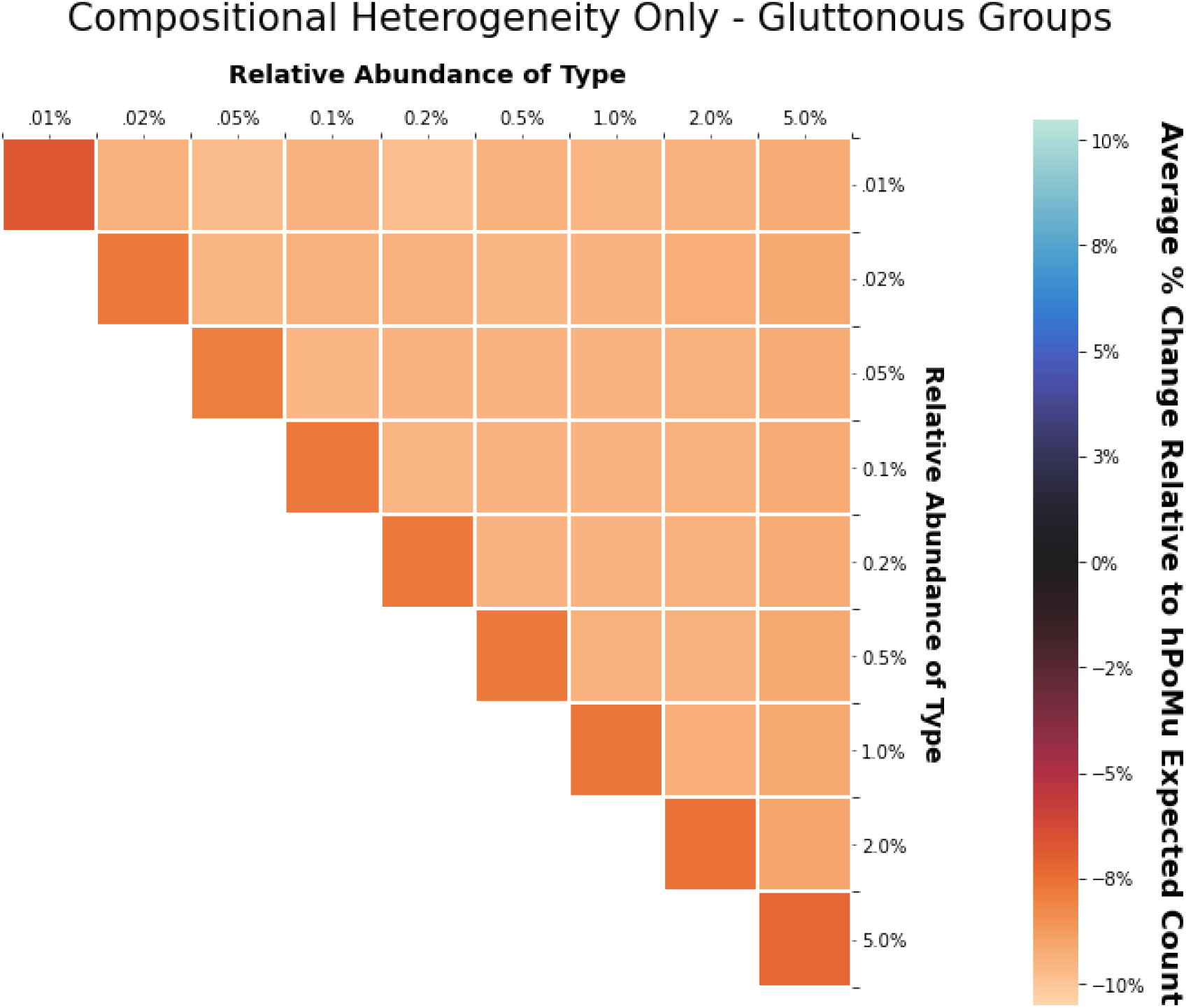
Cf. Supplementary Figure S45.

**Figure 9.**
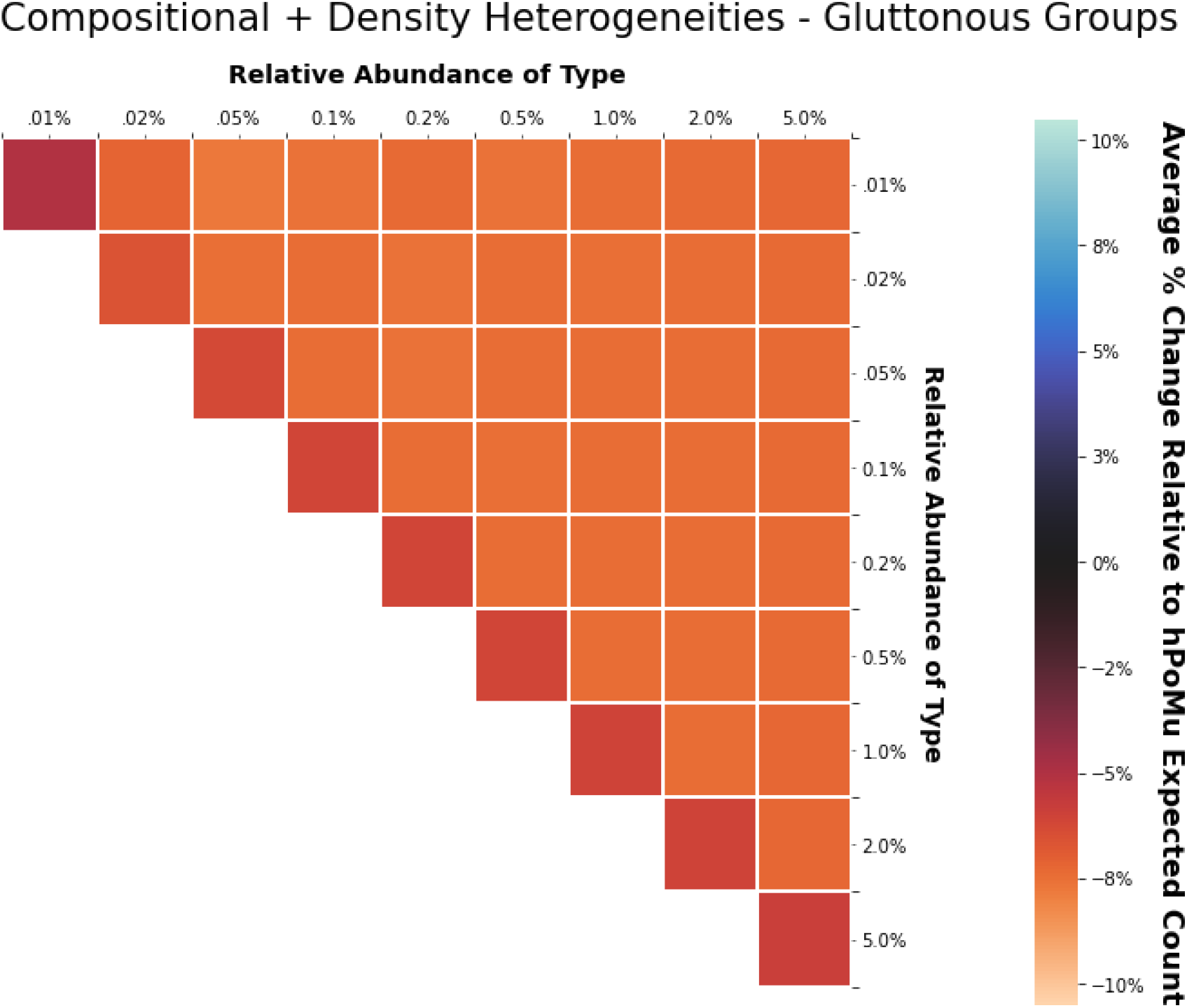
Cf. Supplementary Figure S47.

### Effect of Heterogeneities on Droplets with Two or Fewer Types

How compositional heterogeneity affects the numbers of droplets with two or fewer types depends on whether density heterogeneity is also present.

#### Compositional Heterogeneity Only

In the absence of density heterogeneity, compositional heterogeneity increases the number of droplets with one type. Compositional heterogeneity also causes a smaller increase in the number of droplets with two types. This is most visible in figure 7. This slight increase in the number of droplets with two types appears to cause the slight increase in the number of picky treatments visible in figure 10. As is clear also from figure 6, there is little change in the number of empty droplets.

**Figure 10.**
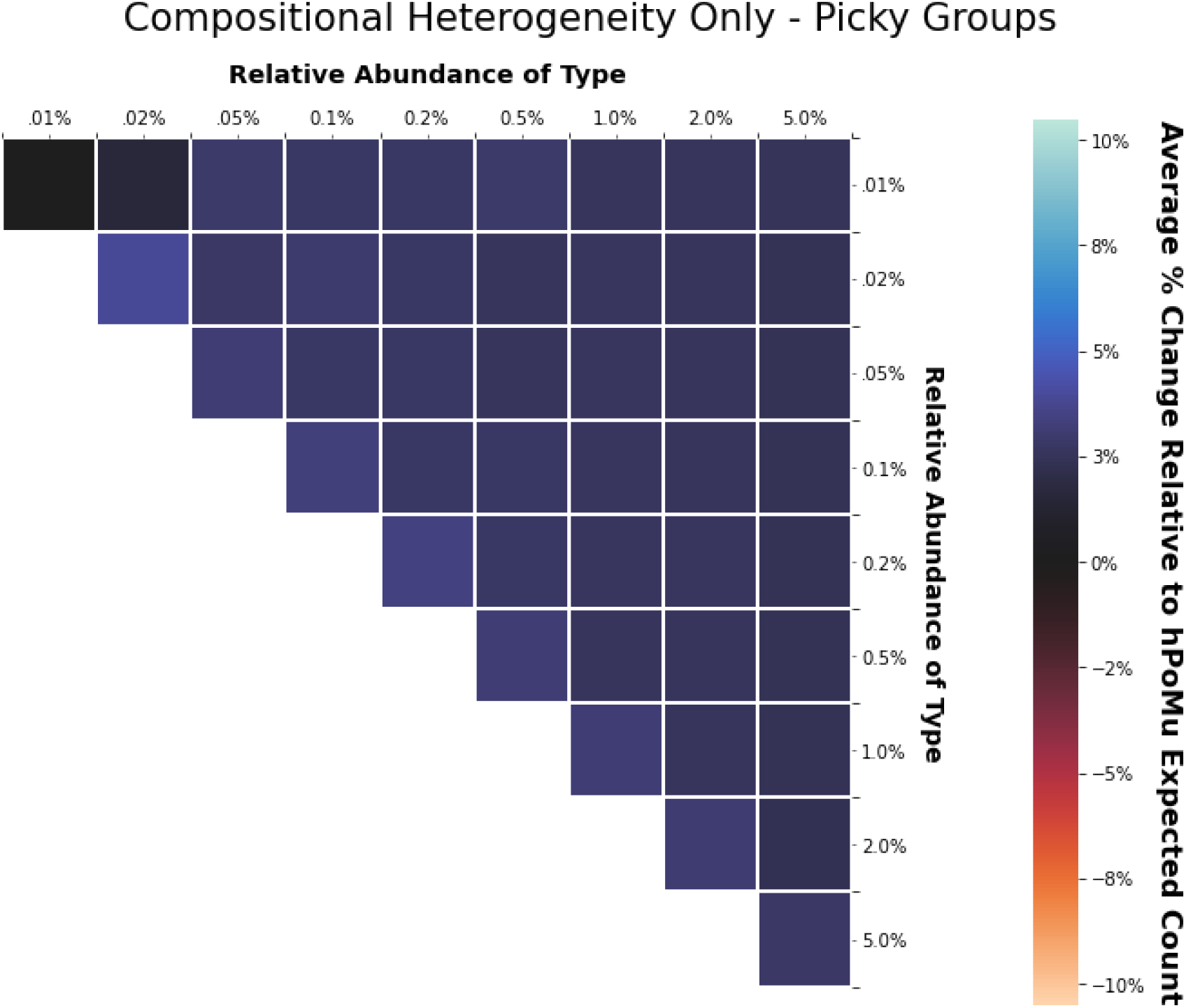
Cf. Supplementary Figure S49.

#### Compositional and Density Heterogeneity

In the presence of density heterogeneity, compositional heterogeneity still increases the number of droplets with a single type, as visible in figure 7. However, in this case compositional heterogeneity also decreases the number of droplets with two types. This is visible in figure 7, but more clearly evident from the decrease in the number of picky treatments visible in figure 11.

**Figure 11.**
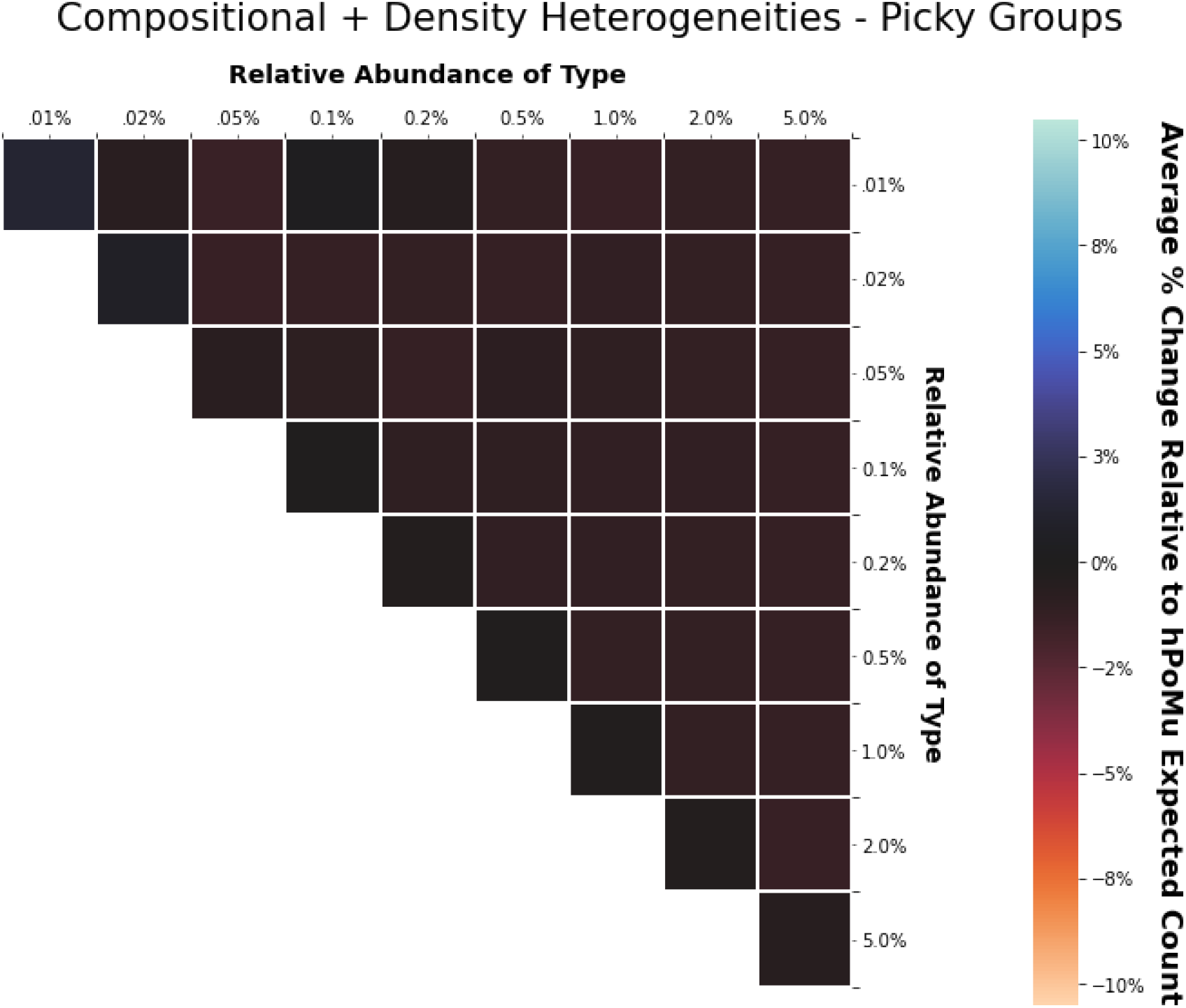
Cf. Supplementary Figure S51.

Nevertheless, it is also clear from figure 7, and from a comparison of figures 11 and 9, that the decrease in the number of droplets with two types is not as pronounced (at least in relative terms) as the decrease in the number of droplets with three or more types. Also in contrast to what happens in the absence of density heterogeneity, in the presence of density heterogeneity compositional heterogeneity markedly increases the number of empty droplets, as shown in figures 6 and 7.

### Estimators Behave Well

The estimators satisfy the two minimal requirements for the behavior of a “decent” estimator. Both their accuracy and precision increase with the size of the batches. The distribution of the estimated concentrations is centered around the true value. Moreover, the variance of the distribution of estimated concentrations decreases as the batch size increases. Cf. figures 12 and 13.

**Figure 12.**
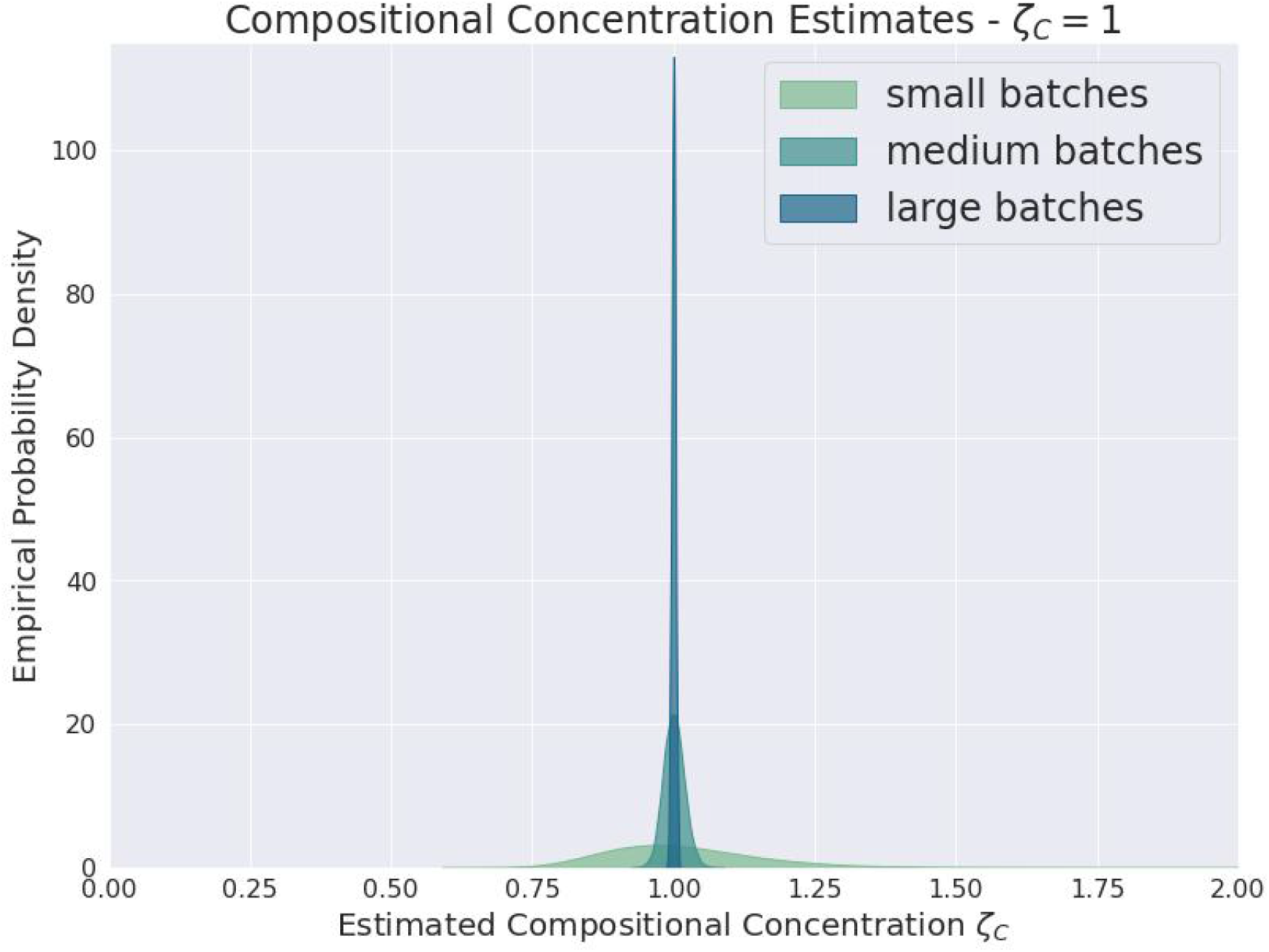
Cf. Supplementary Figure S89.

**Figure 13.**
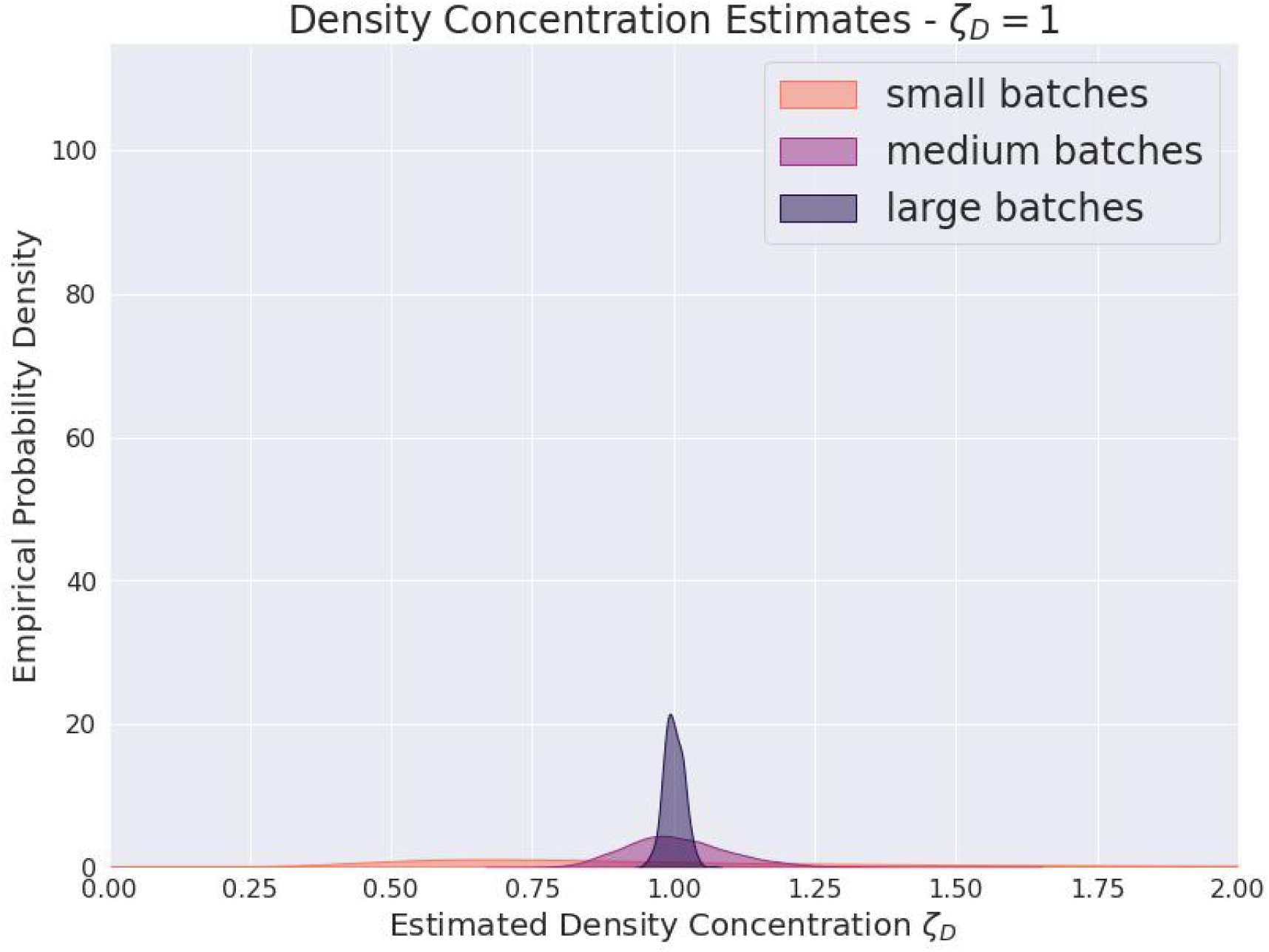
Cf. Supplementary Figure S87.

## DISCUSSION

### The Effect of Heterogeneity on Data Throughput is Equivocal

The general effect that heterogeneities will have on the data throughput of multi-cell, multi-type droplet microfluidics experiments is not straightforward. The answer depends on the particular combination of density heterogeneity and compositional heterogeneity present.

Given both density and compositional heterogeneity, it is clear that the effect on data throughput is negative. Both the numbers of picky treatments and gluttonous treatments decrease. Regardless of whether one uses picky or gluttonous controls, the number of treatments is still the limiting “bottleneck”.

Given only compositional heterogeneity but no density heterogeneity, whether the effect on data throughput is positive depends primarily on our opinion of what is useful data. If our primary concern is avoiding “data starvation” and being able to say anything about the interactions of the rarest types, then the strong decrease in the number of gluttonous treatment groups clearly means that the effect of heterogeneity is negative. On the other hand, if we consider only the data from the picky treatments useful, then in the absence of density heterogeneity the effect of compositional heterogeneity is slightly positive.

### The Estimators are Good Enough for Use in Practice

The above results show that the proposed estimators both (1) converge towards correct answers as the data size increases and (2) have decreasing variance as the data size increases. Therefore they satisfy the minimal requirements one generally expects of “decent” or “adequate” statistical estimators.

The estimators were already guaranteed to be asymptotically consistent when the statistical model was correctly specified. Nevertheless, that theoretical guarantee does not necessarily translate in practice into estimates usefully approximating the truth for realistic data sizes (represented by the medium or large batches). Consistency of an estimator need not imply efficiency of an estimator. The above results show that the proposed estimators are in fact useful in practice. Not only that, but they often manage to give useful results even for unrealistically small data sizes (represented by the small batches).

## CONCLUSIONS

We need to be able to estimate both density and compositional heterogeneity from empirical data produced by multi-cell, multi-type droplet microfluidics experiments in order to predict how these heterogeneities might affect the throughput of a given experimental platform. We have viable statistical estimators for both density heterogeneity and compositional heterogeneity. Therefore, neither quantity needs to be treated as an “unfathomable unknown”. We are able to actively account for the effects of both. Given empirical data, even of only moderate size, we can get estimates for concentration parameters describing levels of heterogeneity that are realistic for the experimental platform.

Having some sense for the amount of either type of heterogeneity one is likely to encounter in practice for any given experimental platform is important. It enables more accurately predicting the throughput from multi-cell, multi-type droplet microfluidics experiments. More accurate predictions of throughput will make droplet microfluidics experiments more viable alternatives to manually co-culturing multiple types of cells. More accurate predictions of throughput are also necessary to accurately simulate multi-cell, multi-type droplet microfluidics experiments. Accurate simulations in turn will help us to identify the statistical analyses that perform best for the novel kinds of datasets produced by multi-cell, multi-type droplet microfluidics experiments. Identifying best performing statistical analyses for these experiments will further make them more viable alternatives to manually co-culturing multiple types of cells.

## Supporting information

Supplementary Sections

## ACKNOWLEDGMENTS

Professor Adam P. Arkin for critical suggestions, essential scientific and technical advice, important introductions, and access to funding and computational resources without which this project would have been impossible. Dr. Fangchao Song for suggesting the topic of the dissertation (Krinsman, 2022) and generous consultations, feedback, and advice throughout. Professor Mark van der Laan for critical suggestions, essential scientific and technical advice, and invaluable recommendations of strategies and resources for scientific writing. Dr. Lauren M. Lui for invaluable recommendations of strategies and resources for scientific writing. Professor Perry de Valpine for the useful suggestion of investigating the Dirichlet-Multinomial and related models for weakening the ‘well-mixed’ droplets assumption.

## Footnotes

1 Cf. sections S3.2.2 and S4.2.2.

2 In practice, given realistic problem sizes, evaluating this exact formula would most likely be intractable, such that one would probably also need to use additional heuristics and approximations. The choice of those heuristics and approximations would depend on the particular causal estimand and estimator thereof, and thus is outside the scope of this discussion.

3 The goal for these experiments is to have one cell per droplet, whence the name “single-cell”. However, because the droplet formation process is random, in practice some droplets in “single-cell” experiments may still end up with more than one cell.

4 Via SciPy (Virtanen et al., 2020), which chooses bandwidths according to Scott’s Rule (Scott, 1992).

