## Supplementary Sections for "Models of Throughput for Multi-Cell, Multi-Type Droplet Microfluidics"

**William Krinsman**[1]

[1]Graduate Group in Biostatistics; University of California, Berkeley
[1]Division of Computing, Data Science, and Society; University of California, Berkeley
[1]Center for Computational Biology; University of California, Berkeley
[1]BioSciences Area; Lawrence Berkeley National Laboratory

Section S1 intends to explore how droplet-based microfluidics technology can be used to infer microbial interactions, in particular why this approach may overcome previous limitations. It does so by providing an overview of the particular experiment motivating (Krinsman, 2022). Section S2 defines and discusses the candidate working statistical models. Section S3 qualitatively compares the candidate working statistical models. We want to determine whether any behave substantially differently from the default working statistical model, which is the simplest. Motivated by the results of section S3, section S4 investigates and compares the target estimands (which quantify the data throughput) for each of the candidate working statistical models. We want to determine whether, in practice, substantially different behavior between the candidate working models translates into substantially different values of the target estimands. Section S5 constructs estimators and obtains inference for all of the candidate working statistical models. Even under model misspecification, these estimators return sensible results.

#### Supplementary Sections

**S1 Using MOREI to Characterize Microbial Interactions** **3**

**Additional References** **141**

### S1 USING MOREI TO CHARACTERIZE MICROBIAL INTERACTIONS

MOREI (pronounced "more-ray") stands for "More Interactions" or (treating MORE itself as a backronym) "Microfluidics Offers Replicate Experiments for Interactions". Starting from a microbial community sample, MOREI creates on the order of $10^5 - 10^6$ microfluidic droplets for characterizing microbial interactions. The droplets are created using droplet-based microfluidics chips.

Several batches of droplets are produced. Droplets within each batch are incubated for the same amount of time, while different batches are incubated for different amounts of time. Cf. figure S1.

The "life cycle" of each droplet can be divided into roughly three phases.

1. The first phase is "**initial droplet formation**". In the first phase, fluid from the microbial community sample is pumped from a syringe into the microfluidics chip. The microfluidics chip uses oil to encapsulate the fluid, as well as any cells that may have been inside the fluid, into a droplet.

2. The second phase is "**growth of cells inside of the droplet**". In the second phase, the droplet is incubated for the amount of time prescribed for its batch, during which any cells inside are able to divide and grow.

3. Finally the third phase is "**preparation for sequencing**". In the third phase, the droplet is fed into another microfluidics chip, where it is merged with another droplet containing materials that enable the sequencing and unique identification of the contents of the original droplet.

After all three phases are completed, all of the (merged) droplets are pooled together to undergo PCR[5] and finally sequencing. Cf. figure S2 below. I will also describe all three phases in more detail below.

#### S1.1 Initial Droplet Formation

The microfluidics chip encapsulates cells into droplets randomly. This makes each droplet's initial condition, both the number of cells in the droplet, as well as which the strains the cells belong to, random. The growth and division of cells during the second phase means that the droplets' random initial conditions are obscured in the final data. Therefore the best we can do is to make predictions of the droplets' unobserved random initial conditions based on average behavior.

We can control the concentration/dilution of the microbial community sample to guarantee a specified average number of cells per droplet. The average number of cells per droplet is usually chosen to be around two. This is intended to guarantee that, when there are two (or more) strains present in a droplet, the most likely scenario is that there is at most one cell of each strain.

The small average number of cells per droplet has other effects. The data is incredibly sparse. Even in a typical non-empty droplet, almost all strains will be absent. This is very much unlike, for example, typical single-cell RNAseq data. Another effect is that for any given interaction, the number of droplets that serve as controls will usually vastly exceed the number of droplets that serve as treatments. Therefore the data throughput bottleneck for inferring interactions becomes the number of treatments.

[5]polymerase chain reaction

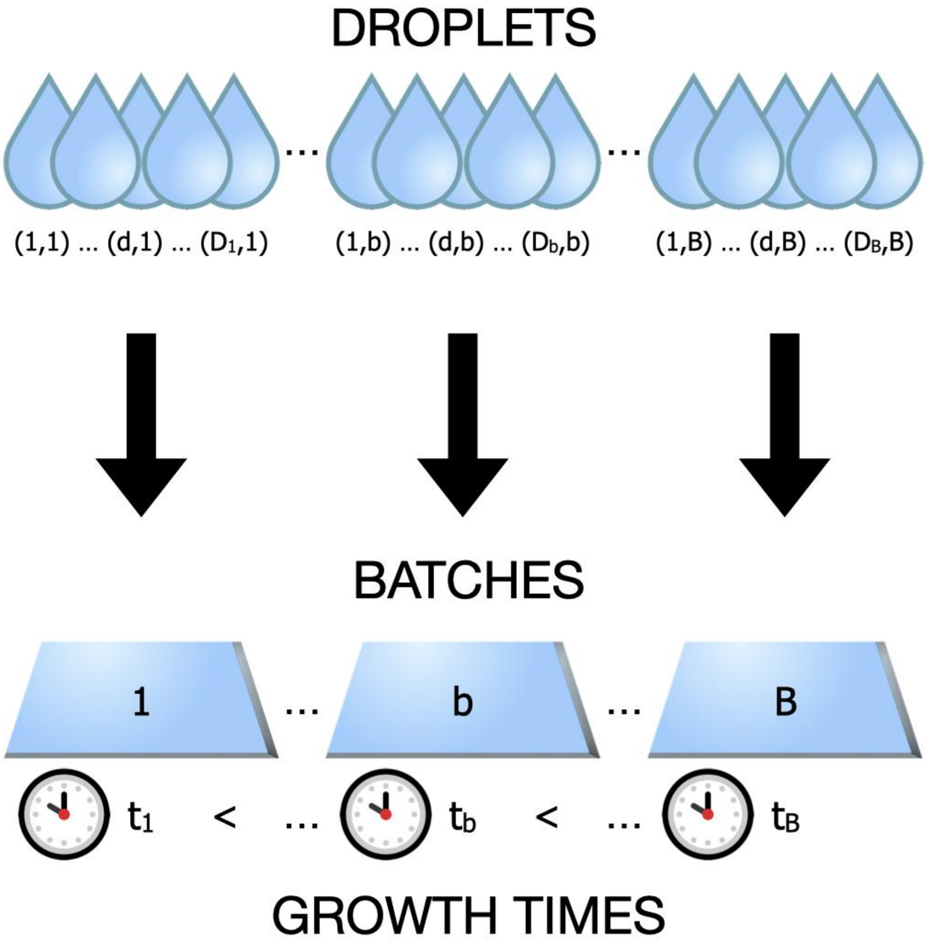

**Supplementary Figure S1.** Notation; Organization of Droplets into Batches

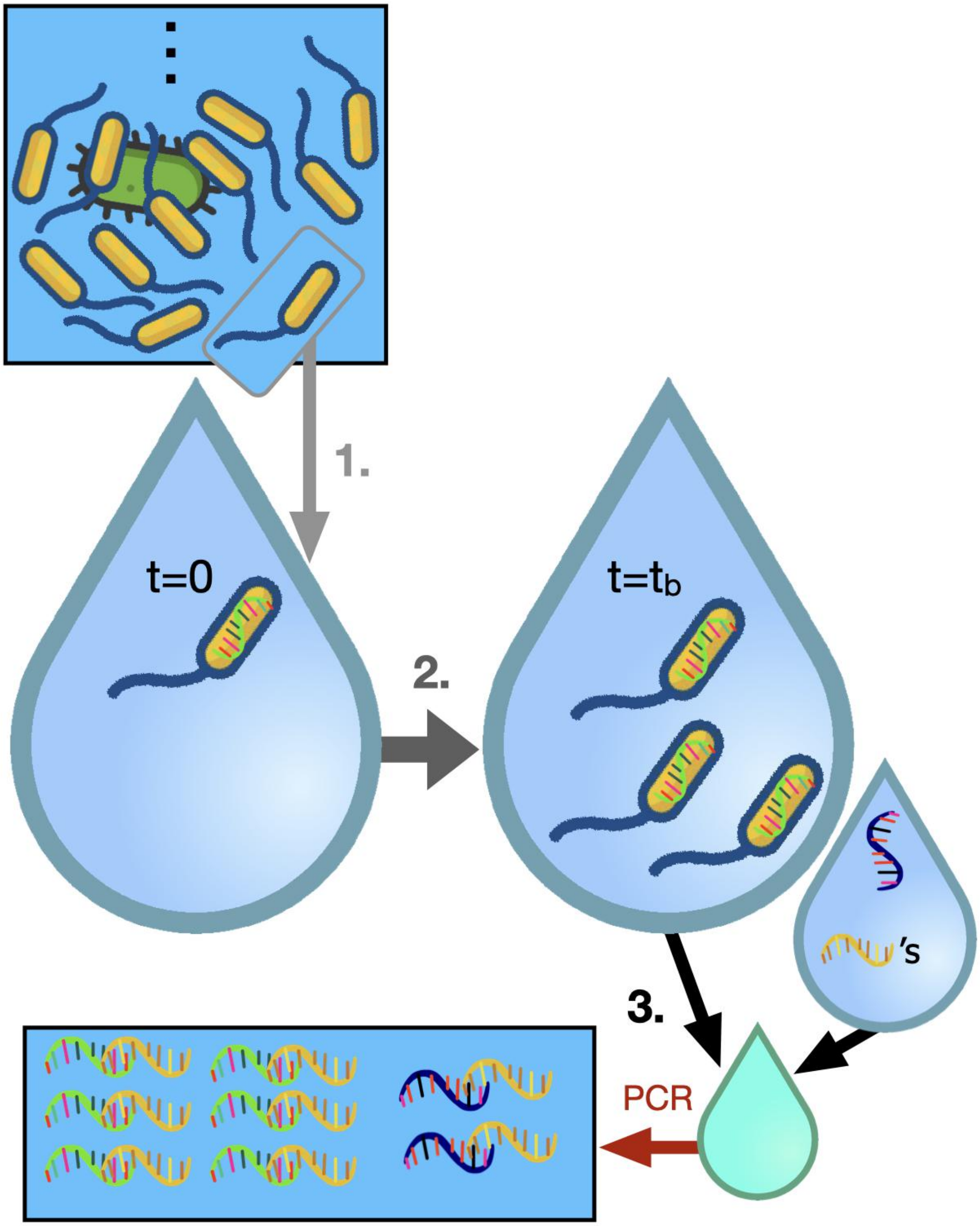

**Supplementary Figure S2.** Schematic of the "life cycle" of a droplet. See text for details.

#### S1.2 Growth of Cells Inside of the Droplet

The contents of the droplets are only observed once, after the end of the second phase. In particular, the details of what occurs during the second phase inside of any given droplet are completely opaque to us. The resulting data is therefore not longitudinal. Thus many standard methods for inferring microbial interactions, which assume time series data, are not applicable. Currently there appears to be no scalable method that can measure the abundances of microbial strains within the same microfluidic droplet at multiple time points (Dressler et al., 2017).

The droplets are all small, and therefore obviously quite nutrient-limited. Therefore models of microbial interactions like the generalized Lotka-Volterra equations, cannot make accurate predictions for the second phase, because they allow for indefinite growth. In reality, the maximum possible number of cells that could exist inside of a droplet at the end of the second phase is modest.

How long the droplets are incubated will determine whether cells are still actively dividing at the end of the second phase. It could be more difficult to infer the strengths of interactions from cells that exhausted the nutrients in their droplet and are undergoing logistic-like growth. This makes it important to incubate distinct batches for distinct amounts of time, to get "pseudo-longitudinal" data. Future work needs to determine how to best combine data from different batches, for example how to best balance averaging results within batches and averaging results between batches, while also accounting for how batches that were incubated longer are more likely to correspond to logistic-like growth.

#### S1.3 Preparation for Sequencing

During the third phase, a second microfluidics chip merges each droplet with a "PCR mix" droplet. The contents of the "PCR mix" droplet prepare the contents of the original droplet for sequencing.

First there is a lysis buffer, which lyses the cells in the droplet. This kills any cells which are still alive, and makes the genetic material inside all cells (living or dead) accessible. The third phase is necessary for observing the contents of the droplet, but the lysis buffer also ensures that it is destructive and cannot be repeated. Therefore the third phase is what forces the data to not be longitudinal. Because dead cells are also lysed and have their genetic material available for sequencing, this also introduces potential biases from counting such "relic" genetic material the same as if it had come from living cells. Cf. figure S3.

Second there are the barcodes, represented by the yellow nucleotides in figure S2. More precisely, they consist of (1) an Illumina Miseq adapter, (2) a barcode sequence, and (3) a 515F primer. The adapter ensures that the sequence and any other nucleotides that bind to it will be sequenced. The barcode sequences are (approximately[6]) unique for each "PCR mix" droplet. Therefore the barcode sequences identify everything they bind to as having come from the same droplet. Finally the 515F primers bind to 16S rRNA genes from the lysed microbial cells. This ensures that primarily the 16S rRNA genes from the cells will be amplified in the PCR step that occurs following the third phase. We use the 16S rRNA gene sequences to distinguish different strains.

Third there are spikein genes, represented by the blue nucleotides in figure S2. A random, but approximately constant[7], number of these are incorporated into each "PCR mix" droplet. Like the 16S genes from inside of the lysed cells, these will also bind to the barcodes, and therefore will also be amplified in the PCR step that occurs following the third phase. Assuming that all sequences from the same droplet that bind to the barcodes will be amplified by roughly the same amount during the PCR step, for each droplet we can divide all of the PCR amplified counts by the ratio of the PCR amplified spikein count to the average number of spikein gene copies originally in each "PCR mix" droplet.

While details of the spikein genes may seem unimportant from a purely statistical perspective, they actually imply a very important feature of the data. Specifically, the spikein gene protocol enables us to normalize by an (approximate) PCR amplification factor, which means that we get **absolute abundance**

[6]Choosing a barcode sequence of $L$ nucleotides, there are $4^L$ possible barcode sequences of length $L$. Therefore, choosing a long enough length $L$ for the barcode sequences, and generating them randomly, in principle one will have with high probability that the barcode sequence in each "PCR mix" droplet is unique, or that almost all are. This is much easier said than done, of course, but still feasible with today's technology.

[7] Assuming a roughly Poisson distribution(Collins et al., 2015) (for which the mean equals the variance), the larger the mean, the smaller the ratio of the standard deviation to the mean. For example, $\frac{\pm 10}{100}$ is a better relative uncertainty than $\frac{\pm 5}{25}$, even though $\pm 10$ is clearly a worse absolute uncertainty than $\pm 5$. Relative uncertainty is all that matters in this context because we use the PCR amplified count of spikein genes as a normalization factor. Thus, as long as the average number of copies of spikein gene per "PCR mix" droplet is large enough, we may treat it as "approximately constant". However, we don't want to add too many copies of the spikein gene on average, because then after PCR amplification the spikein genes will likely "swamp out" any signal from strains that are (relatively) less abundant inside of the droplet.

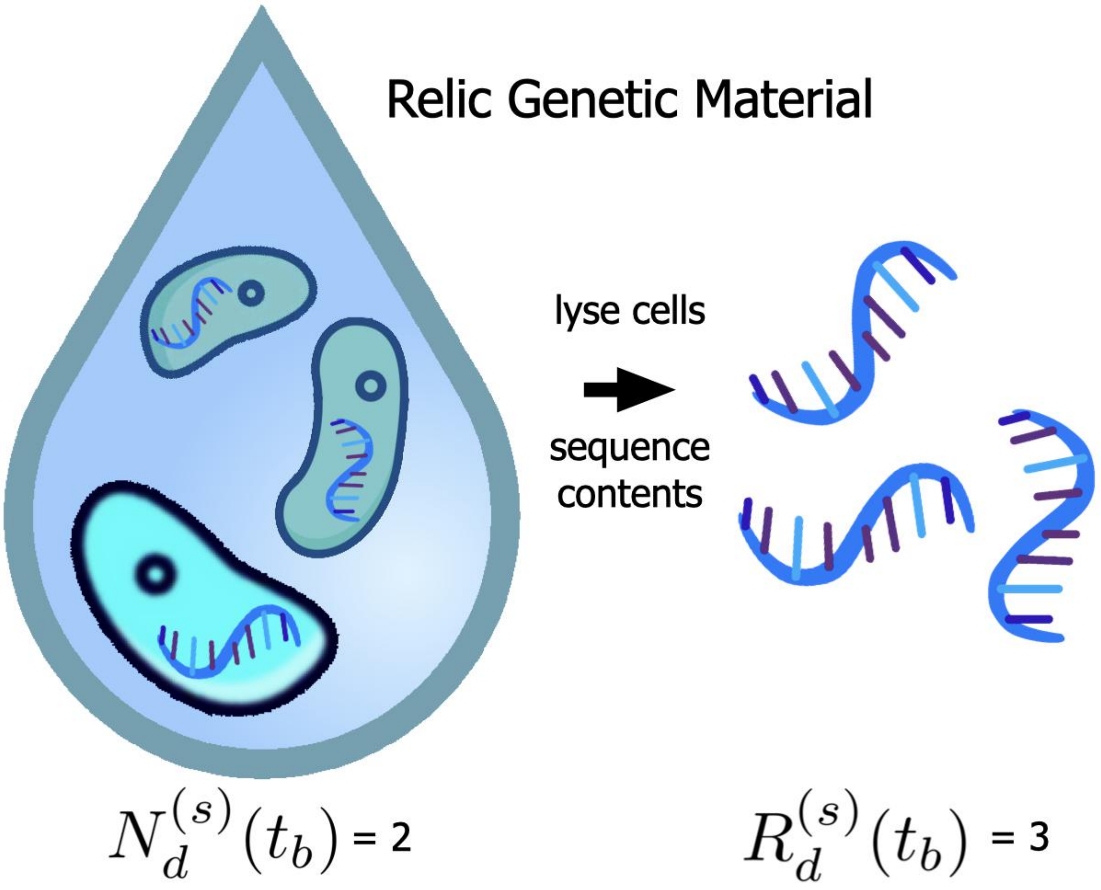

**Supplementary Figure S3.** Relic genetic material can inflate inferred count of cells. This droplet had two living cells and one dead cell after the end of the incubation period for its batch. However, all three cells (living and dead) are lysed and thus also have their contents sequenced.

values from the sequencing results. Without estimates for the PCR amplification factor, normally one can only infer *relative abundance* values. However relative abundance is problematic for inferring microbial interactions (Mounier et al., 2008). A previous attempt (Hsu et al., 2019) to study microbial interactions using microfluidic droplets was limited to 3 distinct strains, due to relying on fluorescent labeling of cells to get absolute abundance measurements. In contrast MOREI is in principle capable of studying microbial communities with many more strains, because strains are distinguished using 16S rRNA gene sequences, while still producing absolute abundance measurements.

##### S1.4 Conclusion

MOREI needs to increase the number of strains whose interactions can be simultaneously characterized in order to advance the state of the art in microbial ecology. Being able to characterize the interactions of more strains corresponds to being able to characterize the interactions of strains with low relative abundance (a.k.a. frequency) in the microbial community sample. For example, if all strains have the same frequency, then simultaneously characterizing the interactions of 20 strains requires characterizing the interactions of strains with 5% relative abundance, while simultaneously characterizing the interactions of $2,000$ strains requires characterizing the interactions of strains with .05% relative abundance.

Thus understanding how to use MOREI to advance the state of the art in microbial ecology requires understanding how to analyze the data produced by MOREI in a way that characterizes the interactions of the least abundant strains as accurately as possible.

Given highly accurate simulations of the data-generating process for MOREI, we could compare the performance of different estimators to see which most accurately characterizes the interactions of the least abundant strains. This would determine the methods that are best for analyzing the data produced by MOREI. That in turn would determine how far we can realistically expect to advance the state of the art in microbial ecology using MOREI. The performance of any statistical method for inferring a microbial interaction, however, will always be limited by the amount of data it receives as input. This corresponds to the throughput, which in turn is determined by the first phase of MOREI. This makes accurate predictions and modeling of the first phase of MOREI particularly important, hence why it is the focus of this paper.

All three phases of the droplet life cycle described in section S1 present potential difficulties for data analysis. Hence we need useful statistical models for all three phases to design accurate simulations of the MOREI data-generating process. Section S2 discusses statistical models one could use for the first phase. Section S3 builds on this by comparing the different models. Section S4 investigates the data throughput predicted by each of the models. Section S5 establishes how to identify which of the models is most realistic. Detailed investigations of statistical issues arising from the second and third phases (as well as the PCR amplification step) unfortunately are deferred to future work.

#### S2 MODELING INITIAL FORMATION OF DROPLETS

Section S2.1 discusses how a statistical model of the initial formation of droplets helps us to characterize microbial interactions. Section S2.2 proposes a general framework for defining any statistical model of the initial formation of droplets. Section S2.3 proposes a sensible default for the statistical model of the initial formation of droplets. In section S2.4 I explain three ways in which the assumptions underlying this default model could fail to be true in practice.

The rest of section explains how to model these assumption failures. Specifically, section S2.5 proposes a model accounting for the finiteness of the sampled pool of microbes. Section S2.6 proposes a model accounting for heterogeneity in the relative abundances of microbial strains throughout the sampled pool of microbes. Section S2.7 proposes a model accounting for heterogeneity in the average number of cells throughout the sampled pool of microbes. Section S2.8 describes a family of distributions subsuming both of the previous two models as well as the default model.

Section S2.9 discusses how this section connects with the others.

##### S2.1 Statistical Motivation

One of the major challenges of the MOREI data is the randomness of the initial droplet formation process. Unlike manually plating everything, with MOREI the scientist has no *direct* control over which microbial interactions are observed. This means a loss of control over which questions that can be answered from the data. The premise of MOREI is that the vastly increased throughput makes the tradeoff of decreased control worthwhile. The more control we can retain for the scientist when using MOREI, compared to

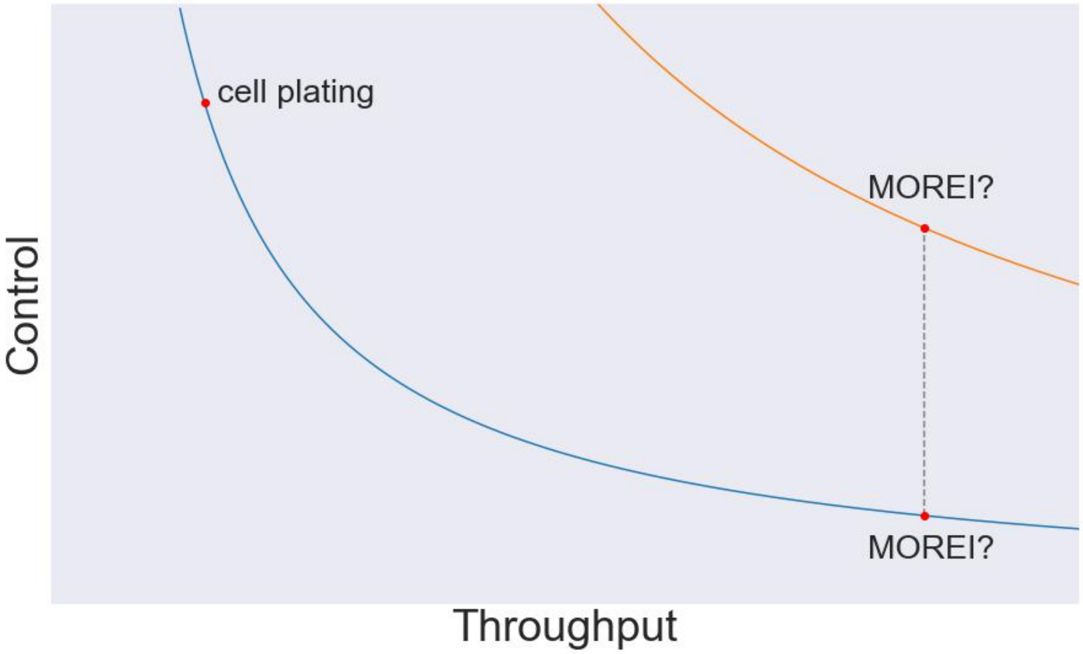

**Supplementary Figure S4.** Control-Throughput Tradeoff Curve. Having a predictive statistical model of initial droplet formation would allow us to increase control without sacrificing throughput, moving us to the higher tradeoff curve.

direct plating, the stronger the case for its superiority over existing methods, and the greater its ability to advance the state of the art. Cf. figure S4.

To restore the scientist's control over which questions they can answer using MOREI data, we want to account for the effects which the randomness of the initial droplet formation process has on the data. To account for these effects, we need to make accurate predictions about what these effects will be. To make such predictions, we need to understand the randomness of the initial droplet formation process. To understand its randomness, we need to have adequate statistical models for describing the initial droplet formation process.

Thus the goal of this section is to review how we might model the initial droplet formation process. I do so both in general terms, as well as derive specific distributions following from specific assumptions we might make about the initial droplet formation process. This is our first step to understanding the randomness of the droplet formation process, to making predictions about its effects, to accounting for its effects, and finally to restoring control to the scientist over which questions they can answer using MOREI data.

While this ultimate goal may sound overly ambitious, we need to keep in mind that the amount of throughput that we can potentially achieve with a MOREI experiment is enormous. With such enormous sample sizes, the effects of asymptotic concentration of measure phenomena (such as the law of large numbers or the central limit theorem) almost certainly come into play. To phrase it very crudely, "randomness + enormous sample sizes = quasi-determinism". To predict where exactly measure will concentrate, and thereby exploit the "quasi-determinism" resulting from enormous sample sizes, all we need is to understand and characterize the underlying probability distribution well enough.

#### S2.2 Hierarchical Count-Categorical Distributions

I define a **count distribution** to be a probability distribution $\mathcal{P}$ taking values in the non-negative integers $\mathbb{N}_{\geq 0}$. For example, $N(0)$ corresponds to a count distribution, as does $N^{(s)}(0)$ for all $s \in [S]$.

Similarly, I define a **multivariate count distribution** to be a probability distribution taking values in $\mathbb{N}_{\geq 0}^S := \bigtimes_{s=1}^S \mathbb{N}_{\geq 0}$. For example, $\vec{\mathbf{N}}(0)$ corresponds to a multivariate count distribution. All of the marginal distributions of any multivariate count distribution are themselves count distributions.

A typical way to specify a multivariate count distribution is to specify the $S$ count distributions that are its marginal distributions along with the statistical dependence structure amongst the marginals. There are infinitely many ways to define such a statistical dependence structure. Arguably the simplest of such ways is to declare the marginal distributions to be mutually independent, but making such a restriction drastically reduces the variety of multivariate count distributions that can be described. In general, directly specifying the statistical dependence structure amongst the marginal distributions can be quite complicated and difficult. Specifying $S$ count distributions can also be cumbersome.

In this thesis I will use another framework for specifying multivariate count distributions. This framework decomposes the multivariate count distribution into a hierarchical[8] distribution. One of the distributions in the hierarchical distribution is always easy to describe. If the other distribution (more properly/technically speaking a family of distributions) in the hierarchical distribution is also easy to describe, then this hierarchical framework allows us to describe a multivariate count distribution easily using two distributions.

Consider a random vector $\vec{\mathbf{X}} \in \mathbb{N}_{\geq 0}^S$ corresponding to a multivariate count distribution. As mentioned before, this always corresponds to $S$ count distributions, one for each of its $S$ marginal distributions $X^{(s)}$. However, there is also always a way to associate with $\vec{\mathbf{X}}$ a single count distribution, namely the sum of the marginal distributions $X := \sum_{s \in [S]} X^{(s)}$. To define a hierarchical distribution from this, we need to specify a conditional distribution for every possible value of $X$. In other words, instead of considering $X^{(s)}$ for every $s \in [S]$ to specify $\vec{\mathbf{X}}$, I propose instead considering the pair $X$ and $\vec{\mathbf{X}}|X$.

For any $n \in \mathbb{N}_{\geq 0}$, $\vec{\mathbf{X}}|X = n$ takes values in $\{0, \ldots, n\}^S$ with the restriction that the sum of its marginals must equal $n$. I define a **categorical distribution** to be any [9] probability distribution $\mathcal{P}_n$ that takes values in $\{0, \ldots, n\}^S$ with the restriction that the sum of its marginals must equal $n$, or which equivalently takes as values partitions of $n$ into $S$ (possibly empty) distinct categories.

Two observations are worth making here. First, technically speaking this framework can be used to characterize any multivariate count distribution. No restrictions[10] on $\vec{\mathbf{X}}$ are required to define $X$ and $\vec{\mathbf{X}}|X$. Second is that $\vec{\mathbf{X}}|X$ is not a proper random variable. At best $\vec{\mathbf{X}}|X$ is a family of random variables or perhaps more accurately a "random random variable". For every $n \in \mathbb{N}_{\geq 0}$, $\vec{\mathbf{X}}|X = n$ is a distinct (proper) random variable with its own distinct probability distribution (a categorical distribution). Thus this framework can only provide a simplification in the case that the infinite family of probability distributions corresponding to the infinite family of random variables $\vec{\mathbf{X}}|X = n$ for all $n \in \mathbb{N}_{\geq 0}$ can itself be described simply (ideally as a straightforward function of $n$). Otherwise this framework clearly complicates[11] the description of $\vec{\mathbf{X}}$.

Given a single count distribution $\mathcal{P}$ and a family of categorical distributions $\mathcal{P}_n$ for all $n \in \mathbb{N}_{\geq 0}$, we can construct a random vector $\vec{\mathbf{X}}$ following what I call a **hierarchical count-categorical distribution** as follows: given a random variable $X \sim \mathcal{P}$, draw from the categorical distribution $\mathcal{P}_n$ whenever the value of $X$ is $n$. Observe how, by construction, the sum of the marginals of $\vec{\mathbf{X}}$ is $X$, and that $(\vec{\mathbf{X}}|X = n) \sim \mathcal{P}_n$ for all $n \in \mathbb{N}_{\geq 0}$. Again, any multivariate count distribution can be constructed this way, but whether this is a simple description of the multivariate count distribution depends entirely on whether the infinite family of categorical distributions $\mathcal{P}_n$ for all $n \in \mathbb{N}_{\geq 0}$ can be simply described.

In what follows, I indulge in an abuse of terminology by referring to the infinite family of categorical distributions corresponding to the $\vec{\mathbf{X}}|X = n$ for all $n \in \mathbb{N}_{\geq 0}$ as "the categorical distribution". This is to emphasize the henceforth implicit assumption that this infinite family of categorical distributions admits a simple parameterization in terms of $n$, simple enough to *not* require much (or any) more effort to describe

[8] In some contexts such distributions are also called "compound". I chose to use "hierarchical" here instead to avoid confusion with another use of the word "compound" that often occurs with Poisson distributions, i.e. a "compound Poisson distribution" in the sense of a sum of $N$ i.i.d. random variables where $N$ itself is Poisson distributed. For example, using the latter notion of "compound", hPoMu is a "compound Poisson Multinoulli", but using the notion of "compound = hierarchical", hPoMu is "compound Poisson Multinomial".

[9] This definition is much more general than what is more commonly referred to as "*the* categorical distribution" (sensu stricto), for which $n = 1$, and which is also called a Multinoulli distribution. Actually all categorical distributions (sensu lato) with $n = 1$ have to reduce to a Multinoulli distribution. This is analogous to how they must reduce to the Dirac Delta at 0 in the case where $n = 0$. Note also that while for $n > 1$ the sum of $n$ independent Multinoulli random variables does correspond to a categorical distribution (sensu lato), specifically the Multinomial distribution, not every categorical distribution (sensu lato) for $n > 1$ is Multinomial. So one could reasonably say that categorical distributions (sensu lato) strictly generalize categorical distributions (sensu stricto) only for $n > 1$.

[10] In fact, even the restriction that $\vec{\mathbf{X}}$ is a multivariate count distribution may be immaterial/unnecessary, although it does imply that $X$ is a discrete distribution, which in turn simplifies considerations of mathematical rigor.

[11] In the worst case this framework requires specifying one count distribution and infinitely many completely unrelated categorical distributions. Compare this to specifying $S$ count distributions and one description of their statistical dependence structure that is required by the other aforementioned framework for specifying a multivariate count distribution.

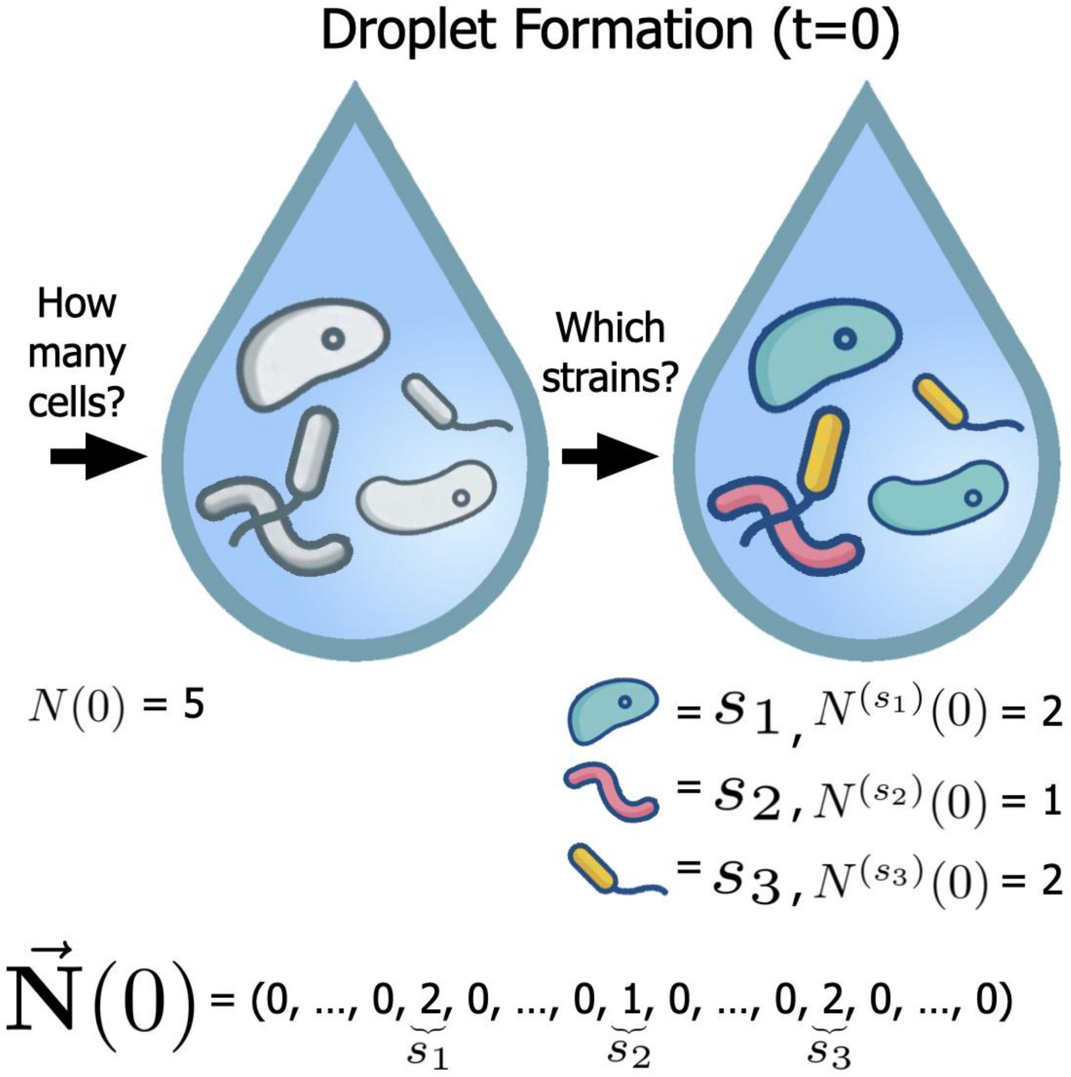

**Supplementary Figure S5.** Schematic of count-categorical distributions.

than a single categorical distribution. When[12] this assumption holds, describing $X$ and $\vec{\mathbf{X}}|X$ is not much more difficult than specifying a single count distribution and a single categorical distribution.

In the context of MOREI, the count distribution part of a hierarchical count-categorical distribution specifies how many cells $N(0)$ are in the droplet, while the categorical distribution part of a hierarchical count-categorical distribution specifies which proportions of cells $\vec{\mathbf{N}}(0)|N(0) = n$ belong to each of the $S$ possible strains. Therefore using the hierarchical count-categorical distribution framework allows us to split the problem of selecting a statistical model for initial droplet formation into two (smaller and easier) parts: (1) selecting which statistical model describes how many cells are in the droplet via the count distribution and (2) selecting which statistical model describes which strains any cells in the droplet belong to via the categorical distribution. Cf. figure S5.

Splitting the problem up this way not only simplifies it, but also makes it easier to connect the model with details of the experimental setup. For example, because we expect the dilution of cells in the microbial community sample to only affect the number of cells that end up in each droplet, we can connect it with a parameter for the count distribution (without affecting the categorical distribution). Similarly, because we expect the relative abundances of the strains in the microbial community sample to only affect

[12] This assumption may not be possible to satisfy for every multivariate count distribution. It is impossible to falsify either way without making the assumption more precise, of course.

the proportions of the cells in each droplet belonging to each strain, we can connect them to parameters for the categorical distribution (without affecting the count distribution).

###### *S2.2.1 Naming Convention*

Throughout, hierarchical count-categorical distributions will be named (or abbreviated) as follows.

- First, a prefix "h" for "hierarchical".
- Two letters corresponding to the count distribution.
- Two letters corresponding to the categorical distribution.

The two letters, for both the count distribution and the categorical distribution, are assigned as follows.

- If the name of the distribution is one word, then the two letters are the first two letters of the word, the first upper case and the second lower case.
- If the name of the distribution is more than one word, then the two letters are the first two initials of the name, both upper case.

For example the "hierarchical geometric multivariate Wallenius' noncentral hypergeometric distribution" would be "hGeMW".

###### *S2.2.2 Connection with Topic Modelling*

Imagine a document as a droplet, and the cells within the droplet as words. We can count the number of the words in the document the same way we can count the number of cells in a droplet. Similarly, the way cells can be assigned to strains, words within a document can be assigned to topics. Imagining the strains of the cells as topics of words, we can also count the number of words corresponding to each given topic. Cf. figure S6. Statistical questions related to modelling documents and the topics of words within them is already a well-studied field, given the name "topic modelling".

Given the above analogy of documents with droplets, words with cells, and topics with strains, a valuable opportunity for future work is to apply what is already known about topic modelling to modelling the initial formation of droplets in MOREI. See e.g. Theorem 1 of (Zhou, 2018) for one relevant example – the paper discusses both the hPoMu and hNBDM distributions (using different names). The analogy genuinely seems to mean that mathematical abstractions relevant for one problem can be relevant for the other problem.

Of course this comes with caveats. One obvious difference is that parameter values for these models which are realistic for topic modelling are unlikely to be realistic for modelling the initial formation of droplets in MOREI, and vice versa. For example, the mean number of words in a document should generally be much larger than the mean number of cells in each droplet. A second caveat is that models like those in (Zhou, 2018) allow different documents to follow different distributions, whereas we might not want different droplets to follow different distributions (although doing so might be useful for e.g. modelling batch effects).

##### S2.3 hPoMu: A Default Model

Section S2.3.1 gives the explicit definition of the hPoMu distribution, mostly for future reference. Section S2.3.2 gives two heuristic arguments which establish hPoMu as the "baseline" or "default" model for the initial formation of droplets against which other models are compared. Finally section S2.3.3 outlines some of the assumptions implicit in the derivation of the hPoMu distribution, hinting at some of its potential weaknesses for accurately modeling the initial formation of droplets in practice.

###### *S2.3.1 hPoMu Definition*

As suggested by the name, the "hierarchical Poisson Multinomial" (hPoMu) distribution is a hierarchical count-categorical distribution whose count distribution is Poisson and whose categorical distribution is multinomial. Therefore its PMF equals

$$\begin{aligned}\mathbb{P}\left(\vec{\mathbf{N}}_d(0)=\vec{\mathbf{n}}\right) &= \mathbb{P}(N_d(0)=n)\cdot\mathbb{P}\left(\vec{\mathbf{N}}_d(0)=\vec{\mathbf{n}} \,\middle|\, N_d(0)=n\right) \\ &= \frac{e^{-\lambda}\lambda^n}{n!}\cdot\binom{n}{n^{(1)}\cdots n^{(S)}}\prod_{s=1}^{S}(f^{(s)})^{n^{(s)}}.\end{aligned} \tag{S1}$$

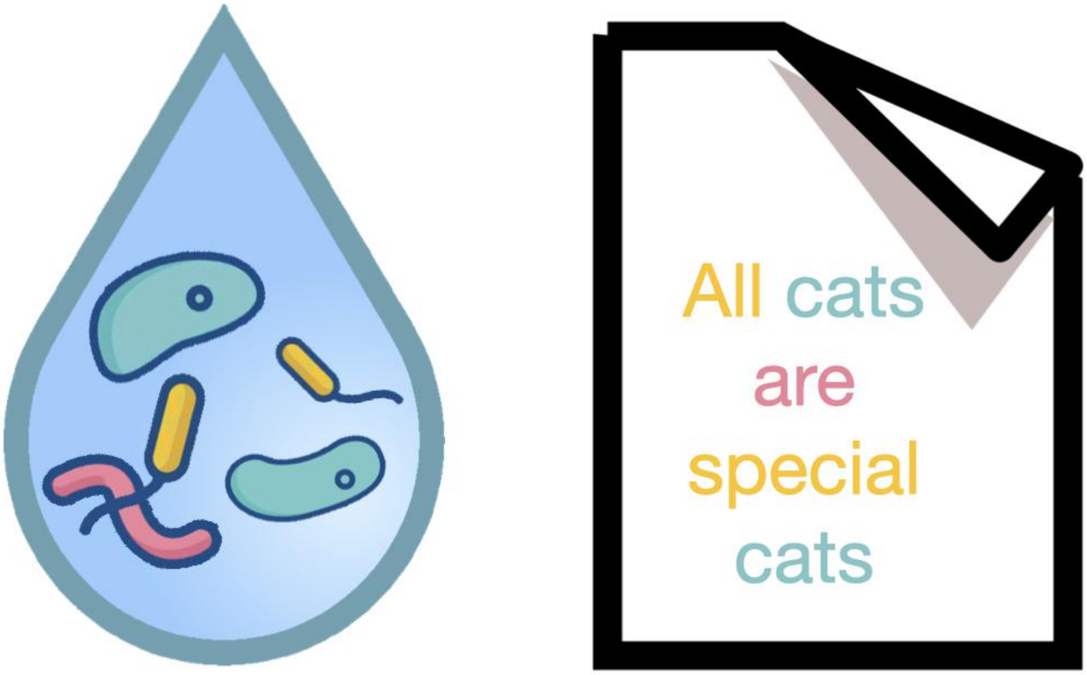

**Supplementary Figure S6.** Analogy between documents with droplets, words with cells, and topics with strains. The droplet has five cells, the document has five words. The droplet has two cells of the yellow strain, the document has two adjectives. The droplet has two cells of the blue strain, the document has two nouns. The droplet has one cell of the pink strain, the document has one verb. Similar families of probability distributions can be used to model both the strains and numbers of cells in droplets as well as the topics and numbers of words in documents.

An alternative form of expressing the PMF of hPoMu, which reveals that its marginals are mutually independent, is

$$\mathbb{P}\left(\vec{\mathbf{N}}_d(0) = \vec{\mathbf{n}}\right) = \prod_{s=1}^{S} \frac{e^{-f^{(s)}\lambda}(f^{(s)}\lambda)^{n^{(s)}}}{(n^{(s)})!} . \tag{S2}$$

Via tedious algebra, one can show that equation (S2) really does equal equation (S1). The $n!$ from the multinomial coefficient cancels with that from the Poisson PMF. The remaining factorials from the multinomial coefficient can be distributed across the product, and $e^{-\lambda}\lambda^n = \prod_{s=1}^{S} e^{-f^{(s)}\lambda}\lambda^{n^{(s)}}$, which can also be distributed across the product. So despite looking different, the first and second definitions are actually consistent with one another.

Heuristically speaking, equation (S2) says that during the formation of each droplet the sampling rate $\lambda$ is evenly spread out among the various strains according to their frequencies in the population.

###### *S2.3.2 Derivations*

The first derivation determines the count distribution and then the marginals. The second derivation determines the marginals and then the count distribution.

Assume we are sampling from a population of $K \in \mathbb{N}$ cells.

There are $f^{(1)}K \in \mathbb{N}$ cells of strain $1, \dots, f^{(s)}K \in \mathbb{N}$ cells of strain $s, \dots,$ and $f^{(S)}K \in \mathbb{N}$ cells of strain $S$. Then clearly $\sum_{\sigma=1}^{S} f^{(\sigma)} = 1$.

When forming a droplet, we assume each cell has the same small probability $p$ of ending up in the droplet. Thus the indicator random variable of ending up in the droplet is a Bernoulli$(p)$ random variable for every cell.

We also assume that whether a given cell ends up in the droplet is completely independent of what happens to any of the other cells.

**First Derivation** The number of cells that end up in the droplet is the sum of all $K$ of the indicator random variables. Because the sum of i.i.d. Bernoullis is binomial, the distribution of the number of cells that end up in the droplet is Binomial$(p, K)$.

Due to the way the experiment was calibrated, we know a priori that the expected number of cells in the droplet is (some fixed constant) $\lambda$. Because the number of cells follows a binomial distribution, the expected number of cells in the droplet is also $pK$. (Thus $p = \frac{\lambda}{K}$.)

Because $K$ is very large, we may as well use the approximating distribution as $K \to \infty$. By the "Law of Small Numbers", a.k.a. the Poisson Limit Theorem, this distribution is Poisson. In other words:

$$\mathbb{P}(N(0) = n) = \frac{e^{-\lambda}\lambda^n}{n!} . \tag{S3}$$

Now that we know the number of cells in each droplet, we need to determine which strain each cell belongs to. Because any cell was equally likely to have been sampled, the strains follow a multivariate hypergeometric distribution with categories of sizes $f^{(1)}K, \dots, f^{(S)}K$. Because the $K$ is very large, it is very unlikely[13] that any given cell would be sampled twice. Therefore we might as well use the approximating distribution as $K \to \infty$, multinomial with category probabilities $f^{(1)}, \dots, f^{(S)}$, which corresponds to sampling with replacement. Thus

$$\mathbb{P}\left(\vec{\mathbf{N}}(0) = \vec{\mathbf{n}} \,\middle|\, N(0) = n\right) = \binom{n}{n^{(1)} \cdots n^{(S)}} \prod_{s=1}^{S} (f^{(s)})^{n^{(s)}} . \tag{S4}$$

Combining the count distribution and categorical distribution derived above, it follows that the joint distribution must approximately equal

$$\begin{aligned}\mathbb{P}\left(\vec{\mathbf{N}}(0) = \vec{\mathbf{n}}\right) &= \mathbb{P}\left(\vec{\mathbf{N}}(0) = \vec{\mathbf{n}} \,\middle|\, N(0) = n\right) \cdot \mathbb{P}(N(0) = n) \\ &= \binom{n}{n^{(1)} \cdots n^{(S)}} \prod_{s=1}^{S} (f^{(s)})^{n^{(s)}} \cdot \frac{e^{-\lambda}\lambda^n}{n!} .\end{aligned} \tag{S5}$$

[13] Even more unlikely than being sampled once, which is already very unlikely.

**Second Derivation** For any given strain $s$, the number of cells *of that strain* that end up in the droplet is the sum of all $f^{(s)}K$ of the indicator random variables belonging to cells *of that strain*. Because the sum of i.i.d. Bernoullis is binomial, the marginal distribution of the number of cells of strain $s$ that end up in the droplet is again binomially distributed, Binomial$(p, f^{(s)}K)$.

Because[14] $p = \frac{\lambda}{K}$, the expected number of cells of strain $s$ that end up in the droplet is $p(f^{(s)}K) = \frac{\lambda}{K}(f^{(s)}K) = f^{(s)}\lambda$. If we assume that $K$ is sufficiently large such that $f^{(s)}K$ is very large, then we may as well use the approximating distribution as[15] $(f^{(s)}K) \to \infty$. As in the first derivation, this distribution is Poisson as a result of the Poisson limit theorem, so for the marginal distributions:

$$\mathbb{P}\left(N^{(s)}(0) = n^{(s)}\right) = \frac{e^{-f^{(s)}\lambda}(f^{(s)}\lambda)^{n^{(s)}}}{(n^{(s)})!}. \tag{S6}$$

Because what happens to any given cell is completely independent of what happens to any other cells, the marginal distributions for each strain are sums of mutually disjoint sets of i.i.d. Bernoulli random variables. Therefore the marginal distributions must be mutually independent Binomial random variables, and thus in the limit approach mutually independent Poisson random variables. It follows that the joint distribution must approximately equal

$$\mathbb{P}\left(\vec{\mathbf{N}}(0) = \vec{\mathbf{n}}\right) = \prod_{s=1}^{S} \frac{e^{-f^{(s)}\lambda}(f^{(s)}\lambda)^{n^{(s)}}}{(n^{(s)})!}. \tag{S7}$$

###### *S2.3.3 Implicit Model Assumptions*

Both of the above two derivations share the following implicit assumptions:

- The experiment can be calibrated to a priori guarantee a certain *average* number of cells per droplet.
- The total number of cells in the sampling pool, $K$, is very large.
- For *every* strain $s$, the number of cells $f^{(s)}K$ in the sampling pool which belong to strain $s$ is very large.
- Each cell has the same small probability (regardless of its strain) of ending up in the droplet as any other cell.
- (Regardless of their strains), whether a given cell ends up in the droplet is completely independent of what happens to any other cell.

The second and third can be thought of as stating that the population size is effectively infinite. The fourth and fifth can be thought of as stating that the sampling pool is completely homogeneous, or perfectly "well-mixed".

My collaborator tells me that in practice, using commercially available microfluidic devices, one can calibrate the experiment to a priori guarantee a certain average number of cells per droplet. My collaborator also tells me that in practice the total number of cells in the sampling pool will be $10^{10}$. However, the remaining three assumptions are more questionable. In the next section I explain three ways these assumptions could fail to be true in practice.

##### S2.4 Failures of hPoMu Model Assumptions

While hPoMu is definitely a sensible model to start with, its implicit assumptions failing to be realistic could undermine its usefulness in practice. In sections S2.4.1, S2.4.2, and S2.4.3 I define three plausible ways that these assumptions could fail to be true in practice. In section S2.4.4 I outline the framework which will be used to describe the last failures of hPoMu assumptions.

[14] Due to the argument found in the first derivation in section S2.3.2.

[15] Equivalently, as $K \to \infty$ along a subsequence such that $f^{(s)}K \in \mathbb{N}$ (i.e. remains an integer).

##### *S2.4.1 Sampling is from a Finite Population without Replacement*

In both of the above derivations from section S2.3.2, we assumed that the number of cells being sampled from is "effectively infinite". This implied that no meaningful loss of accuracy was occurred when invoking the Poisson limit theorem for the number of cells in each droplet. As mentioned before, with $\sim 10^{10}$ cells, this is a reasonable assumption.

However, we assumed more than this. We also assumed that *for each individual strain s* the number of cells is "effectively infinite". This assumption is reasonable and uncontroversial in the case of relatively abundant strains. However, in the case of "rare" strains, whose relative abundance is e.g. 0.1%, 0.01%, or even lower, whether this assumption remains reasonable is a priori unclear. Admittedly 0.01% of $\sim 10^{10}$ cells is still many, but perhaps not enough for the asymptotically approximate distributions to remain accurate.

##### *S2.4.2 Density Heterogeneity*

In both of the above derivations we assumed that each cell everywhere in the sampling pool had the same probability of being sampled for every droplet. Moreover, we also assumed that the probability of ending up in the droplet for any given cell was statistically independent of what happened to any other cell. This would make sense in a theoretical world where each cell corresponded to a scalar field uniformly spread across the entire sampling pool. Of course, in reality cells have finite volume and are limited at any given moment in time to a single location in the sampling pool. So only cells reasonably close to where the droplet is being formed have any nonzero probability of ending up in the droplet, and cells which are closer to one another are more likely to end up in the same droplet than those which are further apart.

These details hypothetically might not matter in the case where the average number of cells per unit volume[16] was constant throughout the entire sampling pool. However in practice we have no guarantee that this would be the case, and conceivably different sections of the sampling pool could have average numbers of cells per unit volume which differ from the average number of cells per unit volume for the entire sampling pool (i.e. the total number of cells divided by the volume of the sampling pool).

Herein I call this phenomenon "density heterogeneity". Compare figures S7 and S8. In those hypothetical examples, the average number of cells per unit volume for the entire sampling pool is 2, but the average number of cells per unit volume differs between the specific sections. In figure S7 the differences are small, introducing not much more variance compared to what would be expected under the uniform/homogeneous density assumed by the default hPoMu model. In figure S8 the differences are larger, introducing much more variance compared to what would be expected under uniform/homogeneous density assumed by the default hPoMu model.

In the hierarchical count-categorical distribution framework, density heterogeneity is an issue which affects the count distribution. The default Poisson model for the count distribution (found in hPoMu) will not account for additional variance which might be introduced in practice by density heterogeneity.

##### *S2.4.3 Compositional Heterogeneity*

In making the homogeneous assumptions that all cells have equal probability of ending up in the droplet, and that what happens to a given cell is completely independent of what happens to any other cells, we implicitly ignore the fact that cells belonging to different strains may tend to behave very differently.

In particular, although the relative abundance of strain $s$ across the entire sampling pool may be $f^{(s)}$, conceivably cells of this strain could be slightly more concentrated or slightly less concentrated than this in different sections of the sampling pool. This means different sections of the sampling pool could have average relative abundances which differ from the average relative abundances for the entire sampling pool. Thus the fact that each droplet is formed only from cells nearby a certain location in the sampling pool could affect not only the number of cells likely to end up in the droplet, but also which strains are most likely to end up in the droplet. This is probably guaranteed to happen to some extent due to random chance given that cells occupy finite, not infinitesimal volume. The probabilities of belonging to given strains cannot be perfectly uniform scalar fields. For an example of one mechanism that could aggravate this, cells of a certain strain could preferentially aggregate with, or avoid, cells of another strain due to their interactions with one another.

Herein I call this phenomenon "compositional heterogeneity". Compare figures S9 and S10. In those hypothetical examples, the average relative abundances of each strain for the entire sampling pool is $\frac{1}{2}$

[16]For volumes "large enough", i.e. this is obviously false for "infinitesimal volumes" as discussed already above.

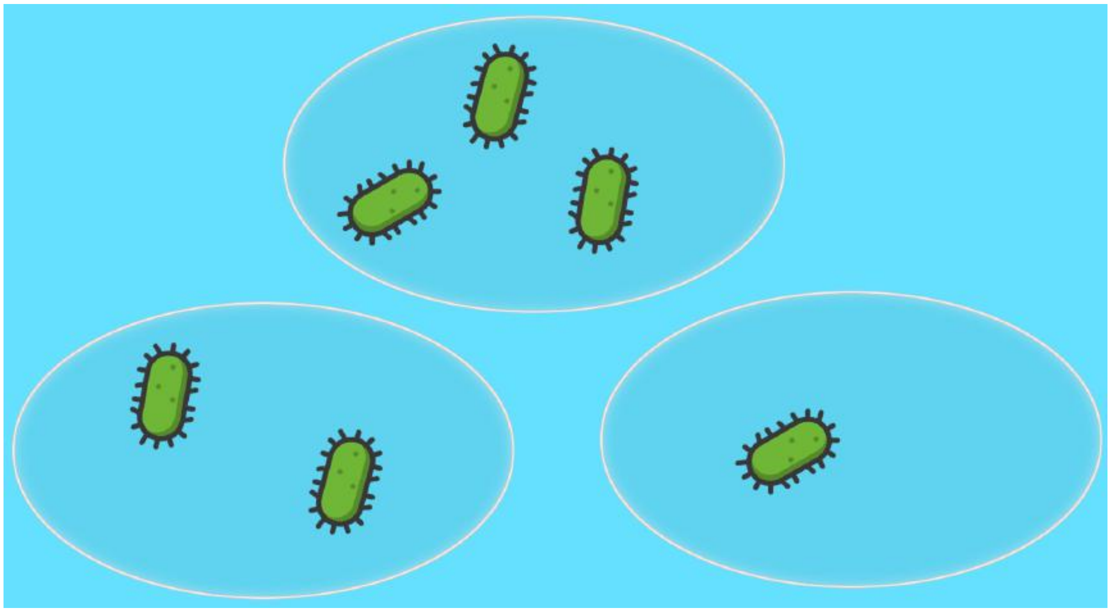

**Supplementary Figure S7.** Low Density Heterogeneity

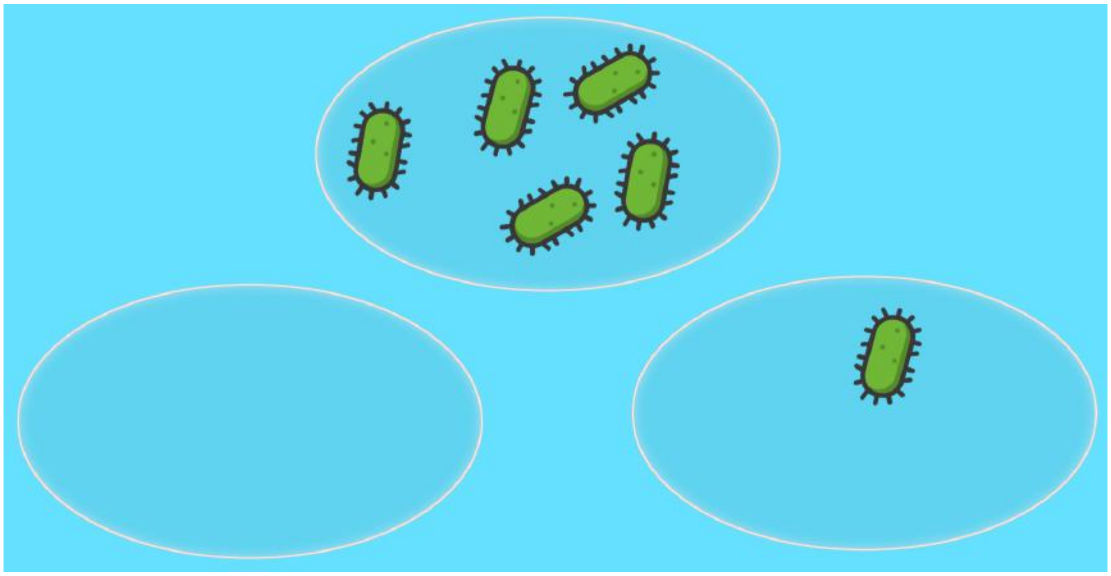

**Supplementary Figure S8.** High Density Heterogeneity

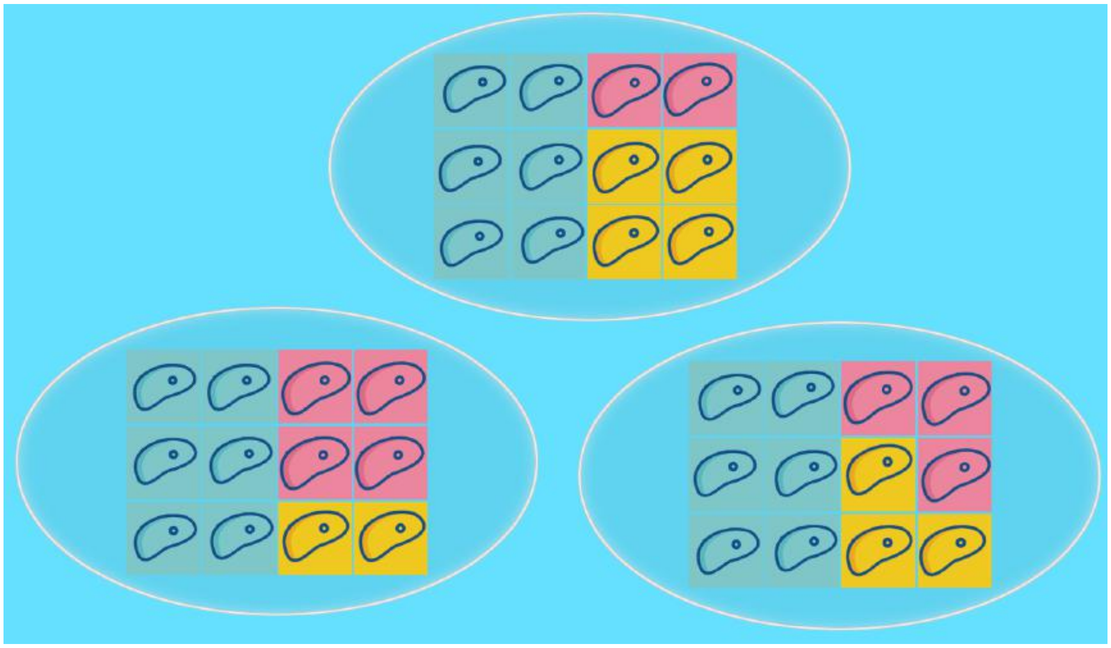

**Supplementary Figure S9.** Low Compositional Heterogeneity

blue, $\frac{1}{4}$ pink, $\frac{1}{4}$ yellow, but the average relative abundances of each strain differs between the specific sections. In figure S9 the differences are small, introducing not much more variance compared to what would be expected under the uniform/homogeneous density assumed by the default hPoMu model. In figure S10 the differences are larger, introducing much more variance compared to what would be expected under uniform/homogeneous density assumed by the default hPoMu model.

In the hierarchical count-categorical distribution framework, compositional heterogeneity is an issue which affects the categorical distribution. The default multinomial model for the categorical distribution (found in hPoMu) is a "null model" that does not account for additional variance that might be introduced in practice by compositional heterogeneity.

###### *S2.4.4 Modeling Heterogeneities*

For both density heterogeneity and compositional heterogeneity, extra variance is introduced that is not accounted for by the default hPoMu model. In trying to predict and model the probability distribution of droplet formation, instead of trying to develop complicated mechanistic models describing these spatial heterogeneities whose impacts on the initial droplet formation distribution we might then infer, we can instead seek the humbler goal of modeling this extra variance. In other words, instead of directly modeling the causes of density and compositional heterogeneity, we can restrict ourselves to modeling their "downstream" effect on variance. This should be enough by itself to make our models for the distribution of initial droplet formation more accurate.

I quantify this extra variance via the over-dispersion (defined below in section S2.4.4), and describe later in sections S2.6.1 and S2.7.1 how the parameters of the proposed models relate to their over-dispersion, and therefore can describe the increased variance caused by either density heterogeneity or compositional heterogeneity.

**Over-Dispersion** Herein we define the over-dispersion of $Y$ with respect to $X$, where $Y$ and $X$ are scalar random variables, as

$$\frac{\mathrm{Var}[Y]-\mathrm{Var}[X]}{\mathrm{Var}[X]}. \tag{S8}$$

If $Y$ and $X$ are random vectors, and if there exists a constant $C$ such that

$$\frac{\mathrm{Cov}[Y_i,Y_j]-\mathrm{Cov}[X_i,X_j]}{\mathrm{Cov}[X_i,X_j]}=C \tag{S9}$$

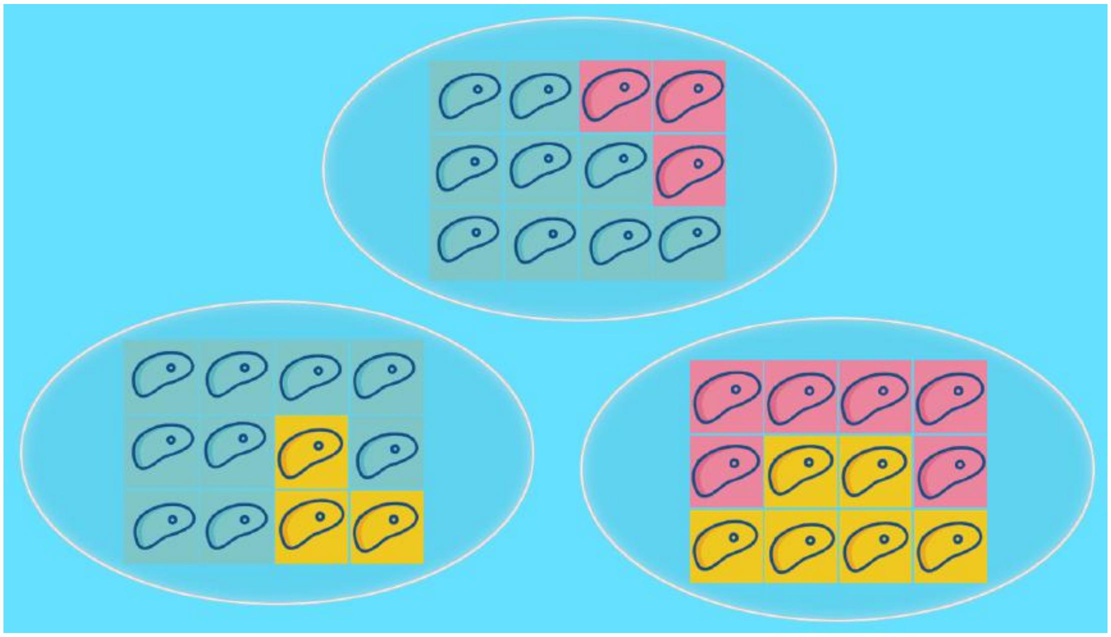

**Supplementary Figure S10.** High Compositional Heterogeneity

for all $i, j$ (including $i = j$), then $C$ is herein defined to be the over-dispersion of $Y$ with respect to $X$.

The over-dispersion of $\mathcal{P}_2$ with respect to $\mathcal{P}_1$, where $\mathcal{P}_2$ and $\mathcal{P}_1$ are probability distributions, is defined to be the over-dispersion of $Y$ with respect to $X$, where the random variables $Y \sim \mathcal{P}_2$ and $X \sim \mathcal{P}_1$ but are otherwise arbitrary.

#### S2.5 Modeling Sampling Without Replacement

The hPoMu model assumes that the number of cells of any given strain $s$ is effectively infinite, and that therefore sampling can effectively be modeled as without replacement. However, for very rare strains the adequacy of this asymptotic approximation becomes questionable. Therefore what we need is a distribution whose definition explicitly acknowledges the finiteness of the populations of cells being sampled. In section S2.5.1 I introduce a model which fills this gap, and then in section S2.5.2 I show that this model extends what can be described using hPoMu alone.

##### *S2.5.1 Introduction to hTPMH*

As suggested by its name, the "hierarchical truncated Poisson multivariate hypergeometric" (hTPMH) distribution uses a multivariate hypergeometric model of sampling without replacement for the categorical distribution. The multivariate hypergeometric distribution is introduced in section S2.5.1. The truncated Poisson distribution is introduced in section S2.5.1, and is a slight modification of the Poisson distribution that explicitly acknowledges that the number of cells to be sampled from is finite. Finally I combine the new information learned to define the hTPMH model in section S2.5.1.

**Multivariate Hypergeometric (MH) Distribution** The PMF of the multivariate hypergeometric distribution can be written as

$$\frac{(K_{d-1}-n)!}{(K_{d-1})!} \cdot \left[ \prod_{s=1}^{S} \frac{(K_{d-1}^{(s)})!}{(K_{d-1}^{(s)} - n^{(s)})!} \right] \cdot \binom{n}{n^{(1)} \cdots n^{(S)}} . \tag{S10}$$

As a reminder $n := \sum_{s=1}^{S} n^{(s)}$ (the sum of the entries of $\vec{\mathbf{n}}$) and $K_{d-1} := \sum_{s=1}^{S} K_{d-1}^{(s)}$ (the sum of the entries of $\vec{\mathbf{K}}_{d-1}$).

**Truncated Poisson (TP) Distribution** The mass function of the truncated Poisson distribution is

$$\frac{1}{\mathbb{P}(\text{Pois}(\lambda) \leq K_{d-1})} \cdot \frac{e^{-\lambda}\lambda^n}{n!} . \tag{S11}$$

Unless $K_{d-1}$ is very small, this is effectively almost identical to the corresponding probabilities from the Poisson distribution.

**hTPMH Definition** Under the hTPMH distribution, for the $d$'th droplet: the strain distribution vector $\mathbb{P}\left(\vec{\mathbf{N}}_d(0) = \vec{\mathbf{n}} \,\middle|\, N_d(0) = n, \vec{\mathbf{K}}_{d-1}\right)$ given the number of cells is multivariate hypergeometric distributed, while the number of cells $\mathbb{P}\left(N_d(0) = n \,\middle|\, \vec{\mathbf{K}}_{d-1}\right)$ is truncated Poisson distributed. Thus for the unconditional probability:

$$\begin{aligned}
&\mathbb{P}\left(\vec{\mathbf{N}}_d(0) = \vec{\mathbf{n}} \,\middle|\, \vec{\mathbf{K}}_{d-1}\right) \\
=\;& \mathbb{P}\left(N_d(0) = n \,\middle|\, \vec{\mathbf{K}}_{d-1}\right) \cdot \mathbb{P}\left(\vec{\mathbf{N}}_d(0) = \vec{\mathbf{n}} \,\middle|\, N_d(0) = n, \vec{\mathbf{K}}_{d-1}\right) \\
=\;& \mathbb{P}(\mathrm{Pois}(\lambda) \le K_{d-1})^{-1} \cdot \frac{e^{-\lambda}\lambda^n}{n!} \frac{(K_{d-1}-n)!}{(K_{d-1})!} \cdot \left[\prod_{s=1}^{S} \frac{(K_{d-1}^{(s)})!}{(K_{d-1}^{(s)} - n^{(s)})!}\right] \cdot \binom{n}{n^{(1)} \cdots n^{(S)}} .
\end{aligned} \tag{S12}$$

##### S2.5.2 hTPMH Generalizes hPoMu

One can show that hTPMH converges to the default hPoMu model under appropriate limits. Therefore when using the hTPMH family of distributions to model the distribution of the initial formation of droplets, we "retain the same language" used by the hPoMu distribution for that task.

In sections S2.5.2, S2.5.2, and S2.5.2 I give the context needed and then show that the hNBDM family really is an extension of hPoMu. Because hTPMH and hPoMu have nearly the same count distribution, at its core this reduces to showing how the multivariate hypergeometric distribution extends the multinomial distribution, the context for which is section S2.5.2.

**Approximate Form of Likelihood Ratio of MH w.r.t. Multinomial and Bounds** Applying Lemma S2.1 and then algebraic manipulations (e.g. multiplying numerator and denominator by the same factor) gives

$$\begin{aligned}
&\frac{(K_{d-1}-n)!}{(K_{d-1})!} \cdot \left[\prod_{s=1}^{S} \frac{(K_{d-1}^{(s)})!}{(K_{d-1}^{(s)} - n^{(s)})!}\right] \\
&= \exp\left(\mathscr{R}_{\mathrm{MHg}}(\vec{\mathbf{n}}, \vec{\mathbf{K}}_{d-1})\right) \cdot \left(\frac{K_{d-1}-n}{K_{d-1}}\right)^{K_{d-1}-n+\frac{1}{2}} \\
&\cdot \prod_{s=1}^{S} \left(\frac{K_{d-1}^{(s)}}{K_{d-1}^{(s)} - n^{(s)}}\right)^{K_{d-1}^{(s)} - n^{(s)} + \frac{1}{2}} \cdot \prod_{s=1}^{S} \left(\frac{K_{d-1}^{(s)}}{f^{(s)} K_{d-1}}\right)^{n^{(s)}} \cdot \prod_{s=1}^{S} (f^{(s)})^{n^{(s)}} .
\end{aligned} \tag{S13}$$

In equation S13, based on (Robbins, 1955), Lemma 1 implies that

$$\underline{\mathscr{R}}\left(\vec{\mathbf{K}}_{d-1} - \vec{\mathbf{n}}, \vec{\mathbf{K}}_{d-1}\right) < \mathscr{R}_{\mathrm{MHg}}\left(\vec{\mathbf{n}}, \vec{\mathbf{K}}_{d-1}\right) < \overline{\mathscr{R}}\left(\vec{\mathbf{K}}_{d-1} - \vec{\mathbf{n}}, \vec{\mathbf{K}}_{d-1}\right) , \tag{S14}$$

with the "lower Robbins function" $\underline{\mathscr{R}}$ defined for convenience as

$$\underline{\mathscr{R}}(\vec{N}, \vec{M}) := \frac{1}{12N+1} - \sum_{s=1}^{S} \frac{1}{12N_s} - \frac{1}{12M} + \sum_{s=1}^{S} \frac{1}{12M_s+1} = -\overline{\mathscr{R}}(\vec{M}, \vec{N}) , \tag{S15}$$

and the "upper Robbins function" $\overline{\mathscr{R}}$ defined for convenience as

$$\overline{\mathscr{R}}(\vec{N}, \vec{M}) := \frac{1}{12N} - \sum_{s=1}^{S} \frac{1}{12N_s+1} - \frac{1}{12M+1} + \sum_{s=1}^{S} \frac{1}{12M_s} = -\underline{\mathscr{R}}(\vec{M}, \vec{N}) . \tag{S16}$$

Above we again used the notational the convention that, given a vector $\vec{X} = (X_1, \ldots, X_S)$, $X := \sum_{s=1}^{S} X_s$.

Equation S13 implies that the likelihood ratio of Multivariate Hypergeometric distribution with respect to the Multinomial distribution may be written

$$\exp\left(\mathscr{R}_{\text{MHg}}(\vec{\mathbf{n}},\vec{\mathbf{K}}_{d-1})\right)\cdot\left(\frac{K_{d-1}-n}{K_{d-1}}\right)^{K_{d-1}-n+\frac{1}{2}}\cdot\prod_{s=1}^{S}\left(\frac{K_{d-1}^{(s)}}{K_{d-1}^{(s)}-n^{(s)}}\right)^{K_{d-1}^{(s)}-n^{(s)}+\frac{1}{2}}\cdot\prod_{s=1}^{S}\left(\frac{K_{d-1}^{(s)}}{f^{(s)}K_{d-1}}\right)^{n^{(s)}}. \tag{S17}$$

In other words, multiplying the PMF of the Multinomial distribution by the quantity in S17 gives the PMF of the Multivariate Hypergeometric Distribution.

**Approximate Form of Likelihood Ratio of hTPMH w.r.t. hPoMu and Bounds** From equation S17 it follows readily that the likelihood ratio of the hTPMH distribution with respect to the hPoMu distribution is

$$\begin{aligned}&\mathbb{P}(\text{Pois}(\lambda)\leq K_{d-1})^{-1}\exp\left(\mathscr{R}_{\text{MHg}}(\vec{\mathbf{n}},\vec{\mathbf{K}}_{d-1})\right)\cdot\left(\frac{K_{d-1}-n}{K_{d-1}}\right)^{K_{d-1}-n+\frac{1}{2}}\\&\cdot\prod_{s=1}^{S}\left(\frac{K_{d-1}^{(s)}}{K_{d-1}^{(s)}-n^{(s)}}\right)^{K_{d-1}^{(s)}-n^{(s)}+\frac{1}{2}}\cdot\prod_{s=1}^{S}\left(\frac{K_{d-1}^{(s)}}{f^{(s)}K_{d-1}}\right)^{n^{(s)}}.\end{aligned} \tag{S18}$$

The factor $\mathbb{P}(\text{Pois}(\lambda)\leq K_{d-1})^{-1}$ reflects the fact that the number of cells in droplet $d$ will be truncated Poisson distributed, not Poisson distributed. Otherwise this is the same as (S17).

**Proof that hTPMH Converges in Distribution to hPoMu** Because the hTPMH and hierarchical Poisson Multinomial (hPoMu) distributions are discrete, to show that the former converges in distribution to the latter, it suffices to show that the probability mass function of the former converges to the latter.

Equation S18 gives a quantity such that, when it is multiplied with the PMF of the hPoMu distribution, the result is the PMF of the corresponding hTPMH distribution.

Fixing the values of $\vec{\mathbf{N}}_{\delta}(0)$ for all $\delta\in[d-1]$, from the above it follows that the quantity in S18 converging to 1 as $K\to\infty$ implies that the hTPMH distribution converges to the hPoMu distribution as $K\to\infty$.

The desired convergence follows from the Squeeze Theorem. Note that both $\underline{\mathscr{R}}\left(\vec{\mathbf{K}}_{d-1}-\vec{\mathbf{n}},\vec{\mathbf{K}}_{d-1}\right)$ and $\overline{\mathscr{R}}\left(\vec{\mathbf{K}}_{d-1}-\vec{\mathbf{n}},\vec{\mathbf{K}}_{d-1}\right)$ converge to 0 as $K\to\infty$ because of the additivity of limits and the consequence of the Archimedean property of the real numbers that $\pm\lim_{N\to\infty}\frac{1}{a_N}=0$ for any sequence $a_N$ such that $\lim_{N\to\infty}a_N=\infty$.

The factor $\mathbb{P}(\text{Pois}(\lambda)\leq K_{d-1})^{-1}$ converges to 1 because of the defintion of $K_{d-1}$ and that any valid cumulative distribution function must converge to 1 as its input goes to $\infty$ (or otherwise there would be nonzero probability mass "at infinity").

The last expression in (S18) converges to 1 because of L'Hôspital's Rule, and the remaining factors converge to 1 because of Lemma S2.3.

We may assume without loss of generality that, when taking the limit as $K\to\infty$, we do so along a subsequence such that all $K^{(s)}=f^{(s)}K$ are in $\mathbb{N}$. This of course also guarantees that all of the $K_{d-1}^{(s)}:=K^{(s)}-\sum_{\delta=1}^{d-1}N_{\delta}^{(s)}(0)$ are also integer-valued.

Instead of showing that the likelihood ratio converges to 1, one can also show (e.g. according to one's personal preference) that the logarithm of the likelihood ratio converges to 0, using Lemma S2.5 (the "logarithmic version" of Lemma S2.3). The "fudge function" $\Phi$ is defined for convenience in equation (S47). Then, starting from equation (S18), tedious algebra shows that the logarithm of the likelihood ratio of hTPMH with respect to hPoMu is

$$\begin{aligned}&-\log\left(\mathbb{P}(\text{Pois}(\lambda)\leq K_{d-1})\right)+\sum_{s=1}^{S}\Phi(K_{d-1}^{(s)}-n^{(s)},n^{(s)},\frac{1}{2})\\&-\Phi(K_{d-1}-n,n,\frac{1}{2})+\sum_{s=1}^{S}n^{(s)}\left(\log(K_{d-1}^{(s)})-\log(f^{(s)}K_{d-1})\right)+\mathscr{R}_{\text{MHg}}(\vec{\mathbf{n}},\vec{\mathbf{K}}_{d-1}).\end{aligned} \tag{S19}$$

The only subtlety involves recalling the definitions of $K_{d-1}^{(s)}$ and $K_{d-1}$ so as to correctly substitute them into (S19):

$$\log(K_{d-1}^{(s)}) - \log(f^{(s)} K_{d-1}) = \log\left( \frac{f^{(s)} K - \sum_{\delta=1}^{d-1} N_\delta^{(s)}(0)}{f^{(s)} K - f^{(s)} \left( \sum_{\delta=1}^{d-1} N(0) \right)} \right) . \tag{S20}$$

#### S2.6 Modeling Compositional Heterogeneity

The hTPMH distribution uses a categorical distribution (the multivariate hypergeometric) which does not capture over-dispersion relative to the multinomial. In fact, its categorical distribution can also be derived using homogeneity assumptions, and only relaxes the assumptions regarding effectively infinite population size. So both hPoMu and hTPMH are unable to model over-dispersion of the categorical distribution caused by compositional heterogeneity. Thus the main change we need to make is to use a categorical distribution over-dispersed relative to the multinomial distribution.

In section S2.6.1 I introduce a model which fills this gap, and then in section S2.6.2 I show that this model extends what can be described using hPoMu alone.

##### *S2.6.1 Introduction to hPoDM*

As suggested by its name, the "hierarchical Poisson Dirichlet-Multinomial" (hPoDM) distribution fills this gap by using a new distribution for the categorical distribution, the Dirichlet-Multinomial.

In section S2.6.1 I introduce the Dirichlet-Multinomial distribution. In section S2.6.1 I explain how the Dirichlet-Multinomial distribution models over-dispersion relative to the multinomial distribution. In section S2.6.1 I use the new information learned to define the hPoDM family of distributions.

**Dirichlet-Multinomial (DM) Distribution** The PMF for the Dirichlet-Multinomial distribution may be written

$$\frac{\Gamma(\zeta_C S)}{\Gamma(\zeta_C S + n)} \cdot \left[ \prod_{s=1}^{S} \frac{\Gamma(\zeta_C S f^{(s)} + n^{(s)})}{\Gamma\left(\zeta_C S f^{(s)}\right)} \right] \cdot \binom{n}{n^{(1)} \cdots n^{(S)}} . \tag{S21}$$

More typically this is parameterized in terms of $\zeta_C S$. I parameterize in terms of $\zeta_C$ instead to facilitate interpretation. Parameterized this way, $\zeta_C = 1$ always corresponds to the same concentration as the uniform distribution on the $(S-1)$-dimensional simplex, regardless of the number of strains $S$.

**Over-Disperson of Dirichlet-Multinomial w.r.t. Multinomial** The over-dispersion of the Dirichlet-Multinomial distribution with concentration $\zeta_C S$, total number $n$, and frequencies $\vec{f}$ with respect to the corresponding Multinomial distribution with total number $n$ and frequencies $\vec{f}$ is

$$\frac{n-1}{1+\zeta_C S} . \tag{S22}$$

Thus the concentration is approximately proportional to the reciprocal of the over-dispersion. (They are asymptotically equivalent up to a constant, i.e. the limit of their ratio as the $\zeta_C \to \infty$ is the constant $n-1$.)

**hPoDM Definition** Under the hPoDM distribution, for the $d$'th droplet: the strain distribution vector $\mathbb{P}\left( \vec{\mathbf{N}}_d(0) = \vec{\mathbf{n}} \,\middle|\, N_d(0) = n \right)$ given the number of cells is Dirichlet-Multinomial distributed, while the number of cells $\mathbb{P}(N_d(0) = n)$ is Poisson distributed. Thus for the unconditional probability:

$$\begin{aligned} & \mathbb{P}\left( \vec{\mathbf{N}}_d(0) = \vec{\mathbf{n}} \right) \\ = \;& \mathbb{P}(N_d(0) = n) \cdot \mathbb{P}\left( \vec{\mathbf{N}}_d(0) = \vec{\mathbf{n}} \,\middle|\, N_d(0) = n \right) \\ = \;& \frac{e^{-\lambda} \lambda^n}{n!} \cdot \frac{\Gamma(\zeta_C S)}{\Gamma(\zeta_C S + n)} \cdot \left[ \prod_{s=1}^{S} \frac{\Gamma(\zeta_C S f^{(s)} + n^{(s)})}{\Gamma\left(\zeta_C S f^{(s)}\right)} \right] \cdot \binom{n}{n^{(1)} \cdots n^{(S)}} . \end{aligned} \tag{S23}$$

##### *S2.6.2 hPoDM Generalizes hPoMu*

Just like for hTPMH, hPoDM converges to the default hPoMu model under appropriate limits. Therefore when using the hPoDM family of distributions to model the distribution of the initial formation of droplets, we "retain the same language" used by the hPoMu distribution for that task.

In sections S2.6.2 and S2.6.2 I give the context needed and then show that the hNBDM family really is an extension of hPoMu. Because hPoDM and hPoMu have the same count distribution, this reduces to showing how the Dirichlet-Multinomial distribution extends the multinomial distribution.

**Approximate Form of Likelihood Ratio of DM w.r.t. Multinomial and hPoDM w.r.t. hPoMu and Bounds** Using Lemma S2.4 we get that

$$
\begin{aligned}
&\frac{\Gamma(\zeta_C S)}{\Gamma(\zeta_C S+n)}\cdot\left[\prod_{s=1}^{S}\frac{\Gamma(\zeta_C S f^{(s)}+n^{(s)})}{\Gamma(\zeta_C S f^{(s)})}\right]\\
&=\exp\left(\mathscr{R}_{\mathrm{DM}}(\vec{\mathbf{n}},\zeta_C S\vec{f})\right)\cdot\left(\frac{\zeta_C S}{\zeta_C S+n}\right)^{\zeta_C S+n-\frac{1}{2}}\\
&\cdot\left[\prod_{s=1}^{S}\left(\frac{\zeta_C S f^{(s)}+n^{(s)}}{\zeta_C S f^{(s)}}\right)^{\zeta_C S f^{(s)}+n^{(s)}-\frac{1}{2}}\right]\cdot\prod_{s=1}^{S}(f^{(s)})^{n^{(s)}},
\end{aligned}
\tag{S24}
$$

where (with the upper $\overline{\mathscr{R}}$ and lower $\underline{\mathscr{R}}$ functions as defined before)

$$
\underline{\mathscr{R}}(\zeta_C S\vec{f},\zeta_C S\vec{f}+\vec{\mathbf{n}})<\mathscr{R}_{\mathrm{DM}}(\vec{\mathbf{n}},\zeta_C S\vec{f})<\overline{\mathscr{R}}(\zeta_C S\vec{f},\zeta_C S\vec{f}+\vec{\mathbf{n}})\,. \tag{S25}
$$

Unfortunately the bounds in (S25) appear to be tight in general only for larger values of $\zeta_C$. (They become tighter as $\zeta_C\to\infty$; cf. section S2.6.2.)

Thus the likelihood ratio of the Dirichlet-Multinomial distribution with respect to the Multinomial distribution, which is the same as the likelihood ratio of the hPoDM distribution with respect to the hPoMu distribution, based on equation S24 above equals

$$
\exp\left(\mathscr{R}_{\mathrm{DM}}(\vec{\mathbf{n}},\zeta_C S\vec{f})\right)\cdot\left(\frac{\zeta_C S}{\zeta_C S+n}\right)^{\zeta_C S+n-\frac{1}{2}}\cdot\left[\prod_{s=1}^{S}\left(\frac{\zeta_C S f^{(s)}+n^{(s)}}{\zeta_C S f^{(s)}}\right)^{\zeta_C S f^{(s)}+n^{(s)}-\frac{1}{2}}\right]. \tag{S26}
$$

**Proof that hPoDM Converges in Distribution to hPoMu** Again, it suffices to show that the probability mass function of hPoDM converges to that of hPoMu. Cf. section S2.5.2.

That the expression in (S26) approaches 1 as $\zeta_C\to\infty$ again follows from the Squeeze Theorem and Lemma S2.3. (As before, that the expressions involving the upper and lower Robbins functions approach 0 is an elementary consequence of the Archimedean property of the real numbers.) Thus the hPoDM distribution can be seen to converge to the hPoMu distribution as $\zeta_C\to\infty$.

Because the likelihood ratio of the Dirichlet-Multinomial distribution with respect to the Multinomial distribution is the same as the likelihood ratio of the hPoDM distribution with respect to the hPoMu distribution, it follows that the above also shows how the Dirichlet-Multinomial distribution converges to the Multinomial distribution as $\zeta_C\to\infty$.

Instead of showing that the likelihood ratio converges to 1, one can also show (e.g. according to one's personal preference) that the logarithm of the likelihood ratio converges to 0, using Lemma S2.5 (the "logarithmic version" of Lemma S2.3). The "fudge function" $\Phi$ is defined for convenience in equation (S47). Then, starting from (S26), tedious algebra shows that the logarithm of the likelihood ratio of (hPo)DM with respect to (hPo)Mu is

$$
\sum_{s=1}^{S}\Phi(\zeta_C S f^{(s)},n^{(s)},n^{(s)}-\frac{1}{2})-\Phi(\zeta_C S,n,n-\frac{1}{2})+\mathscr{R}_{\mathrm{DM}}(\vec{\mathbf{n}},\zeta_C S\vec{f})\,. \tag{S27}
$$

#### S2.7 Modeling Density Heterogeneity

The hPoDM distribution leaves the count distribution unchanged relative to hPoMu, and therefore is unable to model over-dispersion of the count distribution caused by density heterogeneity. Thus the main change we need to make is from using the Poisson distribution as the count distribution to using a distribution over-dispersed relative to the Poisson distribution. In section S2.7.1 I introduce a model which fills this gap, and then in section S2.7.2 I show that this model extends what can be described using hPoMu alone.

###### *S2.7.1 Introduction to hNBDM*

As suggested by its name, the "hierarchical Negative Binomial Dirichlet-Multinomial" (hNBDM) distribution fills this gap by using a new distribution for the count distribution, the negative binomial.

In section S2.7.1 I introduce the negative binomial distribution. In section S2.7.1 I explain how the negative binomial distribution models over-dispersion relative to the Poisson distribution. In section S2.7.1 I use the new information learned to define the hNBDM family of distributions.

**Negative Binomial (NB) Distribution** One may write the PMF of the negative binomial distribution as

$$\frac{\Gamma(\zeta_D S+n)}{\Gamma(\zeta_D S)}\cdot\frac{(\zeta_D S)^{\zeta_D S}}{(\zeta_D S+\lambda)^{\zeta_D S+n}}\cdot\frac{\lambda^n}{n!} \tag{S28}$$

where $\lambda$ denotes the expected value of the distribution.

More typically this is parameterized in terms of $\zeta_D S$ instead of $\zeta_D$. (Indeed in principle it should not even be necessary to specify a number of strains $S$ in order to define this distribution.) I parameterize in terms of $\zeta_D$ to be compatible with the parameterization of the Dirichlet-Multinomial I use in section S2.6.1. See section S2.7.1 for how this pays off later by simplifying the math.

The parameter $\zeta_D$ is herein called the "density concentration". Using the negative binomial distribution for modelling density heterogeneity has precedent in the literature. For macroecological studies, the parameter $\lambda$ "has been defined as the density of organisms in the area of interest" (Willson et al., 1984). Herein $\lambda$ will be interpreted as the *average* cell density in the sampling pool. Previous work has also claimed that $\zeta_D$ should be able to capture variability in the density of organisms: "The practical use of [$\zeta_D S$] ... in ecological studies of aggregation requires caution because [$\zeta_D S$] is usually density-dependent" (Clark and Perry, 1989).

If a Poisson distribution with rate $\lambda$ corresponds to a homogeneous cell density $\lambda$ throughout the sampling pool, a negative binomial distribution with rate $\lambda$ is interpreted to have the same *average* cell density. Larger values of density concentration $\zeta_D$ correspond to *smaller* heterogeneity around the average density, and thus more closely resemble the Poisson case (which is the limit as $\zeta_D \to \infty$), cf. figure S7. Similarly, smaller values of density concentration $\zeta_D$ correspond to *larger* heterogeneity around the average density, cf. figure S8.

**Over-Disperson of Negative Binomial w.r.t. Poisson** The over-dispersion of the negative binomial distribution with concentration $\zeta_D S$ and mean $\lambda$ with respect to the Poisson distribution with mean $\lambda$ is

$$\frac{\lambda}{\zeta_D S}\,. \tag{S29}$$

Thus the concentration is proportional to the reciprocal of the over-dispersion.

**hNBDM Definition** Setting $\zeta_D = \zeta_C =: \zeta$, the PMF of the hierarchical Negative Binomial Dirichlet-Multinomial (hNBDM) distribution is

$$\frac{\Gamma(\zeta S+n)}{\Gamma(\zeta S)}\cdot\frac{(\zeta S)^{\zeta S}}{(\zeta S+\lambda)^{\zeta S+n}}\cdot\frac{\lambda^n}{n!}\,\frac{\Gamma(\zeta S)}{\Gamma(\zeta S+n)}\cdot\left[\prod_{s=1}^{S}\frac{\Gamma(\zeta S f^{(s)}+n^{(s)})}{\Gamma\left(\zeta S f^{(s)}\right)}\right]\cdot\binom{n}{n^{(1)}\cdots n^{(S)}} \tag{S30}$$

which via tedious algebraic manipulations can be seen to equal

$$\prod_{s=1}^{S}\left[\frac{\Gamma(\zeta S f^{(s)}+n^{(s)})}{\Gamma(\zeta S f^{(s)})}\cdot\frac{(\zeta S f^{(s)})^{\zeta S f^{(s)}}}{(\zeta S f^{(s)}+f^{(s)}\lambda)^{\zeta S f^{(s)}+n^{(s)}}}\cdot\frac{(f^{(s)}\lambda)^{n^{(s)}}}{(n^{(s)})!}\right]. \tag{S31}$$

In other words, the marginal distributions of the hNBDM distribution correspond to mutually independent Negative Binomial random variables whose parameters have been rescaled by the $f^{(s)}$.

This makes the relationship of the Negative Binomial distribution to the Dirichlet-Multinomial a direct analogue of the relationship of the Poisson distribution to the Multinomial Distribution. Unsurprisingly these observations have precedent in the literature. Cf. for example Theorem 1 of (Zhou, 2018).

##### *S2.7.2 hNBDM Generalizes hPoMu*

Just like for hTPMH and hPoDM, hNBDM converges to the default hPoMu model under appropriate limits. Therefore when using the hNBDM family of distributions to model the distribution of the initial formation of droplets, we “retain the same language” used by the hPoMu distribution for that task.

In sections S2.7.2 and S2.7.2 I give the context needed and then show that, as a count distribution, the negative binomial distribution truly is a generalization of the Poisson. In sections S2.7.2 and S2.7.2 I give the context needed and then show that the hNBDM family really is an extension of hPoMu.

**Approximate Form of Likelihood Ratio of NB w.r.t. Poisson and Bounds** Applying Lemma S2.4, one has

$$\frac{\Gamma(\zeta_D S+n)}{\Gamma(\zeta_D S)} = e^{-n}\cdot\left(\frac{\zeta_D S+n}{\zeta_D S}\right)^{\zeta_D S+n-\frac{1}{2}}\cdot(\zeta_D S)^n\cdot\exp\left(\mathscr{R}_{\mathrm{NB}}(\zeta_D S,n)\right), \tag{S32}$$

where $\mathscr{R}_{\mathrm{NB}}(\zeta_D S,n)$ satisfies

$$\frac{1}{12(\zeta_D S+n)+1}-\frac{1}{12\zeta_D S} < \mathscr{R}_{\mathrm{NB}}(\zeta_D S,n) < \frac{1}{12(\zeta_D S+n)}-\frac{1}{12\zeta_D S+1}. \tag{S33}$$

It follows from equations (S28) and (S32) that the likelihood ratio of the negative binomial distribution (with mean $\lambda$) with respect to the Poisson distribution (with mean $\lambda$) is

$$e^{-n}\cdot\left(\frac{\zeta_D S+n}{\zeta_D S}\right)^{\zeta_D S+n-\frac{1}{2}}\cdot\left(\frac{\zeta_D S}{\zeta_D S+\lambda}\right)^{\zeta_D S+n}\cdot e^{\lambda}\cdot\exp\left(\mathscr{R}_{\mathrm{NB}}(\zeta_D S,n)\right). \tag{S34}$$

**Proof that NB Converges in Distribution to Poisson** It follows nearly immediately from Lemma S2.3 that the above expression (S34) converges to 1 as $\zeta_D\to\infty$. (Again, both of the bounds for $\mathscr{R}_{\mathrm{NB}}(\zeta_D S,n)$ from (S33) approach 0 for trivial reasons, allowing us to apply the Squeeze Theorem.) In particular, it follows that the negative binomial distribution converges to the Poisson distribution as $\zeta_D\to\infty$.

**Approximate Form of Likelihood Ratio of hNBDM w.r.t. hPoMu and Bounds** Using either of the expressions (S30) or (S31) for the PMF of the hNBDM distribution given above, more tedious algebraic manipulations (including application of Lemma S2.4) allow one to show that the likelihood ratio of the hNBDM distribution relative to the corresponding hPoMu distribution is

$$\prod_{s=1}^{S}\left[e^{-n^{(s)}}\cdot\left(\frac{\zeta S f^{(s)}+n^{(s)}}{\zeta S f^{(s)}}\right)^{\zeta S f^{(s)}+n^{(s)}-\frac{1}{2}}\cdot\left(\frac{\zeta S f^{(s)}}{\zeta S f^{(s)}+f^{(s)}\lambda}\right)^{\zeta S f^{(s)}+n^{(s)}}\cdot e^{f^{(s)}\lambda}\right]\cdot\mathscr{R}_{\mathrm{hNBDM}}(\zeta S\vec{f},\vec{\mathbf{n}}), \tag{S35}$$

where

$$\mathscr{R}_{\mathrm{hNBDM}}(\zeta S\vec{f},\vec{\mathbf{n}}) = \mathscr{R}_{\mathrm{NB}}(\zeta S,n)+\mathscr{R}_{\mathrm{DM}}(\zeta S\vec{f},\vec{\mathbf{n}}) = \sum_{s=1}^{S}\mathscr{R}_{\mathrm{NB}}(\zeta S f^{(s)},n^{(s)}). \tag{S36}$$

Observe that adding the bounds from (S25) and (S33) gives the following bounds for $\mathscr{R}_{\mathrm{hNBDM}}(\zeta S\vec{f},\vec{\mathbf{n}})$:

$$\begin{aligned}
&\sum_{s=1}^{S}\left[\frac{1}{12(\zeta S f^{(s)}+n^{(s)})+1}-\frac{1}{12\zeta S f^{(s)}}\right]+\left(\frac{1}{12\zeta S+1}-\frac{1}{12\zeta S}\right)+\left(\frac{1}{12(\zeta S+n)+1}-\frac{1}{12(\zeta S+n)}\right)\\
=\;& \underline{\mathscr{R}}(\zeta S\vec{f},\zeta S\vec{f}+\vec{\mathbf{n}})+\frac{1}{12(\zeta S+n)+1}-\frac{1}{12\zeta S}\\
<\;& \mathscr{R}_{\mathrm{DM}}(\zeta S\vec{f},\vec{\mathbf{n}})+\mathscr{R}_{\mathrm{NB}}(\zeta S,n)=\mathscr{R}_{\mathrm{hNBDM}}(\zeta S\vec{f},\vec{\mathbf{n}})\\
<\;& \overline{\mathscr{R}}(\zeta S\vec{f},\zeta S\vec{f}+\vec{\mathbf{n}})+\frac{1}{12(\zeta S+n)}-\frac{1}{12\zeta S+1}\\
=\;&\sum_{s=1}^{S}\left[\frac{1}{12(\zeta S f^{(s)}+n^{(s)})}-\frac{1}{12\zeta S f^{(s)}+1}\right]+\left(\frac{1}{12\zeta S}-\frac{1}{12\zeta S+1}\right)+\left(\frac{1}{12(\zeta S+n)}-\frac{1}{12(\zeta S+n)+1}\right).
\end{aligned} \tag{S37}$$

Compared to the above, observe that summing (over $s$) the bounds from (S33) based on (S28) actually gives strictly tighter bounds:

$$\begin{aligned}&\sum_{s=1}^{S}\left[\frac{1}{12(\zeta S f^{(s)}+n^{(s)})+1}-\frac{1}{12\zeta S f^{(s)}}\right]\\ &<\sum_{s=1}^{S}\mathscr{R}_{\text{NB}}(\zeta S f^{(s)},n^{(s)})=\mathscr{R}_{\text{hNBDM}}(\zeta S\vec{f},\vec{\mathbf{n}})\\ &<\sum_{s=1}^{S}\left[\frac{1}{12(\zeta S f^{(s)}+n^{(s)})}-\frac{1}{12\zeta S f^{(s)}+1}\right].\end{aligned} \tag{S38}$$

The additional terms from the bounds in (S37) not present in the bounds from (S38) are the result of twice applying Lemma S2.4 unnecessarily, once to $\frac{\Gamma(\zeta S+n)}{\Gamma(\zeta S)}$ and once to $\frac{\Gamma(\zeta S)}{\Gamma(\zeta S+n)}$. Because they cancel each other out, they obviously do not need to be bounded. Therefore the bounds do not contradict one another.

**Proof that hNBDM Converges in Distribution to hPoMu** Again, it suffices to show that the probability mass function of hNBDM converges to that of hPoMu. Cf. section S2.5.2.

The fact that (S35) converges to 1 as $\zeta\to\infty$ quickly follows from Lemma S2.3. This convergence implies that the hNBDM distribution converges to the hPoMu distribution as $\zeta\to\infty$.

Equivalently, the convergence of the hNBDM distribution to the hPoMu distribution as $\zeta\to\infty$ also follows from the convergence of (S34) to 1 as $\zeta\to\infty$ (replacing $\zeta$, $n$, and $\lambda$ with $\zeta f^{(s)}$, $n^{(s)}$, and $f^{(s)}\lambda$ respectively). Again equivalently, it also follows from the convergence of both (S34) and (S26) to 1 as $\zeta\to\infty$. This is so because (S35) equals the product of (S34) and (S26).

Instead of showing that the likelihood ratio converges to 1, one can also show (e.g. according to one's personal preference) that the logarithm of the likelihood ratio converges to 0, using Lemma S2.5 (the "logarithmic version" of Lemma S2.3). The "fudge function" $\Phi$ is defined for convenience in equation (S47). Then, starting from (S35), tedious algebra shows that the logarithm of the likelihood ratio of hNBDM with respect to hPoMu is

$$\lambda-n+\sum_{s=1}^{S}\Phi(\zeta S f^{(s)},n^{(s)},n^{(s)}-\frac{1}{2})-\sum_{s=1}^{S}\Phi(\zeta S f^{(s)},f^{(s)}\lambda,n^{(s)})+\mathscr{R}_{\text{hNBDM}}(\zeta S\vec{f},\vec{\mathbf{n}}). \tag{S39}$$

#### S2.8 Modeling Arbitrary Combinations of Heterogeneities

While the hNBDM family of distributions is useful for modeling density heterogeneity, it does have the limitation that the value of density heterogeneity is assumed to be coupled with corresponding value of compositional heterogeneity. By relaxing this restriction we can generalize the hNBDM family to the "generalized hNBDM" or "ghNBDM" family of distributions. This makes it possible to model arbitrary combinations of density and compositional heterogeneities.

##### *S2.8.1 ghNBDM Family Definition*

Under a distribution in the ghNBDM family, for the $d$'th droplet: the probability for the strain distribution vector given the number of cells $\mathbb{P}\left(\vec{\mathbf{N}}_d(0)=\vec{\mathbf{n}}\,\middle|\,N_d(0)=n\right)$ is described by a Dirichlet-Multinomial distribution with concentration parameter $\zeta_C$, while the probability for the number of cells $\mathbb{P}(N_d(0)=n)$ is described by a Negative binomial distribution with concentration parameter $\zeta_D$. Thus for the unconditional probability:

$$\begin{aligned}&\mathbb{P}\left(\vec{\mathbf{N}}_d(0)=\vec{\mathbf{n}}\right)\\ =\;&\mathbb{P}(N_d(0)=n)\cdot\mathbb{P}\left(\vec{\mathbf{N}}_d(0)=\vec{\mathbf{n}}\,\middle|\,N_d(0)=n\right)\\ =\;&\frac{\Gamma(\zeta_D S+n)}{\Gamma(\zeta_D S)}\cdot\frac{(\zeta_D S)^{\zeta_D S}}{(\zeta_D S+\lambda)^{\zeta_D S+n}}\cdot\frac{\lambda^n}{n!}\cdot\frac{\Gamma(\zeta_C S)}{\Gamma(\zeta_C S+n)}\cdot\left[\prod_{s=1}^{S}\frac{\Gamma(\zeta_C S f^{(s)}+n^{(s)})}{\Gamma\left(\zeta_C S f^{(s)}\right)}\right]\cdot\binom{n}{n^{(1)}\cdots n^{(S)}}.\end{aligned} \tag{S40}$$

Unlike for the hNBDM family, for arbitrary members of the ghNBDM family one may have that $\zeta_C\neq\zeta_D$. In general the marginal distributions of members of the ghNBDM family are *not* mutually independent (and in fact have non-zero cross-covariance) and need not be negative binomial distributed.

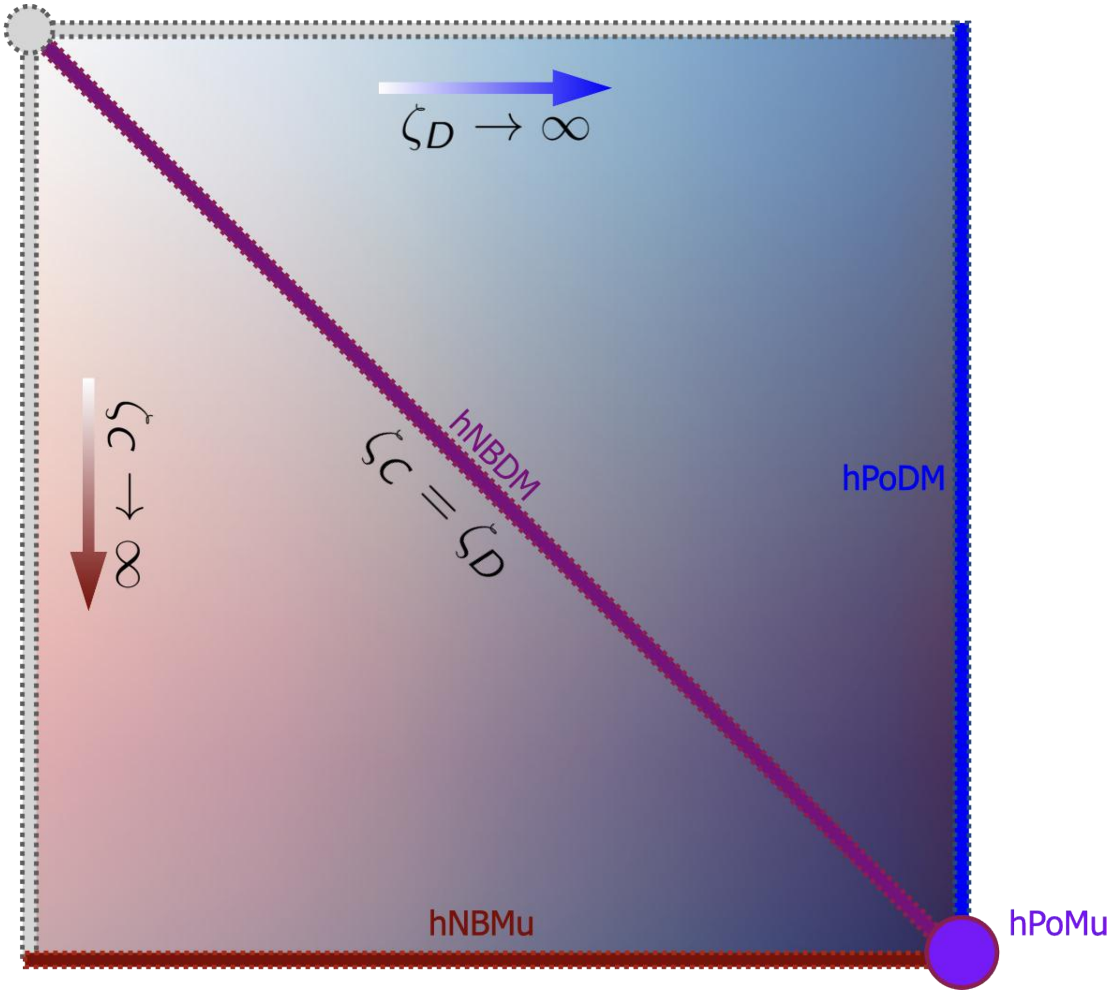

**Supplementary Figure S11.** Schematic of the ghNBDM family of distributions, assuming a fixed $\lambda$ and vector of population strain frequencies $\vec{f}$.

The hNBDM family is the special case of the ghNBDM family when $\zeta_C = \zeta_D$. The hPoDM and hPoMu distributions emerge as limiting cases of the ghNBDM family, or are bona fide members if we allow "$\infty$" as a valid concentration parameter value. Cf. figure S11. The case where $\zeta_C$ is allowed to approach infinity while $\zeta_D$ remains finite corresponds to the hierarchical Negative Binomial Multinomial (hNBMu) distribution.

###### *S2.8.2 Over-Dispersions for these Distributions*

Since the non-diagonal covariances of the hPoMu distribution are 0, it of course does not make sense to speak of over-dispersion relative to them using the definition (S9). However, we may still sensibly speak of the over-dispersion of their marginals with respect to the corresponding marginals of the hPoMu distribution (which unsurprisingly we will now do).

The over-dispersion of the $s$'th marginal of the ghNBDM distribution with respect to the $s$'th marginal of the hPoMu distribution is

$$\frac{[(1+\zeta_D S) + f^{(s)}(\zeta_C S - \zeta_D S)]}{(1+\zeta_C S)(\zeta_D S)}\lambda\,. \tag{S41}$$

Observe how this value depends on the particular $s$. Moreover, the expression (S41) above:

- reduces to (S42) when $\zeta_D = \zeta_C$,
- approaches (S43) when $\zeta_D \to \infty$,

- approaches (S44) when $\zeta_C \to \infty$,
- approaches 0 when both $\zeta_D \to \infty$ and $\zeta_C \to \infty$.

The last corresponds to hPoMu, of course. Details about the remaining special cases (hNBDM, hPoDM, hNBMu) are given below.

**hNBDM Over-dispersion** The over-dispersion of the $s$'th marginal of the hNBDM distribution with respect to the $s$'th marginal of the hPoMu distribution is

$$\frac{\lambda}{\zeta S}, \tag{S42}$$

which interestingly does *not* depend on the particular strain $s$, and moreover equals the over-dispersion of a Negative Binomial distribution with respect to a Poisson distribution, cf. (S29).

The latter is to be expected given that the marginals of the hPoMu distribution are $\mathrm{Pois}(f^{(s)}\lambda)$ distributed and the marginals of the hNBDM distribution are $\mathrm{NB}(\zeta S f^{(s)}, f^{(s)}\lambda)$ distributed. Note that (S42) also tends to 0 as $\zeta \to \infty$ as it should. Finally observe how the over-dispersion (S42) is inversely proportional to the concentration parameter $\zeta$, just like (S29) which it equals.

**hPoDM Over-dispersion** The over-dispersion of the $s$'th marginal of the hPoDM distribution with respect to the $s$'th marginal of the hPoMu distribution is

$$\frac{1-f^{(s)}}{1+\zeta_C S}\lambda\,. \tag{S43}$$

Observe how this value depends on the particular $s$. And of course (S43) tends to 0 as $\zeta_C \to \infty$. Further observe how the over-dispersion (S43) is (asymptotically) proportional to the inverse of the concentration parameter $\zeta_C$, just like (S22).

**hNBMu Over-dispersion** The over-dispersion of the $s$'th marginal of the hNBMu distribution with respect to the $s$'th marginal of the hPoMu distribution is

$$\frac{f^{(s)}\lambda}{\zeta_D S}\,. \tag{S44}$$

Observe how this value depends on the particular $s$. And of course (S44) tends to 0 as $\zeta_D \to \infty$. Further observe how the over-dispersion (S44) is proportional to the inverse of the concentration parameter $\zeta_D$, just like (S22).

#### S2.9 Discussion and Next Steps

Now that we have identified

- a default model we can use to for the initial droplet formation,
- key implicit assumptions made by that default model,
- concrete ways those assumptions could fail to be true in practice, and
- models generalizing the default model which can account for any of those failed assumptions,

we can start to ask how realistic the default model is and what changes might need to be made to use a more realistic model instead.

In section S3 we compare the default model with several of these alternative models to see whether there are noticeable differences. In section S4 we look at the effects of these alternative models on the throughput of data useful for characterizing microbial interactions. Finally in section S5 we propose and investigate the performance of estimators which will allow us to ascertain which of these models is most realistic in practice. All of this means that the data-generating process for MOREI can be simulated more accurately, which will ultimately help to identify best-performing methods for characterizing microbial interactions based on MOREI data.

#### S2.10 Results for Bounding Approximate Likelihood Ratios

**Lemma S2.1.** *Using Stirling's Approximation as given in (Robbins, 1955),*

$$
\begin{aligned}
&\frac{1}{12(n+m)+1} - \frac{1}{12n} \\
&< \log\left( \left( \frac{(n+m)!}{n!} \right) \Big/ \left( e^{-m} \left( \frac{n+m}{n} \right)^{n+m+\frac{1}{2}} n^m \right) \right) \\
&< \frac{1}{12(n+m)} - \frac{1}{12n+1} \,.
\end{aligned}
\tag{S45}
$$

**Proof:** This follows basically immediately from the bounds in (Robbins, 1955) via simple but tedious algebraic manipulations. □

Heuristically, this says that $\frac{(n+m)!}{n!} \approx n^m$ when $n$ is large. This is because the expression $\left(\frac{n+m}{n}\right)^{n+m+\frac{1}{2}}$ approaches $e^m$ as $n \to \infty$ by Lemma S2.3.

**Lemma S2.2.** *Given $\alpha, c, y_1, y_2 \in \mathbb{R}$ (in particular $c$ is constant w.r.t. $x$):*

$$
\lim_{x\to\infty} \left( \frac{\alpha x + y_1}{\alpha x + y_2} \right)^c = 1 \,.
$$

**Proof:** By properties of limits, since $f(x) = x^c$ is continuous, this equals

$$
\left[ \lim_{x\to\infty} \left( \frac{\alpha x + y_1}{\alpha x + y_2} \right) \right]^c ,
$$

which by L'Hôspital's Rule equals $1^c = 1$. □

**Lemma S2.3.** *Given $\alpha, c, y_1, y_2 \in \mathbb{R}$, $\alpha > 0$, one has*

$$
\lim_{x\to\infty} \left( \frac{\alpha x + y_1}{\alpha x + y_2} \right)^{\alpha x + c} = e^{y_1} e^{-y_2} \,.
$$

**Proof:** By properties of limits, the LHS of the above equals

$$
\left[ \lim_{x\to\infty} \left( \frac{\alpha x + y_1}{\alpha x + y_2} \right)^{\alpha x} \right] \left[ \lim_{x\to\infty} \left( \frac{\alpha x + y_1}{\alpha x + y_2} \right)^{c} \right] = \lim_{x\to\infty} \left( \frac{\alpha x + y_1}{\alpha x + y_2} \right)^{\alpha x} ,
$$

where the second equality uses Lemma S2.2. Rearranging further:

$$
= \lim_{x\to\infty} \left[ \left( \left( \frac{x + \frac{y_1}{\alpha}}{x} \right)^x \right)^{\alpha} \left( \left( \frac{x + \frac{y_2}{\alpha}}{x} \right)^x \right)^{-\alpha} \right] ,
$$

from which the result more or less immediately follows from the properties of limits and the definition of the exponential function. *(In particular because $f_1(x) = x^{\alpha}$ and $f_2(x) = x^{-\alpha}$ are continuous.)* □

**Lemma S2.4.** *Using the generalization of the bounds on Stirling's approximation to the entire $\Gamma$ function as given in Theorem 5 of (Gordon, 1994)*

$$
\begin{aligned}
&\frac{1}{12(x+\beta)+1} - \frac{1}{12x} \\
&< \log\left( \left( \frac{\Gamma(x+\beta)}{\Gamma(x)} \right) \Big/ \left( e^{-\beta} \left( \frac{x+\beta}{x} \right)^{x+\beta-\frac{1}{2}} x^{\beta} \right) \right) \\
&< \frac{1}{12(x+\beta)} - \frac{1}{12x+1} \,.
\end{aligned}
\tag{S46}
$$

**Proof:** Lemma S2.4 follows almost immediately from Theorem 5 of (Gordon, 1994) via tedious algebraic manipulations. □

Heuristically, Lemma S2.4 can be interpreted as saying that $\frac{\Gamma(x+\beta)}{\Gamma(x)} \approx x^{\beta}$ for large $x$. Again, Lemma S2.3 implies that $\left(\frac{x+\beta}{x}\right)^{x+\beta-\frac{1}{2}}$ approaches $e^{\beta}$ as $x \to \infty$.

The "fudge function" $\Phi$ is defined as

$$\Phi(x,y,z) := (x+z)\left[\log(x+y) - \log(x)\right]. \tag{S47}$$

**Lemma S2.5.** *Given $y, z \in \mathbb{R}$ which are constant with respect to $x \in \mathbb{R}$,*

$$\lim_{x\to\infty} \Phi(x,y,z) = y.$$

**Proof:** This is a corollary of Lemma S2.3. Specifically, because the logarithm is continuous and continuous functions commute with limits,

$$\lim_{x\to\infty} \Phi(x,y,z) = \log\left(\lim_{x\to\infty} \exp(\Phi(x,y,z))\right) = \log\left(\lim_{x\to\infty}\left(\frac{x+y}{x}\right)^{x+z}\right)$$
$$= \log(e^{y}) = y,$$

where the penultimate equality was an application of S2.3. □

##### *S2.10.1 Comments about Robbins-like Bounds*

The bounds on $n!$ given in (Robbins, 1955) follow from those for the $\Gamma$ function given in Theorem 5 of (Gordon, 1994) using the identity $n! = n\Gamma(n) = n(n-1)!$ (for all $n \in \mathbb{N}$). They are still slightly different though, because for $n \in \mathbb{N}$ the expression from (Gordon, 1994) gives $\Gamma(n) = \frac{n!}{n} = (n-1)! \sim \sqrt{2\pi}n^{n-\frac{1}{2}}e^{-n}$, whereas *directly*[17] applying the expression from (Robbins, 1955) to $(n-1)$ gives $(n-1)! \sim \sqrt{2\pi}(n-1)^{n-\frac{1}{2}}e^{-(n-1)}$. Although these expressions are technically distinct, it follows from Lemma S2.3 that they are asymptotically equivalent ($\sim$, the limit of their ratio approaches 1 as $n \to \infty$). Therefore they are consistent with one another.

#### S2.11 Cumulants of Hierarchical Count-Categorical Distributions

The following are consequences of the law of total covariance (of which the law of total variance is a special case) and in turn from the law of total expectation.

**Lemma S2.6.** *If $\vec{\mathbf{X}}$ is such that $\vec{\mathbf{X}}|N \sim \text{Mult}(N,\vec{f})$ or $\vec{\mathbf{X}}|N \sim \text{DirMult}(\zeta_C S, N, \vec{f})$, then for all $s \in [S]$:*

$$\mathbb{E}[X_s] = f^{(s)}\mathbb{E}[N].$$

**Lemma S2.7.** *If $\vec{\mathbf{X}}$ is such that $\vec{\mathbf{X}}|N \sim \text{Mult}(N,\vec{f})$, then for all $s \in [S]$:*

$$\text{Var}[X_s] = (f^{(s)})^2\,\text{Var}[N] + f^{(s)}(1-f^{(s)})\mathbb{E}[N]. \tag{S48}$$

**Lemma S2.8.** *If $\vec{\mathbf{X}}$ is such that $\vec{\mathbf{X}}|N \sim \text{DirMult}(\zeta_C S, N, \vec{f})$, then for all $s \in [S]$:*

$$\text{Var}[X_s] = \left[\frac{f^{(s)}(1-f^{(s)})}{1+\zeta_C S} + (f^{(s)})^2\right]\text{Var}[N] + \frac{fs(1-f^{(s)})}{1+\zeta_C S}\mathbb{E}[N](\mathbb{E}[N] + \zeta_C S). \tag{S49}$$

Observe how (using L'Hôspital's rule) equation (S49) approaches (S48) as $\zeta_C \to \infty$.

**Lemma S2.9.** *If $\vec{\mathbf{X}}$ is such that $\vec{\mathbf{X}}|N \sim \text{Mult}(N,\vec{f})$, then for all $s_1, s_2 \in [S]$ ($s_1 \neq s_2$):*

$$\text{Cov}[X_{s_1}, X_{s_2}] = f^{(s_1)}f^{(s_2)}\left(\text{Var}[N] - \mathbb{E}[N]\right). \tag{S50}$$

**Lemma S2.10.** *If $\vec{\mathbf{X}}$ is such that $\vec{\mathbf{X}}|N \sim \text{DirMult}(\zeta_C S, N, \vec{f})$, then for all $s_1, s_2 \in [S]$ ($s_1 \neq s_2$):*

$$\text{Cov}[X_{s_1}, X_{s_2}] = f^{(s_1)}f^{(s_2)}\left(\text{Var}[N] - \frac{1}{1+\zeta_C S}\left[\zeta_C S\mathbb{E}[N] + \mathbb{E}[N^2]\right]\right). \tag{S51}$$

Observe how (using L'Hôspital's rule) equation (S51) approaches (S50) as $\zeta_C \to \infty$.

[17] As opposed to indirectly, by dividing the expression for $n!$ by $n$.

###### S2.11.1 Variance and Covariance of ghNBDM

From Lemma S2.8, for the ghNBDM distributions, for each $s \in [S]$:

$$\mathrm{Var}[N^{(s)}(0)] = f^{(s)}\lambda + \frac{[f^{(s)}(1+\zeta_D S) + (f^{(s)})^2(\zeta_C S - \zeta_D S)]}{(1+\zeta_C S)(\zeta_D S)}\lambda^2 \,. \tag{S52}$$

Note how (S52) reduces to (S56) when $\zeta_D = \zeta_C$. Similarly, (S52) approaches (S58) as $\zeta_D \to \infty$, and (S52) approaches (S60) as $\zeta_C \to \infty$, both conclusions following from L'Hôspital's rule.

From Lemma S2.10, for the ghNBDM distributions, for each $s_1, s_2 \in [S]$ ($s_1 \neq s_2$):

$$\mathrm{Cov}[N^{(s_1)}(0), N^{(s_2)}(0)] = f^{(s_1)} f^{(s_2)} \left[\frac{\zeta_C S - \zeta_D S}{(\zeta_D S)(1+\zeta_C S)}\right]\lambda^2 \,. \tag{S53}$$

Note how (S53) reduces to (S57) when $\zeta_D = \zeta_C$. Similarly, (S53) approaches (S59) as $\zeta_D \to \infty$, and (S53) approaches (S61) as $\zeta_C \to \infty$, both conclusions again following from L'Hôspital's rule. Information about special cases (hPoMu, hNBDM, hPoDM, hNBMu) is given below.

**Variance and Covariance of hPoMu** For the hPoMu distribution, for each $s \in [S]$:

$$\mathrm{Var}[N^{(s)}(0)] = f^{(s)}\lambda \,, \tag{S54}$$

and for all $s_1, s_2 \in [S]$ ($s_1 \neq s_2$):

$$\mathrm{Cov}[N^{(s_1)}(0), N^{(s_2)}(0)] = 0 \,. \tag{S55}$$

This follows either from applying Lemmas S2.7 and S2.9 directly, or recalling that the marginals of the hPoMu distribution are mutually independent $\mathrm{Pois}(f^{(s)}\lambda)$ random variables.

**Variance and Covariance of hNBDM** For the hNBDM distribution, for each $s \in [S]$:

$$\mathrm{Var}[N^{(s)}(0)] = f^{(s)}\lambda + \frac{f^{(s)}}{\zeta S}\lambda^2 \,, \tag{S56}$$

and for all $s_1, s_2 \in [S]$ ($s_1 \neq s_2$):

$$\mathrm{Cov}[N^{(s_1)}(0), N^{(s_2)}(0)] = 0. \tag{S57}$$

These follow from applying Lemmas S2.8 and S2.10 directly, or recalling that the marginals of the hNBDM distribution are mutually independent $\mathrm{NB}(\zeta S f^{(s)}, f^{(s)}\lambda)$ random variables. Notice also how (S56) approaches (S54), as $\zeta \to \infty$, as it should. Of course (S57) already equals (S55).

**Variance and Covariance of hPoDM** For the hPoDM distribution, for each $s \in [S]$:

$$\mathrm{Var}[N^{(s)}(0)] = f^{(s)}\lambda + \frac{f^{(s)}(1-f^{(s)})}{1+\zeta_C S}\lambda^2 \,, \tag{S58}$$

and for all $s_1, s_2 \in [S]$ ($s_1 \neq s_2$):

$$\mathrm{Cov}[N^{(s_1)}(0), N^{(s_2)}(0)] = -\frac{f^{(s_1)} f^{(s_2)}}{1+\zeta_C S}\lambda^2 \,. \tag{S59}$$

These follow from applying Lemmas S2.8 and S2.10. Notice also how (S58) approaches (S54), and (S59) approaches (S55), as $\zeta_C \to \infty$, as it should.

**Variance and Covariance of hNBMu** From Lemma S2.7, for the hNBMu distribution, for each $s \in [S]$:

$$\mathrm{Var}[N^{(s)}(0)] = f^{(s)}\lambda + \frac{(f^{(s)})^2}{\zeta_D S}\lambda^2 \,. \tag{S60}$$

As expected, (S60) approaches (S54) as $\zeta_D \to \infty$.

From Lemma S2.9, for the hNBMu distribution, for each $s_1, s_2 \in [S]$ ($s_1 \neq s_2$):

$$\mathrm{Cov}[N^{(s_1)}(0), N^{(s_2)}(0)] = \frac{f^{(s_1)} f^{(s_2)}}{\zeta_D S}\lambda^2 \,. \tag{S61}$$

Again, as expected, (S61) approaches (S55) as $\zeta_D \to \infty$.

### S3 COMPARISON OF MODELS FOR INITIAL FORMATION OF DROPLETS

Section S3.1 explains the motivation for everything else that follows. Section S3.2 explains some of the technical details of the comparisons. Section S3.3 explains the implementation details of the comparisons. Section S3.4 describes and explains the comparisons. Finally section S3.5 interprets the results from section S3.4, explaining what they mean for modeling the initial formation of droplets.

#### S3.1 Statistical Motivation

As explained in section S2.1, we want to find a statistical model capable of describing the initial droplet formation process in order to understand its randomness. Given two models which are both adequate for describing the initial droplet formation process, the simpler model will be easier to understand. Therefore, given two models with the same descriptive power, we should choose the simpler model. It follows that justifying the use of a more complicated model over a simpler model requires showing that the more complicated model has substantially more descriptive power than the simpler model.

Of the models introduced in section S2, hPoMu was (arguably) the simplest. Therefore, to justify potentially using any of the other, more complicated models in place of hPoMu, we must first ascertain whether any of the other models have substantially more descriptive power than hPoMu. In doing so, we will also test which failures of the assumptions motivating hPoMu have the greatest potential to affect the initial droplet formation process. Thus the goal of this section is to ascertain how well hPoMu can describe data generated by the other models. Only those models which can not always be described well by hPoMu have substantially more descriptive power than hPoMu.

#### S3.2 Preliminaries

This section gives precise mathematical definitions for the comparisons of other distributions to hPoMu. Section S3.2.1 defines the divergence used to compare distributions. Section S3.2.2 clarifies the details of the hypothesis test comparing distributions to hPoMu using this divergence. Finally section S3.2.3 presents the formulae for the log likelihood ratios comparing distributions to hPoMu.

##### *S3.2.1 Pearson Categorical Divergence*

The Pearson categorical divergence (more often referred to as the “Pearson $\chi^2$ divergence”, cf. below) of $\vec{\mathbf{v}} = (v^{(1)}, \dots, v^{(I)}) \in \mathbb{R}_{\geq 0}$ relative to $\vec{\mathbf{w}} = (w^{(1)}, \dots, w^{(I)}) \in \mathbb{R}_{\geq 0}$ is defined (Cichocki et al., 2009) as

$$\sum_{i=1}^{I} \frac{(v^{(i)} - w^{(i)})^2}{w^{(i)}}, \tag{S62}$$

where both $\vec{\mathbf{v}}$ and $\vec{\mathbf{w}}$ belong to the unit simplex (have exclusively non-negative entries which sum to 1), and $\vec{\mathbf{w}}$ belongs to the interior of the unit simplex (has all non-zero entries). Note that this definition is not symmetric, i.e. in general the divergence of $\vec{\mathbf{w}}$ relative to $\vec{\mathbf{v}}$ differs from that of $\vec{\mathbf{v}}$ relative to $\vec{\mathbf{w}}$.

Herein I refer to this as the “Pearson categorical divergence” to avoid confusion with either (a) the “Pearson divergence”, $1 - \rho_{\vec{x}\vec{y}}$, where $\rho_{\vec{x}\vec{y}}$ is the Pearson correlation of $\vec{x}$ and $\vec{y}$, and (b) the “Pearson $\chi^2$ distribution”[18], the asymptotic sampling distribution of this statistic under the null distribution of “Pearson-style” goodness of fit tests. In particular it might sound confusing to say (in section S3.3.2) that the “$\chi^2$ divergence” doesn’t have a “$\chi^2$ distribution”.

##### *S3.2.2 Global Picky Goodness of Fit Test*

The strains present in a droplet determine the microbial interactions it can help characterize. Therefore we want to group the droplets according to which strains are present in each droplet. Different models for the initial formation of droplets make different predictions about which such groups will be most common. A hypothesis test with hPoMu as the null distribution compares the predictions made by other models with those made by hPoMu. Rejecting the null hypothesis corresponds to asserting that hPoMu is unable to describe the observed data, so the true model has more “descriptive power”, cf. section S3.1.

The definition of the hypothesis test is split into several parts. Section S3.2.2 defines how the droplets are grouped according to strains for the hypothesis test. Section S3.2.2 gives probabilities for each group under the null hypothesis (hPoMu), while section S3.2.2 clarifies the expected counts. Section S3.2.2 defines the observed counts used in defining the goodness of fit test. Section S3.2.2 explains why some of the groups are redundant. Finally, section S3.2.2 combines the information from the previous sections to define the test statistic.

[18] Technically a parameterized family of distributions and not a single distribution.

**Categories** There are technically exponentially many, $2^S$, potential combinations of strains that could occur in a given droplet. All of these could be used in defining mutually exclusive categories for a goodness of fit test. However, for both reasons of computational feasibility and data availability (most combinations of three or more strains are highly unlikely and will be represented by few or no data points), herein only combinations with two or fewer strains are given distinct categories, with all combinations of three or more strains lumped into a category.

Thus the number of categories for the "global" goodness of fit test is

$$1+S+\binom{S}{2}+1\,. \tag{S63}$$

This is one category for all droplets with zero strains (droplets which are empty), $S$ categories for each of the picky control groups (droplets with exactly one strain), $\binom{S}{2}$ categories for each of the picky treatment groups (droplets with exactly two strains), and one category for all droplets with three or more ("multiple") strains. The groups being "picky" and not "gluttonous" ensures that the categories do not overlap. This makes it feasible to define a single "global" test.

Cf. section  for more details about "picky" and "gluttonous" groups.

**Probabilities** The probabilities associated with each category are

$$\begin{aligned}
p_{\mathcal{G}}^{(\emptyset)} &:= \mathbb{P}(N(0)=0) = e^{-\lambda}\,,\\
p_{\mathcal{G}}^{(s)} &:= \mathbb{P}(N^{(s)}(0)\geq 1, \forall\sigma\neq s: N^{(\sigma)}(0)=0)\\
&= \mathbb{P}(N^{(s)}(0)\geq 1)\cdot\prod_{\sigma\neq s}\mathbb{P}(N^{(\sigma)}(0)=0)\\
&= (1-e^{-f^{(s)}\lambda})\cdot e^{-(1-f^{(s)})\lambda}\,,\\
p_{\mathcal{G}}^{(s_1,s_2)} &:= \mathbb{P}(N^{(s_1)}(0)\geq 1, N^{(s_2)}(0)\geq 1, \forall\sigma\notin\{s_1,s_2\}: N^{(\sigma)}(0)=0)\\
&= \mathbb{P}(N^{(s_1)}(0)\geq 1)\cdot\mathbb{P}(N^{(s_2)}(0)\geq 1)\cdot\prod_{\sigma\notin\{s_1,s_2\}}\mathbb{P}(N^{(\sigma)}(0)=0)\\
&= (1-e^{-f^{(s_1)}\lambda})(1-e^{-f^{(s_2)}\lambda})\cdot e^{-(1-(f^{(s_1)}+f^{(s_2)}))\lambda}\,,\\
p_{\mathcal{G}}^{(3+)} &:= 1-p_{\mathcal{G}}^{(\emptyset)}-\sum_{\sigma=1}^{S}p_{\mathcal{G}}^{(\sigma)}-\sum_{\sigma_1=1}^{S}\sum_{\sigma_2>\sigma_1}p_{\mathcal{G}}^{(\sigma_1,\sigma_2)}\,.
\end{aligned} \tag{S64}$$

**Expected Counts** The expected counts equal the total number of droplets $D$ multiplied by the corresponding probabilities.

$$\begin{aligned}
M_{\mathcal{G}}^{(\emptyset)} &:= D\cdot p_{\mathcal{G}}^{(\emptyset)}\,,\\
M_{\mathcal{G}}^{(s)} &:= D\cdot p_{\mathcal{G}}^{(s)}\,,\\
M_{\mathcal{G}}^{(s_1,s_2)} &:= D\cdot p_{\mathcal{G}}^{(s_1,s_2)}\,,\\
M_{\mathcal{G}}^{(3+)} &:= D\cdot p_{\mathcal{G}}^{(3+)}\,.
\end{aligned} \tag{S65}$$

**Observed Counts** The observed counts for the global goodness of fit test are

$$\begin{aligned}
\hat{M}_{\mathcal{G}}^{(\emptyset)} &:= \sum_{d\in[D]} \mathbb{I}\{N_d(0)=0\}\,, \\
\hat{M}_{\mathcal{G}}^{(s)} &:= \sum_{d\in[D]} \mathbb{I}\left\{N_d^{(s)}(0)\geq 1\right\}\cdot\prod_{\sigma\neq s}\mathbb{I}\left\{N_d^{(\sigma)}(0)=0\right\}\,, \\
\hat{M}_{\mathcal{G}}^{(s_1,s_2)} &:= \sum_{d\in[D]} \Bigg[\mathbb{I}\left\{N_d^{(s_1)}(0)\geq 1\right\}\cdot\mathbb{I}\left\{N_d^{(s_2)}(0)\geq 1\right\} \\
&\qquad\cdot\prod_{\sigma\notin\{s_1,s_2\}}\mathbb{I}\left\{N_d^{(\sigma)}(0)=0\right\}\Bigg]\,, \\
\hat{M}_{\mathcal{G}}^{(3+)} &:= D-\hat{M}_{\mathcal{G}}^{(\emptyset)}-\sum_{\sigma=1}^{S}\hat{M}_{\mathcal{G}}^{(\sigma)}-\sum_{\sigma_1=1}^{S}\sum_{\sigma_2>\sigma_1}\hat{M}_{\mathcal{G}}^{(\sigma_1,\sigma_2)}\,.
\end{aligned} \tag{S66}$$

**Symmetry of Definitions** Observe how, for any given pair of strains $s_1, s_2$:

$$\begin{aligned}
p_{\mathcal{G}}^{(s_1,s_2)} &= p_{\mathcal{G}}^{(s_2,s_1)}\,, \\
M_{\mathcal{G}}^{(s_1,s_2)} &= M_{\mathcal{G}}^{(s_2,s_1)}\,, \\
\hat{M}_{\mathcal{G}}^{(s_1,s_2)} &= \hat{M}_{\mathcal{G}}^{(s_2,s_1)}\,.
\end{aligned} \tag{S67}$$

Thus for any given pair of strains $s_1, s_2$, only the left hand sides of (S67) are considered, avoiding redundancy and overlapping categories.

**Test Statistic** The test statistic is the Pearson categorical divergence of

$$\begin{aligned}
&D^{-1}\cdot\left(\hat{M}_{\mathcal{G}}^{(\emptyset)},\hat{M}_{\mathcal{G}}^{(1)},\dots,\hat{M}_{\mathcal{G}}^{(S)},\hat{M}_{\mathcal{G}}^{(1,2)},\dots,\hat{M}_{\mathcal{G}}^{(S-1,S)},\hat{M}_{\mathcal{G}}^{(3+)}\right) \\
&\text{relative to} \\
&D^{-1}\cdot\left(M_{\mathcal{G}}^{(\emptyset)},M_{\mathcal{G}}^{(1)},\dots,M_{\mathcal{G}}^{(S)},M_{\mathcal{G}}^{(1,2)},\dots,M_{\mathcal{G}}^{(S-1,S)},M_{\mathcal{G}}^{(3+)}\right)\,.
\end{aligned} \tag{S68}$$

We can get an asymptotic $p$-value in the standard way, via the survival function of the $\chi^2$ distribution with $\left(1+S+\binom{S}{2}+1\right)-1$ degrees of freedom.

###### *S3.2.3 Exact Log Likelihood Ratios*

Below I give explicit formulae for the log likelihood ratios w.r.t. hPoMu which do not invoke Stirling's approximation nor the generalization thereof to the $\Gamma$ function. These formulae were applied as described in section S3.3.3.

**hTPMH** The likelihood ratio of the hTPMH distribution with respect to the corresponding hPoMu distribution is

$$\mathbb{P}(\text{Pois}(\lambda)\leq K_{d-1})^{-1}\cdot\frac{(K_{d-1}-n)!}{(K_{d-1})!}\cdot\left[\prod_{s=1}^{S}\frac{(K_{d-1}^{(s)})!}{(K_{d-1}^{(s)}-n^{(s)})!}\right]\cdot\left[\prod_{s=1}^{S}(f^{(s)})^{n^{(s)}}\right]^{-1}\,. \tag{S69}$$

Therefore the log likelihood ratio is

$$\begin{aligned}
&-\log\left(\mathbb{P}(\text{Pois}(\lambda)\leq K_{d-1})\right)+\log\left((K_{d-1}-n)!\right)-\log\left((K_{d-1})!\right)+ \\
&\sum_{s=1}^{S}\left[\log\left((K_{d-1}^{(s)})!\right)-\log\left((K_{d-1}^{(s)}-n^{(s)})!\right)\right]-\sum_{s=1}^{S}n^{(s)}\log(f^{(s)})\,.
\end{aligned} \tag{S70}$$

Using the identity $\Gamma(n+1) = n!$, the above can be rewritten using the log-gamma function[19] (which is directly implemented in SciPy (Virtanen et al., 2020)):

$$-\log\left(\mathbb{P}(\text{Pois}(\lambda) \leq K_{d-1})\right) + \log\left(\Gamma(K_{d-1} - n + 1)\right) - \log\left(\Gamma(K_{d-1} + 1)\right) + \sum_{s=1}^{S}\left[\log\left(\Gamma(K_{d-1}^{(s)} + 1)\right) - \log\left(\Gamma(K_{d-1}^{(s)} - n^{(s)} + 1)\right)\right] - \sum_{s=1}^{S} n^{(s)} \log(f^{(s)}) \,. \tag{S71}$$

As an aside, because the above expression equals (S19) from section S2.5.2, one can derive an explicit formula for $\mathscr{R}_{\text{MHg}}(\vec{\mathbf{n}}, \vec{\mathbf{K}}_{d-1})$. It is not very elegant.

For all observed values of $K_{d-1}$, $\mathbb{P}(\text{Pois}(\lambda) \leq K_{d-1})^{-1}$ was indistinguishable from 1 up to numerical precision. (Indeed for $\lambda = 2$ that probability appears to be indistinguishable from 1 up to numerical precision already for $K_{d-1} \geq 20$, values many orders of magnitude smaller than any observed value.) Therefore the contribution of the logarithm of this term was treated as if it was exactly 0 in the code implementation and not explicitly included.

**hPoDM** The likelihood ratio of the hPoDM distribution with respect to the corresponding hPoMu distribution is

$$\frac{\Gamma(\zeta_C S)}{\Gamma(\zeta_C S + n)} \cdot \left[\prod_{s=1}^{S} \frac{\Gamma(\zeta_C S f^{(s)} + n^{(s)})}{\Gamma\left(\zeta_C S f^{(s)}\right)}\right] \cdot \left[\prod_{s=1}^{S} (f^{(s)})^{n^{(s)}}\right]^{-1} \,. \tag{S72}$$

Therefore the log likelihood ratio is

$$\log\left(\Gamma(\zeta_C S)\right) - \log\left(\Gamma(\zeta_C S + n)\right) + \sum_{s=1}^{S}\left[\log\left(\Gamma(\zeta_C S f^{(s)} + n^{(s)})\right) - \log\left(\Gamma\left(\zeta_C S f^{(s)}\right)\right)\right] - \sum_{s=1}^{S} n^{(s)} \log(f^{(s)}) \,. \tag{S73}$$

As an aside, because the above expression equals (S27) from section S2.6.2, one can derive an explicit formula for $\mathscr{R}_{\text{DM}}(\vec{\mathbf{n}}, \zeta_C S \vec{f})$. Again, it is not elegant.

**hNBDM** The likelihood ratio of the hNBDM distribution with respect to the corresponding hPoMu distribution is

$$\prod_{s=1}^{S}\left[\frac{\Gamma(\zeta S f^{(s)} + n^{(s)})}{\Gamma(\zeta S f^{(s)})} \cdot \frac{(\zeta S f^{(s)})^{\zeta S f^{(s)}}}{(\zeta S f^{(s)} + f^{(s)}\lambda)^{\zeta S f^{(s)} + n^{(s)}}}\right] \cdot \left[\prod_{s=1}^{S} e^{-f^{(s)}\lambda}\right]^{-1} \,. \tag{S74}$$

Therefore the log likelihood ratio is

$$\sum_{s=1}^{S}\left[\log\left(\Gamma(\zeta S f^{(s)} + n^{(s)})\right) - \log\left(\Gamma(\zeta S f^{(s)})\right) + \zeta S f^{(s)} \log\left(\zeta S f^{(s)}\right) - (\zeta S f^{(s)} + n^{(s)}) \log\left(\zeta S f^{(s)} + f^{(s)}\lambda\right)\right] + \sum_{s=1}^{S} f^{(s)} \text{[illegible]} \tag{S75}$$

As an aside, because the above expression equals (S39) from section S2.7.2, one can derive an explicit formula for $\mathscr{R}_{\text{hNBDM}}(\zeta S \vec{f}, \vec{\mathbf{n}})$. Again, it is not elegant.

#### S3.3 Methods

Section S3.3.1 explains which distributions were simulated, why, and how. Section S3.3.2 explains how categorical divergences and corresponding approximate $p$-values were computed. Section S3.3.3 explains how logarithms of the likelihood ratios were computed for five of the distributions. I used NumPy (Harris et al., 2020) version 1.20.2 and SciPy (Virtanen et al., 2020) version 1.6.2 for computations, and Matplotlib (Hunter, 2007) version 3.4.1 and/or Seaborn (Waskom, 2021) version 0.11.1 for plots. Complete implementation details can be found in the code at the relevant GitLab repository. See https://gitlab.com/krinsman/droplets.

[19]Which is most likely more numerically stable and computationally tractable than directly computing the logarithm of the factorial.

###### S3.3.1 Simulated Distributions

To test the extent to which the assumptions of hPoMu may be violated with hPoMu remaining an adequate model, I simulated seven distributions. The first distribution, the hierarchical truncated Poisson Multivariate Hypergeometric (hTPMH) accounts for sampling without replacement (section S3.3.1), removing the sampling with replacement approximation of hPoMu. The remaining six distributions removed the homogeneity assumption of hPoMu by modeling density and compositional heterogeneities via over-dispersion (sections S3.3.1 and S3.3.1). Details of the simulations common to all seven distributions are explained in section S3.3.1.

**Simulating Sampling without Replacement** I used an initial total population size of $500,000,000$ (500 million) cells for defining the parameter $K$ of hTPMH. This is substantially less than the number of cells ($\sim 10^{10}$) which would be sampled from in practice. The NumPy version of the multivariate hypergeometric distribution does not scale beyond a total population size of $10^9$.

**Simulating Compositional Heterogeneity Only** The first three of those model only compositional heterogeneity and have no density heterogeneity. The hierarchical Poisson Dirichlet-Multinomial (hPoDM) with compositional concentration $\zeta_C = 100$ has the lowest compositional heterogeneity, followed by hPoDM with $\zeta_C = 1$. The third such distribution was a hierarchical Exponential hPoDM model (hExhPoDM), with $\zeta_C$ distributed as an Exponential$(1)$ random variable.

**Simulating Compositional and Density Heterogeneities** The final three distributions model both compositional and density heterogeneity, with both assumed to correspond to equal over-dispersion. The hierarchical Negative Binomial Dirichlet-Multinomial (hNBDM) with concentration $\zeta := \zeta_C = \zeta_D = 100$ has the lowest heterogeneities, followed by hNBDM with $\zeta = 1$. The final distribution was again another even more hierarchical model, the hierarchical Exponential hNBDM model (hExhNBDM), and $\zeta$ was distributed as an Exponential$(1)$ random variable.

**Simulation Implementation Details** For all simulations and distributions, the same "simulated community" (or "SimCom") of 91 strains was used. These correspond to 90 strains distributed across 9 distinct relative abundances or population frequencies: 10 strains for each of .01%, .02%, .05%, .1%, .2%, .5%, 1%, 2%, and 5%. The 91st strain was a "remainder" strain with abundance $\approx 12\%$. All distributions also used the same value of the rate parameter, $\lambda = 2$. This equals the expected number of cells for each droplet for all distributions except[20] technically for hTPMH.

Each of the seven distributions was statistically independently simulated 500 times, with each simulation having 15 million statistically independently simulated droplets, except for hTPMH, for which the statistical dependence structure of successive droplets is inherently Markov. (This corresponds to 5 batches, each with 3 million droplets.)

###### S3.3.2 Computing $p$-values and Divergences

Using the categories and expected counts defined for the "global" goodness of fit hypothesis test of the null hPoMu using "picky" groups defined in section , I computed Pearson categorical ("$\chi^2$", cf. section S3.2.1) divergence values and $\chi^2$-approximated $p$-values for all 500 simulations of all 7 distributions.

Despite the fact that each simulation corresponded to an extremely large number of droplets, the expected counts for some categories were very small ($< 5$), making the asymptotic approximation provided by the $\chi^2$ assumption possibly inadequate. To account for this, I generated $10^9$ (one **b**illion) independent replicates of the Multinomial distribution corresponding to the number of droplets per simulation and the probabilities for each category in the goodness of fit test. From this, I generated $10^9$ replicates from the sampling distribution under the null of the Pearson categorical divergence statistic. I then used this Monte Carlo distribution of Pearson categorical divergence statistics to compute "Monte Carlo $p$-values" for all 500 simulations from all 7 distributions. Given the very large number of replicates, I expect these $p$-values to be more accurate than those from the $\chi^2$ asymptotic approximation. Nevertheless the results using the $\chi^2$ approximation were similar, cf. section S3.6.

[20] The expectation of a truncated Poisson distribution will be less than that of the corresponding Poisson distribution. In this context any deficit compared to $\lambda = 2$ is negligible in practice, because the support was always truncated to a value no smaller than $\approx 400$ million.

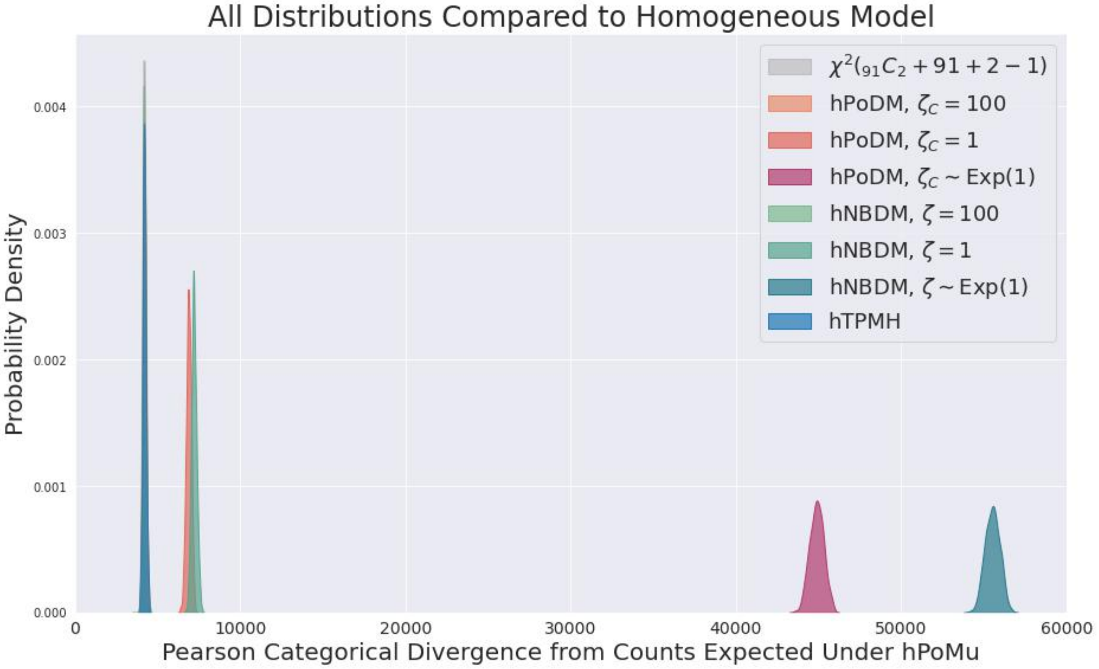

**Supplementary Figure S12**

###### *S3.3.3 Log Likelihood Ratios*

I derived explicit formulae for the logarithms of the likelihood ratios relative to hPoMu for the hTPMH, hPoDM, and hNBDM families of distributions (cf. section S3.2.3). I did not attempt to derive analogous formulae for the hExhPoDM and hExhNBDM families of distributions to avoid making mistakes computing anti-derivatives of the (logarithm of the) Gamma function.

Then, for each of the five distributions (hTPMH, hPoDM $\zeta_C = 100$, hNBDM $\zeta = 100$, hPoDM $\zeta_C = 1$, hNBDM $\zeta = 1$) with explicit formulae for the logarithm of their likelihood ratios with respect to hPoMu, of the 500 simulations I arbitrarily chose one (the 18th) and evaluated the respective formula on all 15 million droplets. For each of the five distributions, I also collected the results of doing so into gluttonous groups[21] (which by definition overlap) and computed the geometric mean of the observed likelihood ratios for all groups.

##### S3.4 Results

Section S3.4.1 discusses evidence indicating that sampling with replacement is the most innocuous of the assumptions behind hPoMu. Section S3.4.2 discusses evidence indicating that ghNBDM distributions with low (but nonzero) heterogeneity may be adequately modeled by hPoMu. Section S3.4.3 discusses evidence indicating that ghNBDM distributions with only moderate heterogeneity can be easily distinguished from hPoMu. Finally, section S3.4.4 discusses evidence indicating that even further heterogeneity can make differences from hPoMu extremely obvious.

###### *S3.4.1 Sampling without Replacement*

All available evidence suggests that, even after $15,000,000$ droplets have been formed, the hTPMH distribution for the chosen value of $K$ is extremely similar to the corresponding hPoMu distribution.

**Goodness of Fit $p$-Values Are Approximately Uniformly Distributed** For the hTPMH distribution, figure S13 shows how the distribution of (Monte Carlo) $p$-values over 500 simulations is approximately uniform, as it would be if we had actually sampled from the true hPoMu null distribution instead.

**Distribution of Pearson Categorical Divergences for Sampling without Replacement** Moreover, as seen clearly in figure S14 (and less so in figure S12), the observed distribution of Pearson categorical

[21] See section S4.3.2 for clarification of the specific meaning of the term "gluttonous groups". See also footnote 24 of section S4.3.2 regarding the groups defining the diagonals of the heatmaps.

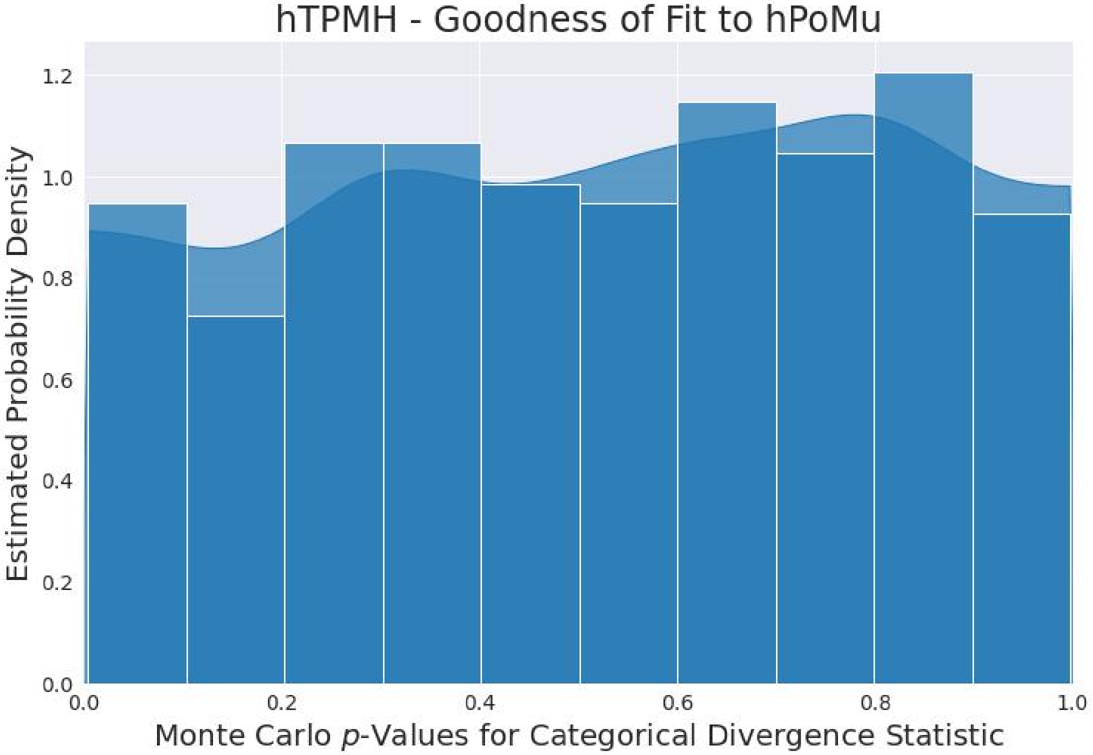

**Supplementary Figure S13**

divergences from the hTPMH distribution overlaps strongly with approximately what would have been expected under the hPoMu null distribution.

**Distribution of Likelihood Ratios w.r.t. hPoMu** When plotted on the same scale as that used for the other distributions, the histogram for hTPMH of observed likelihood ratios relative to hPoMu seen in figure S15 has no right tail whatsoever. This indicates few, if any, of the droplets were substantially "better explained" using the hTPMH distribution compared to using hPoMu.

**Typical Likelihood Ratios as a Function of Strain Frequency** The heat map of typical values for hTPMH of the likelihood ratio across gluttonous groups showin in figure S16 also shows, even at worst, negligible differences from hPoMu. The values are slightly smaller for the least abundant strains, reflecting how the sampling with approximation is weakest for those strains. Even so, the values are still very close to 1 and well within the range observed for other distributions.

##### *S3.4.2 Very Similar Distributions*

All available evidence suggests that the hPoDM with $\zeta_C = 100$ and hNBDM with $\zeta = 100$ distributions are very similar to the corresponding hPoMu distribution. Nevertheless, they also appear to differ slightly more than the hTPMH distribution.

**Goodness of Fit $p$-Values Are Roughly Uniformly Distributed** For the hPoDM distribution with $\zeta_C = 100$ and the hNBDM distribution with $\zeta = 100$, figures S17 and S18 show respectively how their distributions of (Monte Carlo) $p$-values over 500 simulations is roughly similar to the uniform distribution.

The $p$-value distribution for hNBDM with $\zeta = 100$ in figure S18 appears to fit the uniform distribution worse than the corresponding distribution for hPoDM with $\zeta_C = 100$ in figure S17. This could possibly be a reflection of how the hNBDM family incorporates both kind of heterogeneities whereas the hPoDM family does not. However, it could also possibly be a reflection merely of the relatively small ($n = 500$) sample size.

**Distributions of Pearson Categorical Divergences for Very Similar Distributions** Furthermore, figure S14 indicates how the distribution of their Pearson categorical divergence statistics also overlaps well

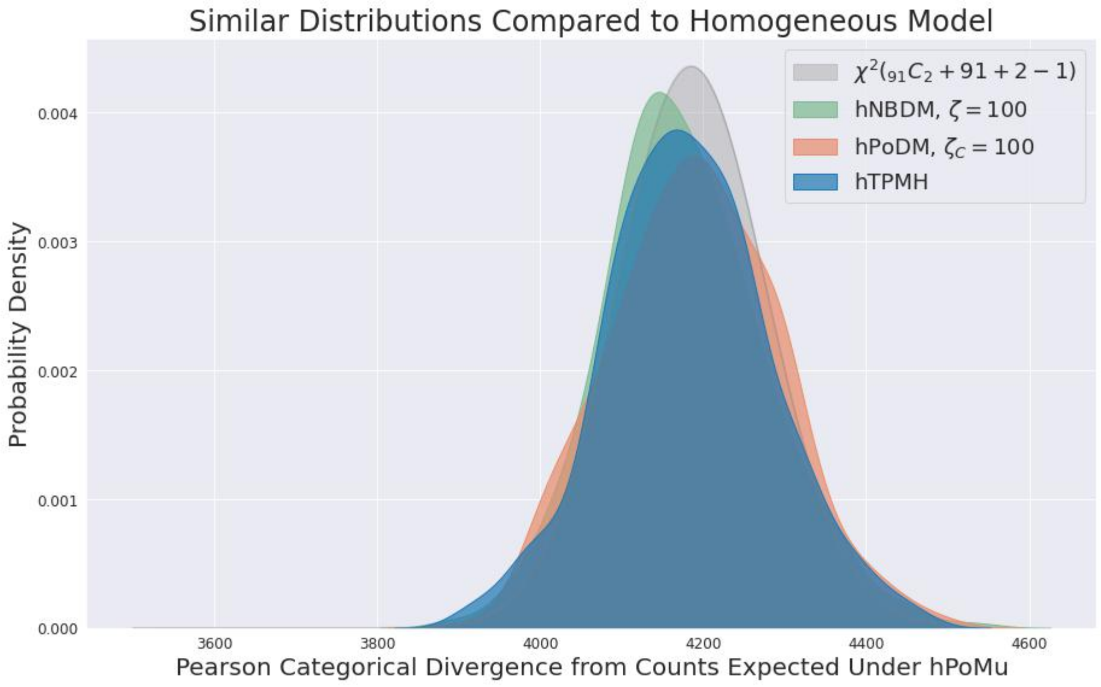

**Supplementary Figure S14**

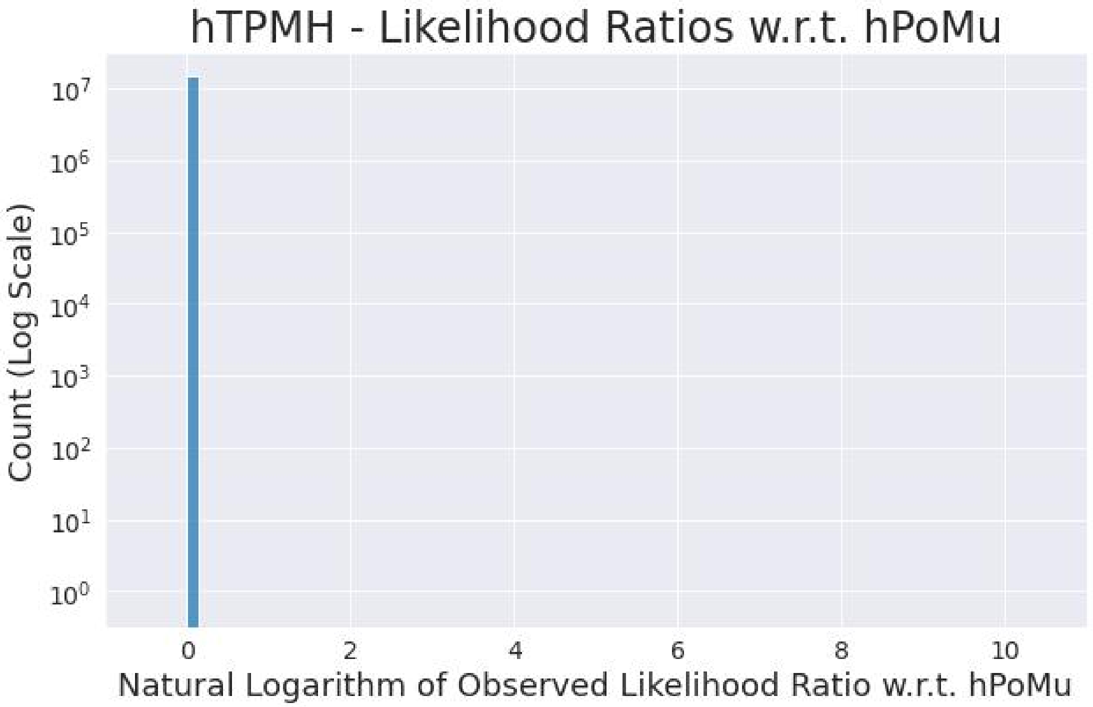

**Supplementary Figure S15**

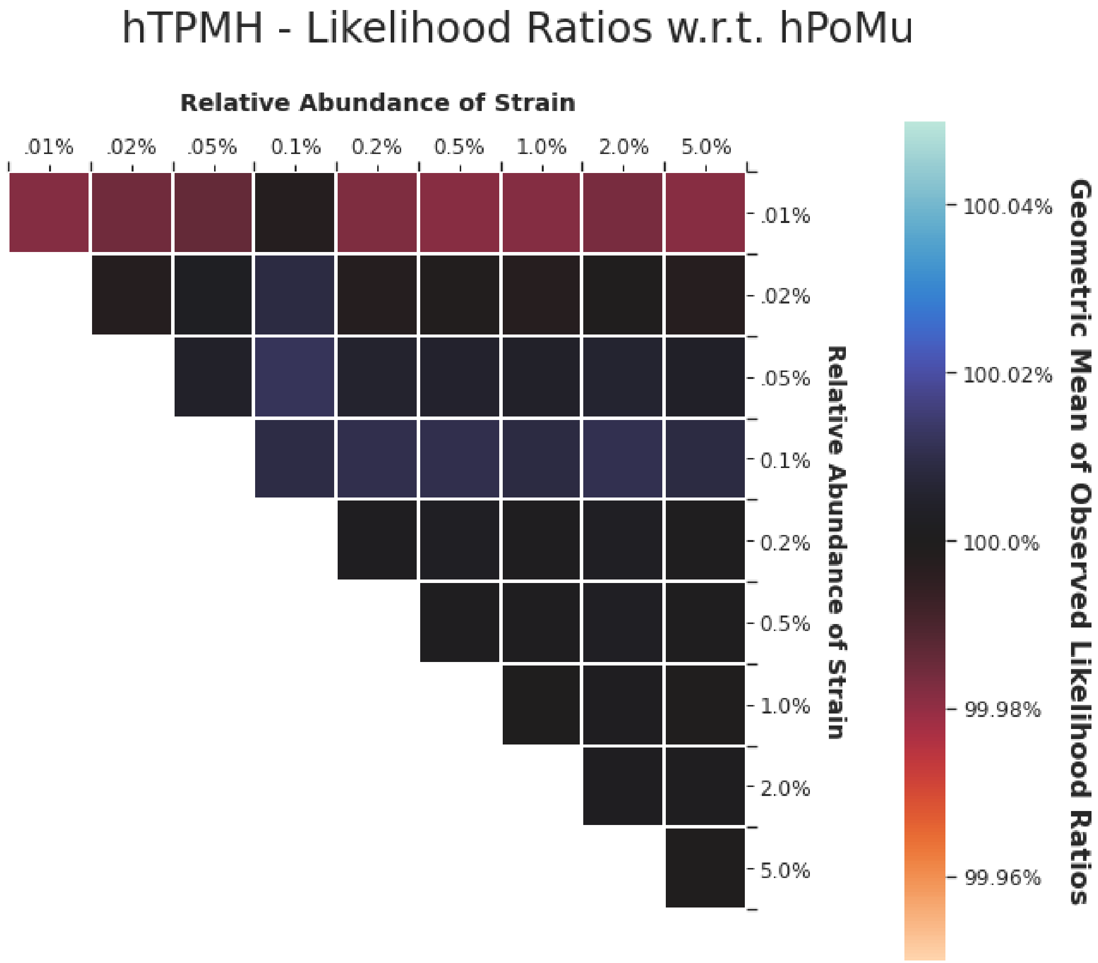

**Supplementary Figure S16**

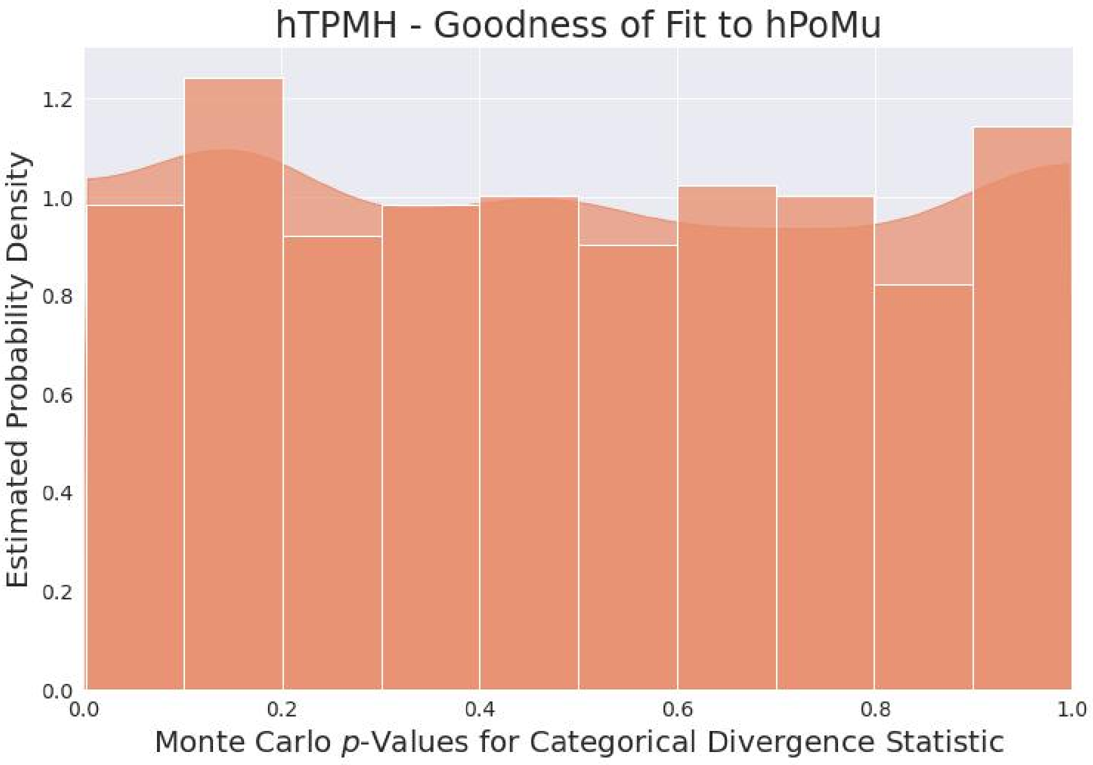

**Supplementary Figure S17**

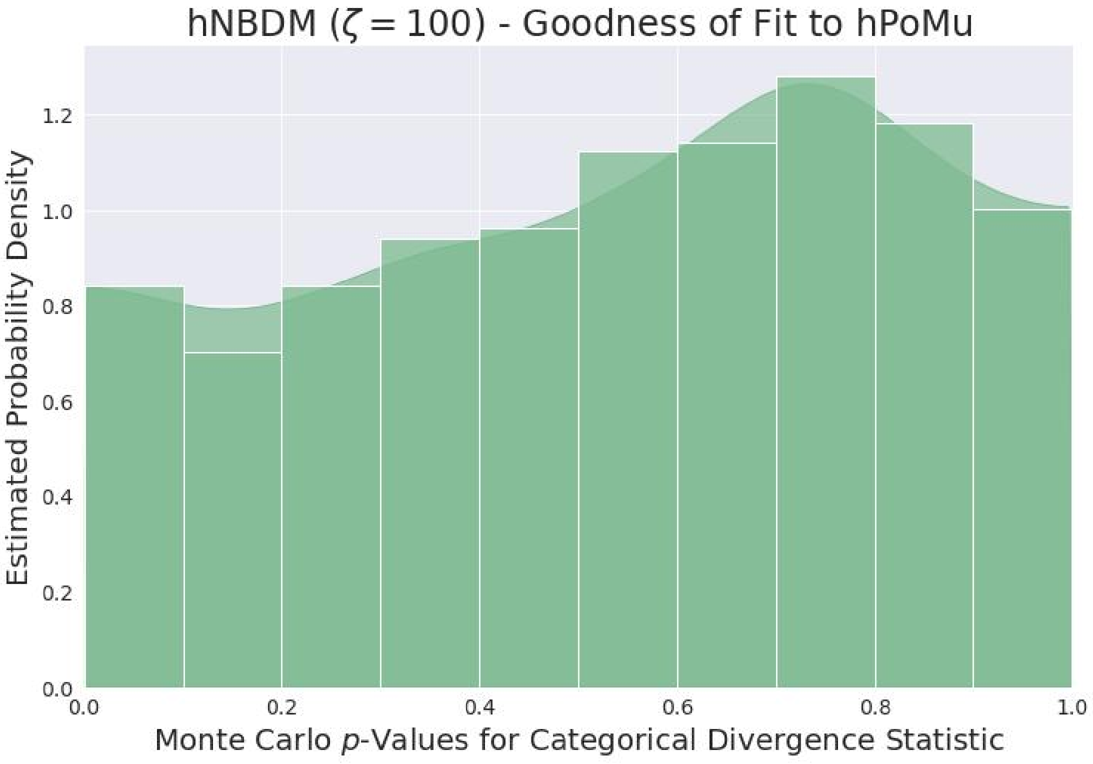

**Supplementary Figure S18**

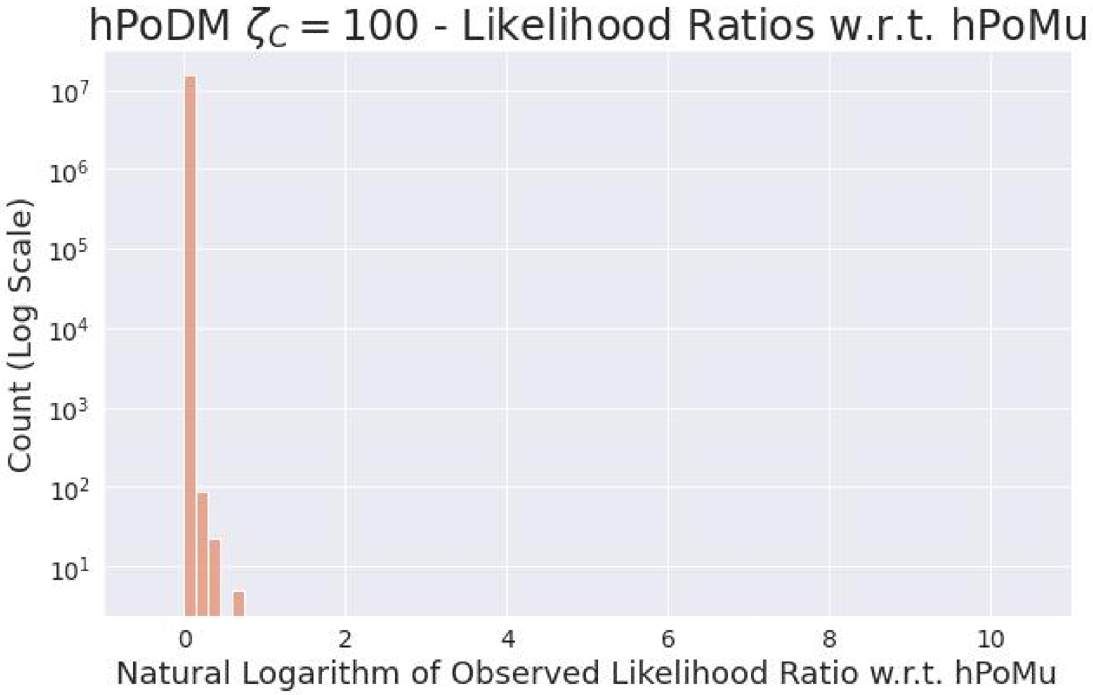

**Supplementary Figure S19**

with roughly what would be expected under the null distribution.

The overlap for hNBDM $\zeta = 100$ seen in figure S14 does not seem to be substantially worse (nor better) than that observed for hPoDM $\zeta_C = 100$, in contrast to the difference between the two distributions which might have been anticipated after comparing figures S17 and S18.

**Distributions of Likelihood Ratios w.r.t. hPoMu** Unlike what occurs for hTPMH, figures S19 and S20 respectively do show the distributions of observed likelihood ratios for hPoDM with $\zeta_C = 100$ and hNBDM with $\zeta = 100$ having right tails. However, these right tails are small.

**Typical Likelihood Ratios as a Function of Strain Frequency** Similarly, while figures S21 and S22 also show greater differences relative to hPoMu compared to those observed for the hTPMH distribution, the differences still trend very small. (That the geometric means across the gluttonous groups tend to be slightly *less* than 1, rather than slightly greater, most likely reflects, in addition to the overlapping nature of the gluttonous groups, the depletion of droplets with 3 or more strains relative to hPoMu that will be observed in these distributions in section S4.)

###### *S3.4.3 Slightly Different Distributions*

All available evidence suggests that the hPoDM with $\zeta_C = 1$ and hNBDM with $\zeta = 1$ distributions are (at least) slightly different from the corresponding hPoMu distribution.

**Goodness of Fit $p$-Values Are All Effectively Zero** For the hPoDM distribution with $\zeta_C = 1$ and the hNBDM distribution with $\zeta = 1$, all of the approximate $p$-values were indistinguishable from 0 up to numerical precision. Therefore their distributions of approximate $p$-values, being a “single point spike” or “Dirac delta”, were not plotted.

**Distributions of Pearson Categorical Divergences for Slightly Different Distributions** The reason behind these low $p$-values is fairly obvious from either figure S12 or S23: their distributions of observed Pearson categorical divergences overlap almost not at all with what would be anticipated under the hPoMu null distribution. Also, the distribution of Pearson categorical divergences for hNBDM with $\zeta = 1$ is slightly further to the right than that of hPoDM with $\zeta_C = 1$, reflecting how the former incorporates both density and compositional heterogeneity while the latter incorporates only compositional heterogeneity.

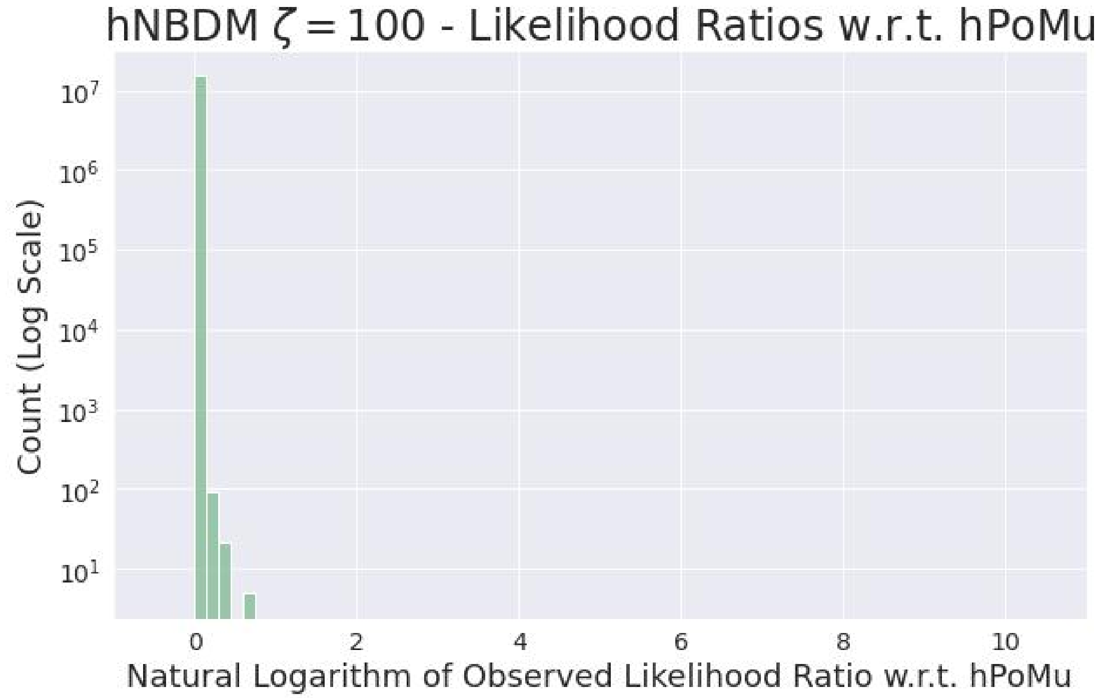

**Supplementary Figure S20**

**Distributions of Likelihood Ratios w.r.t. hPoMu** Figures S24 and S25 show very large right tails for the distributions of observed likelihood ratios for both the hPoDM distribution with $\zeta_C = 1$ and the hNBDM distribution with $\zeta = 1$. This shows that many droplets were substantially "better explained" using the true simulation distribution rather than hPoMu, something which did not occur for any of the less heterogeneous distributions. It is unclear whether either of these two large right tails is "larger" than the other.

**Typical Likelihood Ratios as a Function of Strain Frequency** Finally, figures S26 and S27 show typical values for the likelihood ratios in each gluttonous group for hPoDM $\zeta_C = 1$ and hNBDM $\zeta = 1$ that consistently "float above" 1, in clear contrast to the "hovering around 1" trend observed in figures S16, S21, and S22 that correspond to less heterogeneous distributions. One also sees that the values in figure S27 trend slightly higher than those in figure S26, reflecting again how the hNBDM family of distributions incorporates both kind of heterogeneities whereas the hPoDM family does not.

###### *S3.4.4 Very Different Distributions*

All available evidence suggests that the hExhPoDM with $\mathbb{E}[\zeta_C] = 1$ and hExhNBDM with $\mathbb{E}[\zeta] = 1$ distributions are very different from the corresponding hPoMu distribution.

**Goodness of Fit $p$-Values Are All Effectively Zero** For the hExhPoDM with $\mathbb{E}[\zeta_C] = 1$ and hExhNBDM with $\mathbb{E}[\zeta] = 1$ distributions, all of the (approximate) $p$-values were again indistinguishable from 0 up to numerical precision, as was also the case for hPoDM $\zeta_C = 1$ and hNBDM $\zeta = 1$. The approximate $p$-values were again not plotted for the same reasons as for hPoDM $\zeta_C = 1$ and hNBDM $\zeta = 1$.

**Distributions of Pearson Categorical Divergences for Very Different Distributions** The reason for these small $p$-values is also the same as for hPoDM $\zeta_C = 1$ and hNBDM $\zeta = 1$: as can be seen clearly in either figure S12 or figure S28, the peaks of their distributions of Pearson categorical divergences completely fail to overlap with the range of values that would be anticipated under the hPoMu null distribution. Both peaks are also *much* further to the right than anything observed for any of the other five distributions, indicating that the hExhPoDM $\mathbb{E}[\zeta_C] = 1$ and hExhNBDM $\mathbb{E}[\zeta] = 1$ distributions are (by far) the most heterogeneous of the seven distributions simulated.

Analogous to what occurs for hPoDM $\zeta_C = 1$ and hNBDM $\zeta = 1$, the peak for hExhNBDM $\mathbb{E}[\zeta] = 1$

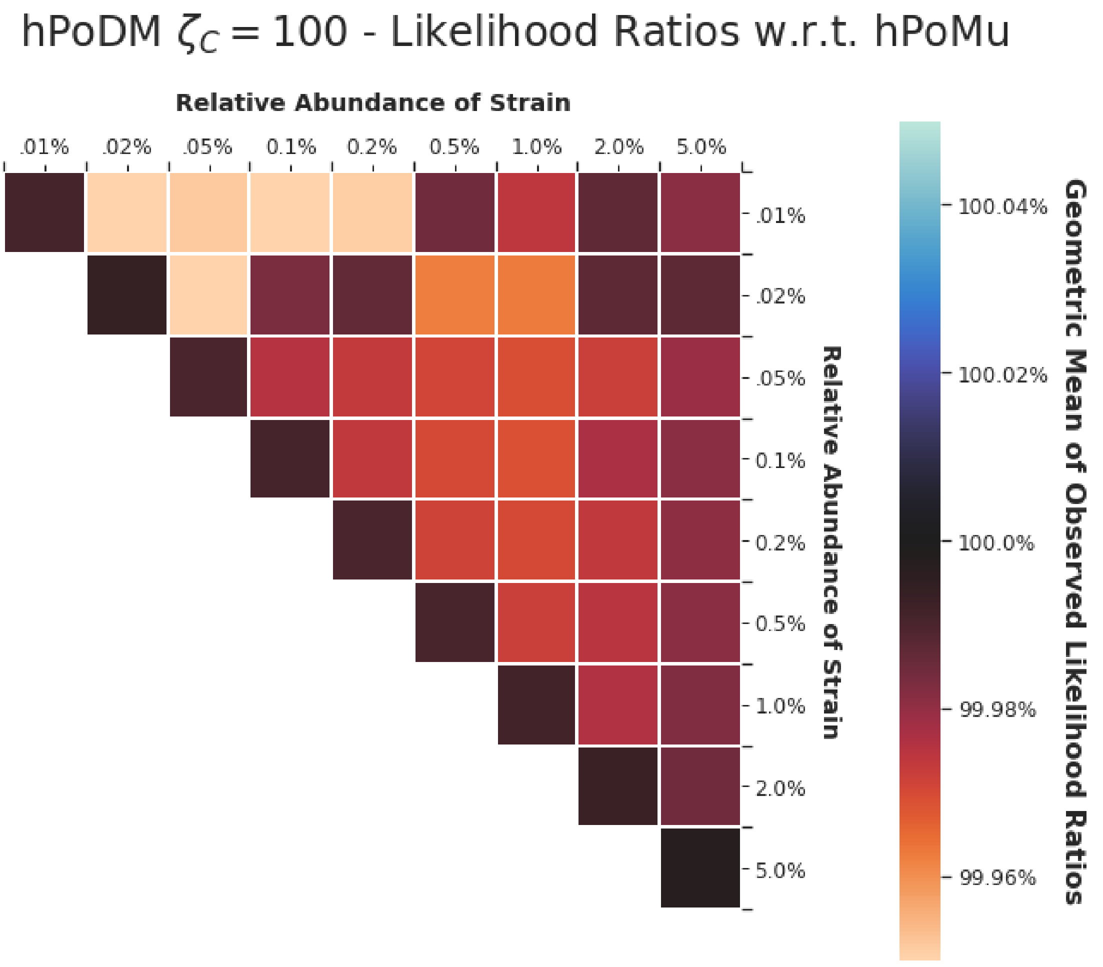

Supplementary Figure S21

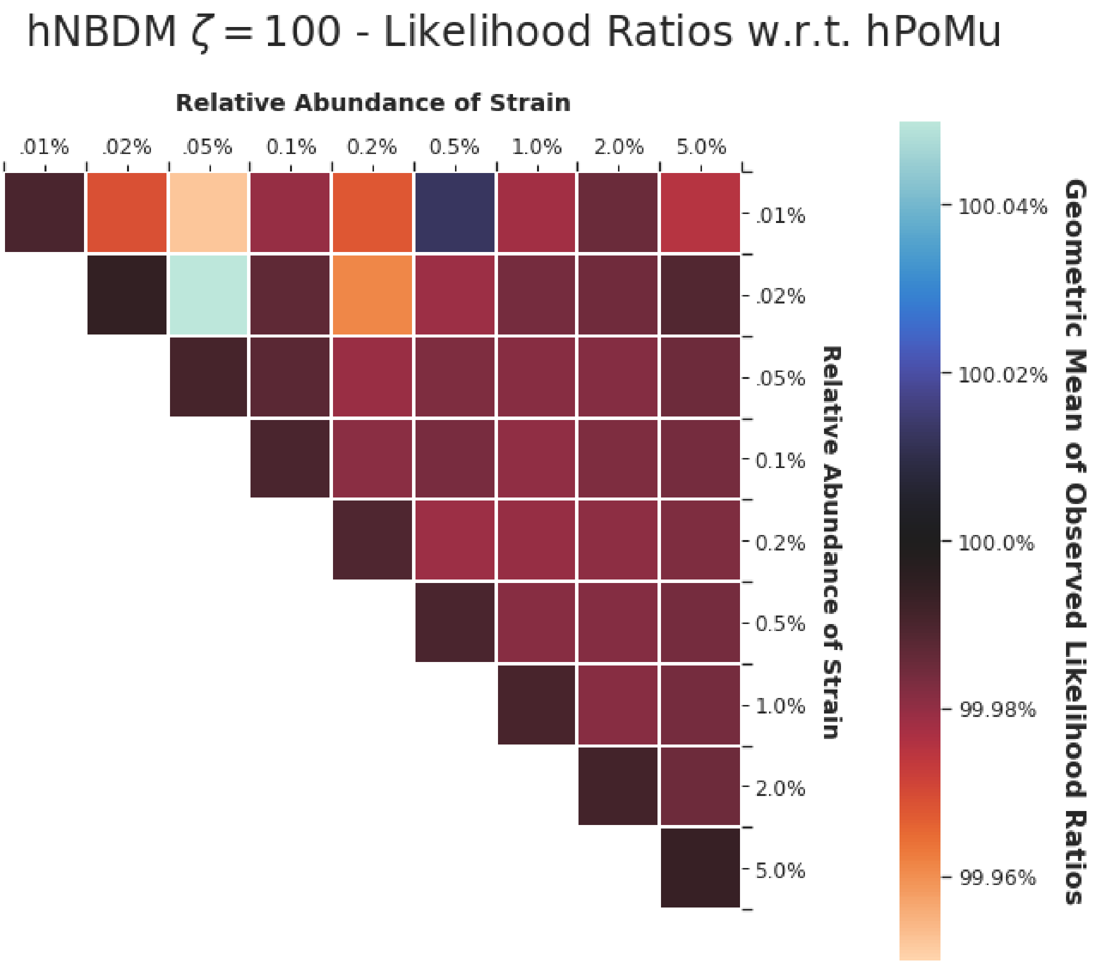

**Supplementary Figure S22**

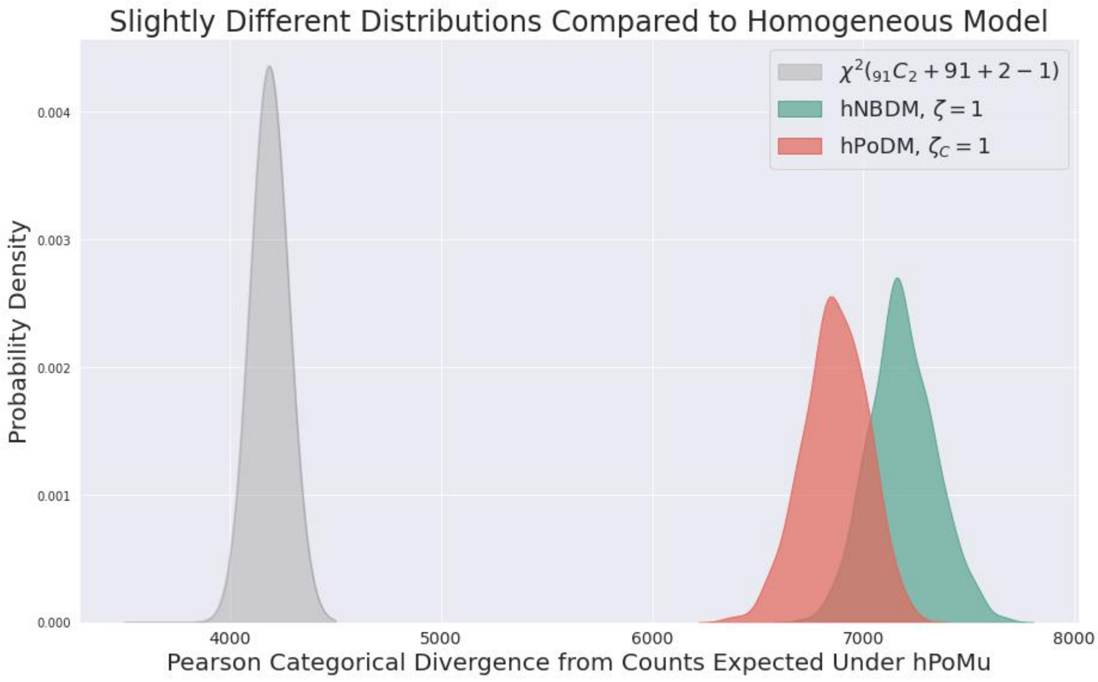

**Supplementary Figure S23**

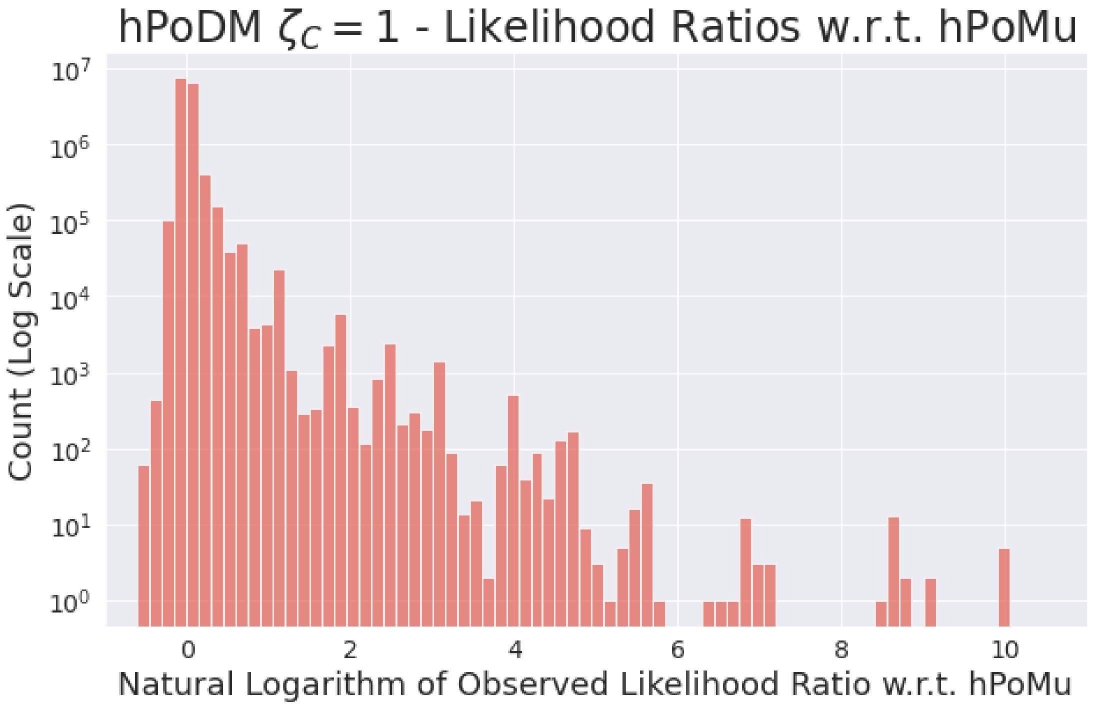

**Supplementary Figure S24**

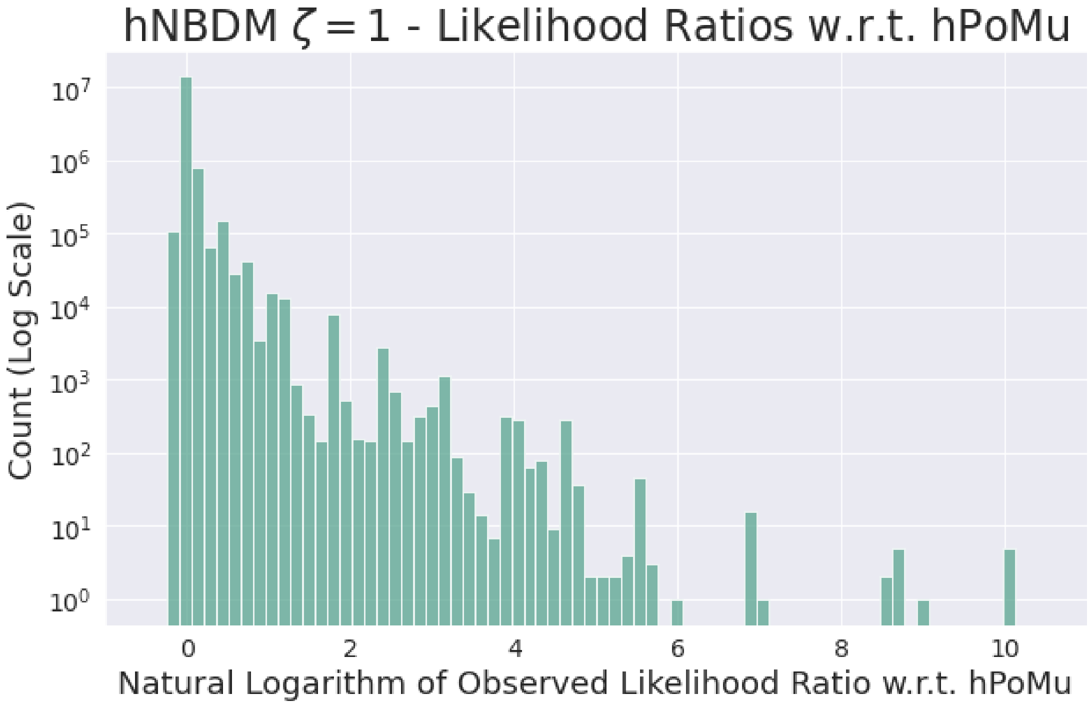

**Supplementary Figure S25**

is further to the right than the peak for hExhPoDM $\mathbb{E}[\zeta_C] = 1$, reflecting how the former incorporates both kind of heterogeneities whereas the latter does not. (Thus hExhNBDM $\mathbb{E}[\zeta] = 1$ can be considered the single most heterogeneous distribution of the seven that were simulated.) However, in contrast to the situation observed for hPoDM $\zeta_C = 1$ and hNBDM $\zeta = 1$, the peaks for hExhNBDM $\mathbb{E}[\zeta] = 1$ and hExhPoDM $\mathbb{E}[\zeta_C] = 1$ do not overlap at all, with the peak for hExhNBDM $\mathbb{E}[\zeta] = 1$ being greatly further to the right than that for hExhPoDM $\mathbb{E}[\zeta_C] = 1$.

#### S3.5 Discussion

Section S3.5.1 discusses the taxonomy of four groups of distributions identified in section S3.4. Section S3.5.2 discusses the two most similar groups, which are either very difficult or impossible to distinguish from hPoMu. Section S3.5.3 discusses the distributions which can be easily distinguished from hPoMu, but whose differences from hPoMu may not be *practically* important. Section S3.5.4 discusses the distributions which are most likely to exhibit differences from hPoMu that are important in practice. Section S3.5.5 reviews the next steps to ascertain which differences from hPoMu might affect our practical goal of being able to infer microbial interactions. Section S3.5.6 overviews the contributions made in this section.

##### *S3.5.1 Overview of Groups of Distributions*

The seven simulated distributions can be grouped into roughly four levels of assumption violation.

For the first two levels, the first consisting of hTPMH and the second consisting of hPoDM $\zeta_C = 100$ and hNBDM $\zeta = 100$, the data produced is difficult to distinguish from data produced by the hPoMu distribution. Therefore hPoMu is most likely an adequate model, and not an overly aggressive approximation, for data generated by distributions from these two levels.

The third level consists of hPoDM $\zeta_C = 1$ and hNBDM $\zeta = 1$, which produce data that can be relatively easily distinguished from data produced by hPoMu. It is nevertheless a priori unclear whether hPoMu is necessarily inadequate as a model for data generated by distributions from this level.

The fourth level consists of hExhPoDM $\mathbb{E}[\zeta_C] = 1$ and hExhNBDM $\mathbb{E}[\zeta] = 1$, which produce data so substantially different from that produced by hPoMu that it is almost impossible to *not* distinguish. It is very difficult to believe that hPoMu could serve as an adequate model for such data.

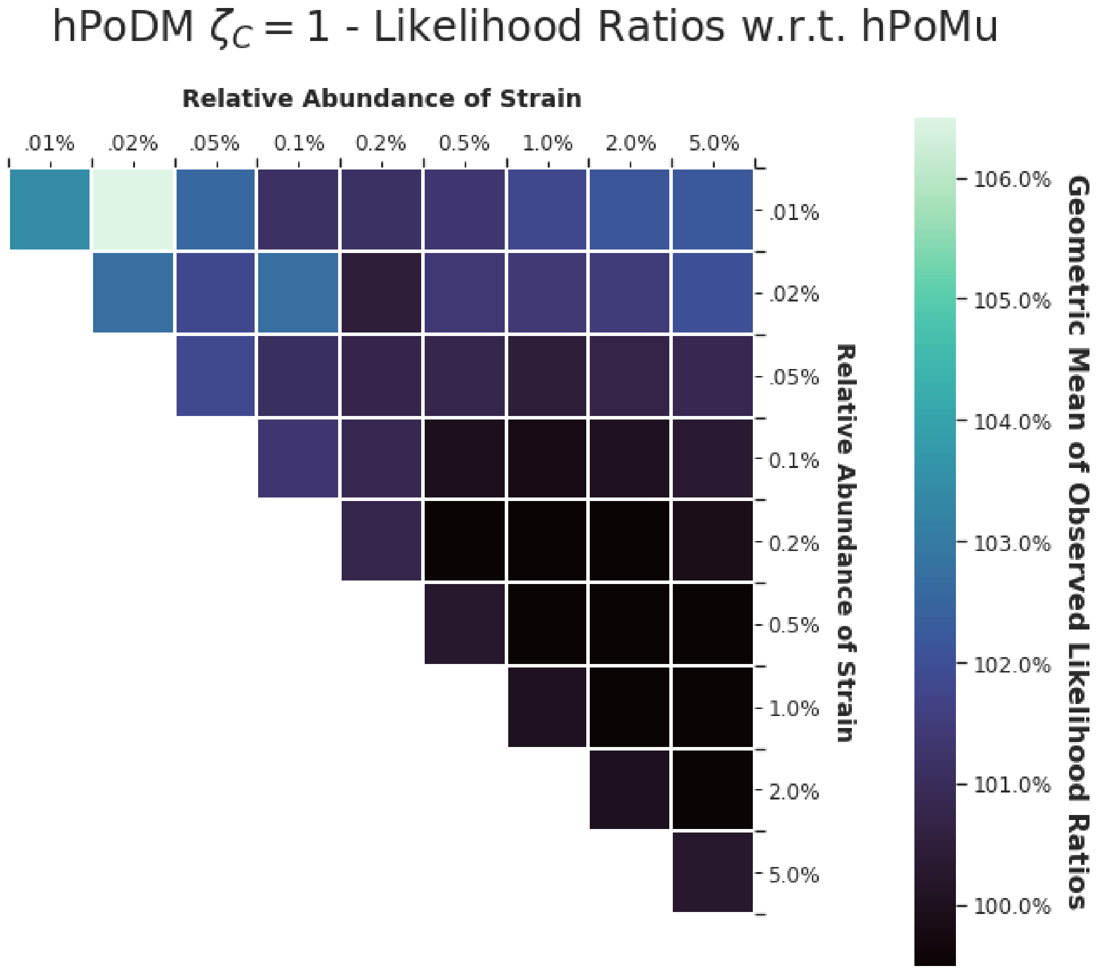

**Supplementary Figure S26**

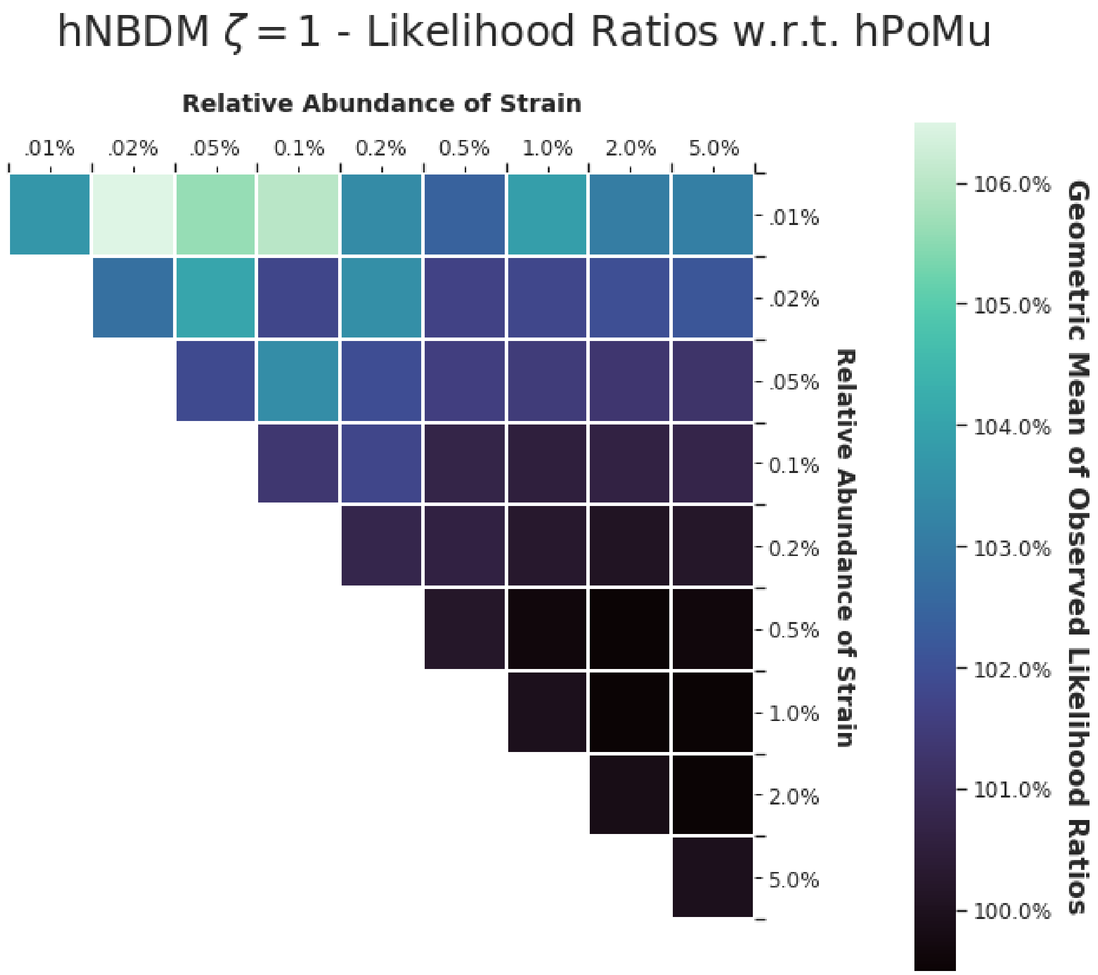

**Supplementary Figure S27**

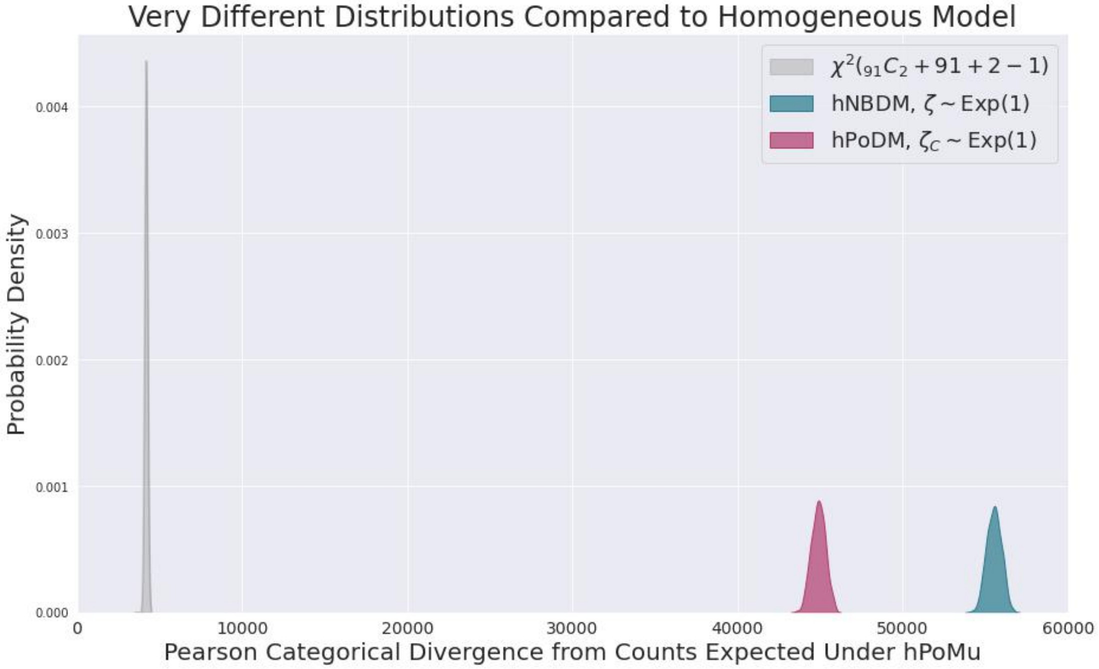

**Supplementary Figure S28**

In any case, we require both additional precision regarding what we mean by "adequately model", as well as additional evidence, to substantiate these claims.

##### *S3.5.2 Very Similar Distributions*

While both the level of consisting of hTPMH and the level consisting of hPoDM $\zeta_C = 100$ and hNBDM $\zeta = 100$ are very similar to, and difficult to distinguish from, hPoMu, the above results tended to indicate that hTPMH was always even more similar than the other two.

It is worth noting that the simulations actually *overstate* how different data generated by hTPMH should be in practice from data generated by hPoMu. I showed in section S2.5.2 that hTPMH converges in distribution to hPoMu as the size of the total population of cells being sampled from increases. While the simulation assumed a population size of 500 **m**illion, or $5 \times 10^8$, cells, a real MOREI experiment is likely to sample from a population containing upwards of 10 **b**illion, or $1 \times 10^{10}$, cells, more than twenty times as many.

Thus, to the extent we consider that data generated by hPoDM $\zeta_C = 100$ and hNBDM $\zeta = 100$ are difficult to distinguish from data generated by hPoMu, data that would be generated by hTPMH in practice should be considered effectively indistinguishable from data generated by hPoMu. In particular, even for extremely low abundance strains, these results show that the sampling with replacement approximation is completely irrelevant in practice to accurate modeling of the data generated by MOREI.

##### *S3.5.3 Slightly Different Distributions*

The above results clearly show that there are "*statistically* significant" differences between data generated by distributions in the third level, hPoDM $\zeta_C = 1$ and hNBDM $\zeta = 1$, and data generated by hPoMu. However, this still is not enough to be certain that there are *practically* significant differences. This is true even when setting aside temporarily the need to be more precise about what would constitute a "practically significant" difference making hPoMu an inadequate model.

For example, the long right tails from figures S24 and S25 are only easily visible when plotting the number of counts on a log scale. The total fraction of droplets represented by those long right tails is actually relatively small. A priori it's unclear whether the existence of a small faction of droplets which are *much* better explained by the true models than by hPoMu is actually an important demerit against hPoMu when the (vast) majority of droplets are nearly equally well explained by both. Focusing on "typical" values of the likelihood ratios as depicted in figures S26 and S27, the advantage of the true models over hPoMu is usually *at most* around 6%.

On the other hand, the "typical" likelihood ratio values in figures figures S26 and S27 also show a distinct upwards trend as the abundance of one of the involved strains decreases. Quantifying the extent of MOREI's ability to advance the state of the art requires focusing on the *least* abundant strains. Therefore, inasmuch as the discrepancies of these models compared to hPoMu may be particularly important for the least abundant strains, the discrepancies may also be important overall.

Thus the indicators from the results in this section seem to point in both directions about whether hPoMu remains an adequate model for data generated by the third level of distributions.

###### S3.5.4 Very Different Distributions

The simulated hExhPoDM and hExhNBDM distributions technically do not belong to the ghNBDM family of distributions, since their concentration "parameters" do not have fixed values, and instead are exponentially distributed. This distinction will be particularly relevant in sections S5.4.2 and S5.4.3.

Moreover, although the mean of these exponential distributions is 1, the behavior of the simulated hExhPoDM with $\mathbb{E}[\zeta_C] = 1$ and hExhNBDM with $\mathbb{E}[\zeta] = 1$ distributions is substantially different from that of the hPoDM $\zeta_C = 1$ and hNBDM $\zeta = 1$ distributions. This is most likely because concentration parameter values sampled from the Exponential$(1)$ are $1 - \frac{1}{e} \approx 63.2\%$ of the time smaller than 1. (The median of the distribution is $\ln(2) \approx .693 < 1$.)

Thus the simulated hExhPoDM with $\mathbb{E}[\zeta_C] = 1$ and hExhNBDM with $\mathbb{E}[\zeta] = 1$ can be assumed a priori to resemble more closely hPoDM and hNBDM distributions with values of $\zeta_C$ and $\zeta$ (substantially) smaller than 1. This corresponds to what we see in practice, because heterogeneities, as measured by over-dispersion, are inversely proportional to the concentration parameters. So for example $\zeta_C = \ln(2)$ would correspond to an hPoDM distribution with $\approx 44\%$ more heterogeneity than the hPoDM $\zeta_C = 1$ distribution.

Considering that the simulated hExhPoDM with $\mathbb{E}[\zeta_C] = 1$ and hExhNBDM with $\mathbb{E}[\zeta] = 1$ correspond $\approx 63.2\%$ to distributions with heterogeneities no less than those of hPoDM $\zeta_C = 1$ and hNBDM $\zeta = 1$, and $1 - \frac{1}{\sqrt{e}} \approx 39.3\%$ of the time to distributions with heterogeneities no less than *twice* those of hPoDM $\zeta_C = 1$ and hNBDM $\zeta = 1$, it is unsurprising that the aggregate/average heterogeneities observed for these distributions is much larger than those observed for either the hPoDM $\zeta_C = 1$ or hNBDM $\zeta = 1$ distributions.

Therefore fact that data generated by these two distributions is very unlikely to be adequately modeled by hPoMu further suggests that data generated by hPoDM or hNBDM distributions with $\zeta_C$ or $\zeta$ values not much smaller than 1 will also be inadequately modeled by hPoMu. It is a priori unclear which, if any, values of the concentration parameters smaller than 1 are realistic in practice.

###### S3.5.5 Next Steps

Although these results do suggest that data generated by certain distributions is unlikely to be adequately modeled by hPoMu, we need to be more precise to substantiate such a claim. We need to decide precisely which aspects of the data we want to be model accurately, or otherwise we have no way to measure whether hPoMu is an adequate model for the data. Only experiments measuring those aspects of the data will give results which could definitively determine whether or when hPoMu is an adequate model. We do this in section S4.

Moreover, even if hPoMu is inaccurate for that desired aspect of the data, it could still be an adequate model for the data as long as its predictions are reserved compared to the truth, rather than overly optimistic. To make that assessment, we need to know what values of heterogeneity might be realistic in practice. That in turn requires having estimators for the heterogeneities to use on empirical data. We fill that gap later in section S5.

###### S3.5.6 Conclusion

In this section we showed that the sampling with replacement assumption of hPoMu is completely harmless for adequately modeling the data. We also showed that, depending on what aspect of the data we wish to model, and what are realistic values of heterogeneities in practice, that the homogeneity assumptions of hPoMu could potentially inadequately model for the data. Therefore we identified density and compositional heterogeneities as the only failure of the hPoMu assumptions with the potential to be important in practice.

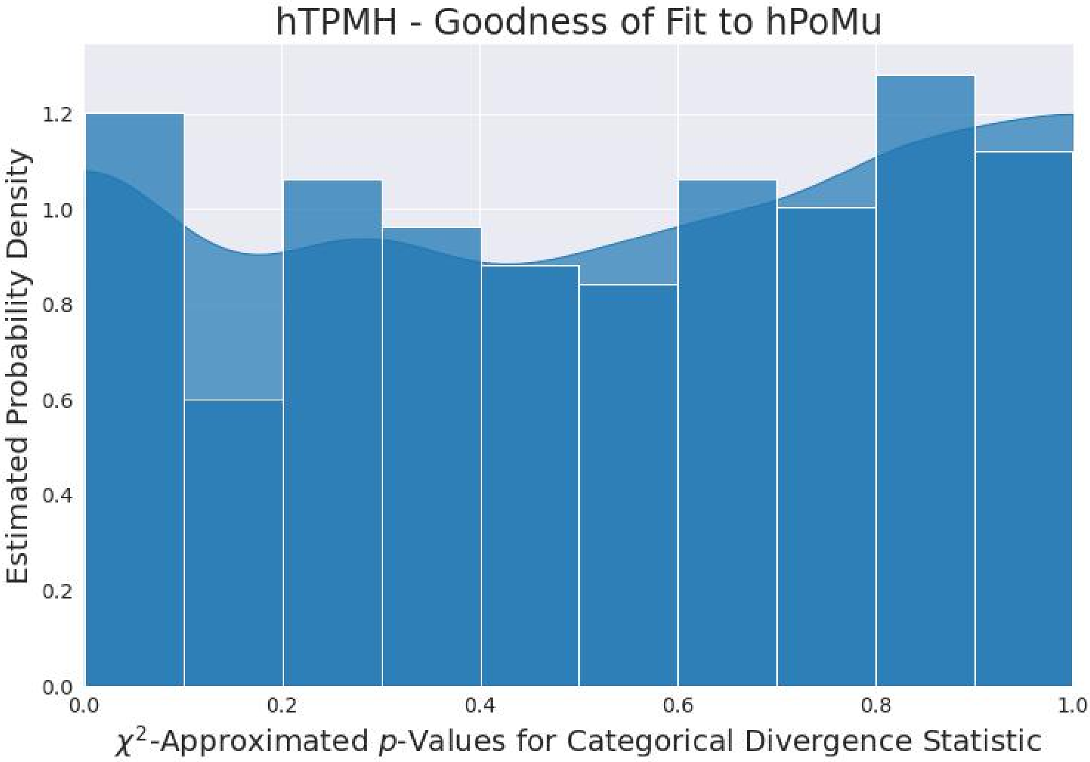

**Supplementary Figure S29**

##### S3.6 $p$-Values from $\chi^2$ Approximation

Even for 15 million droplets, some categories corresponding to the rarest strains had very small expected counts ($< 5$). Thus the $\chi^2$ approximation for the sampling distribution of the categorical divergence under the null distribution was unlikely to be accurate. In contrast, the Monte Carlo approximation to the null distribution is from $10^9$ (one **b**illion) replicates. Therefore the $p$-values from the Monte Carlo approximation are more likely to be representative of the true $p$-values. For completeness, here I report the $p$-value distributions computed from the $\chi^2$ approximation. While similar to the $p$-values from the Monte Carlo approximation, they appear to over-estimate the differences from the null.

##### S3.7 Additional Pearson Categorical Divergence Distributions

Here are two additional plots, one focusing on those distributions with compositional heterogeneity only, and the other on those distributions with both heterogeneities.

#### S4 EFFECTS OF HETEROGENEITIES ON DATA THROUGHPUT

Section S4.1 explains the motivation for the analyses performed in this section. Section S4.2 explains some of the technical details of the analyses. Section S4.3 explains implementation details of the analyses. Section S4.4 explains the results of the analyses. Section S4.5 interprets the results and explains why they are relevant to modeling the initial formation of droplets.

##### S4.1 Statistical Motivation

We want to restore control to the scientist about which questions they can answer using MOREI data by accounting for the effects caused by the randomness of the initial droplet formation process. The main effect we care about in this regard is the data throughput of the experiment in terms of the number of droplets that can potentially inform about microbial interactions. Without enough such droplets, even the best possible estimation methods might be sufficiently starved of data to make any useful inferences about microbial interactions, or even any inferences at all. As long as the average number of cells per droplet is not too large, in general the number of droplets available to serve as controls will be much larger than the

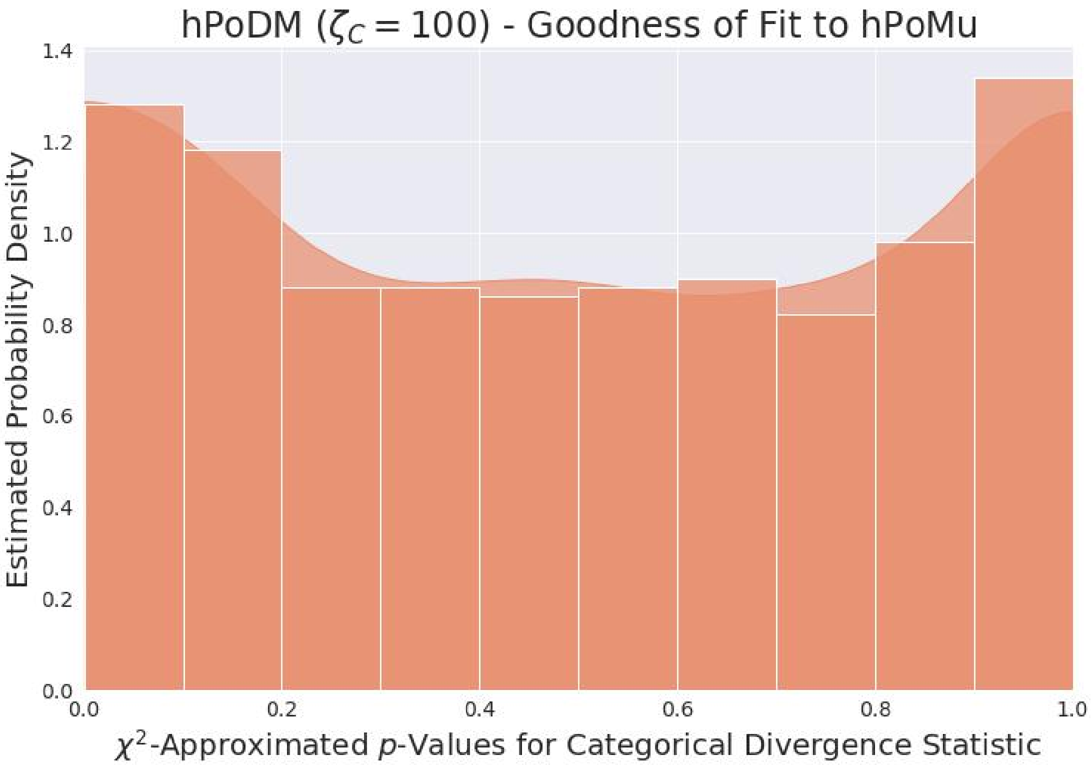

**Supplementary Figure S30**

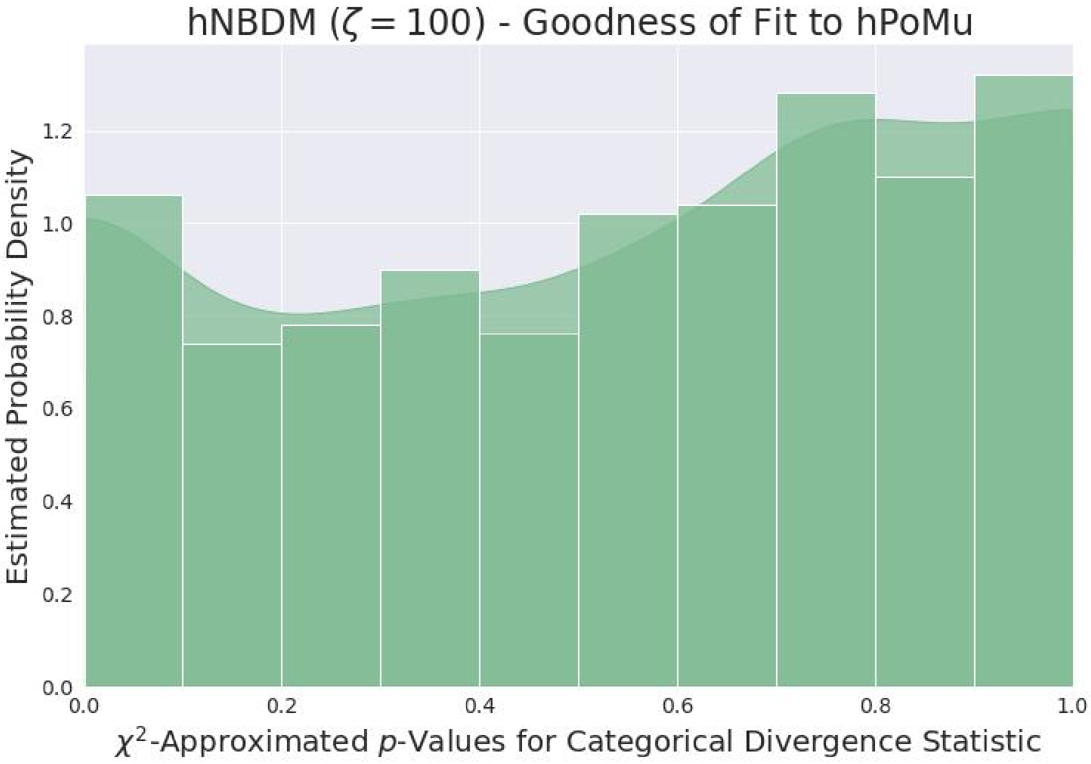

**Supplementary Figure S31**

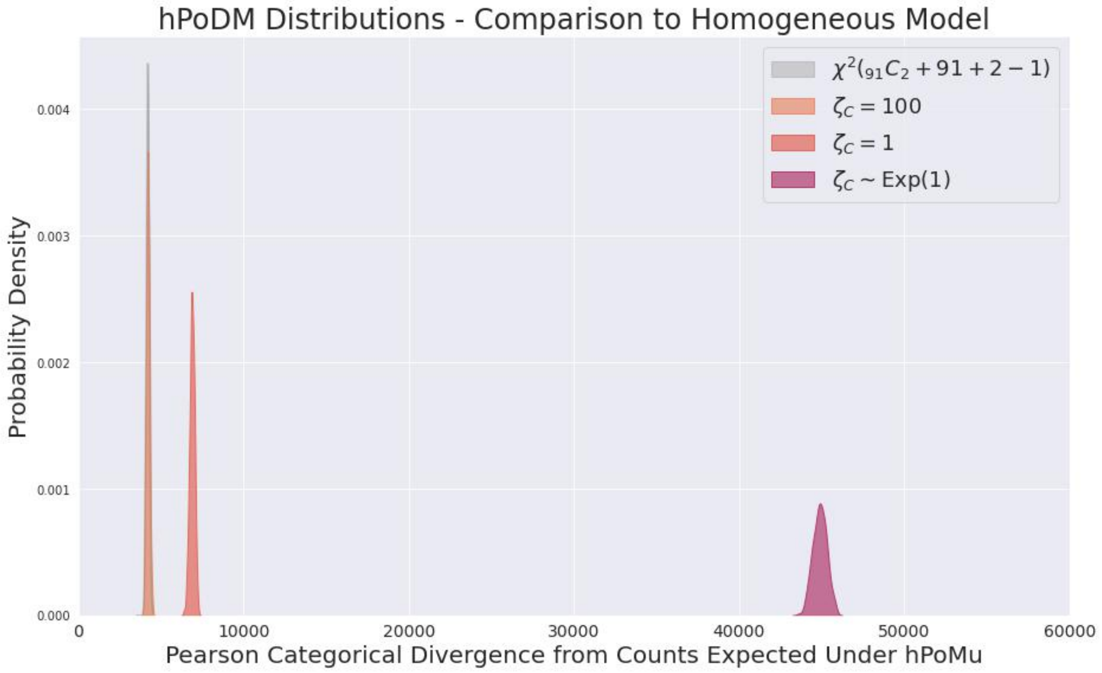

**Supplementary Figure S32**

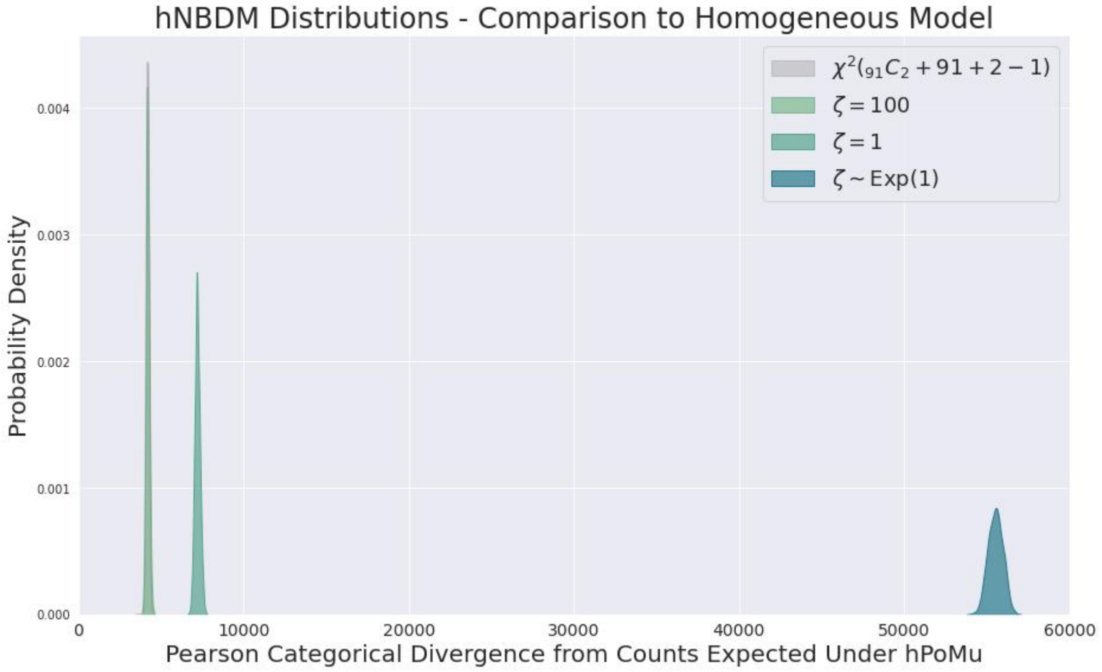

**Supplementary Figure S33**

number of droplets available to serve as treatments. If an estimation method is too starved of data to make a useful inference about a given microbial interaction, more likely than not the reason will be that there are too few treatment droplets where the strains co-occur. Therefore, in trying to account for the effects caused by the randomness of the initial droplet formation process, I argue that our main goal should be to understand the bottlenecks in the number of treatment droplets predicted by the various models. Thus understanding these predicted bottlenecks is the main goal of this section.

Another potential challenge arising from the randomness of the initial droplet formation process is the "multiple representatives problem", where within a given droplet there is initially a strain that is represented by multiple cells. Cf. figure S34. Even if we were more concerned about the effects randomness of the initial droplet formation process causes via the "multiple representatives problem" than the effects it has on data throughput, I argue we should still first focus on the effects it has on data throughput, for two reasons.

First, to the extent that the "multiple representatives problem" might create "noise" or "fluctuations" in the MOREI data, these effects will on average "cancel out" if the data throughput is great enough to ensure large sample sizes for both the treatment and control groups. Second, even to the extent the "multiple representatives problem" might create systematic errors which do not "cancel out" given large enough sample sizes, these systematic errors should become easier to identify, and therefore also easier to account for, with greater data throughput. In either case, figuring out where we can and need to (or where we can't) account for effects caused by the "multiple representatives problem" begins with identifying the bottlenecks in data throughput that the particular form of randomness of the initial droplet formation process causes.

#### S4.2 Preliminaries

In this section I clarify details needed to analyze the data throughput predicted by each model of initial droplet formation. Section  gives precise definitions of the statistics that are relevant for studying data throughput (following the argument given in section S4.1). In addition to the "picky" and "global" way of grouping of droplets by strain defined already in section S3.2.2, section S4.2.2 defines a "gluttonous" and "local" way of grouping droplets by strain.

##### *S4.2.1 Definition of Throughput*

Previously we have been looking at distributions of numbers of cells in a given droplet (of any strain or of a given strain). As argued in section S4.1, I believe that understanding the distributions of numbers of *strains* in a given droplet is the most relevant for predicting data throughput. In section S4.2.1 I clarify how to define these as statistics computed from the numbers of cells. Then sections S4.2.1 and S4.2.1 give two different possible definitions of throughput.

**Strain Count Distribution** The notation $\mathbb{I}\{A\}$ denotes the indicator function for the event $A$. For a given strain $s$, the **strain presence** RV (random variable):

$$\mathbb{S}^{(s)}(0) := \mathbb{I}\left\{N^{(s)}(0) \geq 1\right\} = \mathbb{I}\left\{N^{(s)}(0) > 0\right\} . \tag{S76}$$

indicates whether any cells of strain $s$ belong to the droplet. It takes values in $\{0,1\}$ like any indicator random variable. Specifically, it equals 1 if strain $s$ is present in the droplet and 0 if strain $s$ is absent.

Similarly, for a given strain $s$, the **strain absence** RV:

$$\overline{\mathbb{S}}^{(s)}(0) := 1 - \mathbb{S}^{(s)}(0) = \mathbb{I}\left\{N^{(s)}(0) = 0\right\} , \tag{S77}$$

takes values in $\{0,1\}$ like any indicator random variable, equalling 1 if strain $s$ is absent in the droplet and 0 if strain $s$ is present.

For a given droplet, the joint distribution of the strain presence RVs for all strains corresponds to a binary random vector:

$$\vec{\mathbb{S}}(0) := (\mathbb{S}^{(1)}(0), \dots, \mathbb{S}^{(S)}(0)) . \tag{S78}$$

Its values are in $\bigtimes_{s=1}^{S}\{0,1\}$ (the $S$-fold Cartesian product of $\{0,1\}$ with itself).

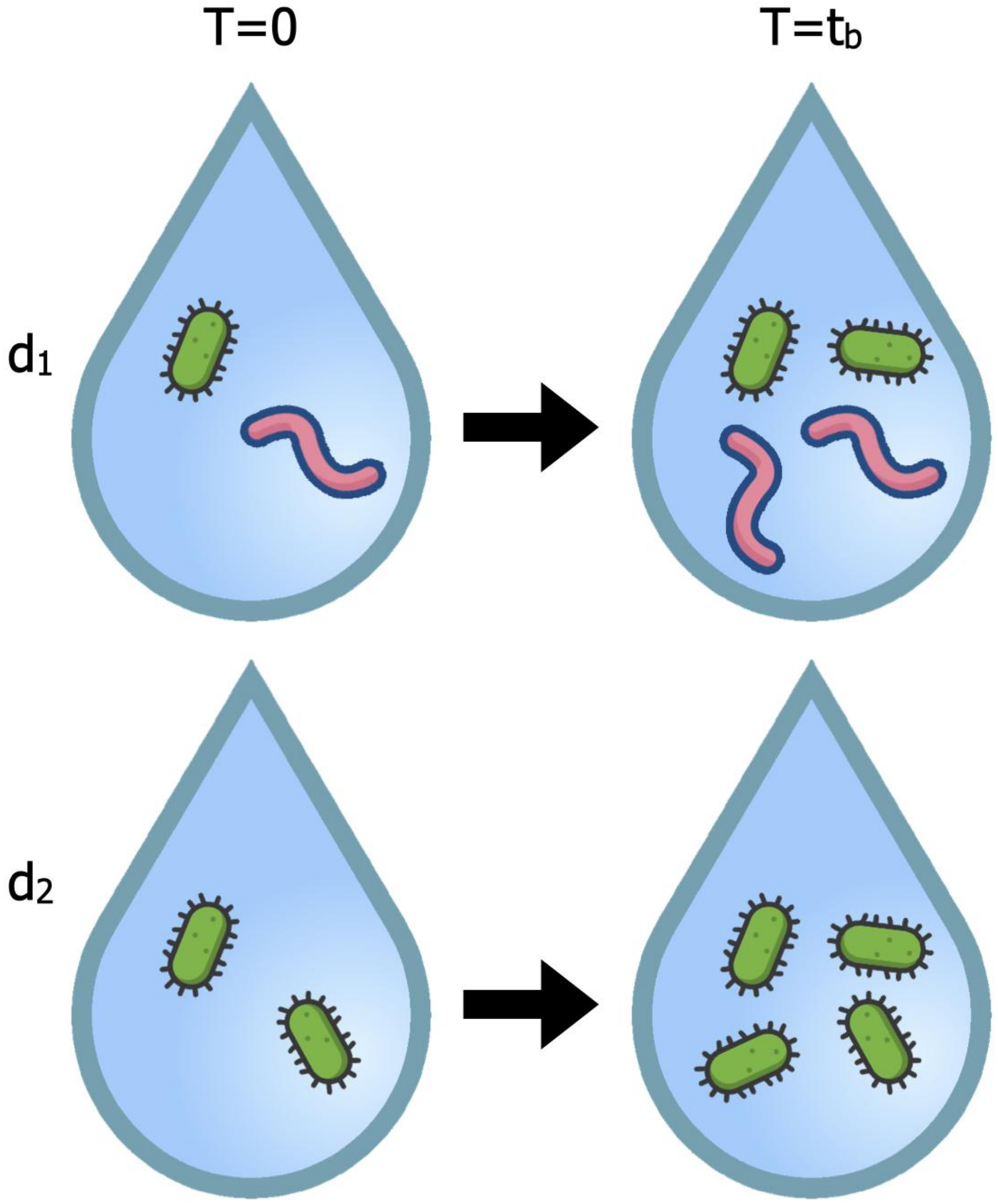

**Supplementary Figure S34.** Multiple Representatives Problem. The droplets are only observed at time $t_b$. If we incorrectly assume both droplets began with the same number of green cells, then we reach the wrong conclusion that the pink cells halve the growth rate of the green cells, when in fact they have no effect.

For a given droplet, its **strain count** $\mathbb{S}(0)$ is the number of strains present in the droplet. The strain count equals the sum of the strain presence RVs:

$$\mathbb{S}(0) := \sum_{s\in[S]} \mathbb{S}^{(s)}(0) = \sum_{s\in[S]} \mathbb{I}\left\{N^{(s)}(0) \geq 1\right\}. \tag{S79}$$

It is an RV taking values in $\{0\} \cup [S]$.

Another useful notion is the **support** of a (non-negative) vector $\vec{\mathbf{v}} \in \mathbb{R}^S_{\geq 0}$:

$$\operatorname{supp}(\vec{\mathbf{v}}) := \{s \in [S] : [\vec{\mathbf{v}}]_s > 0\} \subseteq [S]. \tag{S80}$$

It follows directly from the definitions that $\operatorname{supp}(\vec{\mathbf{N}}(0)) = \operatorname{supp}(\vec{\mathbb{S}}(0))$ always, corresponding to the set of $s \in [S]$ such that $\mathbb{S}^{(s)}(0) = 1$. Moreover we also always have[22] that $\mathbb{S}(0) = |\operatorname{supp}(\vec{\mathbf{N}}(0))| = |\operatorname{supp}(\vec{\mathbb{S}}(0))|$.

As argued already in section S4.1, I believe that the notion of "throughput" that is important for this problem corresponds to $\vec{\mathbb{S}}(0)$ and not (directly[23]) to $\vec{\mathbf{N}}(0)$. While choosing $\lambda \approx 2$ is intended to make $\vec{\mathbf{N}}(0)$ as similar to $\vec{\mathbb{S}}(0)$ as possible, they are definitely not the same, something which e.g. the multiple representatives problem from section S4.1 makes clear.

Even when the first moments of $\vec{\mathbf{N}}(0)$ remain the same, changes to the second moments of $\vec{\mathbf{N}}(0)$ can still cause changes in the expected numbers of droplets serving as treatments or controls. A heuristic way to understand this is to observe that the expected data throughput corresponds to the second (or higher order) moments of $\vec{\mathbb{S}}(0)$; see sections S4.2.1 and S4.2.1. In particular, although for any member of the ghNBDM family (see section S2.8.1) the expected cell counts for each strain are the same as those under hPoMu, the choice of distribution from the ghNBDM family still affects the predicted data throughput.

**Gluttonous Definition of Data Throughput** The treatment group for inferring the effect of strain $s_1$ on the growth of strain $s_2$ corresponds to droplets where $s_1$ and $s_2$ co-occur. The set of all droplets satisfying this condition is

$$\begin{aligned}
\mathcal{T}_g^{(s_1,s_2)} &:= \left\{d \in [D] : \mathbb{S}_d^{(s_1)}(0) \cdot \mathbb{S}_d^{(s_2)}(0) = 1\right\} \\
&= \left\{d \in [D] : \mathbb{I}\left\{N_d^{(s_1)}(0) \geq 1 \;\&\; N_d^{(s_2)}(0) \geq 1\right\} = 1\right\} \\
&= \left\{d \in [D] : \operatorname{supp}(\vec{\mathbb{S}}_d(0)) \supseteq \{s_1, s_2\}\right\}.
\end{aligned} \tag{S81}$$

This is called the "**gluttonous**" **treatment** group because *all* droplets where $s_1$ and $s_2$ co-occur are included. Cf. figure S35.

The control group for inferring the effect of strain $s_1$ on the growth of strain $s_2$ corresponds to droplets where $s_2$ occurs but $s_1$ does not. The set of all droplets satisfying this condition is

$$\begin{aligned}
\mathcal{C}_g^{(s_1,s_2)} &:= \left\{d \in [D] : \overline{\mathbb{S}}_d^{(s_1)}(0) \cdot \mathbb{S}_d^{(s_2)}(0) = 1\right\} \\
&= \left\{d \in [D] : \mathbb{I}\left\{N_d^{(s_1)}(0) = 0 \;\&\; N_d^{(s_2)}(0) \geq 1\right\} = 1\right\} \\
&= \left\{d \in [D] : \operatorname{supp}(\vec{\mathbb{S}}_d(0)) \supseteq \{s_2\},\, s_1 \notin \operatorname{supp}(\vec{\mathbb{S}}_d(0))\right\}.
\end{aligned} \tag{S82}$$

This is called the "**gluttonous**" **control** group because *all* droplets where $s_2$ occurs but $s_1$ does not are included. Cf. figure S35.

These definitions place no constraints on the presence or absence of strains $s \in [S]$ besides $s_1$ and $s_2$. This can be favorable for example when studying very rare strains, for which there may be no or very few droplets containing $s_1$ and $s_2$ only, but some containing $s_1$ and $s_2$ along with other strains. These definitions give us the best possible chance of avoiding "data starvation".

From equation (S81) we get (assuming the droplets are identically distributed) that the expected number of droplets for the gluttonous treatment group for inferring the effect of strain $s_2$ on the growth of $s_1$ is

$$\mathbb{E}\left[\left|\mathcal{T}_g^{(s_1,s_2)}\right|\right] = D \cdot \mathbb{E}\left[\mathbb{S}^{(s_1)}(0)\, \mathbb{S}^{(s_2)}(0)\right]. \tag{S83}$$

[22] The ***cardinality*** $|\mathcal{S}|$ of a finite set $\mathcal{S}$ is the number of elements of the set. E.g. $|[S]| = S$.

[23] Only indirectly as mediated via $\vec{\mathbb{S}}(0)$, which of course can be computed from $\vec{\mathbf{N}}(0)$.

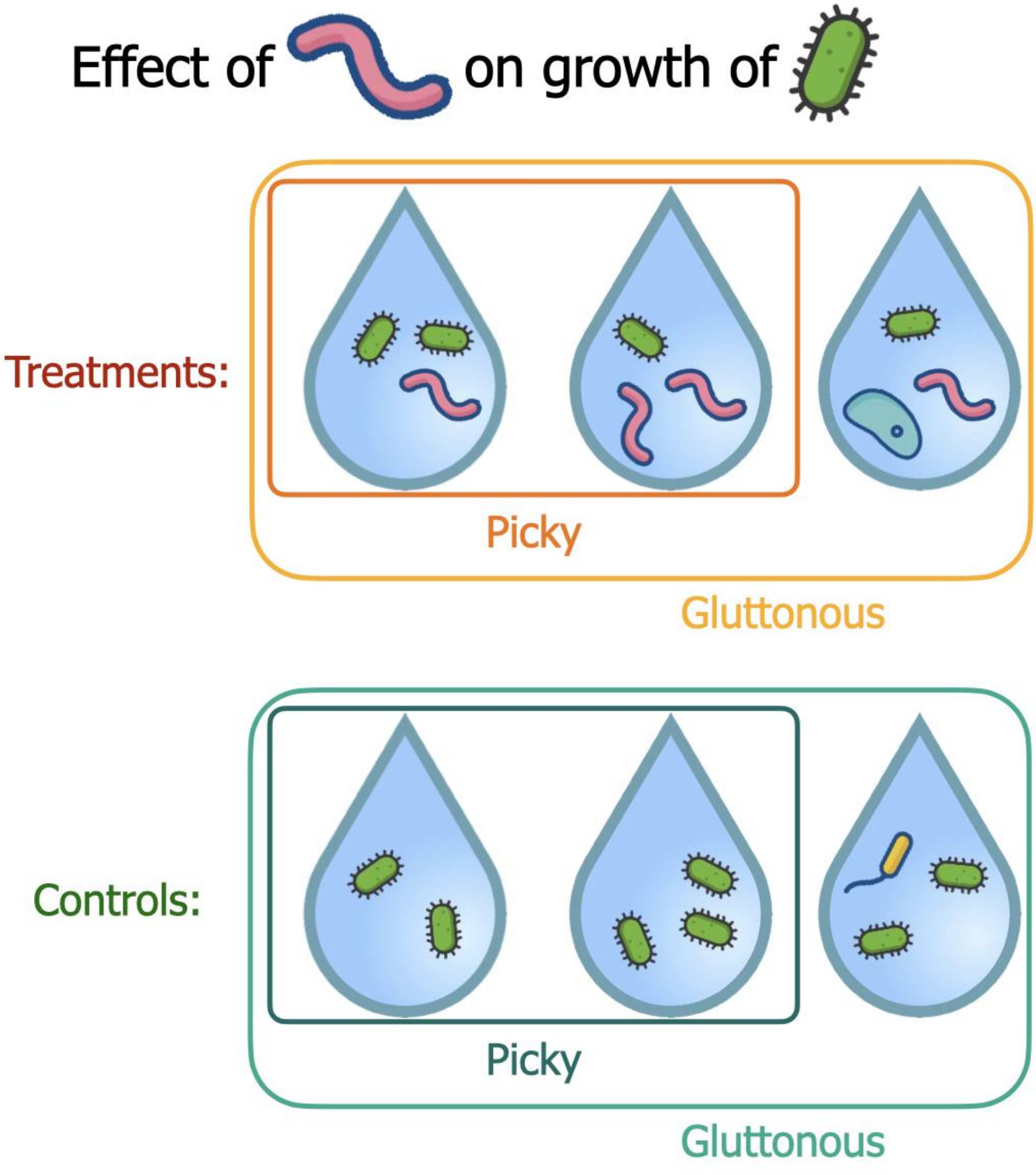

**Supplementary Figure S35.** Gluttonous vs. Picky Definitions of Data Throughput. Cf. text.

Similarly, from equation (S82) we get (assuming the droplets are identically distributed) that the expected number of droplets for the corresponding gluttonous control group for inferring the effect of strain $s_2$ on the growth of $s_1$ is

$$\mathbb{E}\left[\left|\mathcal{C}_g^{(s_1,s_2)}\right|\right] = D \cdot \mathbb{E}\left[\overline{\mathbb{S}}^{(s_1)}(0) \cdot \mathbb{S}^{(s_2)}(0)\right] = D \cdot \mathbb{E}\left[\mathbb{S}^{(s_2)}(0) - \mathbb{S}^{(s_1)}(0) \cdot \mathbb{S}^{(s_2)}(0)\right]. \tag{S84}$$

As claimed before in section S4.2.1, these expressions for the expected gluttonous data throughput involve the second moments of $\vec{\mathbb{S}}(0)$.

**Picky Definition of Data Throughput** We see that the above "gluttonous" definitions correspond to more possible combinations of strains than we probably would have included when studying the effect of strain $s_1$ on the growth of strain $s_2$ by manually plating cells. This motivates the following definitions.

The set of droplets where $s_1$ and $s_2$ co-occur in a way corresponding to the combination of strains we would use if we were manually plating cells is

$$\begin{aligned}\mathcal{T}_p^{(s_1,s_2)} &:= \left\{ d \in [D] : \mathbb{S}_d^{(s_1)}(0) \cdot \mathbb{S}_d^{(s_2)}(0) \prod_{\sigma \notin \{s_1,s_2\}} \overline{\mathbb{S}}_d^{(\sigma)}(0) = 1 \right\} \\ &= \left\{ d \in [D] : \mathbb{I}\left\{ N_d^{(s_1)}(0) \geq 1 \;\&\; N_d^{(s_2)}(0) \geq 1 \;\&\; \forall \sigma \notin \{s_1,s_2\},\ N_d^{(\sigma)}(0) = 0 \right\} = 1 \right\} \\ &= \left\{ d \in [D] : \operatorname{supp}(\vec{\mathbb{S}}_d(0)) = \{s_1,s_2\} \right\}.\end{aligned} \tag{S85}$$

This is called the "**picky**" **treatment** group because droplets where $s_1$ and $s_2$ co-occur are included *only when all other strains are absent*. Cf. figure S35.

The set of droplets where *only* $s_2$ occurs, corresponding to the combination of strains we would use if we were manually plating cells, is

$$\begin{aligned}\mathcal{C}_p^{(s_1,s_2)} &:= \left\{ d \in [D] : \overline{\mathbb{S}}_d^{(s_1)}(0) \cdot \mathbb{S}_d^{(s_2)}(0) \prod_{\sigma \notin \{s_1,s_2\}} \overline{\mathbb{S}}_d^{(\sigma)}(0) = 1 \right\} \\ &= \left\{ d \in [D] : \mathbb{I}\left\{ N_d^{(s_1)}(0) = 0 \;\&\; N_d^{(s_2)}(0) \geq 1 \;\&\; \forall \sigma \notin \{s_1,s_2\},\ N_d^{(\sigma)}(0) = 0 \right\} = 1 \right\} \\ &= \left\{ d \in [D] : \operatorname{supp}(\vec{\mathbb{S}}_d(0)) = \{s_2\} \right\}.\end{aligned} \tag{S86}$$

This is called the "**picky**" **control** group because droplets where $s_2$ occurs but $s_1$ does not occur are included *only when all other strains are absent*. It is not enough for $s_1$ alone to be absent. Cf. figure S35.

Unlike the gluttonous groups, the picky groups *do* place constraints on the presence or absence of strains $s \in [S]$ besides $s_1$ and $s_2$. For both the treatments and the controls, for "picky" groups all strains besides $s_1$ or $s_2$ must have zero counts. Every picky group is by definition a subset of its gluttonous counterpart.

The picky definitions can be favorable when we have plenty of droplets from which to make estimates. They reduce the possibility of confounding effects on the growth of strain $s_2$ that could be caused by strains that are not strain $s_1$. On the other hand, in instances where there are only very few or no droplets without "extra" strains, the picky definitions could lead to "data starvation".

From equation (S85) we get (assuming the droplets are identically distributed) that the expected number of droplets for the picky treatment group for inferring the effect of strain $s_2$ on the growth of $s_1$ is

$$\mathbb{E}\left[\left|\mathcal{T}_p^{(s_1,s_2)}\right|\right] = D \cdot \mathbb{E}\left[\mathbb{S}^{(s_1)}(0) \cdot \mathbb{S}^{(s_2)}(0) \prod_{\sigma \notin \{s_1,s_2\}} \left(1 - \mathbb{S}^{(\sigma)}(0)\right)\right]. \tag{S87}$$

Similarly, from equation (S86) we get (assuming the droplets are identically distributed) that the expected number of droplets for the corresponding picky control group for inferring the effect of strain $s_2$ on the growth of $s_1$ is

$$\mathbb{E}\left[\left|\mathcal{C}_p^{(s_1,s_2)}\right|\right] = D \cdot \mathbb{E}\left[\left(1 - \mathbb{S}^{(s_1)}(0)\right) \cdot \mathbb{S}^{(s_2)}(0) \prod_{\sigma \notin \{s_1,s_2\}} \left(1 - \mathbb{S}^{(\sigma)}(0)\right)\right]. \tag{S88}$$

As claimed before in section S4.2.1, these expressions for the expected picky data throughput involve the higher order moments of $\vec{\mathbb{S}}(0)$.

##### S4.2.2 Pairwise Gluttonous Goodness of Fit Tests

In section S3.2.2 droplets were grouped according to strain in a non-overlapping way. This made it possible to compare other distributions to hPoMu using a single hypothesis test, but for practical reasons it also required lumping together all droplets with three or more strains. However, if we are willing to perform multiple hypothesis tests, we can better utilize the information contained in those droplets. By performing a separate hypothesis test for each pair of strains, we can include more droplets in both our "treatment" and "control" groups.

The definition of the hypothesis test corresponding to each pair of strains is split into several parts. Section S4.2.2 defines how the droplets are grouped. Section S4.2.2 gives probabilities for each group under the null hypothesis (hPoMu), while section S4.2.2 clarifies the expected counts. Section S4.2.2 defines the observed counts used in defining the goodness of fit test. Section S4.2.2 explains why some of the groups are redundant. Finally, section S4.2.2 combines the information from the previous sections to define the test statistic.

**Categories** Note that the above definitions correspond to "gluttonous" groups, unlike the "global" test defined before which corresponds to "picky" groups. The "gluttony" means that the categories for distinct combinations of strains can overlap. This is what necessitates separate tests for each combination of strains, i.e. why they are "pairwise".

**Probabilities** The probabilities $p_{0,0}^{(s_1,s_2)}$,$p_{1,0}^{(s_1,s_2)}$, $p_{1,1}^{(s_1,s_2)}$, $p_{0,1}^{(s_1,s_2)}$, for the pairwise tests are

$$\begin{aligned}
p_{0,0}^{(s_1,s_2)} &:= \mathbb{P}_{\text{hPoMu}(\lambda,\vec{f})}\left(N^{(s_1)}(0)=0, N^{(s_2)}(0)=0\right) \\
&= e^{-f^{(s_1)}\lambda} \cdot e^{-f^{(s_2)}\lambda}, \\
p_{1,0}^{(s_1,s_2)} &:= \mathbb{P}_{\text{hPoMu}(\lambda,\vec{f})}\left(N^{(s_1)}(0)\geq 1, N^{(s_2)}(0)=0\right) \\
&= \left(1-e^{-f^{(s_1)}\lambda}\right) \cdot \cdot e^{-f^{(s_2)}\lambda}, \\
p_{1,1}^{(s_1,s_2)} &:= \mathbb{P}_{\text{hPoMu}(\lambda,\vec{f})}\left(N^{(s_1)}(0)\geq 1, N^{(s_2)}(0)\geq 1\right) \\
&= \left(1-e^{-f^{(s_1)}\lambda}\right) \cdot \left(1-e^{-f^{(s_2)}\lambda}\right), \\
p_{0,1}^{(s_1,s_2)} &:= \mathbb{P}_{\text{hPoMu}(\lambda,\vec{f})}\left(N^{(s_1)}(0)=0, N^{(s_2)}(0)\geq 1\right) \\
&= e^{-f^{(s_1)}\lambda} \cdot \left(1-e^{-f^{(s_2)}\lambda}\right).
\end{aligned} \tag{S89}$$

Note the implicit dependence on an assumed $\lambda$ in the definitions of $p_{0,0}^{(s_1,s_2)}$, $p_{1,0}^{(s_1,s_2)}$, $p_{1,1}^{(s_1,s_2)}$, $p_{0,1}^{(s_1,s_2)}$. Thus the scientist needs an estimate, or some a priori knowledge, of the value of $\lambda$ to use these tests.

**Expected Counts** The expected counts for the pairwise tests are

$$\begin{aligned}
M_{0,0}^{(s_1,s_2)} &:= D \cdot p_{0,0}^{(s_1,s_2)}, \\
M_{1,0}^{(s_1,s_2)} &:= D \cdot p_{1,0}^{(s_1,s_2)}, \\
M_{1,1}^{(s_1,s_2)} &:= D \cdot p_{1,1}^{(s_1,s_2)}, \\
M_{0,1}^{(s_1,s_2)} &:= D \cdot p_{0,1}^{(s_1,s_2)}.
\end{aligned} \tag{S90}$$

**Observed Counts** The observed counts are defined as

$$\begin{aligned}
\hat{M}_{0,0}^{(s_1,s_2)} &:= \left|\left\{d : N_d^{(s_1)}(0)=0, N_d^{(s_2)}(0)=0\right\}\right|, \\
\hat{M}_{1,0}^{(s_1,s_2)} &:= \left|\left\{d : N_d^{(s_1)}(0)\geq 1, N_d^{(s_2)}(0)=0\right\}\right|, \\
\hat{M}_{1,1}^{(s_1,s_2)} &:= \left|\left\{d : N_d^{(s_1)}(0)\geq 1, N_d^{(s_2)}(0)\geq 1\right\}\right|, \\
\hat{M}_{0,1}^{(s_1,s_2)} &:= \left|\left\{d : N_d^{(s_1)}(0)=0, N_d^{(s_2)}(0)\geq 1\right\}\right|.
\end{aligned} \tag{S91}$$

**Symmetry of Definitions** Of course,

$$\begin{aligned}
\hat{M}_{1,0}^{(s_1,s_2)} &= \hat{M}_{0,1}^{(s_2,s_1)}, & p_{1,0}^{(s_1,s_2)} &= p_{0,1}^{(s_2,s_1)}, \\
\hat{M}_{0,0}^{(s_1,s_2)} &= \hat{M}_{0,0}^{(s_2,s_1)}, & p_{0,0}^{(s_1,s_2)} &= p_{0,0}^{(s_2,s_1)}, \\
\hat{M}_{1,1}^{(s_1,s_2)} &= \hat{M}_{1,1}^{(s_2,s_1)}, & p_{1,1}^{(s_1,s_2)} &= p_{1,1}^{(s_2,s_1)}.
\end{aligned} \tag{S92}$$

Therefore the results of the test for $(s_2, s_1)$ will always be the same as those for $(s_1, s_2)$, and so at most one of the two tests needs to be performed.

**Test Statistic** The test statistic is the Pearson categorical divergence of

$$\begin{aligned}
& D^{-1} \cdot \left( \hat{M}_{0,0}^{(s_1,s_2)}, \hat{M}_{1,0}^{(s_1,s_2)}, \hat{M}_{1,1}^{(s_1,s_2)}, \hat{M}_{0,1}^{(s_1,s_2)} \right) \\
\text{relative to} \quad & D^{-1} \cdot \left( M_{0,0}^{(s_1,s_2)}, M_{1,0}^{(s_1,s_2)}, M_{1,1}^{(s_1,s_2)}, M_{0,1}^{(s_1,s_2)} \right),
\end{aligned} \tag{S93}$$

namely

$$\sum_{i=0}^{1} \sum_{j=0}^{1} \frac{\left( \hat{M}_{i,j}^{(s_1,s_2)} - M_{i,j}^{(s_1,s_2)} \right)^2}{M_{i,j}^{(s_1,s_2)}}. \tag{S94}$$

One can then get an asymptotic $p$-value for this test in the standard way, via the survival function of the $\chi^2$ distribution with 3 degrees of freedom.

#### S4.3 Methods

Section S4.3.1 explains how percentages and percent changes for droplets with a given number of strains were computed. Section S4.3.2 explains how I computed the percent changes for gluttonous groups. Section S4.3.3 explains how I computed the percent changes for picky groups. I plotted results using Matplotlib (Hunter, 2007) version 3.4.1 and Seaborn (Waskom, 2021) version 0.11.1. The bar plot (Supplementary Figure S36) was made using the *microbiome* (Lahti and Shetty, 2019), *phyloseq* (McMurdie and Holmes, 2013), and *ggplot2* (Wickham, 2016) packages for the *R* programming language. Complete implementation details can be found in the code at the relevant GitLab repository. See https://gitlab.com/krinsman/droplets.

##### S4.3.1 Numbers of Droplets with $n$ Strains

Using the relationships:

$$\begin{aligned}
\mathbb{E}[|\{d \in [D] : \mathbb{S}_d(0) = n\}|] =& D \cdot \mathbb{P}(\mathbb{S}(0) = n) \\
=& D \cdot \left[ \sum_{\mathcal{S} \subseteq [S] : |\mathcal{S}| = n} \mathbb{P}(\operatorname{supp}(\vec{\mathbb{S}}(0)) = \mathcal{S}) \right],
\end{aligned} \tag{S95}$$

I computed via brute force these quantities for $n \in [5]$ for hPoMu:

$$\begin{aligned}
& D \cdot \left[ \sum_{\mathcal{S} \subseteq [S] : |\mathcal{S}| = n} e^{-\sum_{s \notin \mathcal{S}} f^{(s)} \lambda} \cdot \prod_{s \in \mathcal{S}} \left( 1 - e^{-f^{(s)} \lambda} \right) \right] \\
=& D \cdot \left[ \sum_{\mathcal{S} \subseteq [S] : |\mathcal{S}| = n} e^{-(1 - \sum_{s \in \mathcal{S}} f^{(s)}) \lambda} \cdot \prod_{s \in \mathcal{S}} \left( 1 - e^{-f^{(s)} \lambda} \right) \right].
\end{aligned} \tag{S96}$$

The value for 6+ strains is just $D$ minus the sum of the values for all $n \in [5]$.

For each of the seven distributions, I stratified and then counted the droplets from each of the 500 simulations according to how many strains were present. I then computed the average values for each distribution as the arithmetic mean over the 500 simulations. (By the law of large numbers the arithmetic mean is consistent for the true expectation and the only reasonable choice of measure of "central tendency" in this context.) I then computed the average percent change for each distribution by subtracting the value expected under hPoMu from the mean observed value, and then dividing by the expected value.

###### S4.3.2 Gluttonous Groups

Using the formulae (S89) and (S90), I computed the expected count for each treatment gluttonous group[24] under hPoMu. For each of the seven distributions, for each of the 500 simulations using NumPy (Harris et al., 2020) version 1.20.2 I computed via brute force the number of occurrences of each of the gluttonous groups, and then reported the arithmetic means over all 500 simulations. I again computed the average percent change as the observed value minus the value expected under hPoMu divided by the value expected under hPoMu.

Initially I considered "filtering" differences depending on whether they were "significant" according to the (asymptotic) $p$-values of the hypothesis tests defined in section S4.2.2. However, I found that doing so made the results more difficult to interpret by obscuring overall trends. The inferred overall trends were qualitatively the same in either case. Moreover, because each gluttonous pair corresponds to a distinct hypothesis test, and the gluttonous groups overlap, this leads to a difficult multiple comparisons problem. One has to not only decide how to account for multiple testing between simulations but also "within" simulations. Each of the multiple testing correction schemes I tried seemed objectionable in some way. Given all of the above, I decided to report the "raw" average percent change without "filtering" for "significance".

###### S4.3.3 Picky Groups

Using the formulae (S64) and (S65), cf. also the equations (S87) and (S88), I computed the expected count for each picky group under hPoMu. Note that, unlike for gluttonous treatment groups, picky treatment groups do not overlap. Moreover, unlike for gluttonous control groups, the picky control group for investigating the effect of strain $s_1$ on the rate of growth of strain $s_2$ is the same regardless of what $s_1$ is. So the values shown on the diagonals of the heat maps really correspond to the actual picky control groups. The procedure was then otherwise exactly the same as for the gluttonous groups. I did not "filter" according to "significance".

##### S4.4 Results

Section S4.4.1 explains how the observed effects on data throughput support the "taxonomy" of distributions previously proposed in section S3.4. Section explains the evidence indicating that increased heterogeneity decreases the average number of droplets with 3 or more strains. Section S4.4.3 explains the effect of increased heterogeneity on data throughput for droplets with 2 or fewer strains, with section S4.4.3 explaining what happens for compositional heterogeneity without density heterogeneity, and section S4.4.3 explaining what happens when both types of heterogeneity are present.

###### S4.4.1 Distributions Again Fall Into Same Groups

From figures S36, S37, S38, S39, S40, S41, S42, and S43 we see that hTPMH, hPoDM with $\zeta_C = 100$, and hNBDM with $\zeta_C = 100$ are again (cf. section S3.4) either effectively indistinguishable or very similar to hPoMu. Any differences in average composition are only visible for medium and high heterogeneities, and easily visible usually only for high heterogeneity.

###### S4.4.2 Increased Heterogeneity Decreases Number of Droplets with 3 or More Strains

From figures S36 and S37 we see that, both for the hPoDM and hNBDM families, as the compositional heterogeneity increases, the number of droplets with 3 or more strains decreases in both absolute and relative terms. Since droplets with 3 or more strains are a component of the gluttonous groups, their decrease with increased compositional heterogeneity is also reflected in figures S44, S46, S45, and S47. We can also see from figures S37, S44, S46, S45, and S47 that this decrease in the number of droplets with 3 or more strains is more pronounced in the absence of density heterogeneity.

###### S4.4.3 Effect of Increased Heterogeneity on Droplets with Fewer than 3 Strains

For droplets with fewer than 3 strains, the effects that occur as compositional heterogeneity increases depend on whether density heterogeneity is present.

[24] For the gluttonous "controls", i.e. corresponding to the diagonals of the heatmaps, I actually used the sets $\{d \in [D] : N^{(s)}(0) \geq 1\}$, which is a superset of all gluttonous controls for experiments measuring effects on the growth of strain $s$. The expected size of these sets is $D \cdot \mathbb{E}[\mathbb{S}^{(s)}(0)] = D \cdot \mathbb{P}(N^{(s)}(0) \geq 1)$, which under hPoMu equals $D \cdot (1 - e^{-f^{(s)}\lambda})$.

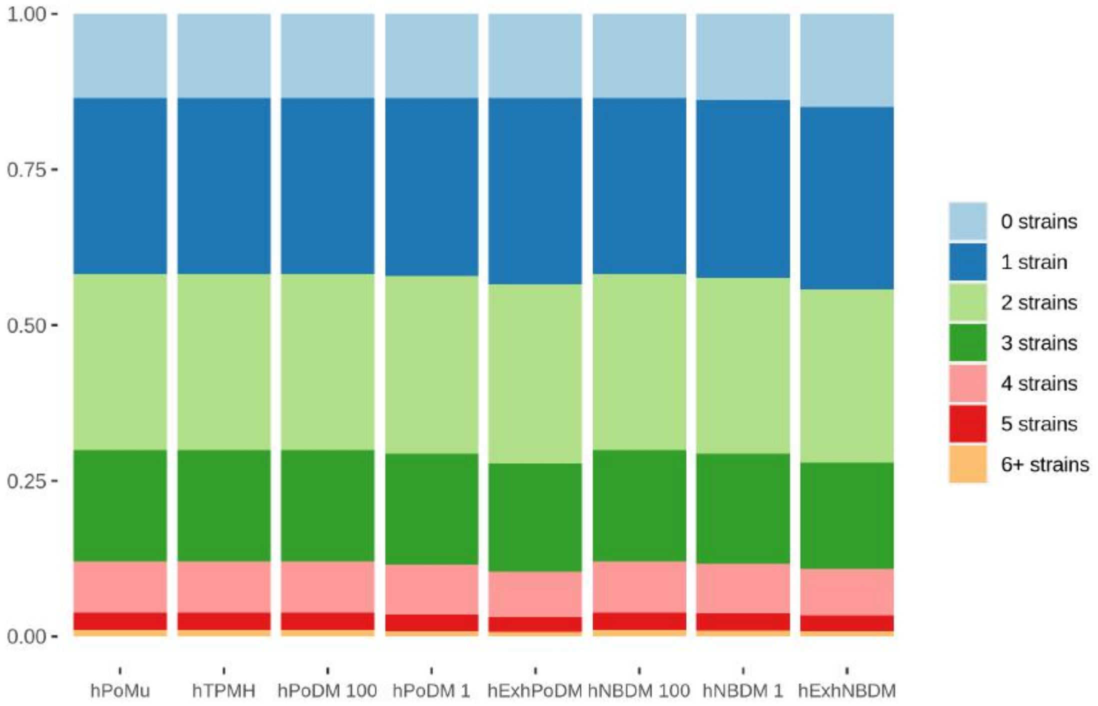

**Supplementary Figure S36**

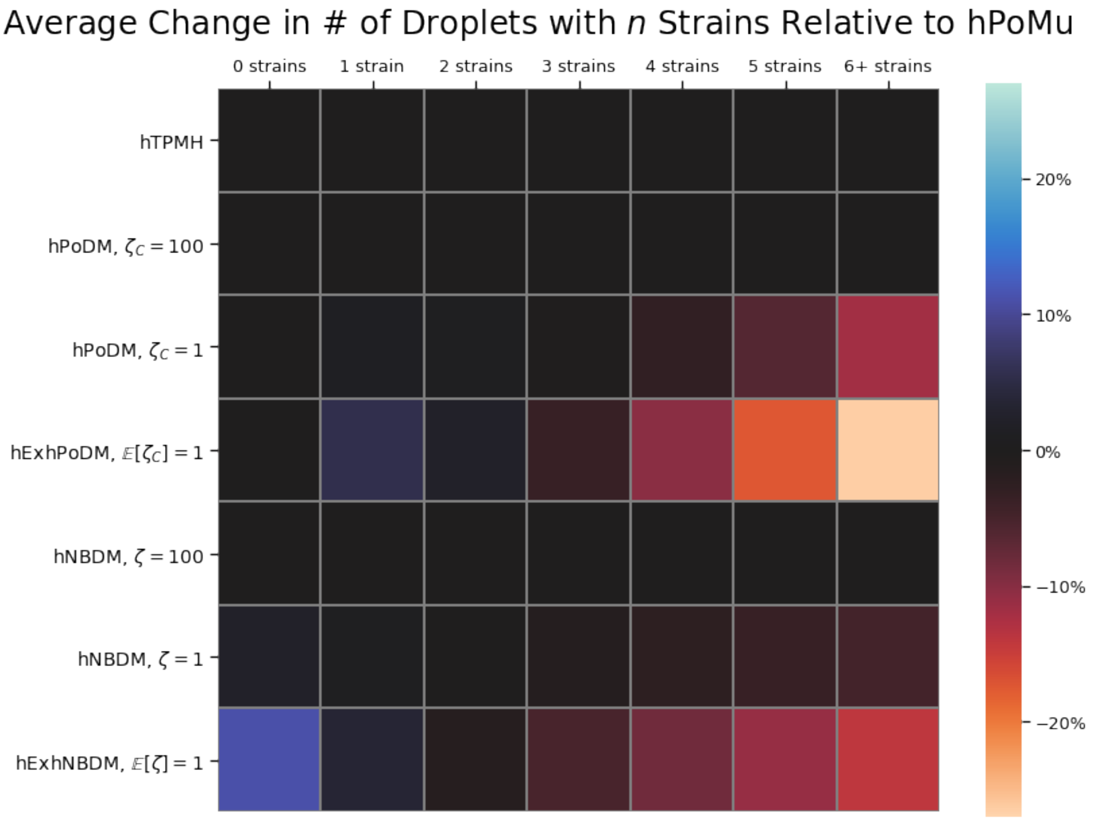

**Supplementary Figure S37**

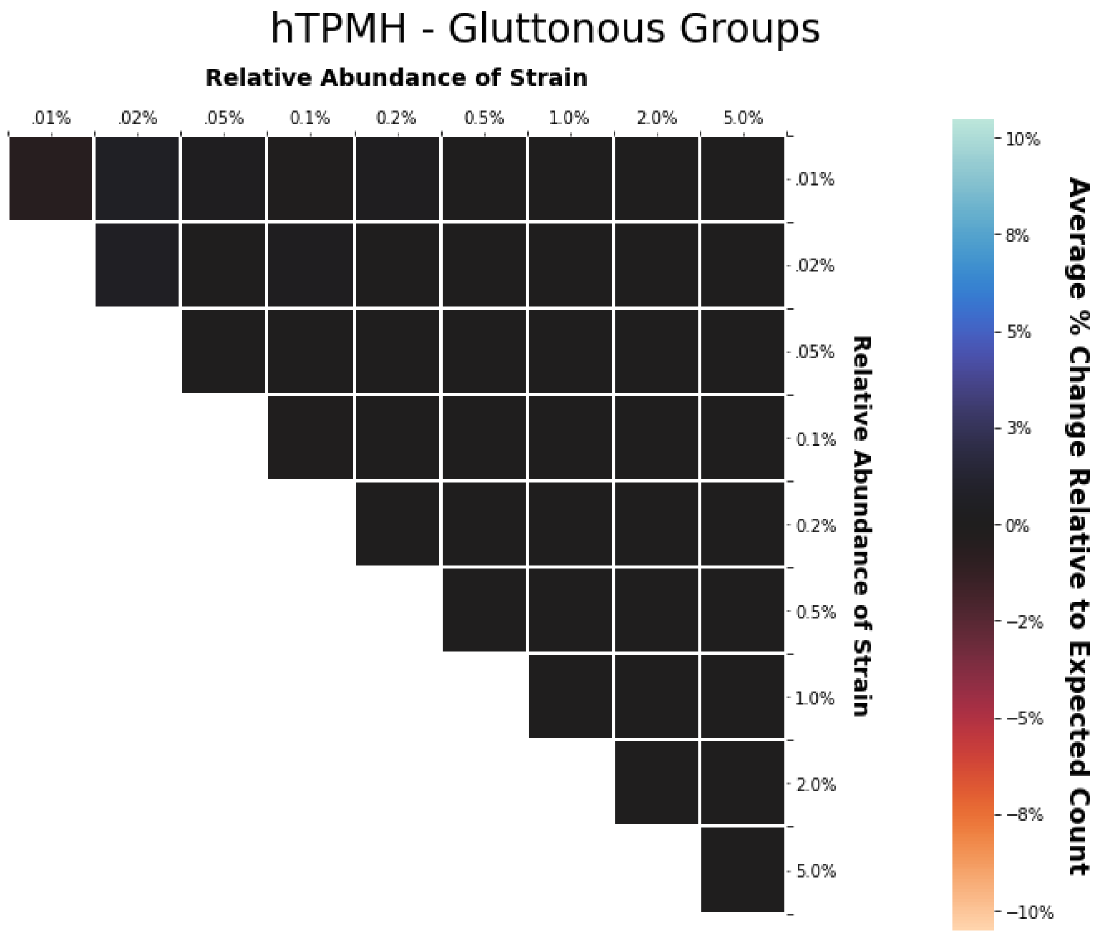

**Supplementary Figure S38**

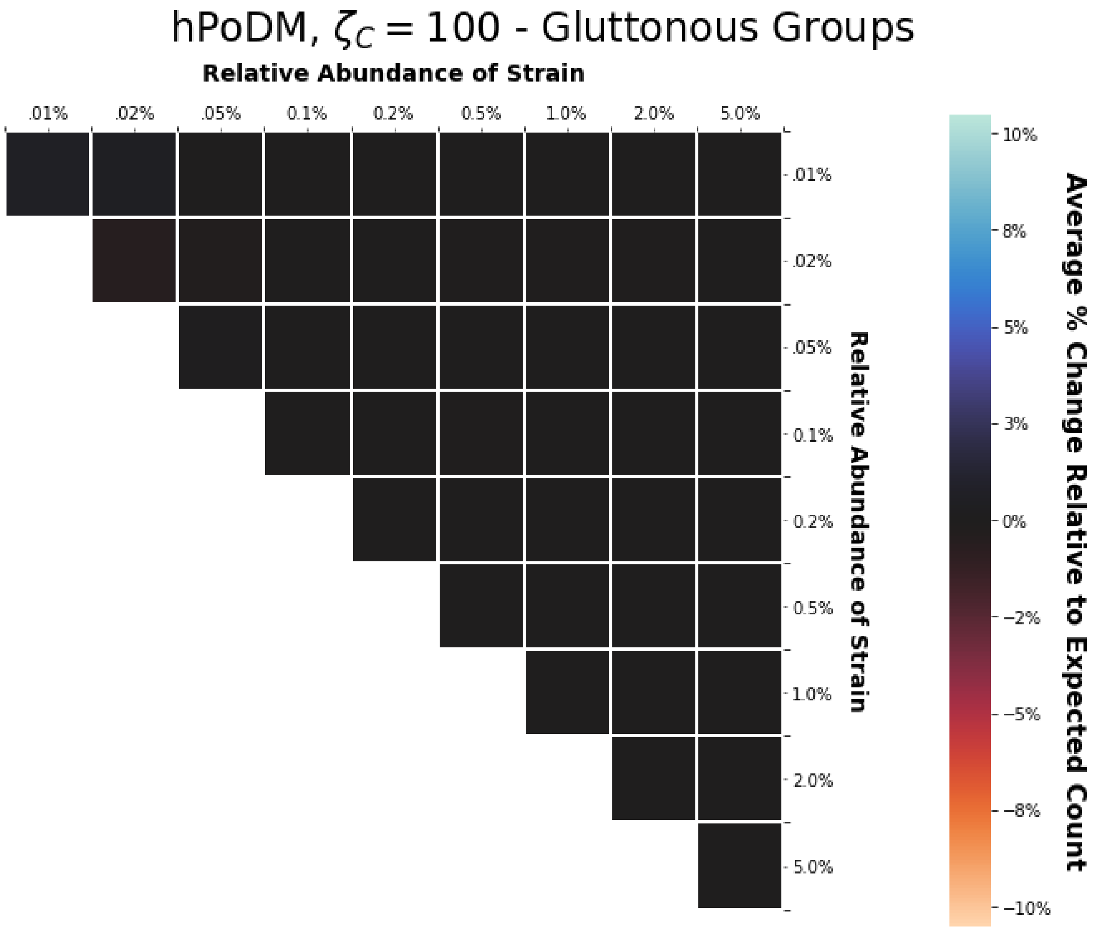

**Supplementary Figure S39**

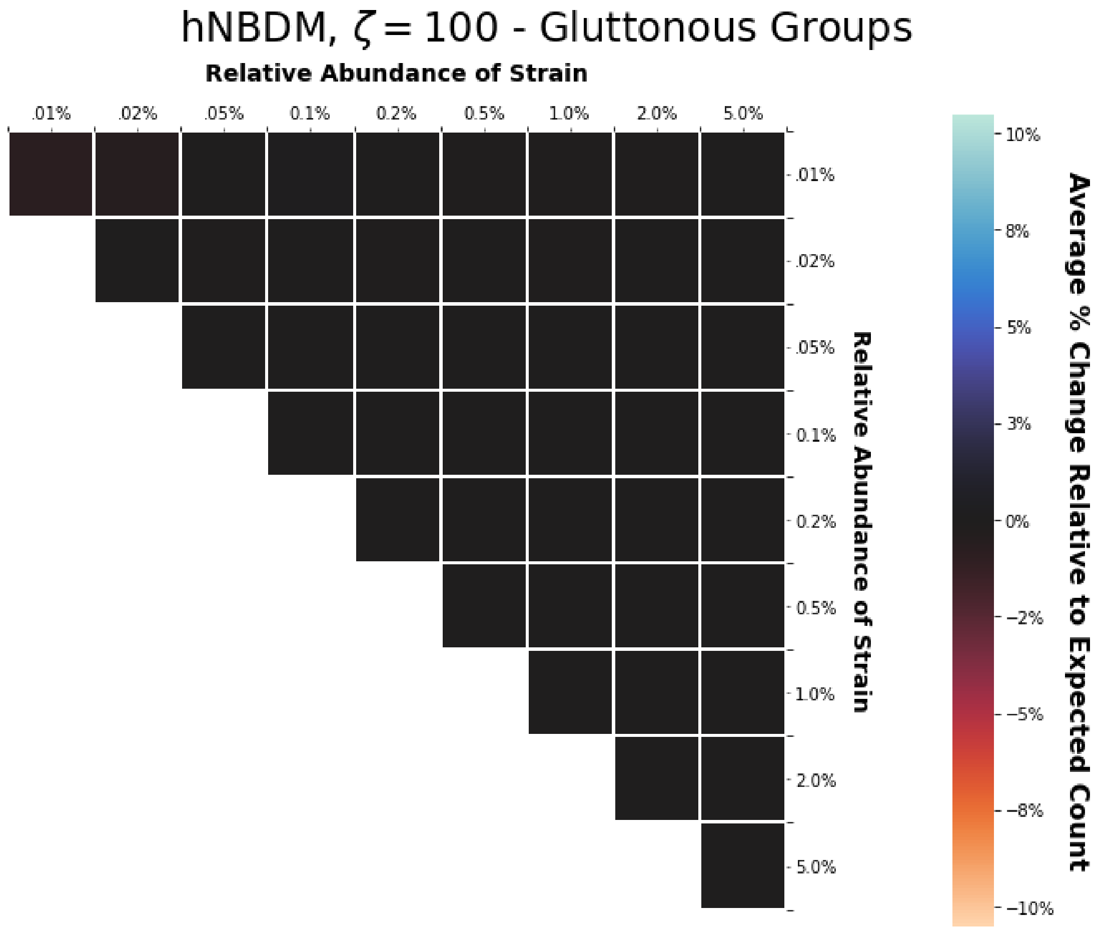

Supplementary Figure S40

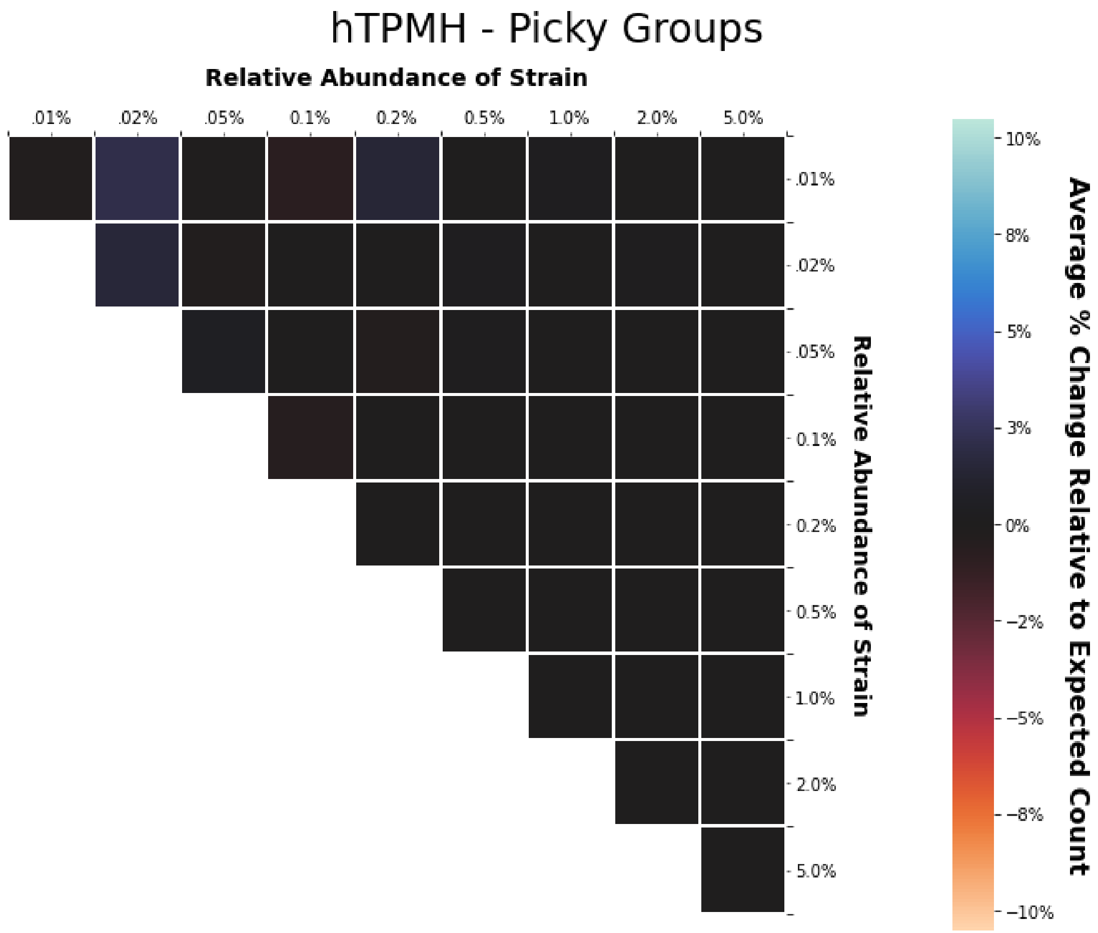

**Supplementary Figure S41**

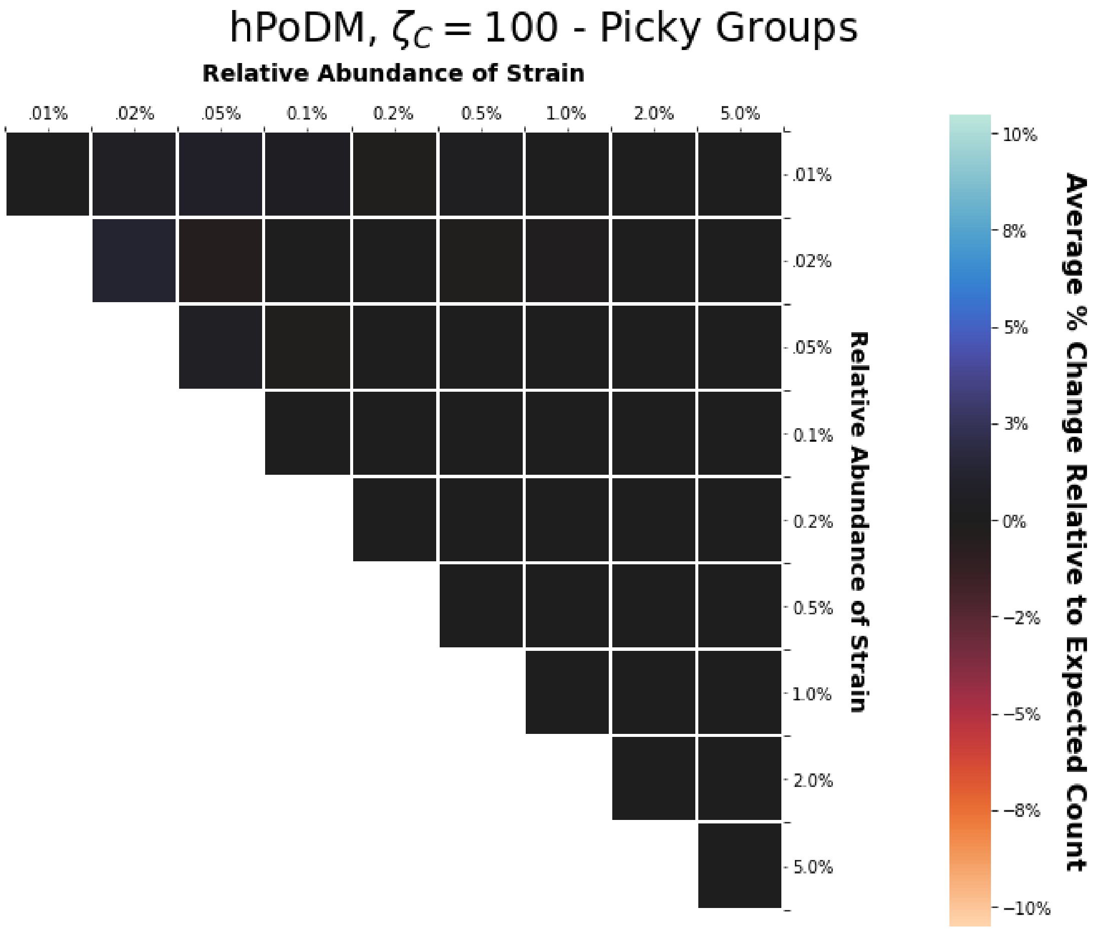

**Supplementary Figure S42**

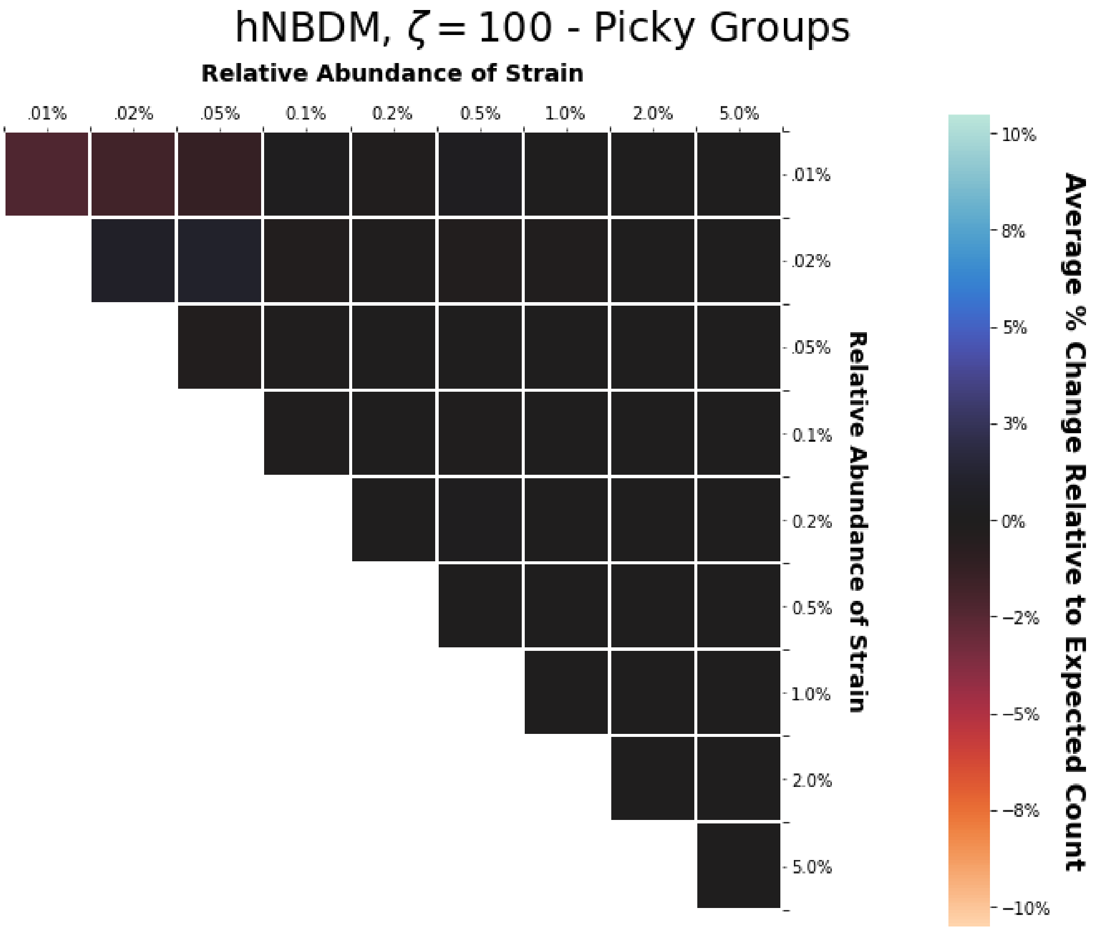

**Supplementary Figure S43**

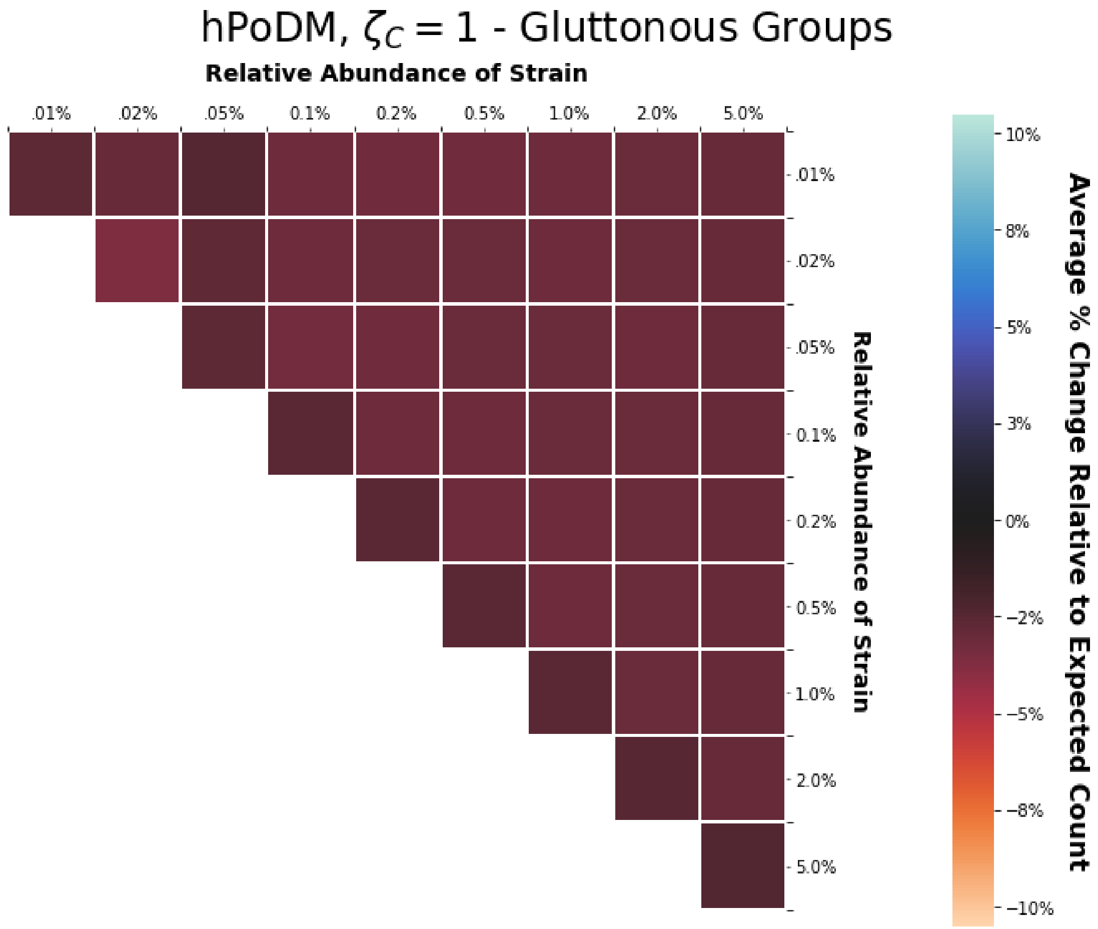

**Supplementary Figure S44**

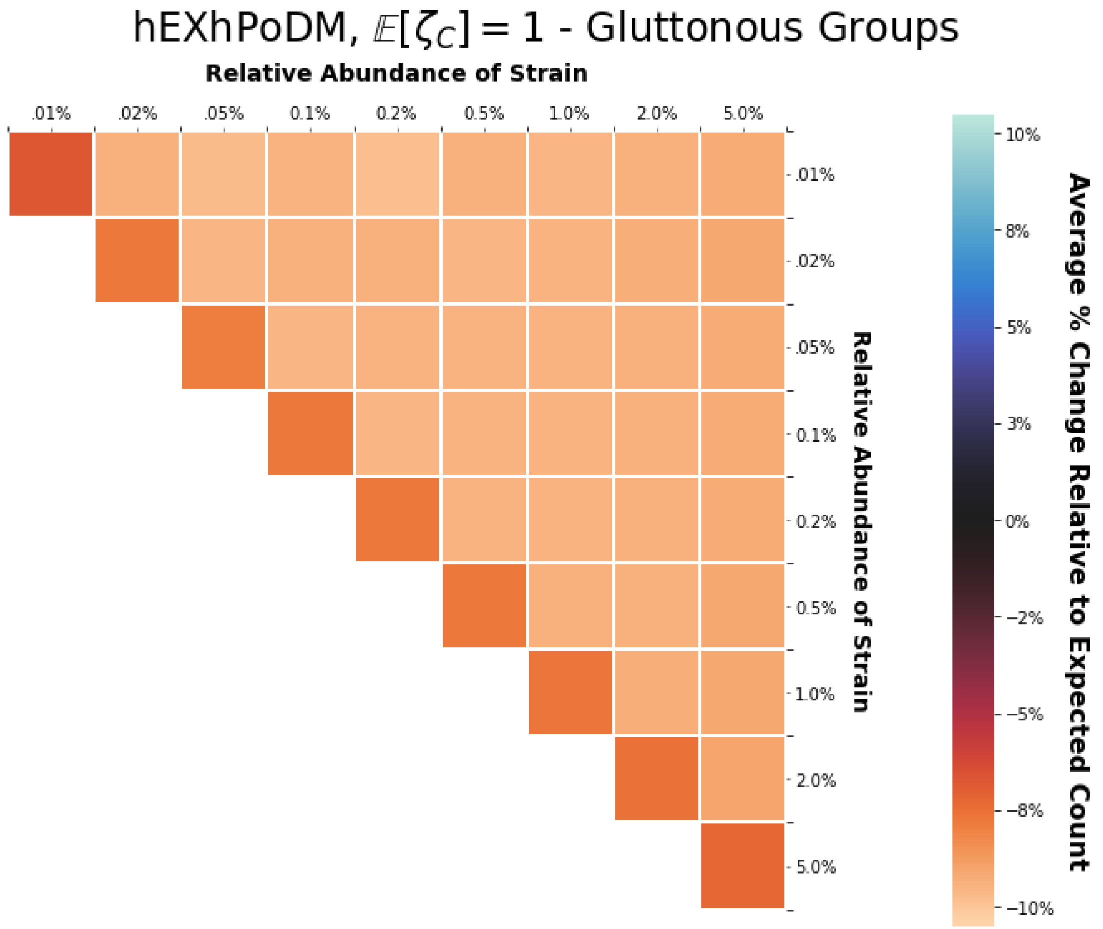

**Supplementary Figure S45**

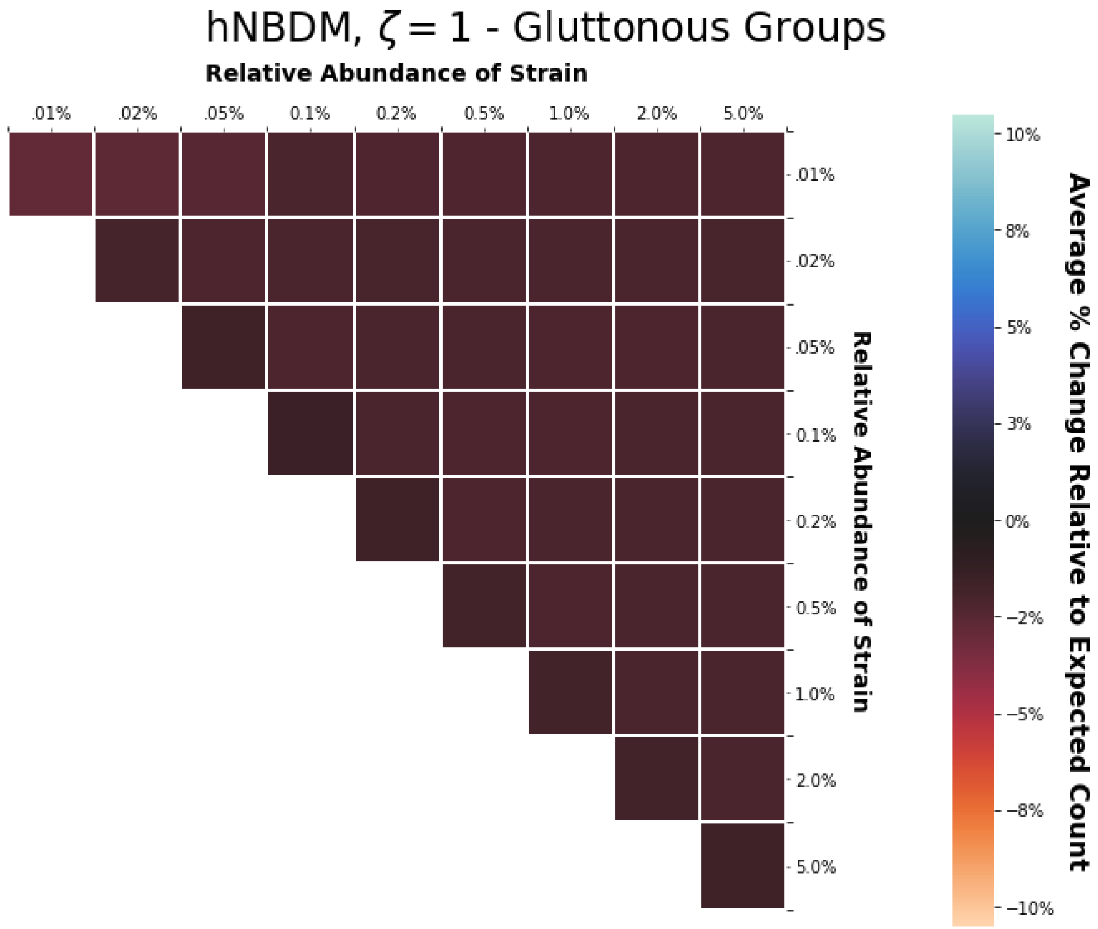

**Supplementary Figure S46**

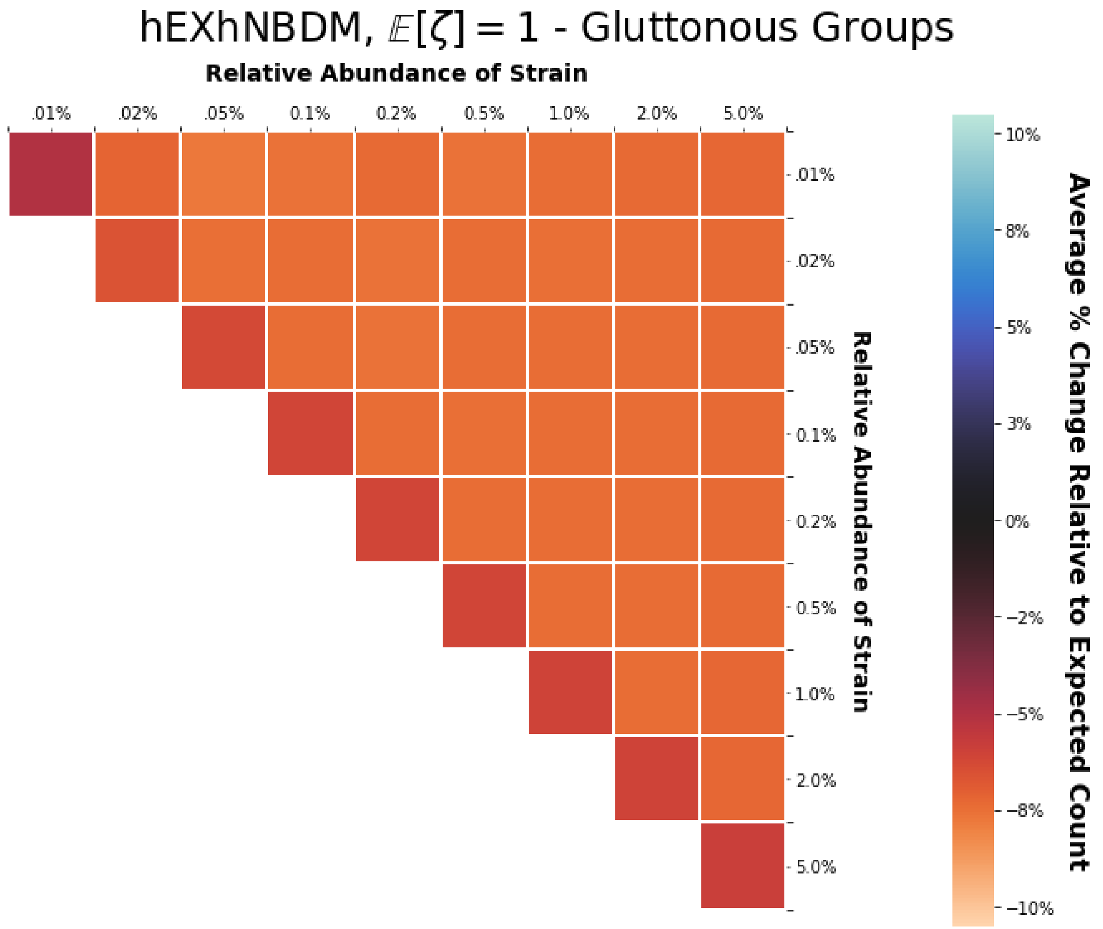

**Supplementary Figure S47**

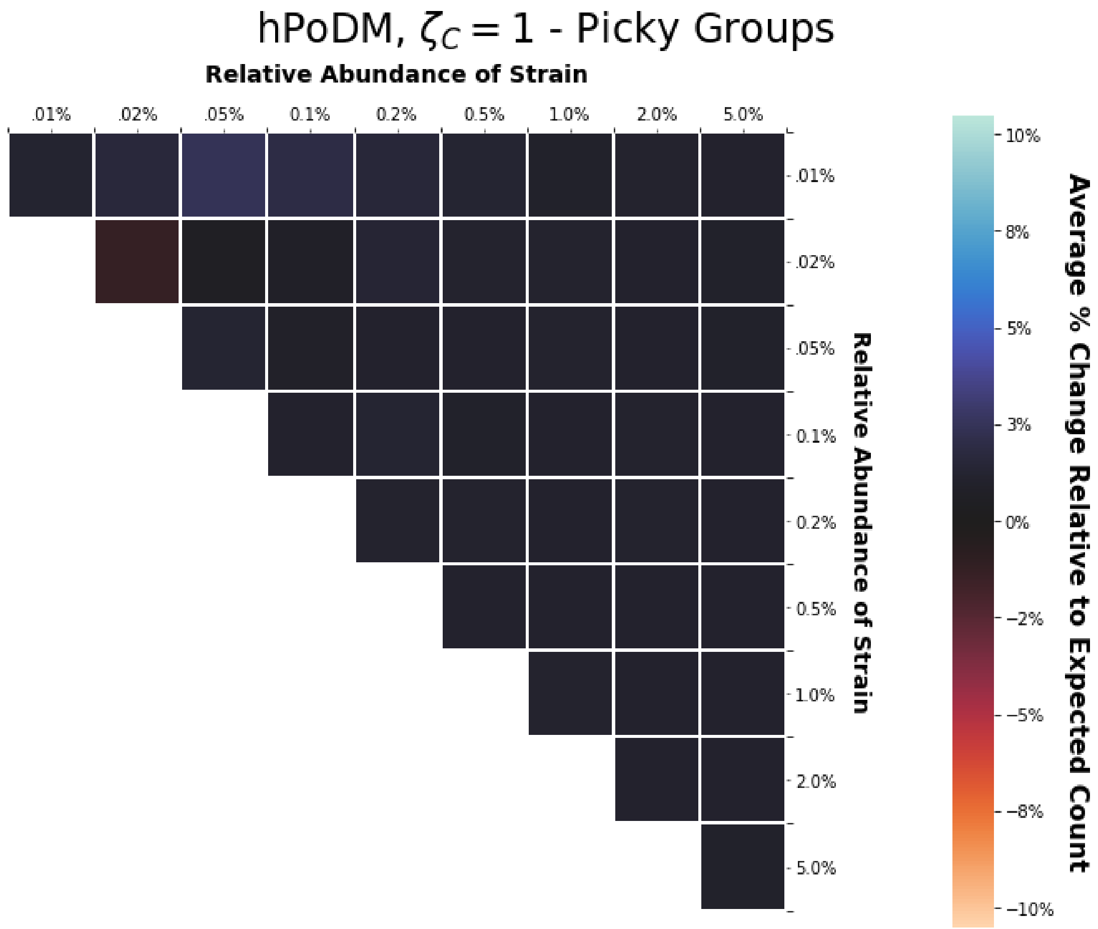

**Supplementary Figure S48**

**Increased Compositional Heterogeneity Only** In the absence of density heterogeneity, we see an increase in the number of droplets with one strain, with a smaller increase in the number of droplets with two strains. This is best evidenced in figure S37. This slight increase in the number of droplets with two strains causes the slight increase in the number of picky treatments shown in figures S48 and S49. As is clear also from figure S36, there is little change in the number of empty droplets with zero strains regardless of the level of compositional heterogeneity.

**Increased Compositional and Density Heterogeneity** In the presence of density heterogeneity, as seen in figure S37 the number of droplets with a single strain does still increase slightly. However, in contrast to what happens in the absence of density heterogeneity, in the presence of density heterogeneity there is an observable decrease in the number of droplets with two strains. While this trend is less obvious based only on figure S37, it is clear based on the observed decrease in the number of picky treatments visible in figures S50 S51. Nevertheless it is also clear from figure S37 and from a comparison of figures S51 and S47 (and S50 and S46) that the decrease in the number of droplets with two strains is not as pronounced (at least in relative terms) as the decrease in the number of droplets with three or more strains. Also in contrast to what happens in the absence of density heterogeneity, in the presence of density heterogeneity figures S36 and S37 show a marked increase in the number of empty droplets with zero strains as the compositional heterogeneity increases.

#### S4.5 Discussion

Section S4.5.1 discusses how the interpretation of the results from section S4.4 depends fundamentally on our opinion of which droplets constitute "good data". Section S4.5.2 discusses how the results from section S4.4 indicate what the next step is to figure out how to recover true interaction networks from

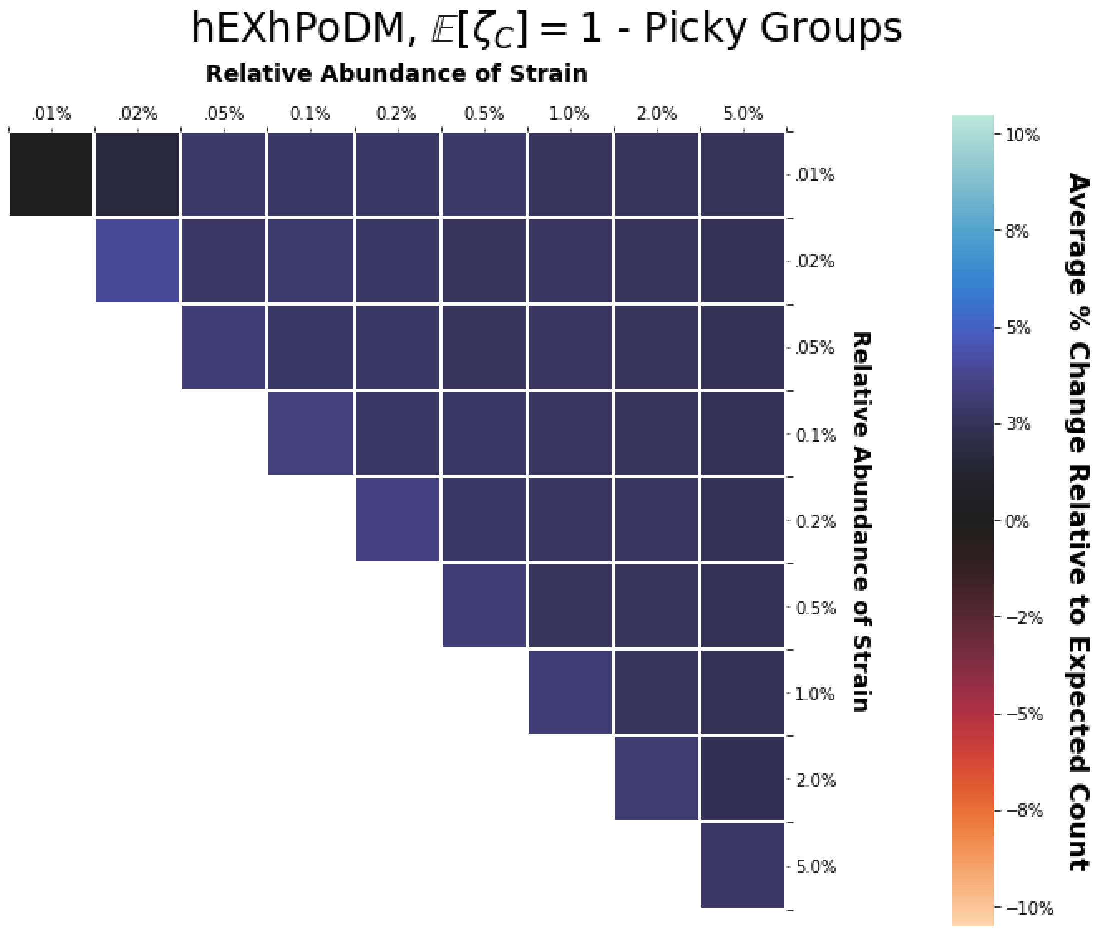

**Supplementary Figure S49**

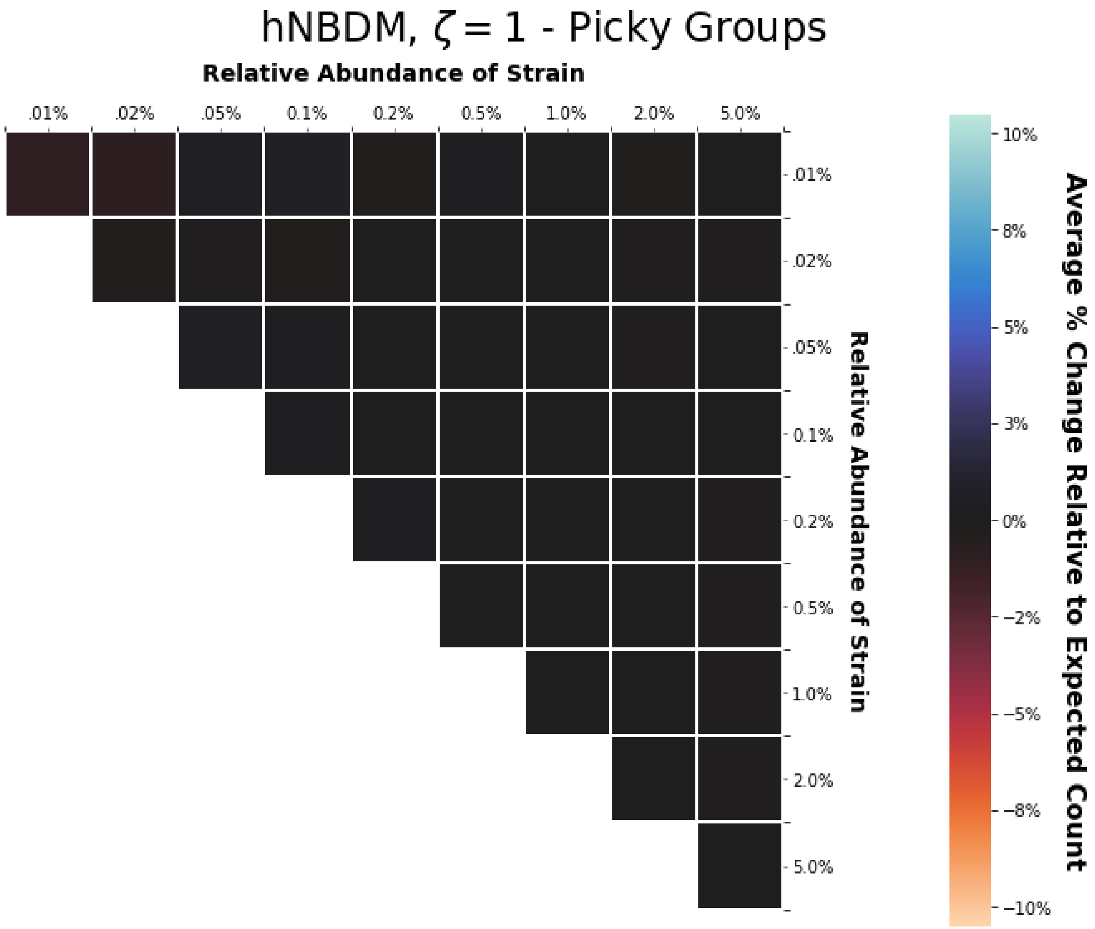

**Supplementary Figure S50**

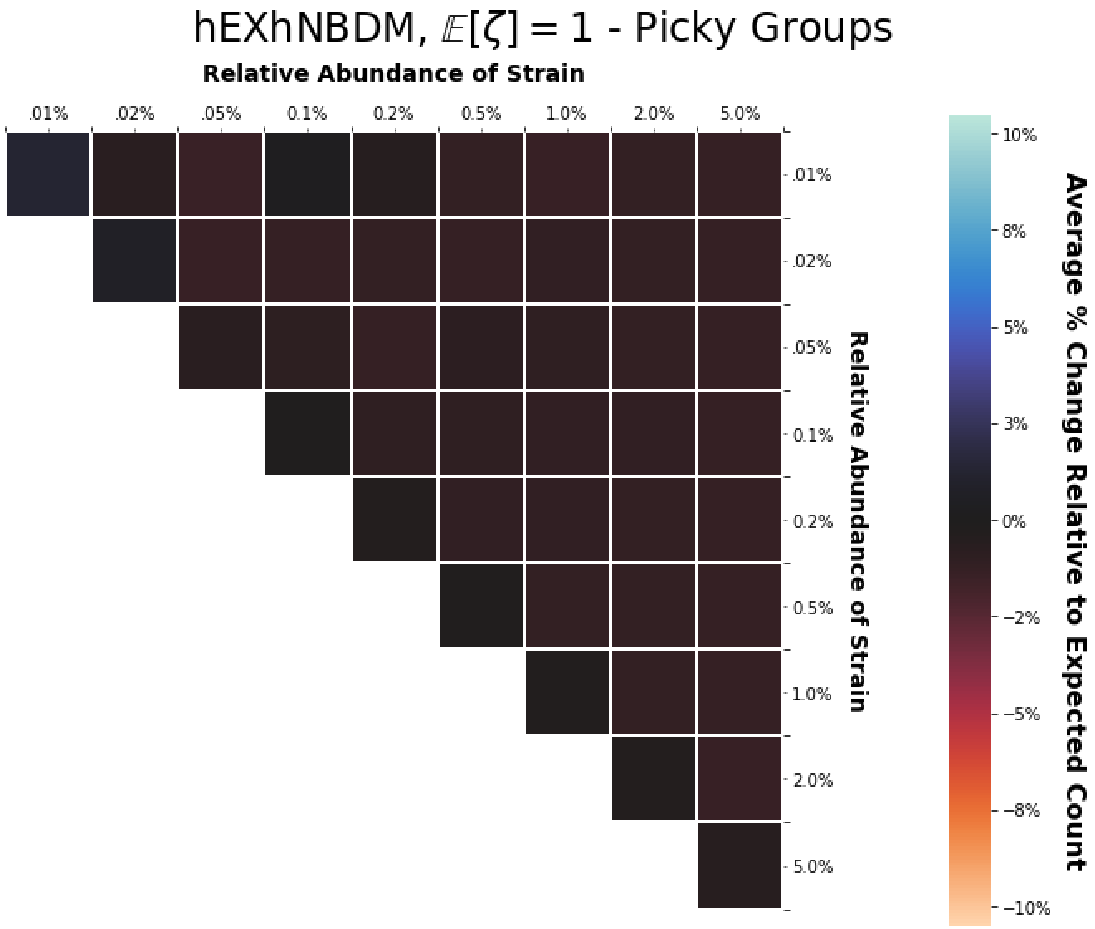

**Supplementary Figure S51**

MOREI data.

###### S4.5.1 The Effect of Heterogeneity on Data Throughput is Equivocal

The general effect that heterogeneities will have on the data throughput of MOREI is not straightforward. The answer depends on the particular combination of density heterogeneity and compositional heterogeneity present.

Given both density and compositional heterogeneity, it is clear that the effect on data throughput is negative. Both the numbers of picky treatments and gluttonous treatments decrease.

However, in the presence of only compositional heterogeneity but no density heterogeneity, whether the effect on data throughput is positive depends primarily on our opinion of what is useful data. If our primary concern is avoiding "data starvation" and being able to say anything about the interactions of the rarest strains, then the strong decrease in the number of gluttonous treatment groups clearly means that the effect of heterogeneity is negative. On the other hand, if we consider only the data from the picky treatments useful, then in the absence of density heterogeneity the effect of compositional heterogeneity is slightly positive.

###### S4.5.2 Next Steps

What is also clear is that, if we wish to be able to predict generally what effects heterogeneities may have on the data throughput of MOREI in practice, we must be able to estimate both density heterogeneity and compositional heterogeneity from empirical data. This is the topic of section S5. Given such estimates from real-world data, we can get some sense for the amount of either type of heterogeneity one is likely to encounter in practice. That makes it possible to create more accurate simulations. More accurate simulations in turn better assess which inference methods are most capable of recovering true interaction networks from MOREI data.

#### S5 ESTIMATORS FOR DENSITY AND COMPOSITIONAL HETEROGENEITIES

Section S5.1 explains the motivation for studying this estimation problem. Section S5.2 goes over technical details related to defining the estimators. Section S5.3 describes some implementation details of the analyses. Section S5.4 explains the results of the analyses. Finally section S5.5 interprets the results and explains how they are relevant to modeling the initial formation of droplets.

##### S5.1 Statistical Motivation

We want to restore control to the scientist about which questions they can answer using MOREI data by accounting for the effects of the randomness of the initial droplet formation process. To account for these effects we need to make realistic predictions about what these effects will be. I showed in previous sections that different statistical models for the initial droplet formation process predict substantially different effects. Therefore, to have any hope of making realistic predictions about what these effects will be, we need to be able to evaluate which of these statistical models is most realistic.

To evaluate which of these statistical models is most realistic, we need to estimate the parameters of the ghNBDM family from real data. We need to show that estimation of these parameters from MOREI data is even theoretically possible, and we need to develop estimators for these parameters whose effectiveness we can demonstrate in practice. Those are the goals of this section.

##### S5.2 Preliminaries

Using the hierarchical count-categorical framework defined in section S2.2, I treat the estimation of the density concentration $\zeta_D$ and compositional concentration $\zeta_C$ parameters of the ghNBDM family (cf. section S2.8) as two separate estimation problems. The first seeks to estimate the density concentration $\zeta_D$ from the empirical count distribution. The second seeks to estimate the compositional concentration $\zeta_C$ from the empirical categorical distributions[25].

For both estimation problems, I employ two strategies: (1) plugin ("method of moments") estimation, and (2) maximum likelihood ("ML") estimation. Both approaches give statistically consistent and asymptotically normal estimators. The maximum likelihood strategy has the advantage of being more efficient and asymptotically optimal (in the sense of satisfying the Cramer-Rao lower bound). The plugin

[25] Distribution*s* (plural) because again technically speaking there is a different categorical distribution for each possible value of the count distribution.

strategy has the advantage of being computationally simpler (and thus also easier for non-specialists to implement). I argue in sections S5.2.1, S5.2.3, and S5.5.2 that the plugin strategy also has the advantage of being "semi-parametric"[26], at least in the sense that its estimand is still straightforward to interpret even under model misspecification, although this is debatable.

Sections S5.2.1 and S5.2.2 discuss the estimation problem corresponding to the count distributions and the density concentration parameter $\zeta_D$. This is formulated as a parametric estimation problem involving the negative binomial distribution. Section S5.2.1 provides details of the plugin strategy, while section S5.2.2 provides details of the ML strategy.

Sections S5.2.3 and S5.2.4 discuss the estimation problem corresponding to the categorical distributions and the compositional concentration parameter $\zeta_C$. This is formulated as a univariate parametric estimation problem involving the Dirichlet-Multinomial distribution. The remaining nuisance parameters (the strain frequencies) can be assumed to be either already known or estimated separately (cf. section S5.6). Section S5.2.3 provides details of the plugin strategy, while section S5.2.4 provides details of the ML strategy.

#### Notation

The notation $\mathbb{I}\{A\}$ denotes the indicator function for the event $A$. Given any estimand which is a conditional expectation, i.e. of the form $\mathbb{E}[X \mid A]$ for some random variable $X$ and some event $A$ in the underlying sigma-algebra, the corresponding empirical (arithmetic) mean is denoted

$$\hat{\mu}[X \mid A] := \frac{\sum_{d \in [D]} X_d \mathbb{I}\{A\}}{\sum_{d \in [D]} \mathbb{I}\{A\}} . \tag{S97}$$

In particular the unconditional empirical mean is

$$\hat{\mu}[X] := \frac{1}{D} \sum_{d \in [D]} X_d . \tag{S98}$$

A conditional variance estimand $\mathrm{Var}[X \mid A]$ has a corresponding empirical conditional variance:

$$\hat{\mathrm{var}}[X \mid A] := \frac{\sum_{d \in [D]} (X_d - \hat{\mu}[X \mid A])^2 \mathbb{I}\{A\}}{\sum_{d \in [D]} \mathbb{I}\{A\}} , \tag{S99}$$

and an unconditional variance estimand $\mathrm{Var}[X]$ has the empirical counterpart:

$$\hat{\mathrm{var}}[X] := \frac{1}{D} \sum_{d \in [D]} (X_d - \hat{\mu}[X])^2 . \tag{S100}$$

Generalizing the above, for a conditional *co*variance estimand $\mathrm{Cov}[X, Y \mid A]$ its empirical counterpart is denoted and defined as

$$\hat{\mathrm{cov}}[X, Y \mid A] := \frac{\sum_{d \in [D]} (X_d - \hat{\mu}[X \mid A])(Y_d - \hat{\mu}[Y \mid A]) \mathbb{I}\{A\}}{\sum_{d \in [D]} \mathbb{I}\{A\}} , \tag{S101}$$

and likewise for an unconditional covariance estimand $\mathrm{Cov}[X, Y]$ and its empirical counterpart:

$$\hat{\mathrm{cov}}[X, Y] := \frac{1}{D} \sum_{d \in [D]} (X_d - \hat{\mu}[X])(Y_d - \hat{\mu}[Y]) . \tag{S102}$$

[26] Section 4 of (Nakashima, 1997) argues (at least for density concentration) that the plugin estimator is also "semi-parametric" when compared to the MLE due to having less bias and standard error. This is because the plugin estimator retains consistency (and supposedly also efficiency) for any distribution with the same overdispersion relative to the Poisson while the ML estimator does not. Cf. sections S2.4.4 and S2.7.1 of this work with formula (2.3) and section 4 of (Nakashima, 1997). This work does not investigate the estimators' behavior under model misspecification enough to evaluate this robustness claim. The true values of the effective density concentration (cf. section S5.2.1) and the effective compositional concentration (cf. section S5.2.3) are never computed for the misspecifed models. Moreover the misspecified models (hExhPoDM and hExhNBDM) are only slight modifications of the ghNBDM family. An investigation of the estimators' behavior under model misspecification thorough enough to make a legitimate and informed judgment about this issue is a worthwhile subject for future work.

##### *S5.2.1 Plugin Estimator for Density Concentration*

Section S5.2.1 derives the formula for the plugin estimator. Section S5.2.1 discusses how the plugin strategy implies an estimand that is defined for count distributions more general than the negative binomial distribution alone.

**Derivation of Plugin Estimator for Density Concentration** Recall that when $N(0)$ is negative binomial distributed[27], the variance equals

$$\mathrm{Var}[N(0)] = \lambda \left(1 + \frac{\lambda}{\zeta_D S}\right), \tag{S103}$$

while the variance that would have been expected under the Poisson distribution equals

$$\mathbb{E}[N(0)] = \lambda\,. \tag{S104}$$

Thus when $N(0)$ is negative binomial distributed its over-dispersion relative to the Poisson distribution equals

$$\frac{\mathrm{Var}[N(0)] - \mathbb{E}[N(0)]}{\mathbb{E}[N(0)]} = \frac{\lambda}{\zeta_D S} = \frac{\mathbb{E}[N(0)]}{\zeta_D S}\,. \tag{S105}$$

Solving for $\zeta_D$ leads to

$$\zeta_D = \frac{1}{S} \cdot \frac{(\mathbb{E}[N(0)])^2}{\mathrm{Var}[N(0)] - \mathbb{E}[N(0)]}\,. \tag{S106}$$

The above relationship (S106) motivates the following plugin estimator for $\zeta_D$:

$$\hat{\zeta_D} := \begin{cases} \infty & \widehat{\mathrm{var}}[N(0)] \le \hat{\mu}[N(0)] \\ \frac{1}{S}\frac{(\hat{\mu}[N(0)])^2}{\widehat{\mathrm{var}}[N(0)] - \hat{\mu}[N(0)]} & \text{else} \end{cases}\,. \tag{S107}$$

Cf. the derivation of the analogous equation (3.10) in (Anscombe, 1950).

**Effective Density Concentration** If one defines for any possible distribution of $N(0)$ with finite variance (e.g. including the Poisson) the "effective density concentration" as

$$\zeta_{D_{eff}} := \begin{cases} \infty & \mathrm{Var}[N(0)] \le \mathbb{E}[N(0)] \\ \frac{1}{S}\frac{(\mathbb{E}[N(0)])^2}{\mathrm{Var}[N(0)] - \mathbb{E}[N(0)]} & \text{else} \end{cases}, \tag{S108}$$

then the above plugin estimator (S107) is always consistent for the "effective density concentration" $\zeta_{D_{eff}}$. Consistency follows from the weak law of large numbers and the continuous mapping theorem for convergence in probability.

The effective density concentration $\zeta_{D_{eff}}$ being infinite merely flags that the distribution of $N(0)$ is not over-dispersed relative to the Poisson. An obvious limitation of this is that provides no quantification of the extent to which the distribution might be *under*-dispersed relative to the Poisson. Such under-dispersed distributions are not necessarily "exotic", consider for example the distribution of $Y = \frac{1}{2}X$ when $X \sim$ Pois. Then $\mathrm{Var}[Y] = \left(\frac{1}{2}\right)^2 \mathrm{Var}[X] = \frac{1}{4}\mathbb{E}[X] < \frac{1}{2}\mathbb{E}[X] = \mathbb{E}[Y]$. This of course works also for $pX$ for any $0 < p < 1$. Considering the possible ramifications of under-dispersion is left to future work.

When the distribution of $N(0)$ *is* over-dispersed relative to the Poisson, the effective density concentration measures the over-dispersion on a normalized and inverted scale. Because the effective density concentration $\zeta_{D_{eff}}$ equals the density concentration parameter $\zeta_D$ whenever $N(0)$ is negative binomial distributed, the estimand $\zeta_{D_{eff}}$ may be considered a generalization of the parameter $\zeta_D$ to other distributions. This is similar to the argument made in section 4 of (Nakashima, 1997) that the negative binomial plugin estimator[28] is only "semi-parametric".

---

[27] Using the parametrization for the negative binomial described in section S2.7.1.

[28] (Nakashima, 1997) considers a reparameterized version of the negative binomial distribution such that the corresponding plugin estimator is the reciprocal of that considered herein. This would correspond to an estimand that is the reciprocal of the effective density concentration.

##### S5.2.2 Maximum Likelihood Estimator for Density Concentration

This section explains how to use the maximum likelihood strategy to estimate the density concentration $\zeta_D$. Section S5.2.2 computes the relevant likelihood and log likelihood for the count distributions, while section S5.2.2 computes the score. Section S5.2.2 uses those results to derive the maximum likelihood estimators for the negative binomial distribution. Sections S5.2.2 and S5.2.2 survey what is already known about this estimator from previous literature.

**Negative Binomial (Log) Likelihood** Starting from equation (S28),

- if we assume for mathematical convenience that the count distributions of all droplets are mutually independent,
- using the identity[29] $\frac{\Gamma(x+M)}{\Gamma(x)} = \prod_{m=0}^{M-1}(x+m)$ for all positive integers $M$,

then the following is the full likelihood for a batch of $D$ droplets:

$$\mathcal{L}(\lambda, \zeta_D) = \prod_{d \in [D]} \left[ \left( \prod_{\nu=0}^{n_d - 1} (\zeta_D S + \nu) \right) \cdot \left(1 + \frac{\lambda}{\zeta_D S}\right)^{-\zeta_D S} \cdot \left(\frac{1}{\zeta_D S + \lambda}\right)^{n_d} \cdot \frac{\lambda^{n_d}}{n_d!} \right]. \tag{S109}$$

(Recall that $[D] := \{1, \dots, D\}$.) Thus this is the log likelihood:

$$\begin{aligned} \ell(\lambda, \zeta_D) &:= \\ \log\left(\mathcal{L}(\lambda, \zeta_D)\right) &= \sum_{d \in [D]} \left[ \sum_{\nu=0}^{n_d - 1} \log\left(\zeta_D S + \nu\right) \right] - D\zeta_D S \log\left(1 + \frac{\lambda}{\zeta_D S}\right) + \\ & D\overline{n} \log\left(\frac{1}{\zeta_D S + \lambda}\right) + D\overline{n}\log(\lambda) - \sum_{d \in [D]} \log\left(n_d!\right), \end{aligned} \tag{S110}$$

where I have defined $\overline{n} := \frac{1}{D}\sum_{d \in [D]} n_d$. (Each variable $n_d$ corresponds to a possible value of the random variable $N_d(0)$, the number of cells in droplet $d$.)

**Negative Binomial Score** It follows that the score with respect to $\lambda$ is

$$\frac{\partial \ell(\lambda, \zeta_D)}{\partial \lambda} = -\frac{D\zeta_D S}{\zeta_D S + \lambda} + \frac{\overline{n}}{\lambda} \cdot \left(\frac{D\zeta_D S}{\zeta_D S + \lambda}\right). \tag{S111}$$

Similarly, from (very) tedious calculus we get the following expression for the score with respect to $\zeta_D$:

$$\begin{aligned} \frac{\partial \ell(\lambda, \zeta_D)}{\partial \zeta_D} &= \sum_{d \in [D]} \left[ \sum_{\nu=0}^{n_d - 1} \frac{S}{\zeta_D S + \nu} \right] + \frac{DS\lambda}{\zeta_D S + \lambda} - \\ & DS \log\left(1 + \frac{\lambda}{\zeta_D S}\right) - \frac{DS\overline{n}}{\zeta_D S + \lambda}. \end{aligned} \tag{S112}$$

**Negative Binomial Maximum Likelihood Estimator** Setting $\frac{\partial \ell(\lambda, \zeta_D)}{\partial \lambda} = 0$, equation (S111) gives us that the maximum likelihood estimator $\hat{\lambda}$ for $\lambda$ is $\hat{\lambda} = \overline{n}$. This means that the plugin and maximum likelihood estimators for $\lambda$ coincide.

Setting $\frac{\partial \ell(\lambda, \zeta_D)}{\partial \zeta_D} = 0$, and substituting into (S112) the relationship $\lambda = \hat{\lambda} = \overline{n}$, we get that, if the maximum likelihood estimator $\hat{\zeta}_D$ of $\zeta_D$ exists, then it must be a solution of the following equation:

$$D \log\left(1 + \frac{\overline{n}}{\hat{\zeta}_D S}\right) = \sum_{d \in [D]} \left[ \sum_{\nu=0}^{n_d - 1} \frac{1}{\hat{\zeta}_D S + \nu} \right]. \tag{S113}$$

Rearranging terms for the purposes of computational efficiency, the equation (S113) above can be rewritten

$$D \log\left(1 + \frac{\overline{n}}{\hat{\zeta}_D S}\right) = \sum_{m=1}^{M} \frac{D_{\geq m}}{\hat{\zeta}_D S + (m-1)}, \tag{S114}$$

where $M := \max_{d \in [D]} N_d(0)$ and $D_{\geq m} := |\{d : N_d(0) \geq m\}|$.

[29]This follows from how $\Gamma$ interpolates the factorials, i.e. $\Gamma(x+1) = x\Gamma(x)$.

**Existence, Uniqueness, and Finiteness Conditions** It is believed(Anscombe, 1950)(Willson et al., 1984)(Willson et al., 1986) that a solution to the above equation (S114) exists if and only if $\hat{\text{var}}[N(0)] > \hat{\mu}[N(0)]$ (i.e. if and only if the empirical distribution is over-dispersed with respect to the Poisson), and that if a solution exists it is always unique. A purported[30] proof of these claims is given in (Aragón et al., 1992).

Note how the above purported conditions for the existence and uniqueness of the negative binomial MLE are the same as the conditions for the finiteness of the plugin estimator. When the MLE does not exist, it is because the likelihood is increasing but has no maximum. Therefore in those situations it makes sense to define the MLE to be infinite (cf. (Anscombe, 1950)). Thus the plugin estimator and MLE agree on when the estimate should be finite or infinite. Just like for the plugin estimator, the MLE being infinite has the interpretation that the Poisson distribution fits the data better than any negative binomial distribution.

**Estimation Difficulties** The two parameter (i.e. both $\lambda$ and $\zeta_D$ unknown) estimation problem for the negative binomial may be intrinsically difficult due to there being no complete sufficient statistic, corresponding to how the two parameter negative binomial family is not an exponential family (Willson et al., 1986).

At least for (relatively) small samples (Bowman, 1984)(Clark and Perry, 1989)(Piegorsch, 1990)(Lloyd-Smith, 2007), previous work has raised the concern that the MLE has unusually large bias (Shenton and Wallington, 1962) (Willson et al., 1984)(Bowman, 1984)(Piegorsch, 1990)(Lloyd-Smith, 2007). However at least one source (Piegorsch, 1990) conjectures that the MLE becomes a more viable option for large sample sizes.

The maximum likelihood estimation of $\zeta_D$ is believed to be particularly difficult for "small" values of $\lambda$ and large values of $\zeta_D$ (Shenton and Wallington, 1962)(Willson et al., 1984)(Bowman, 1984)(Lloyd-Smith, 2007). This may be due to the contours of the likelihood function being relatively "flat" or slowly increasing with respect to $\zeta_D$(Willson et al., 1986). However, other work (Clark and Perry, 1989) has found that the small $\lambda$ and large $\zeta_D$ regime is also the most difficult for other kinds of estimators of $\zeta_D$. (Bowman, 1984) found that $\hat{\text{var}}[N(0)] \leq \hat{\mu}[N(0)]$ occurred most often in the small $\lambda$, large $\zeta_D$ regime, a condition which necessitates infinite estimates for both the plugin and ML estimators (cf. section S5.2.2).

Because of this, some authors have suggested that the MLE does not perform much better than the corresponding plugin estimator (despite having lower variance) (Anscombe, 1950)(Shenton and Wallington, 1962)(Willson et al., 1984) and thus that the plugin estimator should be preferred due to its greater computational simplicity (Anscombe, 1950)(Willson et al., 1984)(Yu et al., 2013).

Other authors have suggested instead reparameterizing the negative binomial in terms of $\frac{1}{\zeta_D S}$ (Ross and Preece, 1985)(Clark and Perry, 1989)(Piegorsch, 1990)(Nakashima, 1997)(Lloyd-Smith, 2007). This makes sense because the difficulties occur mostly for large values of $\zeta_D$ for which the resulting distributions are all very similar to the corresponding Poisson(Bowman, 1984)(Lloyd-Smith, 2007) and because this removes the need for infinite estimates (Clark and Perry, 1989)(Lloyd-Smith, 2007). (Ross and Preece, 1985) argue that reparameterizing the negative binomial using $\frac{1}{\zeta_D S}$ allows a continuous transition between the negative binomial, which is over-dispersed relative to the Poisson, and the ("positive") binomial, which is under-dispersed relative to the Poisson. Nevertheless even the reparameterized MLE was not always necessarily found to behave much better than the reparameterized plugin estimator (Piegorsch, 1990).

Exact formulae for the asymptotic variance of both the plugin and ML estimators were given in (Anscombe, 1950), and exact formulae for the asymptotic bias of both the plugin and ML estimators were given in (Shenton and Wallington, 1962). ((Nakashima, 1997) gives exact formulae for the asymptotic variances of the $\frac{1}{\zeta_D S}$ reparameterized versions of the plugin and ML estimators.) Because the asymptotic formulae of the plugin and ML estimators have "the same general structure"(Shenton and Wallington, 1962) for both the bias and variance[31], the larger sample sizes typical for this problem may lead to the performance of the MLE and plugin estimator being comparable.

[30]The proof is probably correct, but I have inspected it only cursorily and not in detail. Therefore I feel I cannot legitimately attest to its validity firsthand.

[31]Cf. Fig.3 of (Shenton and Wallington, 1962) and equations 3.6 and 3.11 of (Anscombe, 1950).

##### S5.2.3 Plugin Estimator for Compositional Concentration

Section S5.2.3 outlines the big picture ideas behind the plugin estimator's somewhat complicated derivation. Section S5.2.3 provides the details behind most of the derivation. Section S5.2.3 discusses a possible ambiguity and choices for resolving it. Section S5.2.3 gives the formal definition of the final plugin estimator used for all later analyses. Section S5.2.3 discusses how the plugin strategy implies an estimand that is defined for categorical distributions more general than the Dirichlet-Multinomial distribution alone.

**Overview of Derivation of Plugin Estimator for Compositional Concentration** The general idea is as follows:

- For each number of cells $n \geq 2$ and each tuple of strains $(s_1, s_2)$ get a plugin estimator for $\frac{1}{1+\zeta_C}$.
- Take a weighted average to get a single plugin estimator for $\frac{1}{1+\zeta_C}$.
- Solve to get a plugin estimator for $\zeta_C$.

The motivation for averaging before solving for $\zeta_C$ in the expression $\frac{1}{1+\zeta_C S}$ is because I anticipate the former operation to be "better behaved" than the reciprocals involved in the latter. Solving after averaging makes it possible to take reciprocals only once. This hopefully leads to a plugin estimator for $\zeta_C$ "better behaved" than plugin estimators derived by solving (multiple times) first and then averaging. Nevertheless the latter in principle is possible too.

Note that when $n = 1$, the Dirichlet-Multinomial distribution reduces completely to the corresponding Multinomial ("Multinoulli") distribution and any compositional concentration parameter is completely unidentifiable. (This corresponds to how $n - 1 = 0$ in the over-dispersion formula, but follows ultimately from the probability mass function of the Dirichlet-Multinomial distribution.)

**Solving for Compositional Concentration Parameter** Whenever $\mathbb{E}[\vec{\mathbf{N}}(0) \mid N(0)]$ is Dirichlet-Multinomial distributed, for all fixed $n \geq 2$ one has, it follows from (S49) that for any strain $s$:

$$\frac{1}{1+\zeta_C S} = \frac{\operatorname{Var}[N^{(s)}(0) \mid N(0) = n] - nf^{(s)}(1-f^{(s)})}{n(n-1)f^{(s)}(1-f^{(s)})}. \tag{S115}$$

Similarly, it follows from (S51) that for any pair of strains $s_1$, $s_2$:

$$\frac{1}{1+\zeta_C S} = \frac{-\operatorname{Cov}[N^{(s_1)}(0), N^{(s_2)}(0) \mid N(0) = n] - nf^{(s_1)}f^{(s_2)}}{n(n-1)f^{(s_1)}f^{(s_2)}}. \tag{S116}$$

Therefore, for any convex combination $c_\sigma$, $c_{\sigma_1,\sigma_2}$ of the right hand sides of (S115) and (S116) one has

$$\begin{aligned}\frac{1}{1+\zeta_C S} =& \frac{1}{n(n-1)}\left[\sum_{\sigma\in[S]} c_\sigma \left[\frac{\operatorname{Var}[N^{(\sigma)}(0)] - nf^{(\sigma)}(1-f^{(\sigma)})}{f^{(\sigma)}(1-f^{(\sigma)})}\right]\right.\\ &\left.+\sum_{\sigma_1\in[S]}\sum_{\sigma_2\neq\sigma_1} c_{\sigma_1,\sigma_2}\left[\frac{-\operatorname{Cov}[N^{(\sigma_1)}(0), N^{(\sigma_2)}(0)] - nf^{(\sigma_1)}f^{(\sigma_2)}}{f^{(\sigma_1)}f^{(\sigma_2)}}\right]\right].\end{aligned} \tag{S117}$$

Therefore one has further that

$$\begin{aligned}\frac{1}{1+\zeta_C S} =& \sum_{n\geq 2}\left(\left[\frac{1}{n(n-1)}\left[\sum_{\sigma\in[S]} c_\sigma\left[\frac{\operatorname{Var}[N^{(\sigma)}(0) \mid N(0)=n] - nf^{(\sigma)}(1-f^{(\sigma)})}{f^{(\sigma)}(1-f^{(\sigma)})}\right]\right.\right.\right.\\ &\left.\left.+\sum_{\sigma_1\in[S]}\sum_{\sigma_2\neq\sigma_1} c_{\sigma_1,\sigma_2}\left[\frac{-\operatorname{Cov}[N^{(\sigma_1)}(0), N^{(\sigma_2)}(0) \mid N(0)=n] - nf^{(\sigma_1)}f^{(\sigma_2)}}{f^{(\sigma_1)}f^{(\sigma_2)}}\right]\right]\right]\\ &\left.\cdot\,\mathbb{P}(N(0) = n \mid N(0) \geq 2)\right),\end{aligned} \tag{S118}$$

allowing one to solve for the compositional concentration parameter

$$\zeta_C = \frac{1}{S}\left(\left(\sum_{n\geq 2}\left(\left[\frac{1}{n(n-1)}\left[\sum_{\sigma\in[S]} c_\sigma\left[\frac{\mathrm{Var}[N^{(\sigma)}(0)\mid N(0)=n]-nf^{(\sigma)}(1-f^{(\sigma)})}{f^{(\sigma)}(1-f^{(\sigma)})}\right]\right.\right.\right.\right.\right.$$
$$\left.\left.\left. + \sum_{\sigma_1\in[S]}\sum_{\sigma_2\neq\sigma_1} c_{\sigma_1,\sigma_2}\left[\frac{-\mathrm{Cov}[N^{(\sigma_1)}(0),N^{(\sigma_2)}(0)\mid N(0)=n]-nf^{(\sigma_1)}f^{(\sigma_2)}}{f^{(\sigma_1)}f^{(\sigma_2)}}\right]\right]\right] \tag{S119}$$
$$\left.\left.\left.\cdot\,\mathbb{P}(N(0)=n\mid N(0)\geq 2)\right)\right)^{-1}-1\right).$$

**Choice of Convex Coefficients** Choosing $c_\sigma, c_{\sigma_1,\sigma_2} = \frac{1}{S^2}$ for all strains (and pairs thereof) corresponds to the $L_2$ projection of the covariance matrices onto the vector space of matrices whose entries all equal a common value. Thus the corresponding plugin estimator might initially appear attractive as a ”least-squares” estimator.

However, the corresponding plugin estimator can have an extremely slow rate of convergence, e.g. when there are many ”rare” strains. Preliminary investigations (data not shown) found that the corresponding plugin estimator effectively failed to converge towards the true value at all even after sampling 15 million droplets.

A more pragmatic strategy is to choose the weights $c_\sigma, c_{\sigma_1,\sigma_2}$ such that estimates with the most data behind them also receive the most weight. This means assigning higher weights to (pairs of) more common strains. Using the choices $c_\sigma = (f^{(\sigma)})^2$ and $c_{\sigma_1,\sigma_2} = f^{(\sigma_1)}f^{(\sigma_2)}$, the resulting formula for $\zeta_C$ is

$$\zeta_C = \frac{1}{S}\left(\left(\sum_{n\geq 2}\left(\left[\frac{1}{n(n-1)}\left[\sum_{\sigma\in[S]} (f^{(\sigma)})^2\left[\frac{\mathrm{Var}[N^{(\sigma)}(0)\mid N(0)=n]-nf^{(\sigma)}(1-f^{(\sigma)})}{f^{(\sigma)}(1-f^{(\sigma)})}\right]\right.\right.\right.\right.\right.$$
$$\left.\left.\left. + \sum_{\sigma_1\in[S]}\sum_{\sigma_2\neq\sigma_1} f^{(\sigma_1)}f^{(\sigma_2)}\left[\frac{-\mathrm{Cov}[N^{(\sigma_1)}(0),N^{(\sigma_2)}(0)\mid N(0)=n]-nf^{(\sigma_1)}f^{(\sigma_2)}}{f^{(\sigma_1)}f^{(\sigma_2)}}\right]\right]\right] \tag{S120}$$
$$\left.\left.\left.\cdot\,\mathbb{P}(N(0)=n\mid N(0)\geq 2)\right)\right)^{-1}-1\right).$$

The corresponding plugin estimator converges much faster to the true value $\zeta_C$ (see results section).

**Plugin Estimator Definition** The resulting plugin estimator for $\zeta_C$ is

$$\hat{\zeta}_{C\,final} = \begin{cases} \infty & \frac{1}{1+\hat{\zeta}_C S} \leq 0 \\ 0 & \frac{1}{1+\hat{\zeta}_C S} \geq 1 \\ \hat{\zeta}_C & \text{else} \end{cases} \tag{S121}$$

where $\hat{\zeta}_C$ is defined as

$$\hat{\zeta}_C := \frac{1}{S}\left(\left(\sum_{n\geq 2}\left(\left[\frac{1}{n(n-1)}\left[\sum_{\sigma\in[S]} (\hat{f}^{(\sigma)})^2\left[\frac{\widehat{\mathrm{var}}[N^{(\sigma)}(0)\mid N(0)=n]-n\hat{f}^{(\sigma)}(1-\hat{f}^{(\sigma)})}{\hat{f}^{(\sigma)}(1-\hat{f}^{(\sigma)})}\right]\right.\right.\right.\right.\right.$$
$$\left.\left.\left. + \sum_{\sigma_1\in[S]}\sum_{\sigma_2\neq\sigma_1} \hat{f}^{(\sigma_1)}\hat{f}^{(\sigma_2)}\left[\frac{-\widehat{\mathrm{cov}}[N^{(\sigma_1)}(0),N^{(\sigma_2)}(0)\mid N(0)=n]-n\hat{f}^{(\sigma_1)}\hat{f}^{(\sigma_2)}}{\hat{f}^{(\sigma_1)}\hat{f}^{(\sigma_2)}}\right]\right]\right] \tag{S122}$$
$$\left.\left.\left.\cdot\,\mathbb{P}(N(0)=n\mid N(0)\geq 2)\right)\right)^{-1}-1\right).$$

In the above definition, $\hat{f}^{(\sigma)}$ denote statistically consistent estimators of the frequencies $f^{(\sigma)}$. One possibile definition for such estimators (which was used for the implementation seen in section S5.4) is given in section S5.6. I did not want to assume that the scientist had, or wanted to rely on, a priori accurate estimates of the true frequency values. In principle, if such knowledge were available and trusted, then one would use those values for $\hat{f}^{(\sigma)}$ in equation S121.

Note that when $\frac{1}{1+\hat{\zeta}_C} > 1$, the result of solving for $\hat{\zeta}_C$ is less than 0. The interpretation in this case is extremely unclear given that the limit as $\zeta_C \to 0$ corresponds to "infinite heterogeneity". This is the reason for thresholding the values of $\hat{\zeta}_{C_{final}}$ below by 0.

**Effective Compositional Concentration** Using the chosen values of $c_\sigma$ and $c_{\sigma_1,\sigma_2}$ gives one possible choice for a definition of "effective compositional concentration", with the resulting estimand existing for any distribution of $\vec{\mathbf{N}}(0)$ with finite covariances, not just when $\mathbb{E}[\vec{\mathbf{N}}(0) \mid N(0)]$ is Dirichlet-Multinomial distributed:

$$(\zeta_{C_{eff}})_{final} = \begin{cases} \infty & \frac{1}{1+\zeta_{C_{eff}} S} \leq 0 \\ 0 & \frac{1}{1+\zeta_{C_{eff}} S} \geq 1 \\ \zeta_{C_{eff}} & \text{else} \end{cases} \tag{S123}$$

with $\zeta_{C_{eff}}$ defined (when $\frac{1}{1+\zeta_{C_{eff}} S} \neq 0$) as

$$\begin{aligned} \zeta_{C_{eff}} = \frac{1}{S} \Bigg( \Bigg( \sum_{n \geq 2} \Bigg( \Bigg[ \frac{1}{n(n-1)} \Bigg[ \sum_{\sigma \in [S]} (f^{(\sigma)})^2 \Bigg[ \frac{\operatorname{Var}[N^{(\sigma)}(0) \mid N(0)=n] - nf^{(\sigma)}(1-f^{(\sigma)})}{f^{(\sigma)}(1-f^{(\sigma)})} \Bigg] \\ + \sum_{\sigma_1 \in [S]} \sum_{\sigma_2 \neq \sigma_1} f^{(\sigma_1)} f^{(\sigma_2)} \Bigg[ \frac{-\operatorname{Cov}[N^{(\sigma_1)}(0), N^{(\sigma_2)}(0) \mid N(0)=n] - nf^{(\sigma_1)} f^{(\sigma_2)}}{f^{(\sigma_1)} f^{(\sigma_2)}} \Bigg] \Bigg] \Bigg] \\ \cdot \mathbb{P}(N(0)=n \mid N(0) \geq 2) \Bigg) \Bigg)^{-1} - 1 \Bigg), \end{aligned} \tag{S124}$$

The law of large numbers combined with the continuous mapping theorem for convergence in probability guarantees the consistency of the estimator (S121) for the effective compositional concentration $(\zeta_{C_{eff}})_{final}$ whenever, under the given distribution for $\vec{\mathbf{N}}(0)$, the chosen estimators $\hat{f}^{(\sigma)}$ are also consistent for the true frequencies. (Asymptotic consistency is actually guaranteed, due to the same reasons, for the analogous plugin estimator and estimand given any choice of convex coefficients $c_\sigma, c_{\sigma_1,\sigma_2}$. The result does *not* depend on the chosen definitions $c_\sigma := f^{(\sigma)}$, $c_{\sigma_1,\sigma_2} := f^{(\sigma_1)} f^{(\sigma_2)}$.)

Note also that $(\zeta_{C_{eff}})_{final}$ is thresholded below by 0 for the same reason that $\hat{\zeta}_{C_{final}}$ is thresholded below by 0.

Because the effective compositional concentration, as well as $\zeta_{C_{eff}}$, both equal the compositional concentration parameter $\zeta_C$ whenever $\mathbb{E}[\vec{\mathbf{N}}(0) \mid N(0)]$ is Dirichlet-Multinomial distributed, the estimand $(\zeta_{C_{eff}})_{final}$ may be considered a generalization of the parameter $\zeta_C$ to other distributions.

**Effective Compositional Concentration without Frequencies** There could be distributions which are not parameterized in terms of frequencies $f^{(s)}$ to which we might still want to apply the notion of effective compositional concentration. One strategy we could employ is based on the expectations of the multinomial and Dirichlet-Multinomial distributions. For the marginal distribution corresponding to any strain $s$, when $\vec{\mathbf{N}}(0)$ conditional on $N(0) = n$ is multinomial or Dirichlet-Multinomial distributed, then one always has

$$\mathbb{E}\Big[N^{(s)}(0) \,\Big|\, N(0) = n\Big] = nf^{(s)}. \tag{S125}$$

Rearranging the relationship from (S125) gives us a proxy which we can use in place of the frequency parameters $f^{(s)}$ (at least when $n \geq 1$):

$$f^{(s)} = \frac{\mathbb{E}\Big[N^{(s)}(0) \,\Big|\, N(0) = n\Big]}{n}, \tag{S126}$$

related to the so-called "effective frequency", see section S5.6 for a derivation. Note this has the drawback of being dependent on the value of $n$.

Using these proxies for the frequencies (S126), under the above distributional assumptions we can rewrite equation (S115) as

$$\frac{1}{1+\zeta_C S} = \frac{1}{n-1} \cdot \left[ \frac{n \operatorname{Var}[N^{(s)}(0) \mid N(0)=n] - \mathbb{E}\left[N^{(s)}(0) \,\middle|\, N(0)=n\right] \left(n - \mathbb{E}\left[N^{(s)}(0) \,\middle|\, N(0)=n\right]\right)}{\mathbb{E}\left[N^{(s)}(0) \mid N(0)=n\right] \left(n - \mathbb{E}\left[N^{(s)}(0) \mid N(0)=n\right]\right)} \right], \tag{S127}$$

and rewrite equation (S116) as

$$\frac{1}{1+\zeta_C S} = \frac{1}{n-1} \cdot \left[ \frac{-n \operatorname{Cov}[N^{(s_1)}(0), N^{(s_2)}(0) \mid N(0)=n] - \mathbb{E}\left[N^{(s_1)}(0) \,\middle|\, N(0)=n\right] \mathbb{E}\left[N^{(s_2)}(0) \,\middle|\, N(0)=n\right]}{\mathbb{E}\left[N^{(s_1)}(0) \mid N(0)=n\right] \mathbb{E}\left[N^{(s_2)}(0) \mid N(0)=n\right]} \right]. \tag{S128}$$

Based on this we can easily generalize the formulae from sections S5.2.3-S5.2.3 to define a notion of effective compositional concentration that does not require the distribution to have frequency parameters $f^{(s)}$. Again, cf. section S5.6.

I do not claim that this is the only possibly proxy one could use in place of the frequency parameters for more general distributions. However, it does have the benefit of being extremely widely applicable. Any definition of effective compositional concentration will (seemingly) require the distribution to have finite second (conditional) cumulants, which in turn guarantees the finiteness of the first (conditional) cumulants in formulae (S127) and (S128) above.

##### *S5.2.4 Maximum Likelihood Estimator for Compositional Concentration*

This section explains how to use the maximum likelihood strategy to estimate the compositional concentration $\zeta_C$. Section S5.2.4 computes the relevant likelihood and log likelihood for the categorical distributions, while section S5.2.4 computes the score. Section S5.2.4 uses those results to derive the maximum likelihood estimators for the Dirichlet-Multinomial distribution.

**Dirichlet-Multinomial (Log) Likelihood** Starting from equation (S21),

- if we assume for mathematical convenience that the distributions of all droplets are mutually independent,
- using the identity[32] $\frac{\Gamma(x+M)}{\Gamma(x)} = \prod_{m=0}^{M-1}(x+m)$ for all positive integers $M$,
- if we assume that the droplets are Dirichlet-Multinomial distributed conditional on their count distributions,

then this is the full likelihood for an entire batch of $D$ droplets:

$$\begin{aligned} \mathcal{L}(\vec{f}, \zeta_C) = \prod_{d\in[D]} \Bigg[ & \mathbb{P}(N_d(0) = n_d) \cdot \left( \prod_{\nu=0}^{n_d-1} (\zeta_C S + \nu)^{-1} \right) \cdot \\ & \prod_{s\in[S]} \left( \prod_{\nu=0}^{n_d^{(s)}-1} (\zeta_C S f^{(s)} + \nu) \right) \cdot \binom{n_d}{n_d^{(1)} \cdots n_d^{(S)}} \Bigg], \end{aligned} \tag{S129}$$

where (as a reminder) by definition $n_d = \sum_{s\in[S]} n_d^{(s)}$.

[32]Cf. footnote 29.

Therefore this is the full log likelihood:

$$\begin{aligned} &\ell(\vec{f},\zeta_C) := \\ &\log\left(\mathcal{L}(\vec{f},\zeta_C)\right) = \sum_{d\in[D]}\left[\log\left(\mathbb{P}(N_d(0)=n_d)\right) - \sum_{\nu=0}^{n_d-1}\log(\zeta_C S+\nu) + \right. \\ &\qquad \left. \sum_{s\in[S]}\left(\sum_{\nu=0}^{n_d^{(s)}-1}\log(\zeta_C S f^{(s)}+\nu)\right) + \log\left(\binom{n_d}{n_d^{(1)}\cdots n_d^{(S)}}\right)\right] . \end{aligned} \tag{S130}$$

**Dirichlet-Multinomial Score** Assuming that the count distribution has no dependence[33] on $\zeta_C$, it follows that the score with respect to $\zeta_C$ is

$$\frac{\partial\ell(\vec{f},\zeta_C)}{\partial\zeta_C} = \sum_{d\in[D]}\left[-\sum_{\nu=0}^{n_d-1}\frac{S}{\zeta_C S+\nu} + \sum_{s\in[S]}\left(\sum_{\nu=0}^{n_d^{(s)}-1}\frac{Sf^{(s)}}{\zeta_C S f^{(s)}+\nu}\right)\right] . \tag{S131}$$

Using the commutativity of addition (i.e. rearranging terms) for purposes of computational efficiency, the above may be rewritten

$$\frac{\partial\ell(\vec{f},\zeta_C)}{\partial\zeta_C} = -\sum_{m=1}^{M}\frac{D_{\geq m}S}{\zeta_C S+(m-1)} + \sum_{s\in[S]}\left(\sum_{m=1}^{M^{(s)}}\frac{D_{\geq m}^{(s)}Sf^{(s)}}{\zeta_C S f^{(s)}+(m-1)}\right) , \tag{S132}$$

where I have defined

$$\begin{aligned} M &:= \max_{d\in[D]} N_d(0) , \\ \forall s\in[S] \quad M^{(s)} &:= \max_{d\in[D]} N_d^{(s)}(0) , \\ D_{\geq m} &:= \left|\{d : N_d(0)\geq m\}\right| , \\ \forall s\in[S] \quad D_{\geq m}^{(s)} &:= \left|\left\{d : N_d^{(s)}(0)\geq m\right\}\right| . \end{aligned} \tag{S133}$$

This is basically the same approach to grouping terms as suggested in (Sklar, 2014).

**Dirichlet-Multinomial Maximum Likelihood Estimator** Assuming that the frequencies $\vec{f} := (f_1,\ldots,f^{(S)})$ are already known or estimated (and thus don't need to be treated as nuisance parameters), it follows from the above formula (S132) for the score that a maximum likelihood estimator $\hat{\zeta_C}$ for the compositional concentration $\zeta_C$ must be a solution of the following equation:

$$\sum_{m=1}^{M}\frac{D_{\geq m}}{\hat{\zeta_C}S+(m-1)} = \sum_{s\in S}\left(\sum_{m=1}^{M^{(s)}}\frac{D_{\geq m}^{(s)}f^{(s)}}{\hat{\zeta_C}Sf^{(s)}+(m-1)}\right) . \tag{S134}$$

I only claim that satisfying the above equation is necessary, not sufficient. In particular, I make no claims regarding the conditions under which a root for the above equation exists, nor regarding the conditions under which a root (if it exists) will be unique. These conditions appear to be unknown.

#### S5.3 Methods

Section S5.3.1 explains how the empirical distributions of the plugin heterogeneity estimators were computed as a function of batch size. Section S5.3.2 provides analogous details for the ML estimators. Section S5.3.3 explains how these results were depicted. Complete implementation details can be found in the code at the relevant GitLab repository. See https://gitlab.com/krinsman/droplets.

[33] Or ignoring such a dependence if it exists, e.g. as in the case of hNBDM, and choosing to consider the maximum *conditional* likelihood estimator instead of the MLE sensu stricto.

##### *S5.3.1 Computing Plugin Estimator Distributions*

For each of the seven simulated distributions, for each of the 500 simulations, for each of the three batch sizes (small = $10,000$ droplets, medium = $500,000$ droplets, large = $15,000,000$ droplets), using NumPy (Harris et al., 2020) version 1.20.2 I partitioned the simulation results into equal parts of the given batch size. Since each simulation contains $15,000,000$ droplets, this corresponds to $1,500$ small batches per simulation, 30 medium batches per simulation, and 1 large batch per simulation. Then for each of the batches, I evaluated the plugin density heterogeneity estimator (S107) and the plugin compositional heterogeneity estimator (S121) and stored the results.

##### *S5.3.2 Computing MLE Distributions*

Simulated droplets were partitioned into batches exactly as was done for the plugin estimators. Cf. section S5.3.1 above for details.

I tried to use Brent's method (Brent, 1973, Ch. 3-4) whenever possible to estimate roots of the score functions, because Powell's method (Powell, 1970) was often unstable. Preliminary investigations (data not shown) suggested that Powell's method often either failed to converge, or failed to converge to a reasonable value (for example returning a negative answer, or one that is far too large), under conditions where Brent's method converged to a reasonable answer similar to the truth. While it was always possible to supply a reasonable interval of values for Brent's method to restrict its search, the method requires the signs of the function to differ at both ends of the interval. It was difficult to systematically choose (i.e. without manual intervention) interval endpoints which would guarantee opposite signs at both ends of the interval for all possible simulated datasets. Therefore Powell's method often had to be used as an alternative, which may have introduced many inaccurate estimates. While gradient-based methods probably would have been more accurate and converged faster, this level of accuracy seemed sufficient for a proof of principle. Estimates that were less than zero were obviously erroneous and therefore discarded – this affected fewer than 3% of the estimates in general and usually less (data not shown).

**For the Negative Binomial (Density Concentration):**

- Check that the sample variance is larger than the sample mean, otherwise return $\infty$ and terminate.
- Set initial guess value for the root-finding algorithm to be the value of the plugin estimator[34], equation (S107).
- If the score function, computed using the terms of equation (S114) rearranged to one side to match (S112), had different signs at $10^{-4}$ and $10^{4}$:
  - Use Brent's method (Brent, 1973, Ch. 3-4) implemented in SciPy (Virtanen et al., 2020) with brackets $10^{-4}$ and $10^{4}$ to look for a root of the score function.
- If the score function, computed using the terms of equation (S114) rearranged to one side to match (S112), had the same sign at $10^{-4}$ and $10^{4}$:
  - Use Powell's hybrid method (Powell, 1970) as implemented in the HYBRD routine of MINPACK (Moré et al., 1980) (Cowell, 1984, Ch. 5) and made available through SciPy (Virtanen et al., 2020) to search for a root of the score function.
  - If the method fails to converge and the final guess is greater than $10^{4}$, return $\infty$.
  - Otherwise (even if the method fails to converge) return the final guess (or discard if the final guess is less than zero).

**For the Dirichlet-Multinomial (Compositional Concentration):**

- Set the initial guess value for the root-finding algorithm to be the value of the plugin estimator, equation (S121).
- If the plugin estimator was larger than $10^{6}$ (including $\infty$), set the initial guess value to be $0.5 \times 10^{6}$.

[34] This never exceeded 10,000 when the sample variance was larger than the sample mean.

- If the score function, computed using equation (S132), had different signs at $10^{-6}$ and $10^{6}$:
  - Use Brent's method (Brent, 1973, Ch. 3-4) implemented in SciPy (Virtanen et al., 2020) with brackets $10^{-6}$ and $10^{6}$ to look for a root of the score function.
- If the score function, computed using equation (S132), had the same sign at $10^{-6}$ and $10^{6}$:
  - Use Powell's hybrid method (Powell, 1970) as implemented in the HYBRD routine of MINPACK (Moré et al., 1980) (Cowell, 1984, Ch. 5) and made available through SciPy (Virtanen et al., 2020) to search for a root of the score function.
  - If the method fails to converge and the final guess is greater than $10^{6}$, return $\infty$.
  - Otherwise (even if the method fails to converge) return the final guess (or discard if the final guess is less than zero).

###### S5.3.3 Plotting Estimator Distributions

I plotted results using Matplotlib (Hunter, 2007) version 3.4.1 and Seaborn (Waskom, 2021) version 0.11.1. Kernel density estimates were made using the default Seaborn (Waskom, 2021) settings[35]. I computed empirical cumulative density functions using StatsModels (Seabold and Perktold, 2010) version 0.12.2, which essentially implements a wrapper for the binary search algorithm from NumPy (Harris et al., 2020). I then computed the empirical survival functions from the empirical cumulative density functions.

##### S5.4 Results

Section S5.4.1 explains what indicates that the plugin estimators are consistent in practice when the concentration parameters are infinite. Section S5.4.2 explains the analogous evidence when the concentration parameters are finite.

###### S5.4.1 Estimators are Consistent for Infinite Concentrations

When the true value of the estimand is infinite, we see the correct behavior for a consistent estimator. Section S5.4.1 explains how the survival functions support this conclusion. Section S5.4.1 explains how the distributions of incorrect finite estimates also supports this conclusion.

**Survival Functions Show Anticipated Behavior** When the true value of the concentration estimand is infinite, we see for both the density (as in figures S52, S54, S56, S58 for the plugin estimators and figures S53, S55, S57, S59 for the ML estimators) and compositional concentration estimators (as in figure S60 for the plugin estimator and figure S61 for the ML estimator) that the empirical survival function generally moves to the upper right as the batch size increases. This is the correct behavior for a consistent estimator, since in the case that all estimates were infinite the survival function would be the horizontal line $y = 1$.

**Incorrect Finite Estimates Shift Further to the Right as Batch Size Increases** Moreover, as the size of the batches increases (cf. figures S62, S64, S66, S68, and S70 for the plugin estimators and figures S63, S65, S67, S69, and S71 for the ML estimators) the distribution of the remaining estimates shifts further and further to the right. In other words, even when the estimators incorrectly estimate a finite value, the estimated finite value still tends to increase (and thus "better approximates infinity") as the data size increases.

###### S5.4.2 Estimators are Consistent for Finite Concentrations

When the true value of the concentration estimand is finite, we also see the correct behavior for a consistent estimator. Section S5.4.2 explains how the pattern of infinite estimates supports this conclusion. Section S5.4.2 explains how the estimators satisfy the two minimal requirements for the behavior of a "decent" estimator when the model is correctly specified. Section S5.4.2 explains why the estimates are also good even in the case when the model is not correctly specified.

[35] Via SciPy (Virtanen et al., 2020), which chooses bandwidths according to Scott's Rule (Scott, 1992).

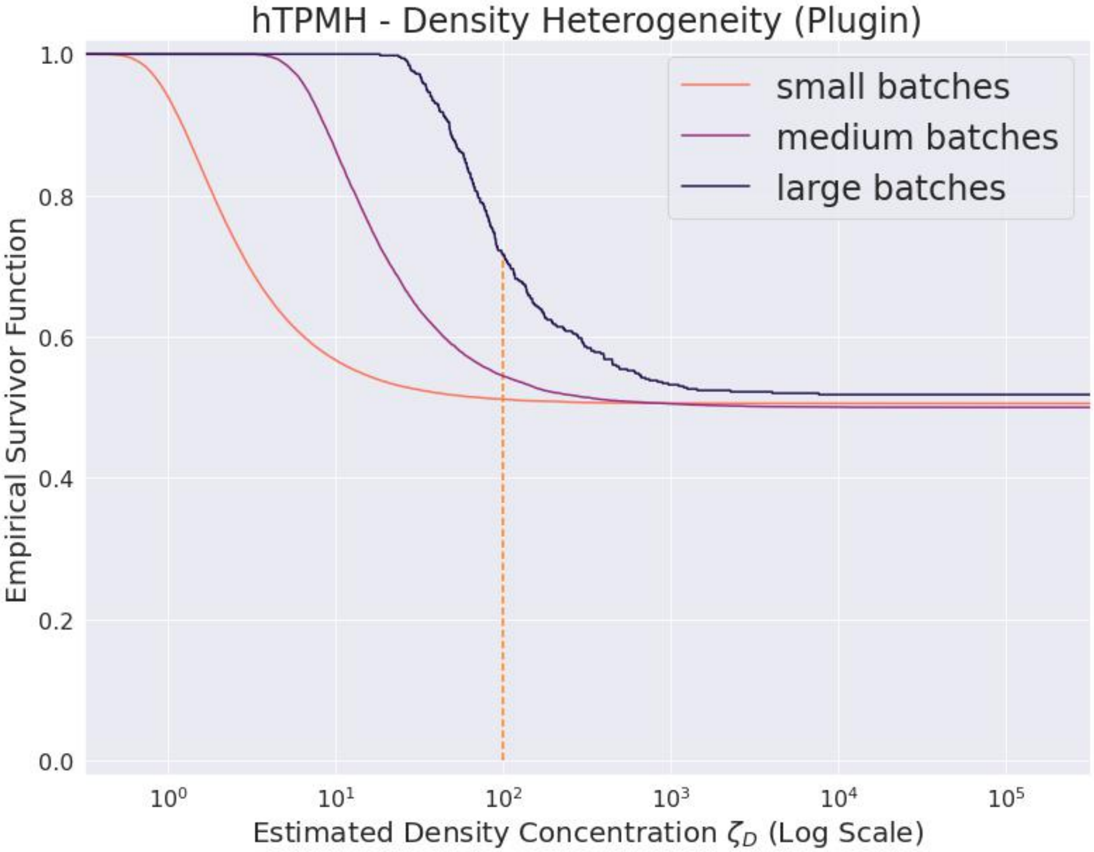

**Supplementary Figure S52**

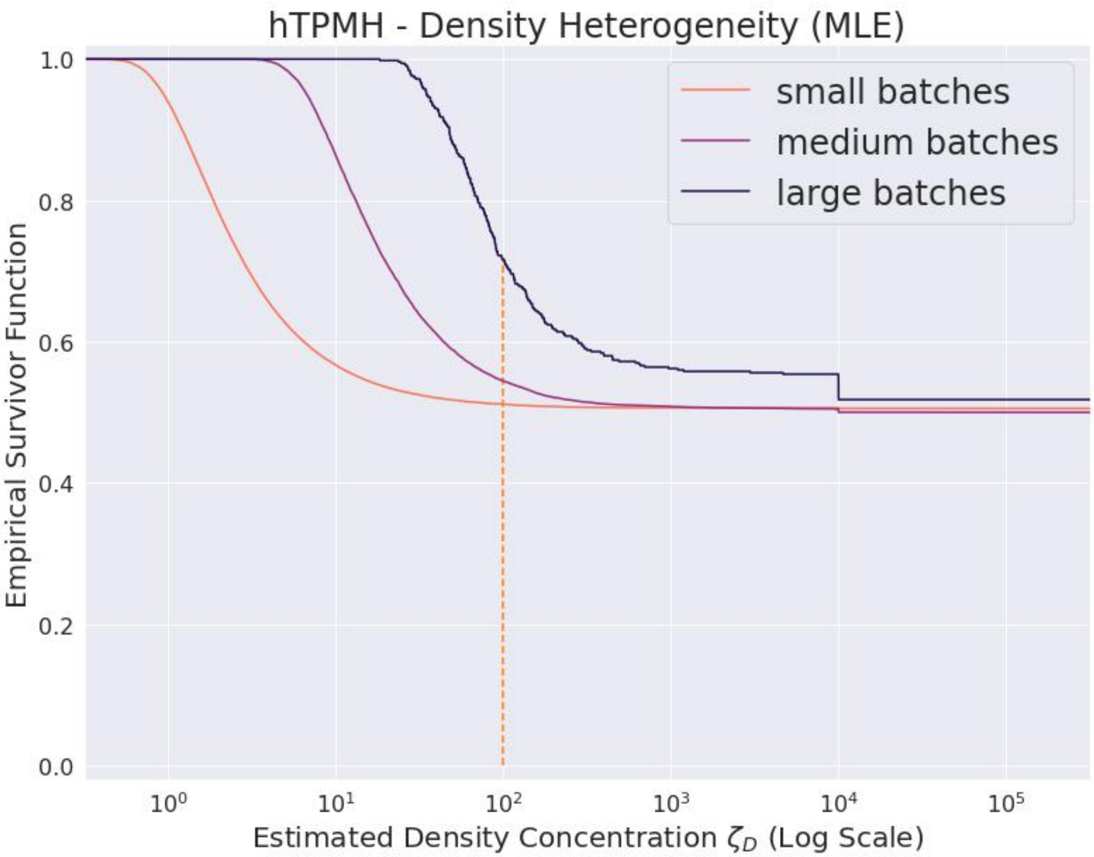

**Supplementary Figure S53**

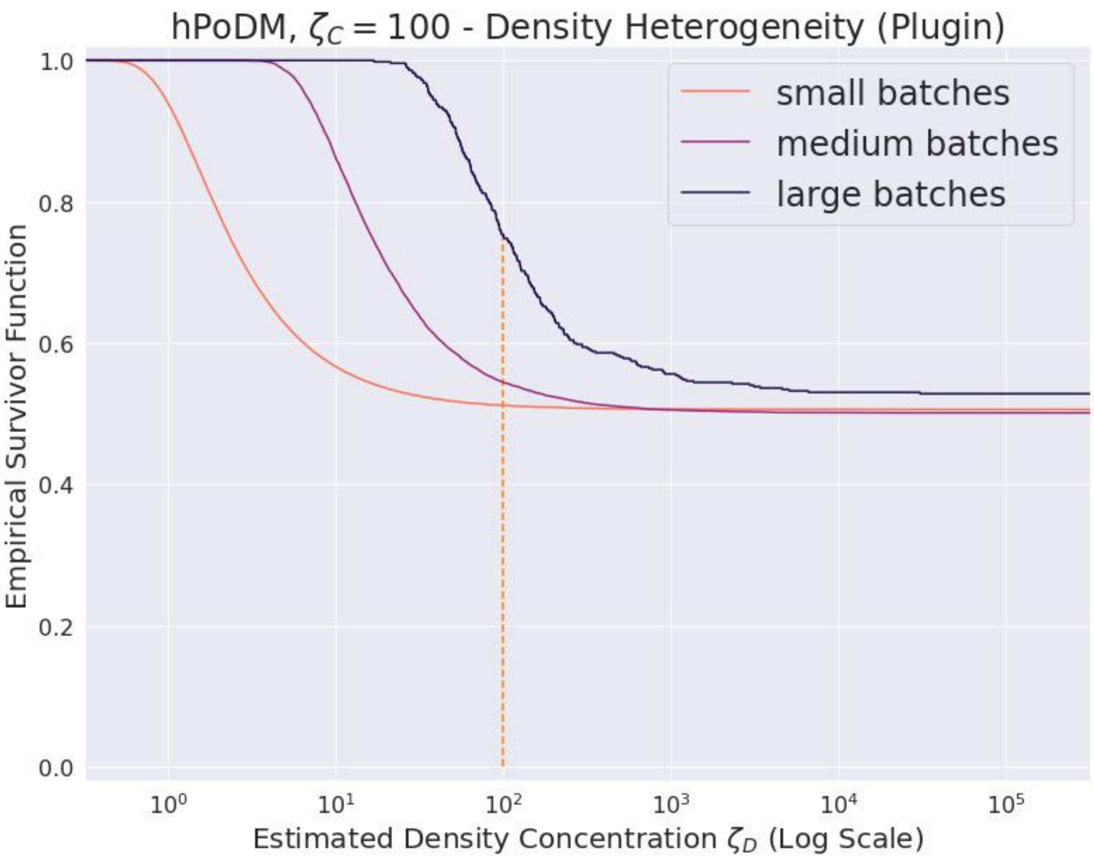

**Supplementary Figure S54**

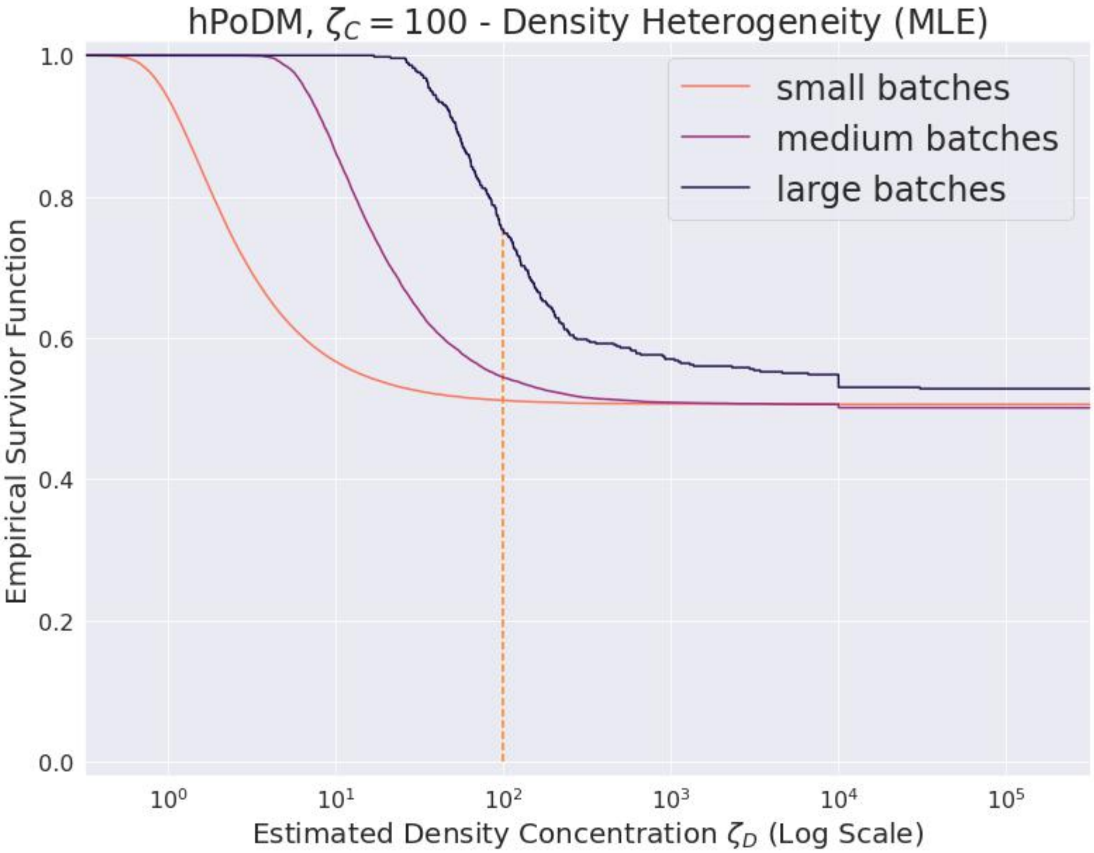

**Supplementary Figure S55**

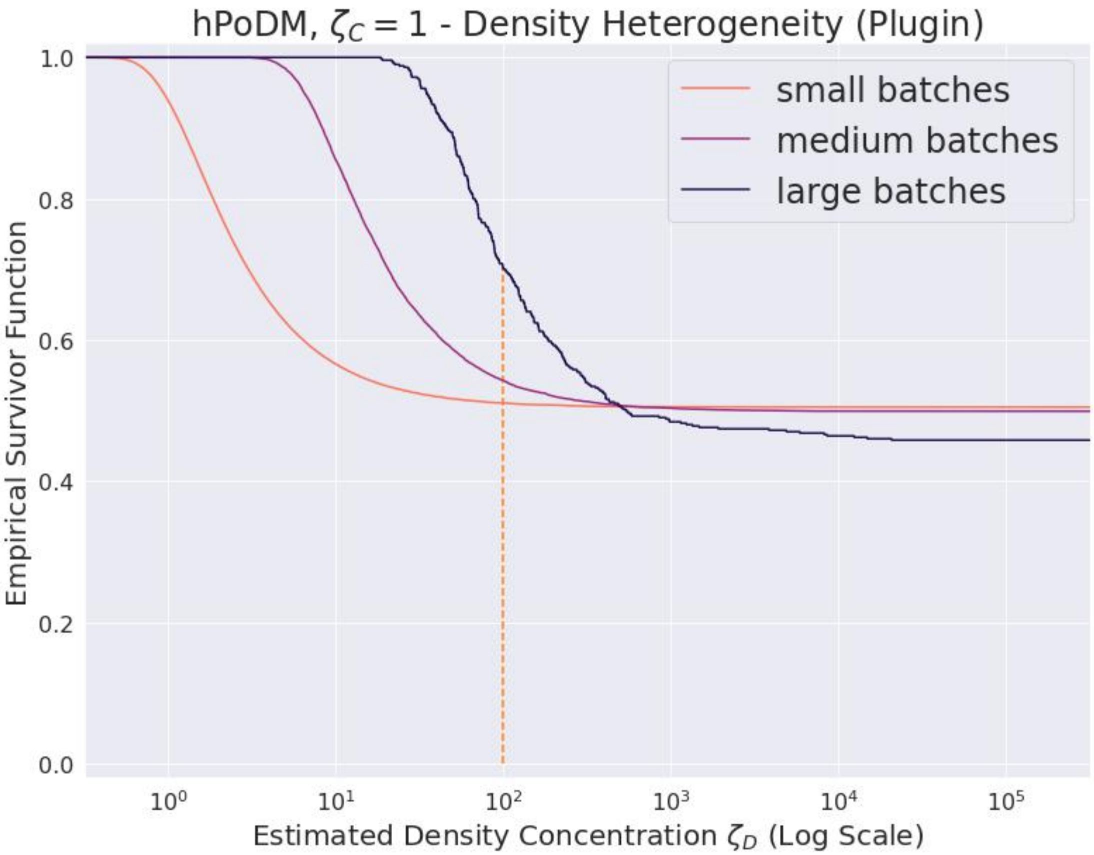

**Supplementary Figure S56**

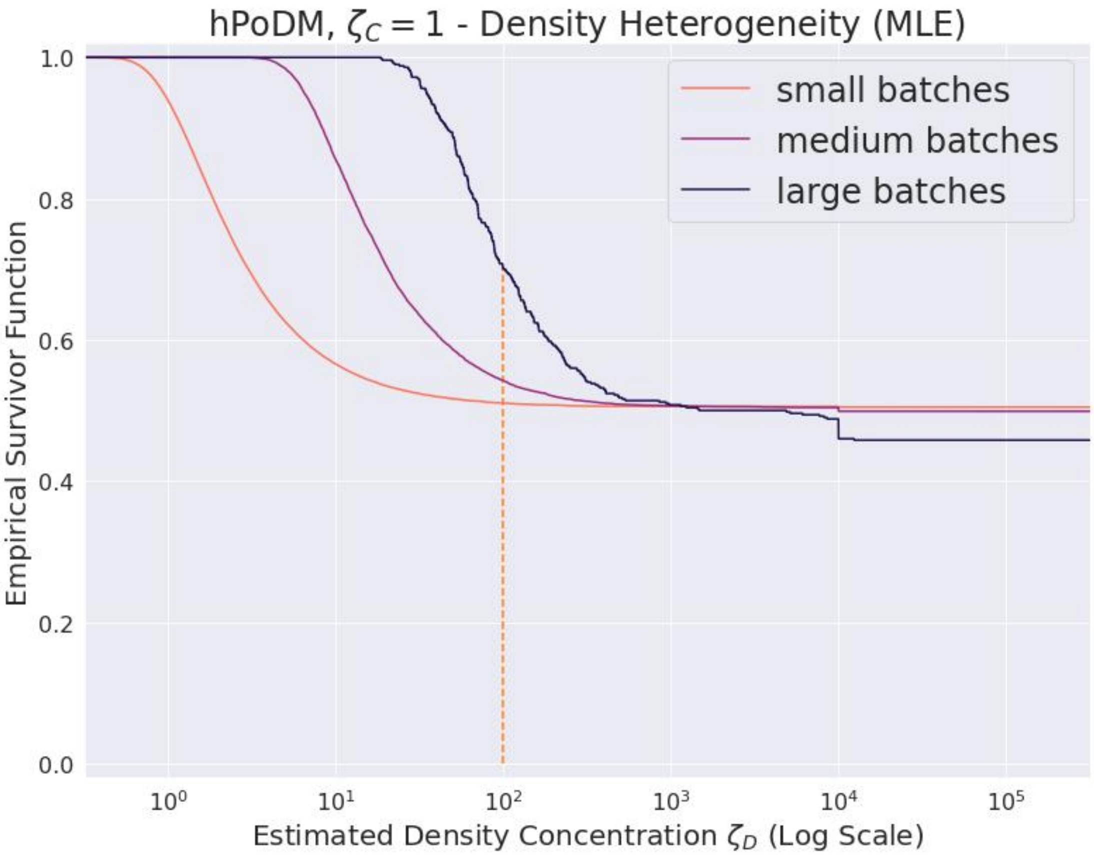

**Supplementary Figure S57**

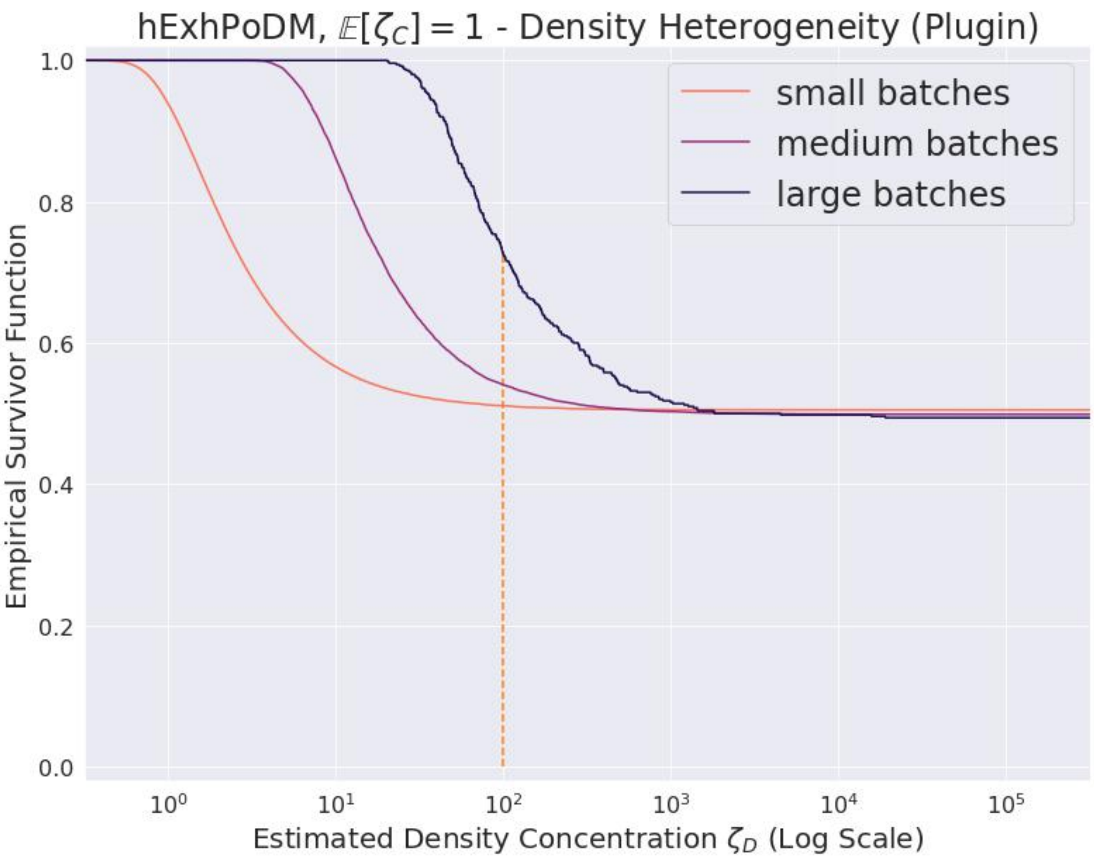

**Supplementary Figure S58**

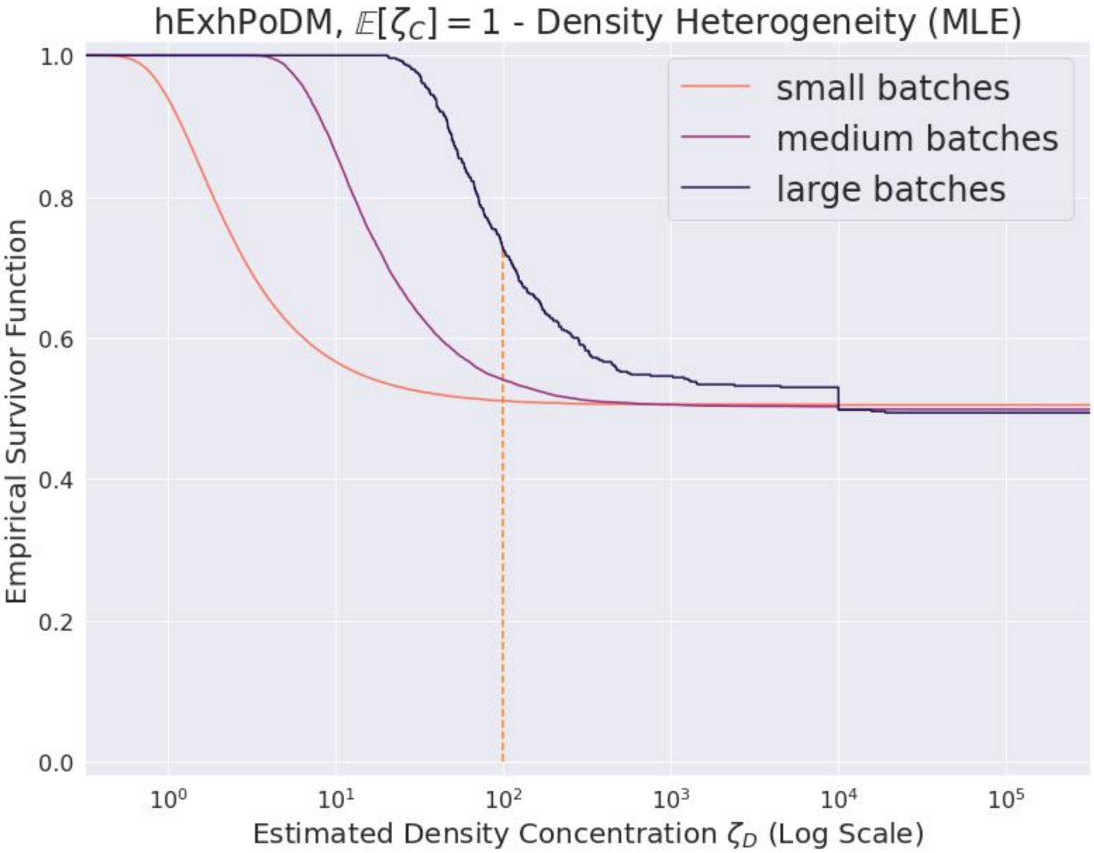

**Supplementary Figure S59**

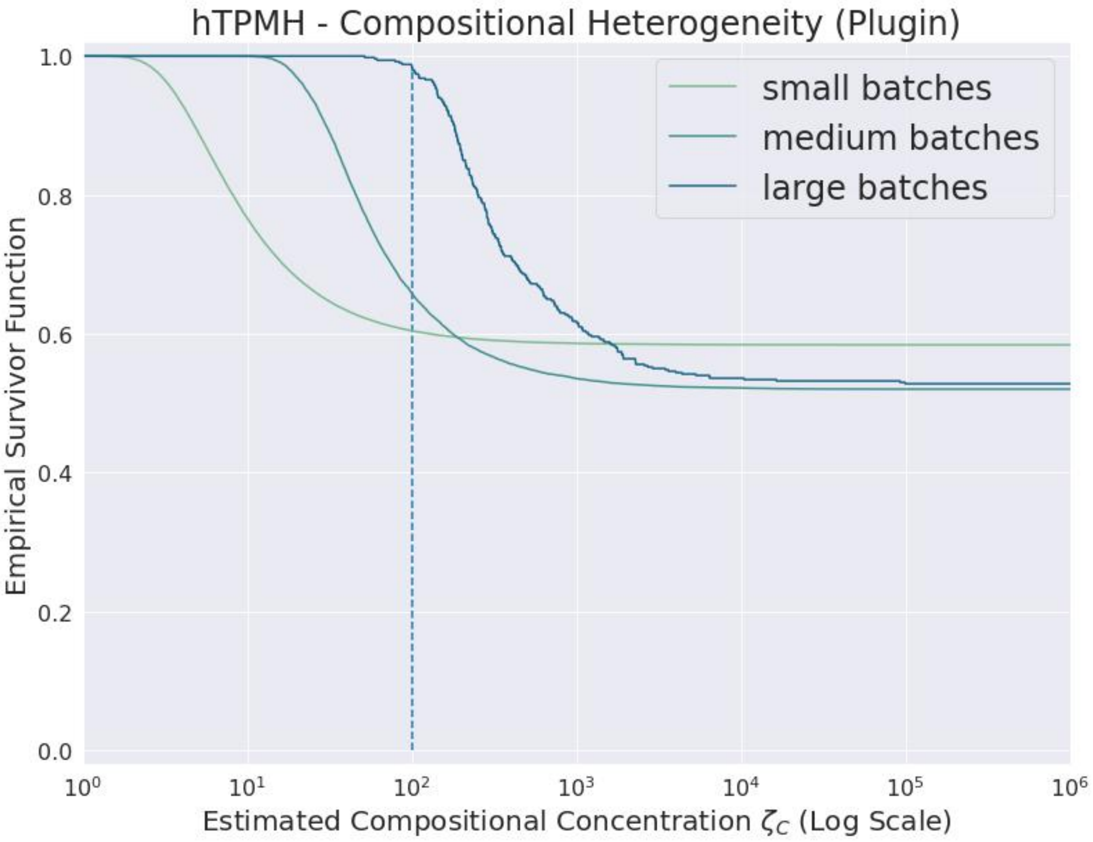

**Supplementary Figure S60**

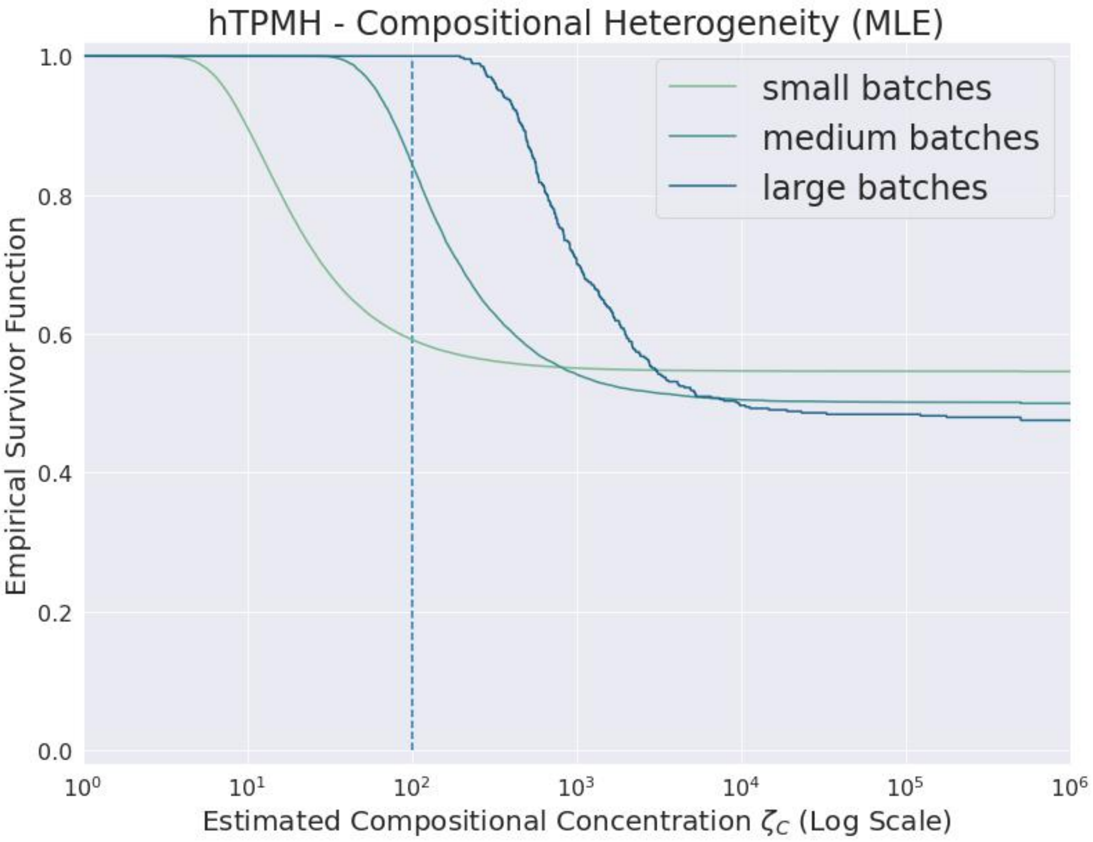

**Supplementary Figure S61**

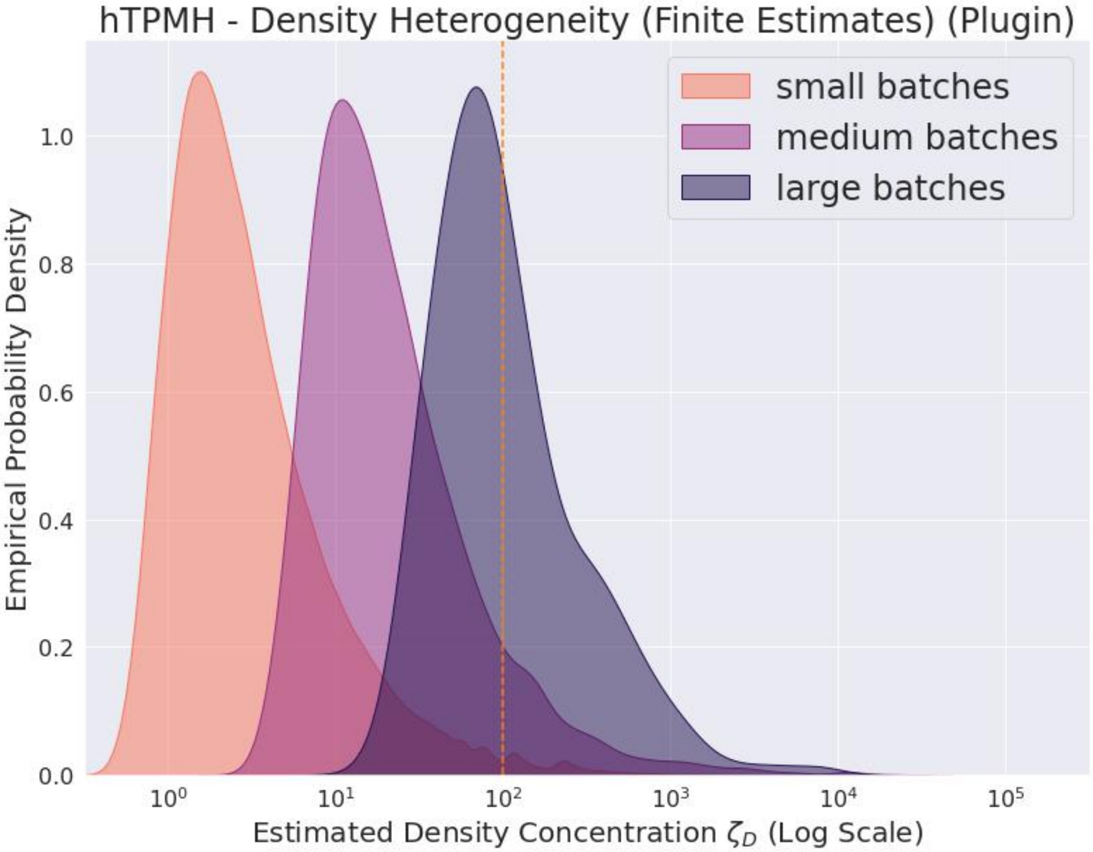

**Supplementary Figure S62**

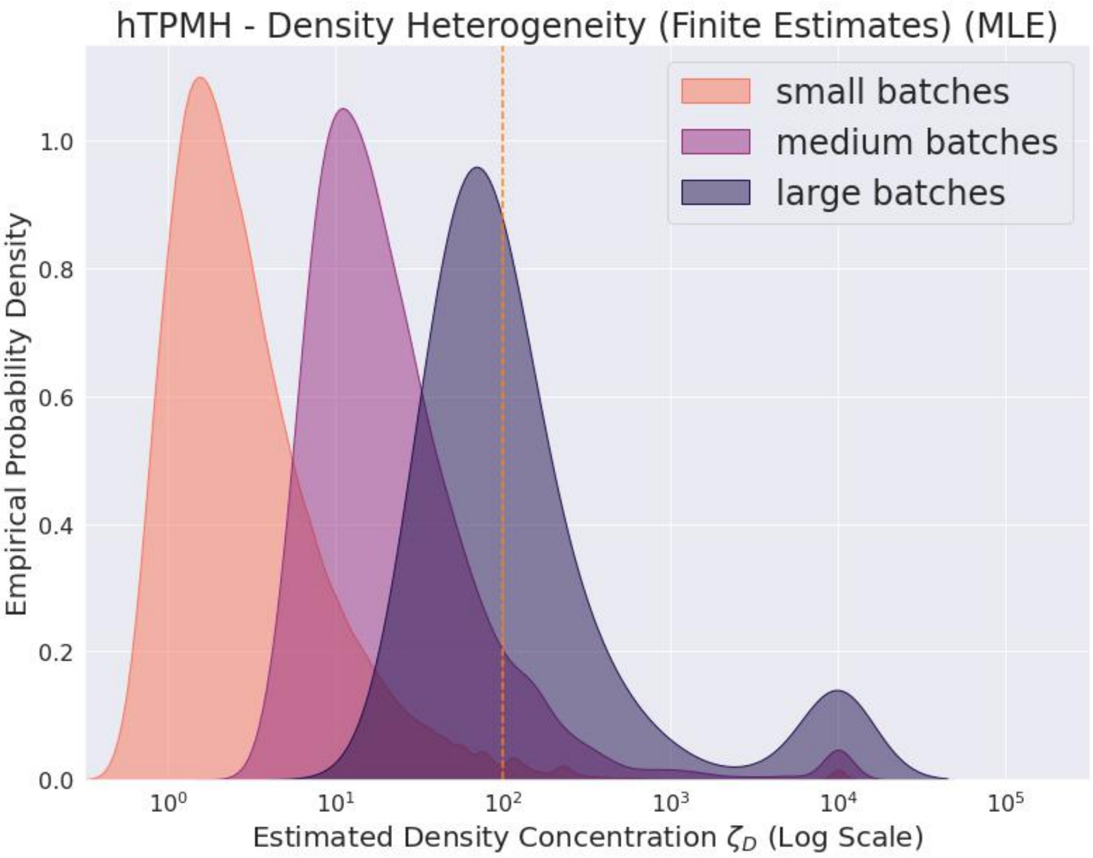

**Supplementary Figure S63**

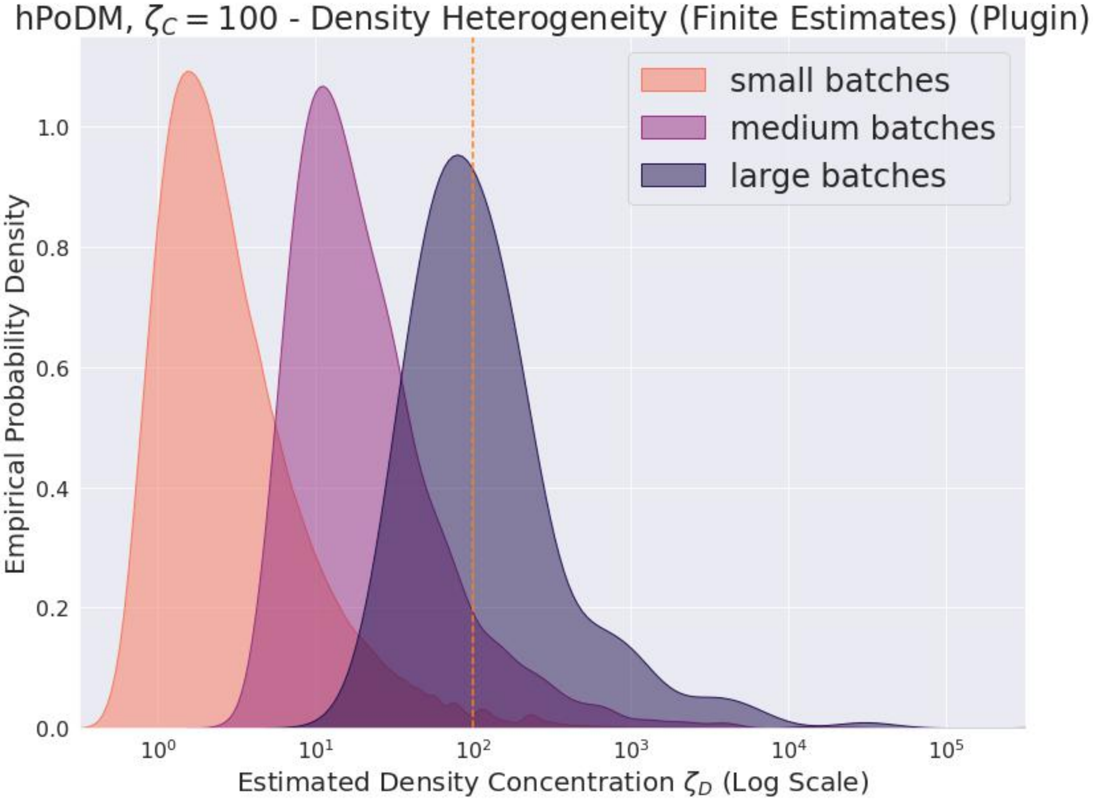

**Supplementary Figure S64**

**Incorrect Infinite Estimates Only Occur for Distributions Very Similar to hPoMu** For the distributions which are very similar to the homogeneous case with infinite concentration, the concentration estimates are sometimes infinite (cf. figures S72, S74, and S76 for the plugin estimators and figures S73, S75, and S77 for the ML estimators). Nevertheless, these figures also show that, as the batch size increases, the proportion of concentration estimates which are infinite decreases. This appears to occur more quickly for the compositional concentration estimators (see figures S72 and S76 for the plugin estimators and figures S73 and S77 for the ML estimators) than for the density concentration estimators (see figure S74 for the plugin estimator and figure S75 for the ML estimator).

For the distributions which are less similar to hPoMu, when the true value of the concentration estimand is finite, the estimated values are either never or almost never infinite. This is why survival functions are not reported neither for the density concentration estimates of hNBDM with $\zeta = 1$ and hExhNBDM with $\mathbb{E}[\zeta] = 1$ nor for the compositional concentration estimates of hPoDM with $\zeta_C = 1$, hNBDM with $\zeta = 1$, hExhPoDM with $\mathbb{E}[\zeta_C] = 1$, and hExhNBDM with $\mathbb{E}[\zeta] = 1$.

**Accuracy and Precision Both Increase with Batch Size** When the true value of the concentration estimand is finite, and the model is correctly specified, we see (cf. figures S78, S80, S82, S84, S86, and S88 for the plugin estimators and figures S79, S81, S83, S85, S87, and S89 for the ML estimators) that the distribution of the (finite) estimates is centered around the true value.

When the limit approached by the estimators is finite, even when the model is *not* correctly specified, the variance of the distribution of estimated concentrations decreases as the batch size increases. Cf. figures S78, S80, S82, S84, S86, S88, S90, S92, and S94 for the plugin estimators and figures S79, S81, S83, S85, S87, S89, S91, S93, and S95 for the ML estimators.

**Results are Qualitatively Correct under Model Misspecification** For hExhPoDM with $\mathbb{E}[\zeta_C] = 1$, and hExhNBDM with $\mathbb{E}[\zeta] = 1$, the model is misspecified[36], since these distributions do not have single

[36] Note that technically for hExhPoDM, only the categorical distribution, corresponding to the compositional concentration, is misspecified relative to the chosen estimators. For hExhNBDM both the count distribution, corresponding to the density

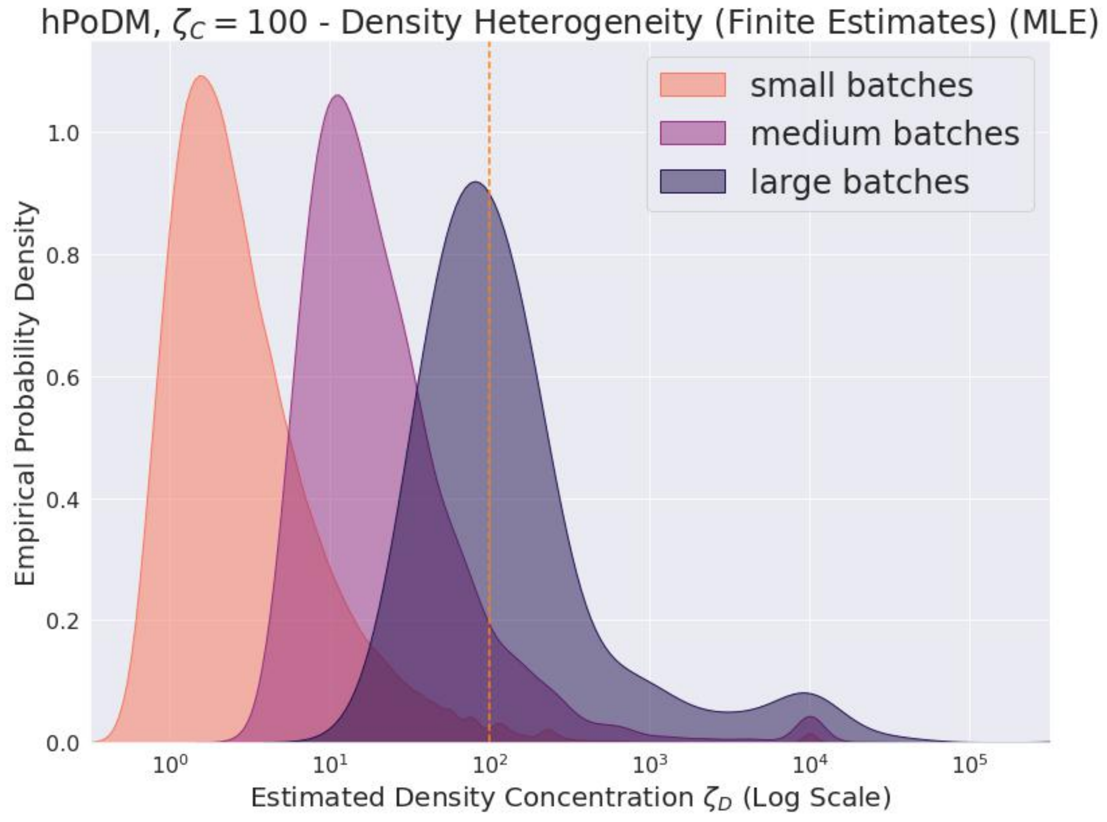

**Supplementary Figure S65**

values of concentration parameters. Nevertheless, from the results in previous sections it is also clear that these are the most heterogeneous distributions. These estimators therefore give qualitatively correct answers for these distributions, despite the model being misspecified, because (cf. figures S90, S92, and S94 for the plugin estimators and figures S91, S93, and S95 for the ML estimators) the lowest concentration estimates occur for these distributions.

##### *S5.4.3 Comparing ML Estimator and Plugin Estimator Results*

An important reason for using ML estimators over plugin estimators is increased efficiency. Cf. the discussion from the beginning of section S5.2. Therefore I evaluate whether the precision of the ML estimator appears to increase much more rapidly than that of its plugin counterpart for the density estimators in section S5.4.3 and for the concentration estimators in section S5.4.3. One reason for using the plugin estimators over ML estimators is because the plugin estimators arguably make fewer model assumptions. Again, cf. the discussion at the beginning of section S5.2. Therefore in section S5.4.3 I compare the behaviors of the plugin and ML estimators for the misspecified models.

**Precision of ML Estimator vs. Plugin Estimator for Density Concentration** For density concentration, both ML and plugin appear to demonstrate effectively the same efficiency in practice. See figures S52, S53, S54, S55,S56, S57, S58, and S59 for survival curves when the true value of the estimand is infinite, figures S74 and S75 for survival curves when the true value of the estimand is finite, figures S62, S63, S64, S65, S66, S67, S68, and S69 for KDE plots when the true value of the estimand is infinite, and figures S80, S81, S86, and S87 for KDE plots when the true value of the estimand is finite.

I double-checked the exact values and the estimates produced by the two estimators really are different, even though these differences are not apparent[37] from the aforementioned graphs. This makes sense given that the median difference in the estimated values was never greater than $3 \times 10^{-4}$ (data not shown) and was often smaller (especially for the larger batches). Evidently the difference in the asymptotic efficiency

---

concentration, and the categorical distribution are misspecified relative to the chosen estimators.

[37] Except for the bumps around $10^4$, see below.

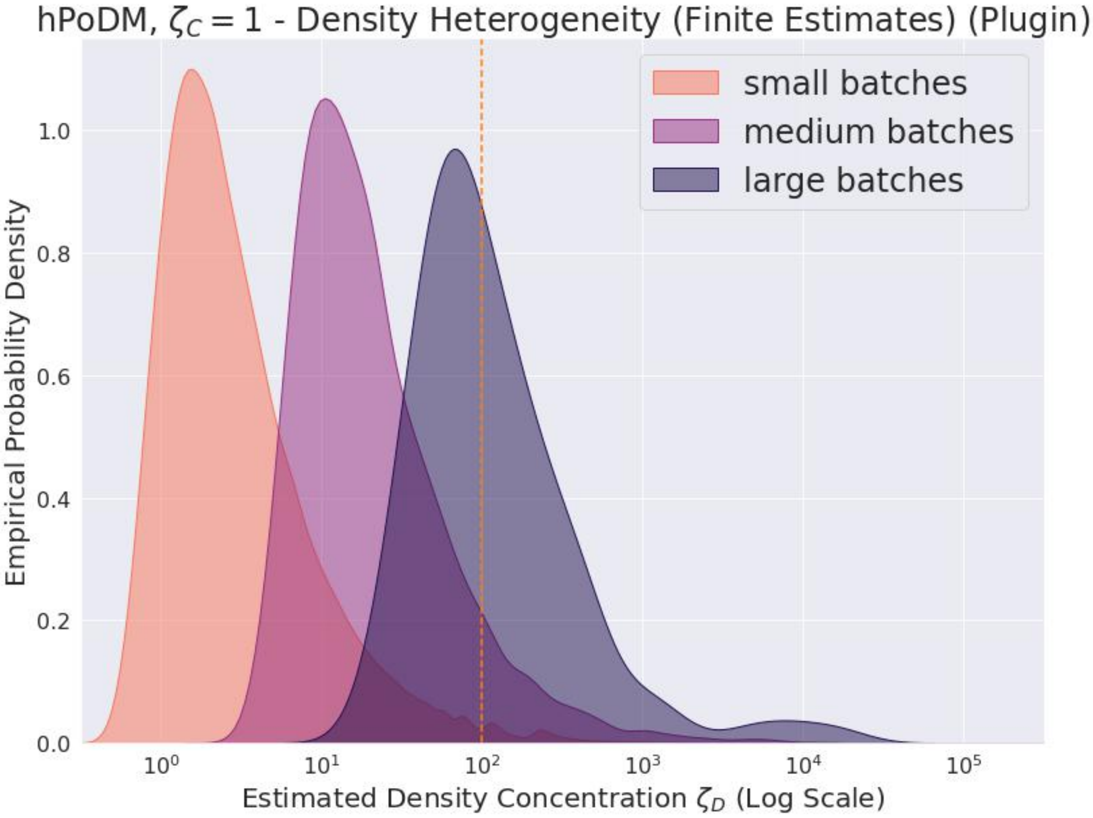

**Supplementary Figure S66**

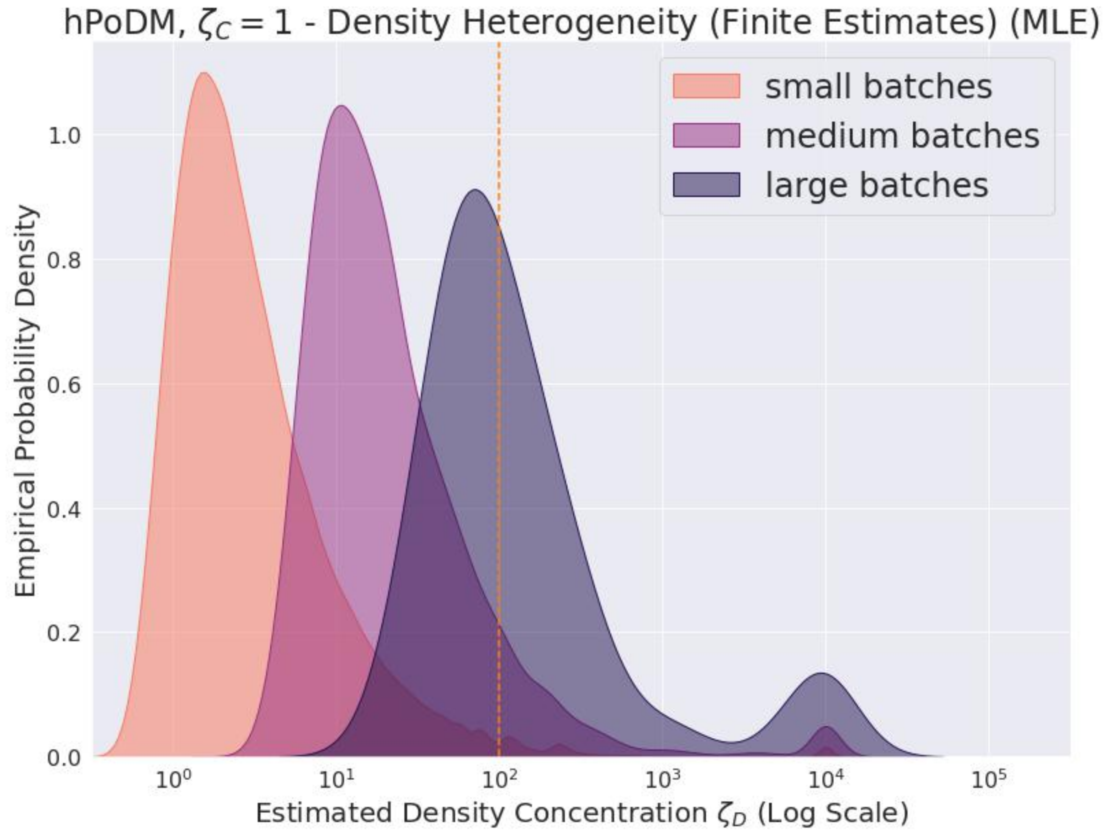

**Supplementary Figure S67**

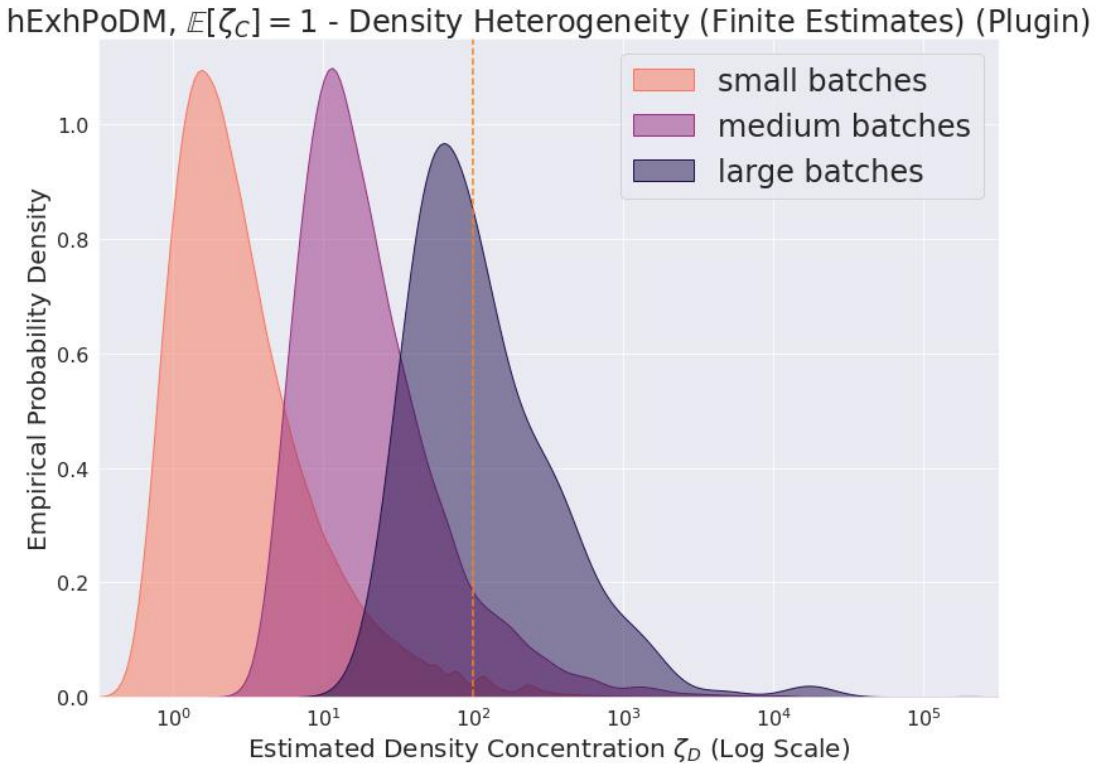

**Supplementary Figure S68**

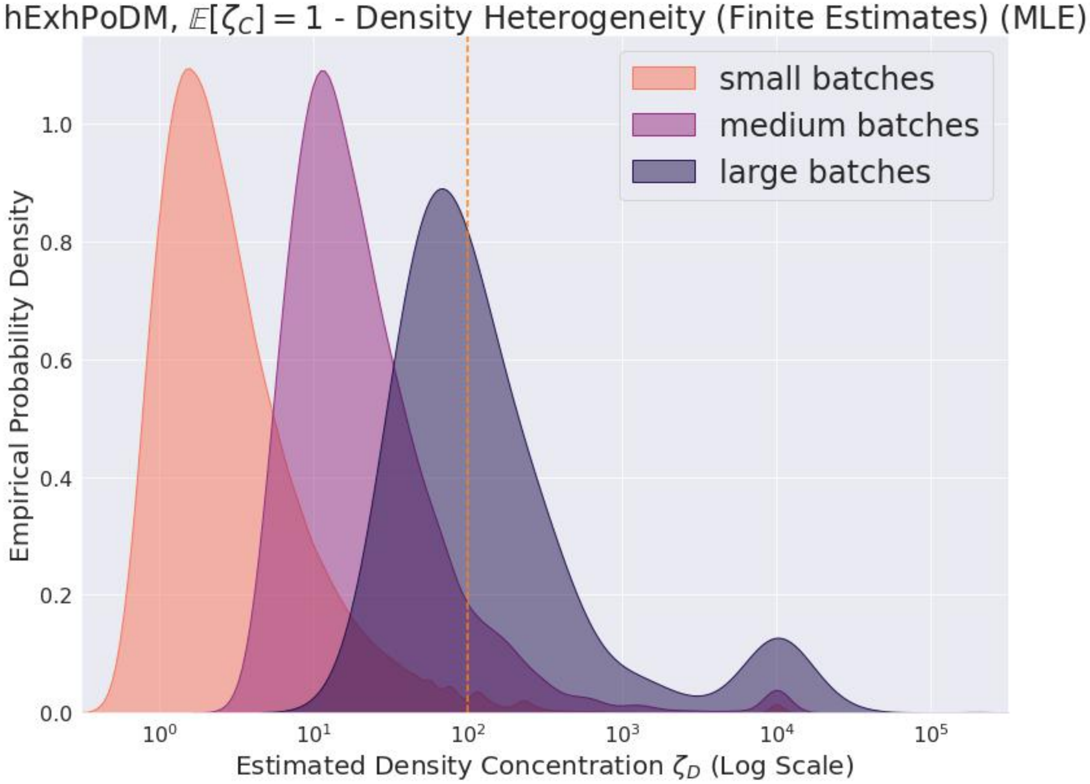

**Supplementary Figure S69**

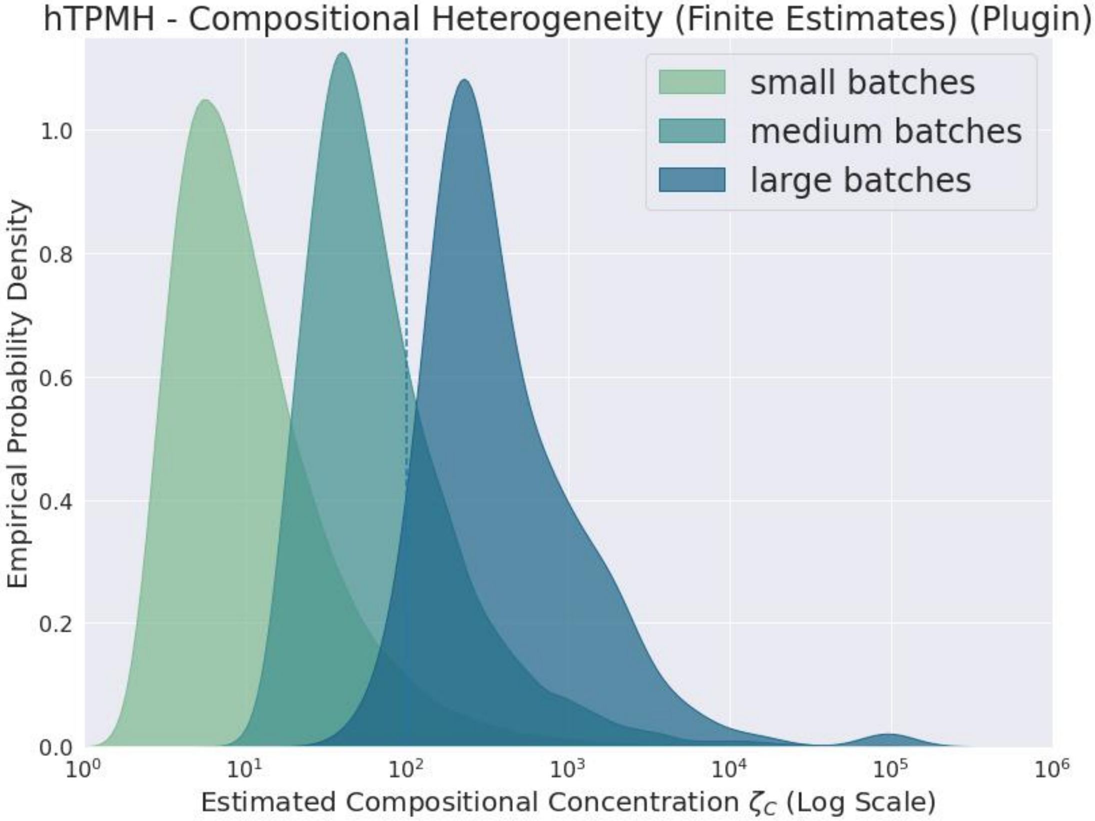

**Supplementary Figure S70**

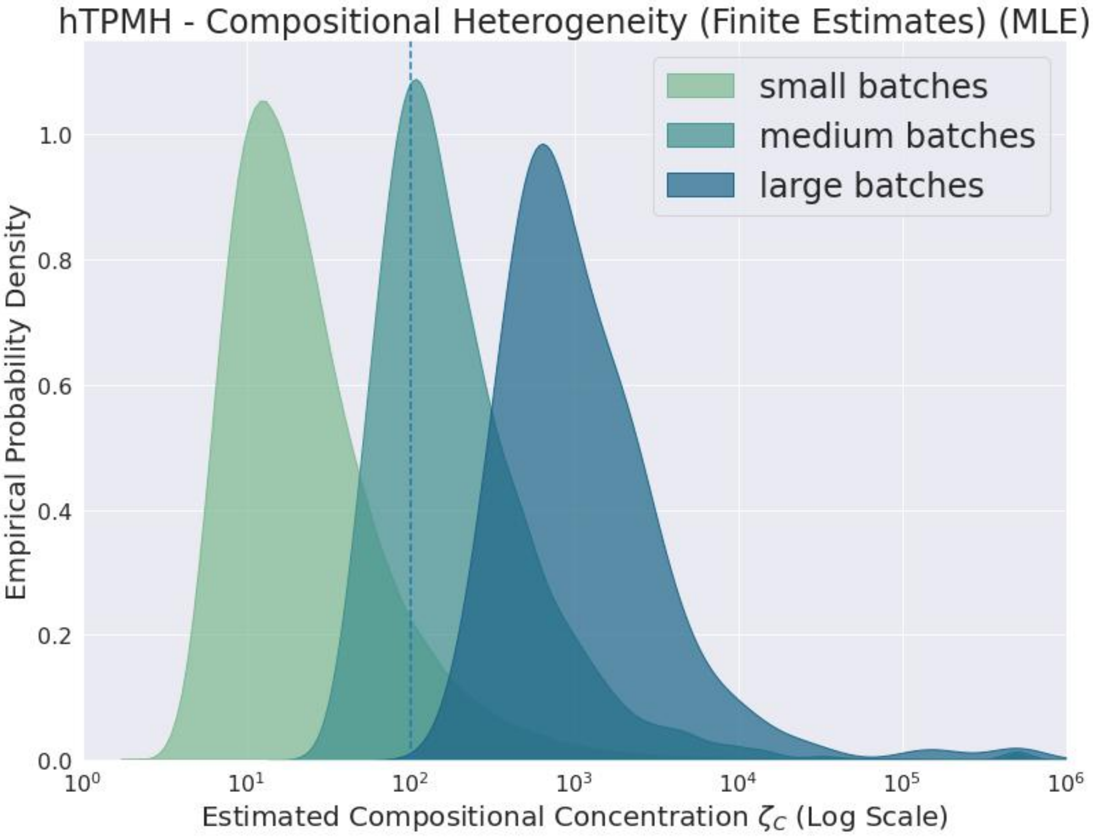

**Supplementary Figure S71**

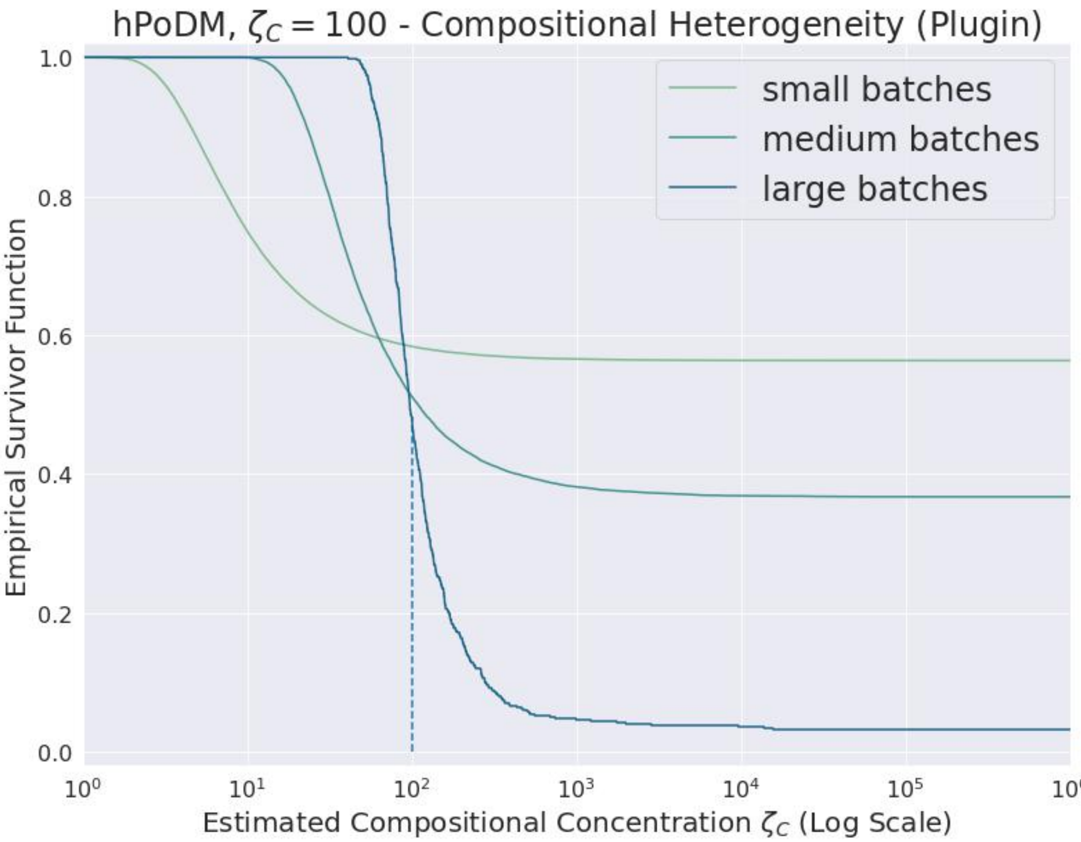

**Supplementary Figure S72**

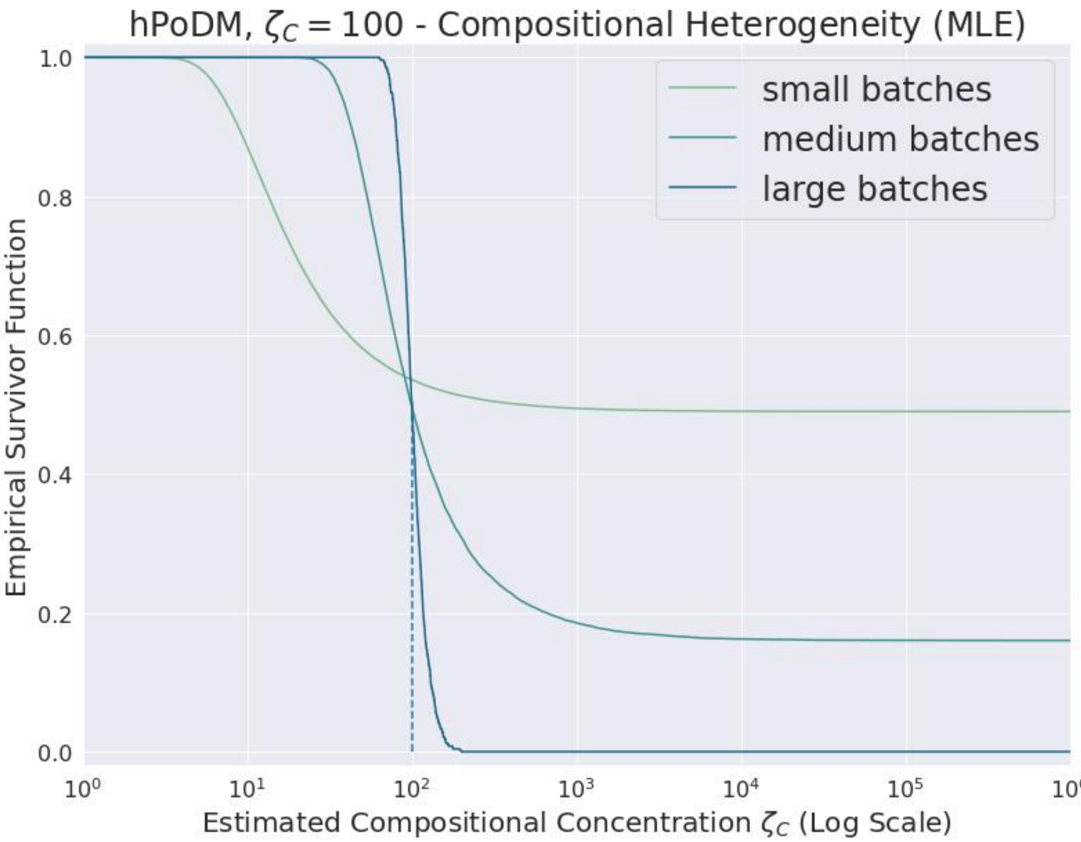

**Supplementary Figure S73**

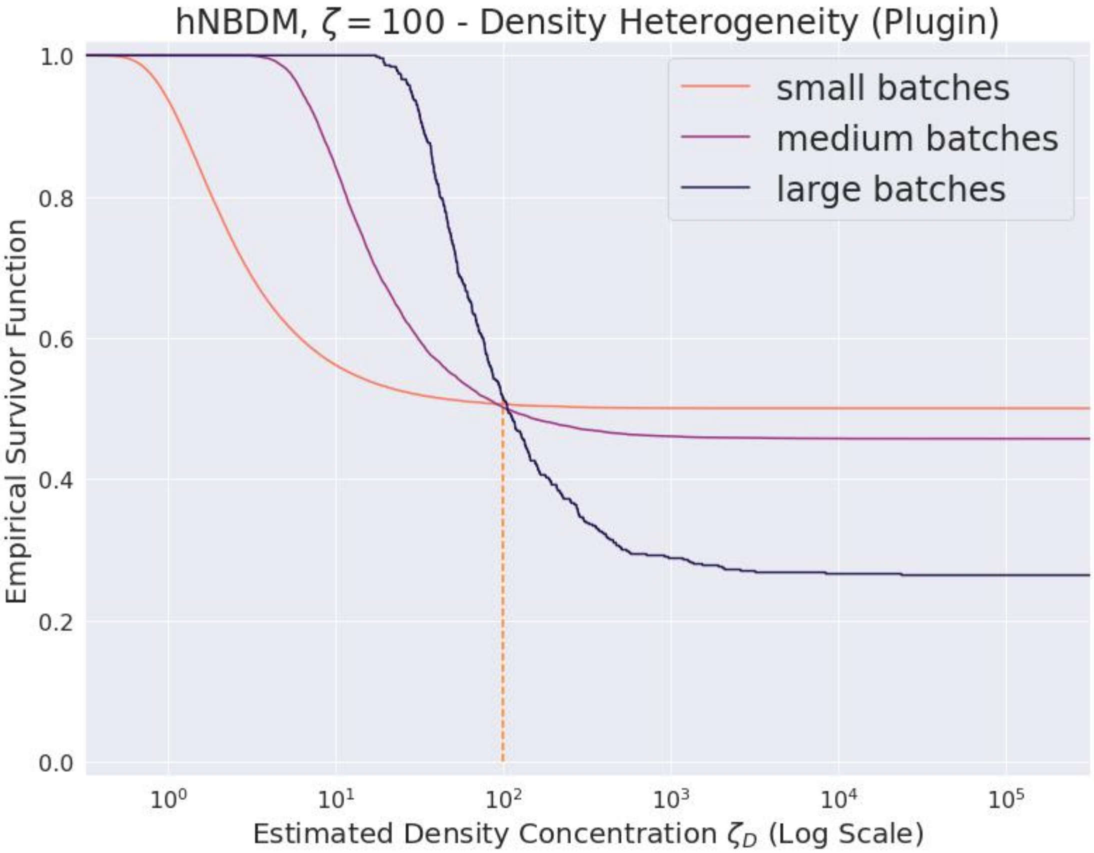

**Supplementary Figure S74**

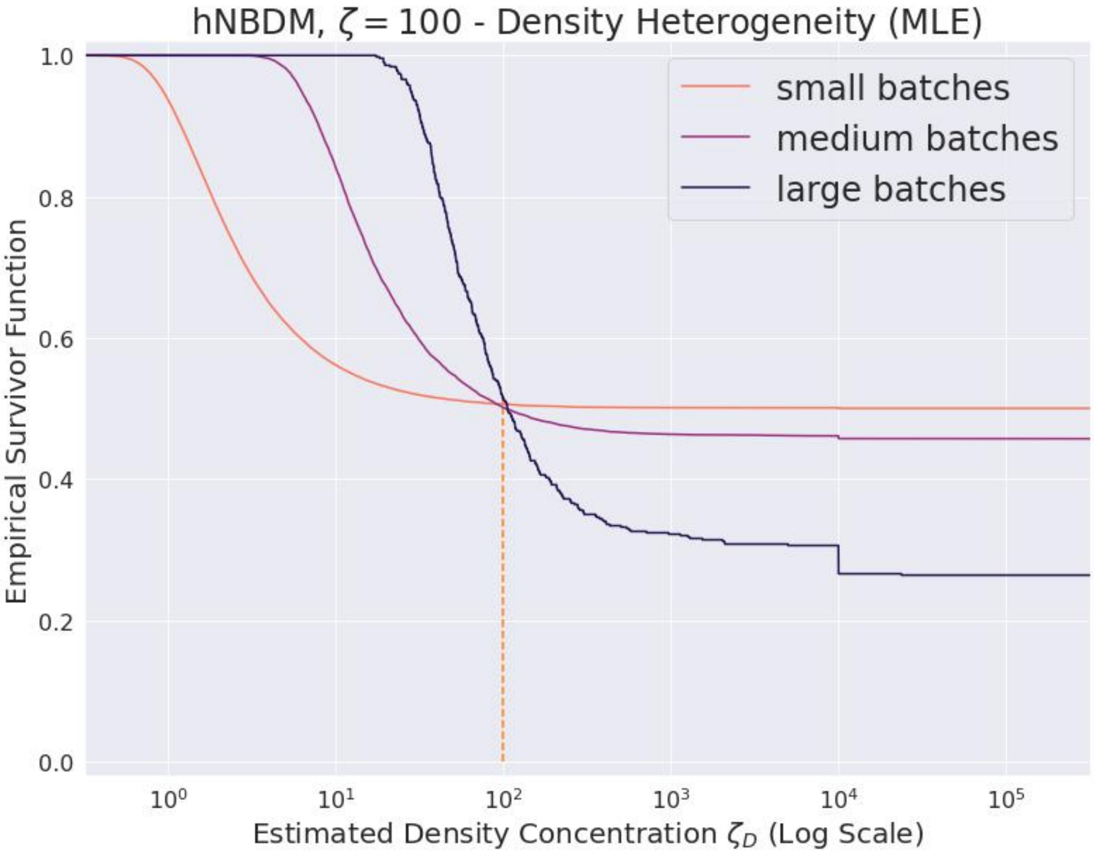

**Supplementary Figure S75**

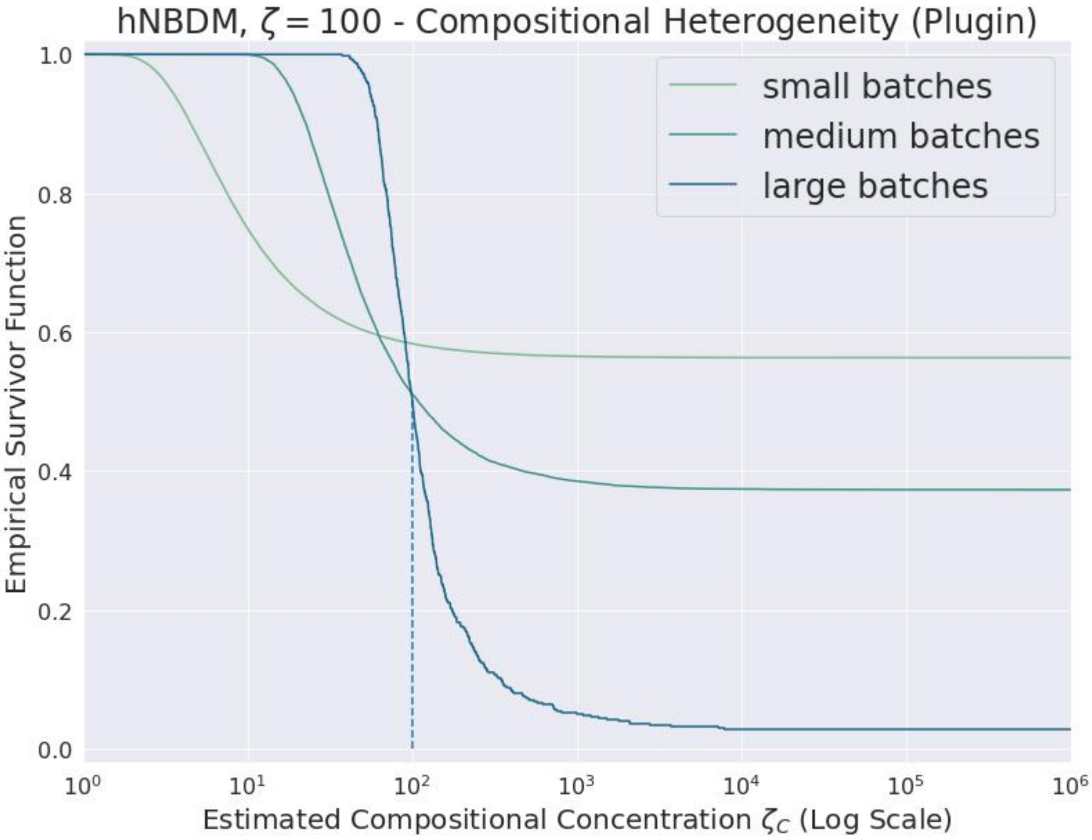

**Supplementary Figure S76**

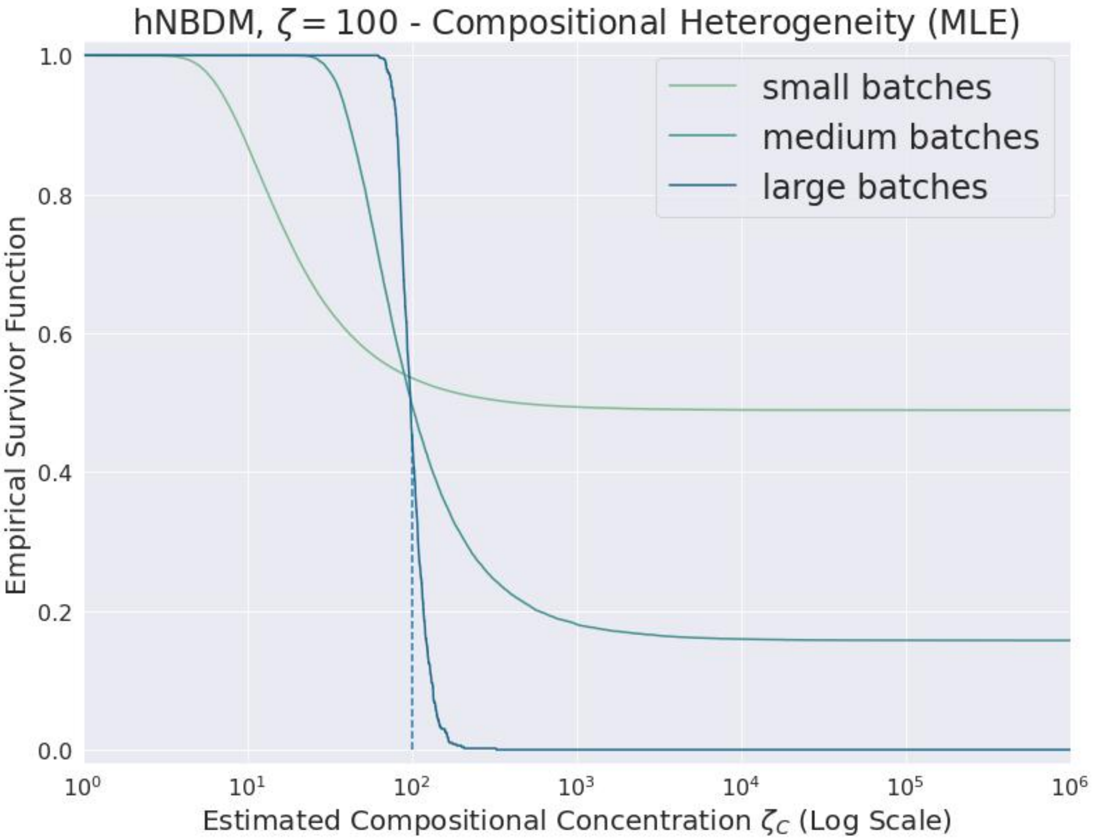

**Supplementary Figure S77**

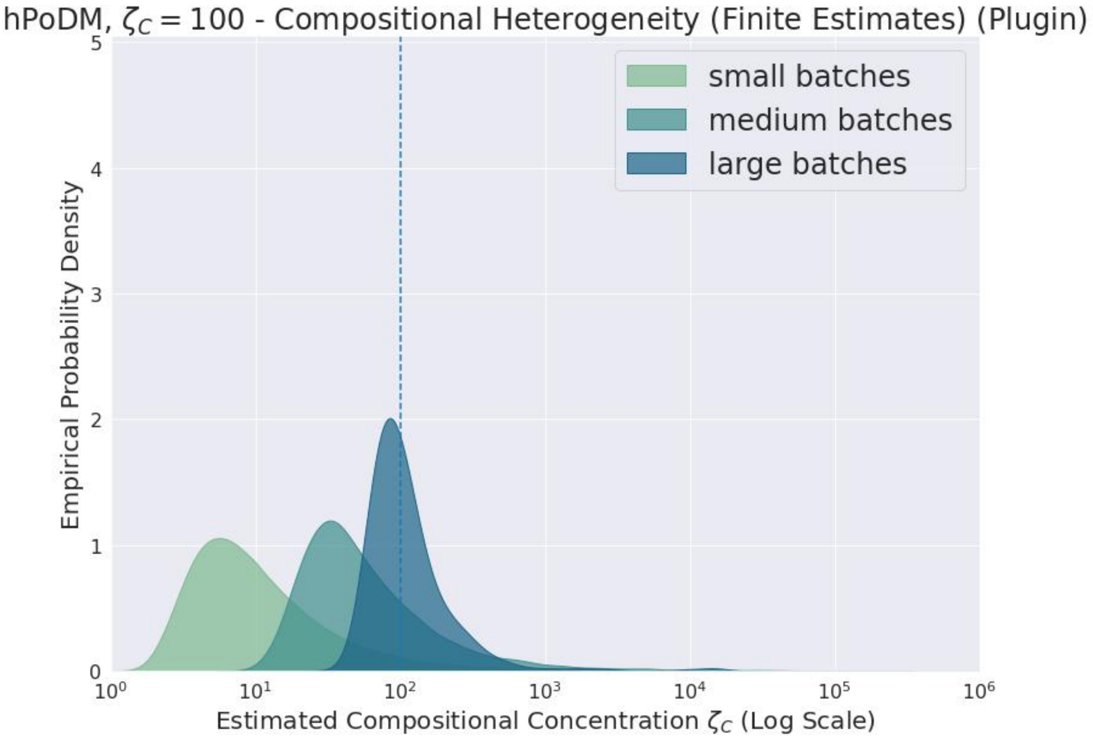

**Supplementary Figure S78**

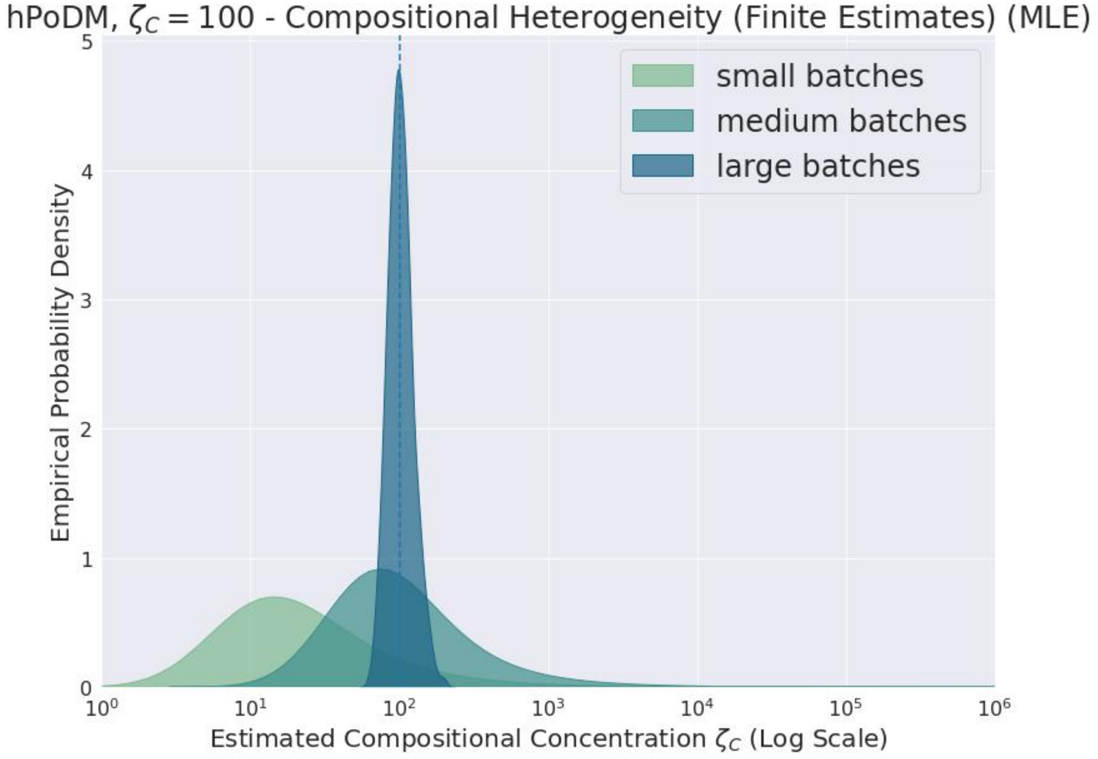

**Supplementary Figure S79**

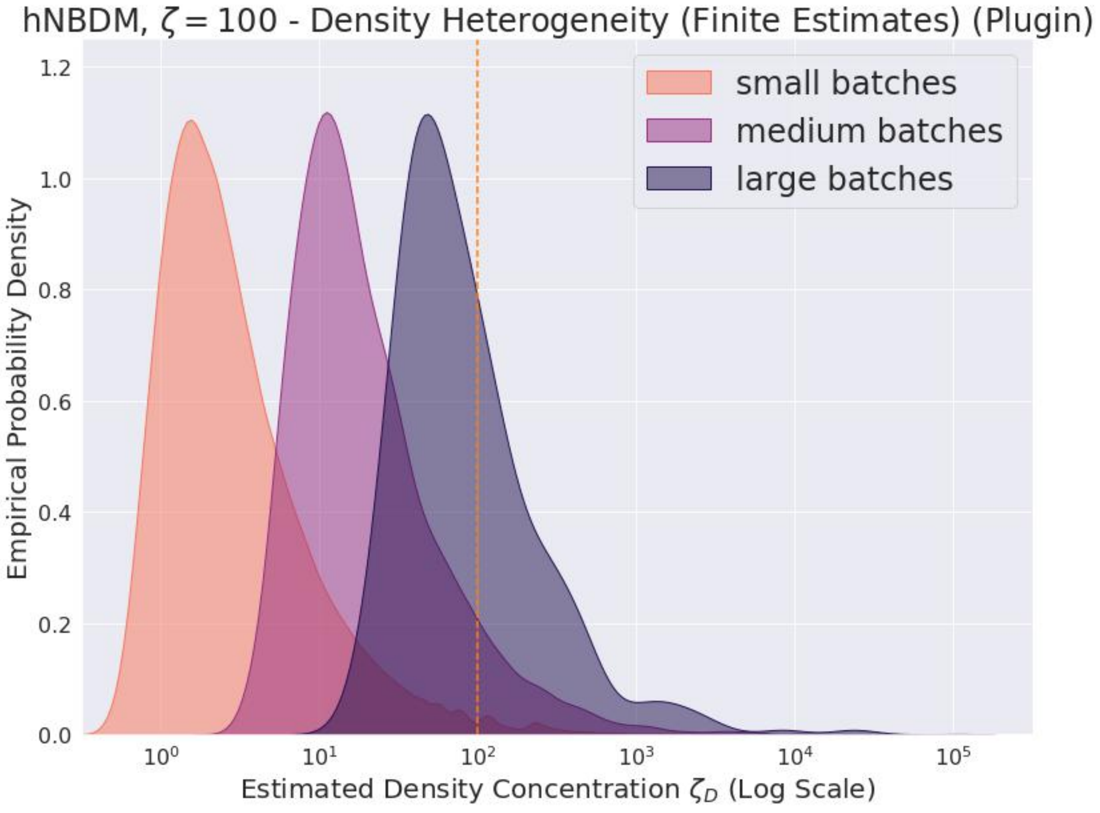

**Supplementary Figure S80**

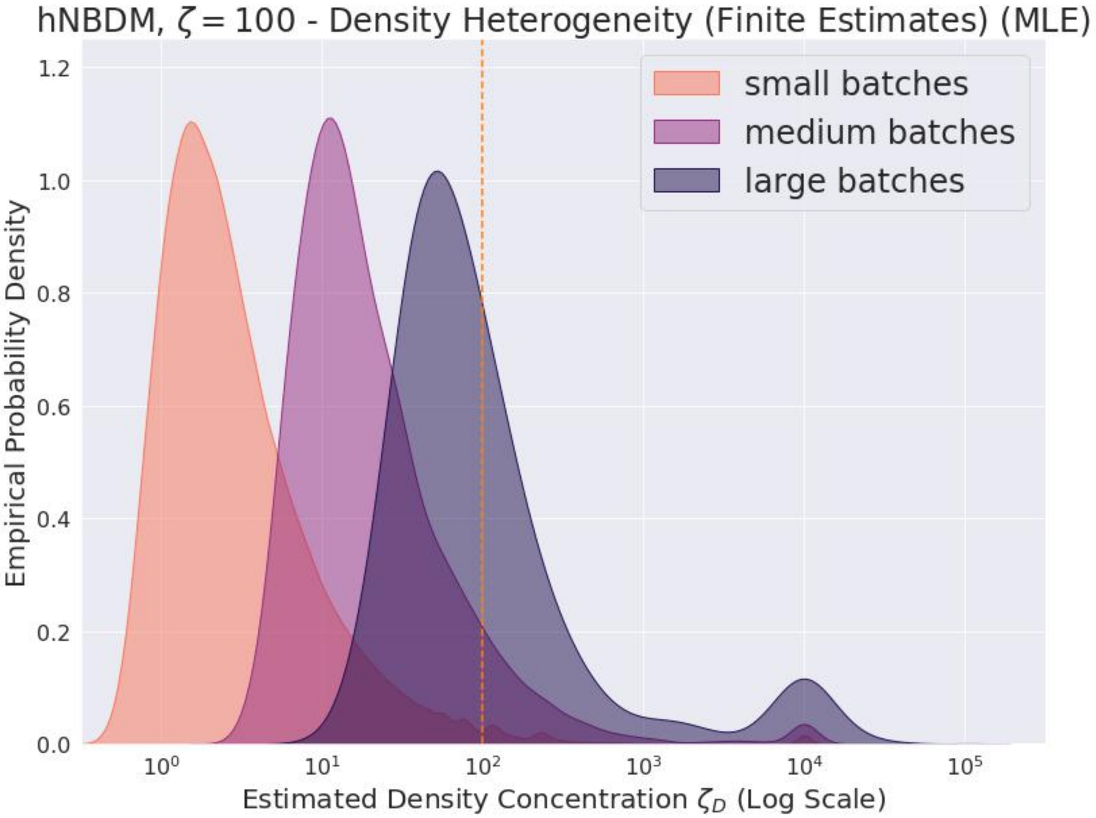

**Supplementary Figure S81**

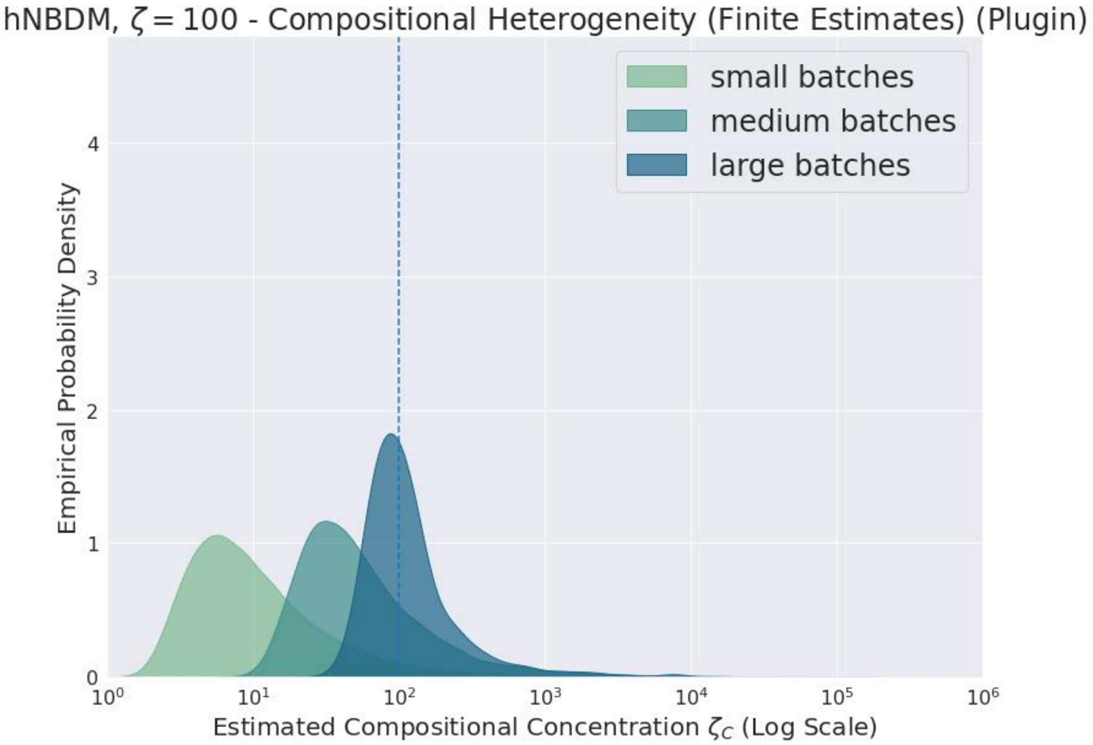

**Supplementary Figure S82**

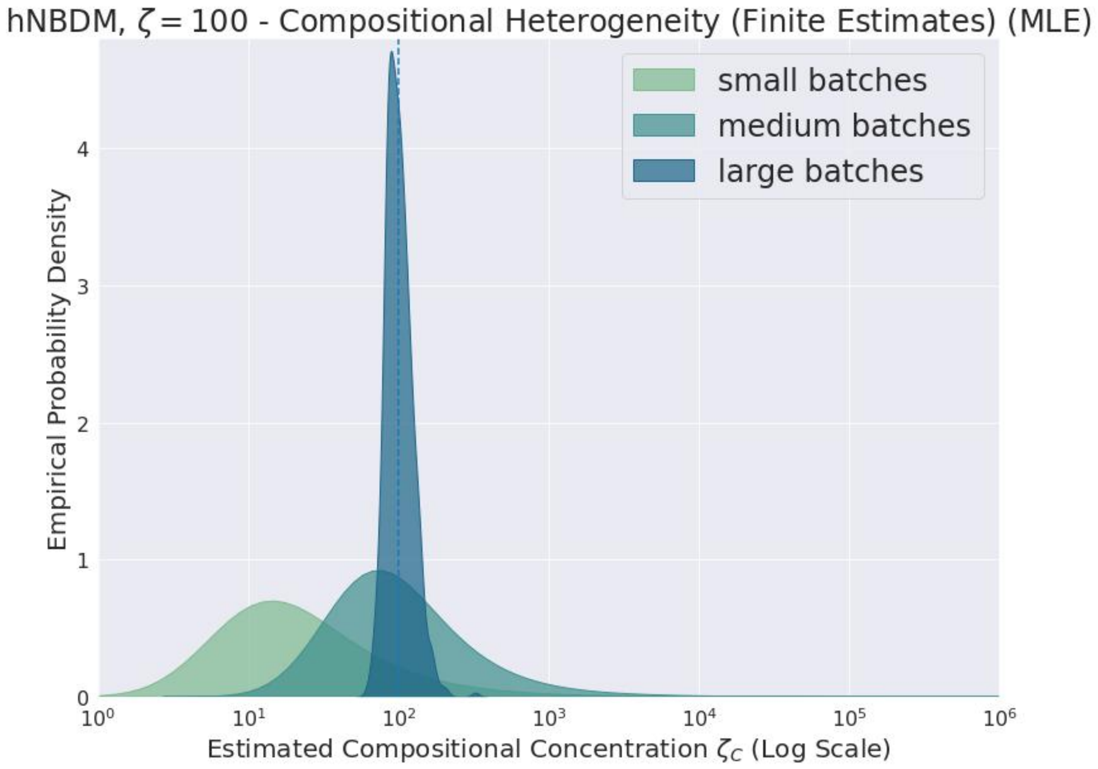

**Supplementary Figure S83**

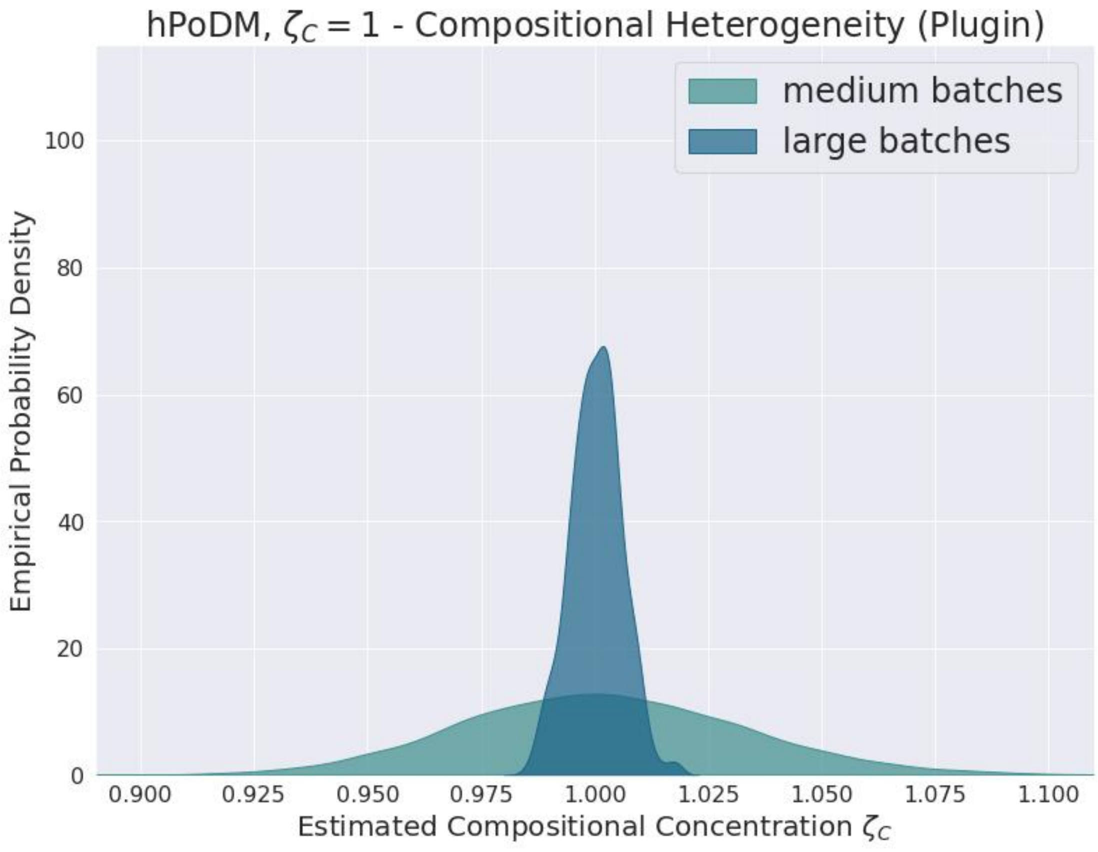

**Supplementary Figure S84**

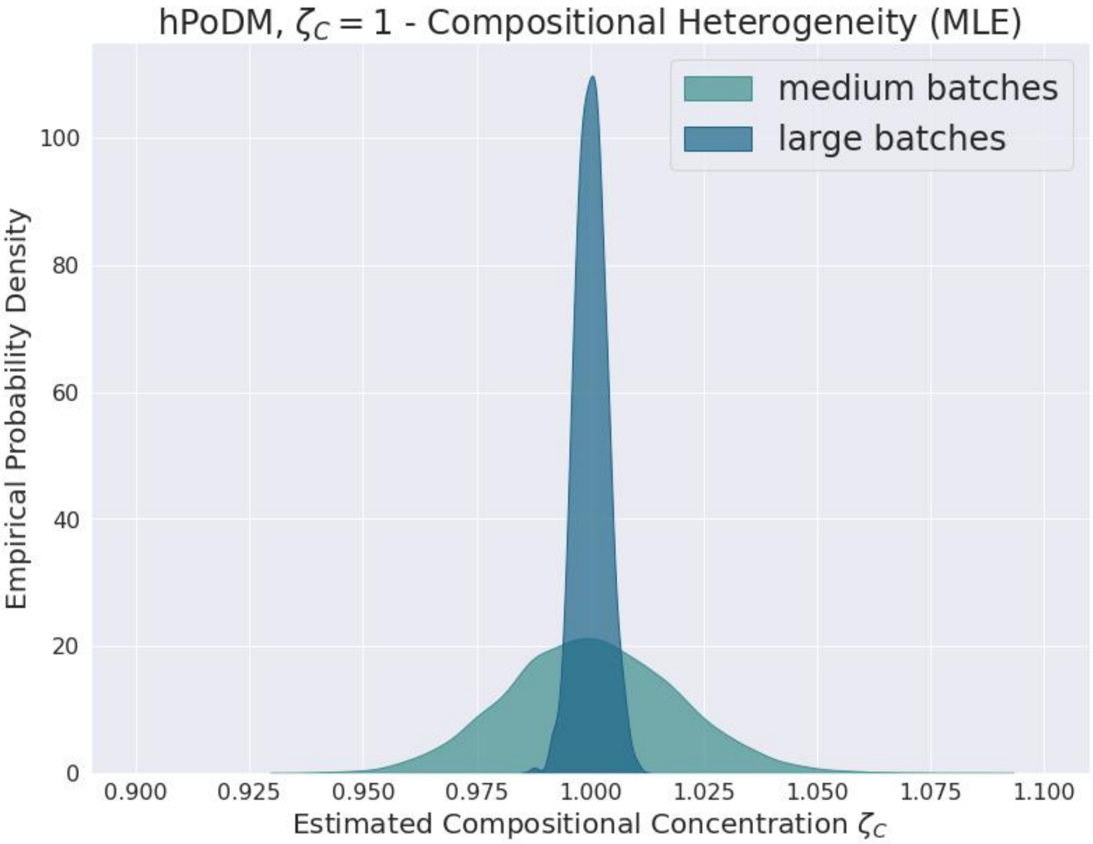

**Supplementary Figure S85**

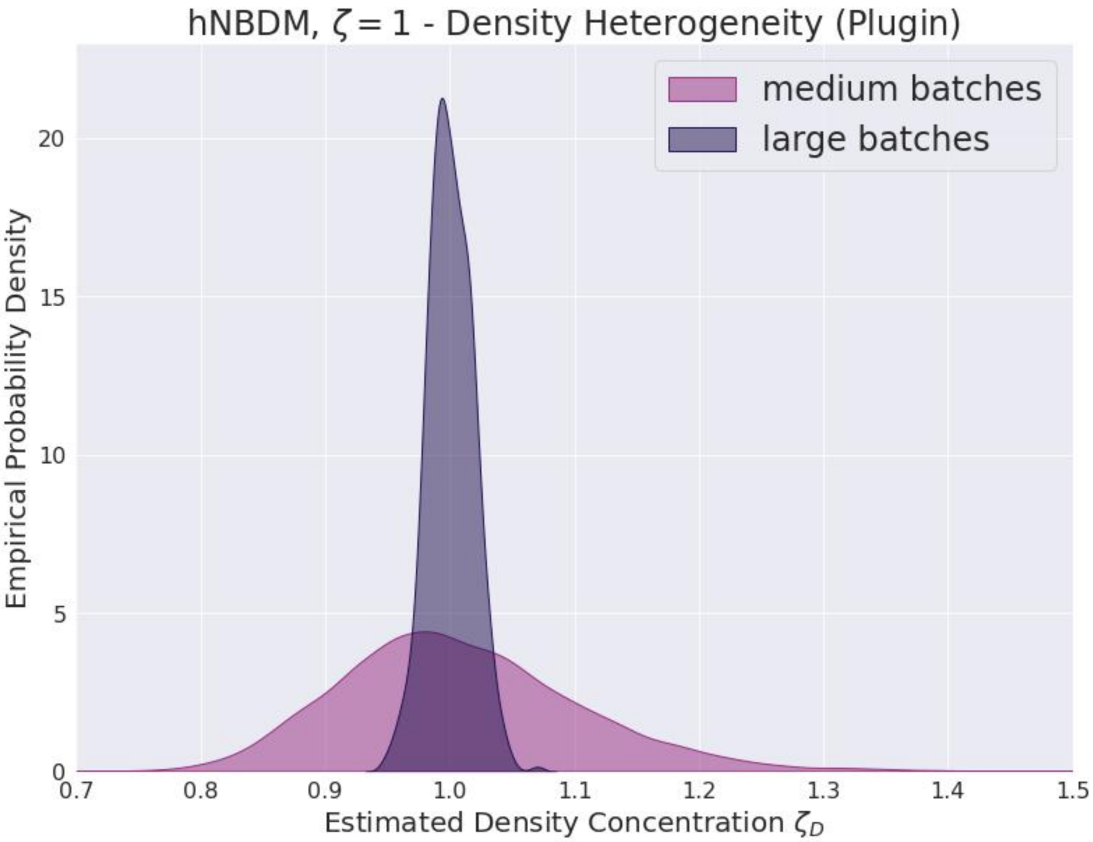

**Supplementary Figure S86**

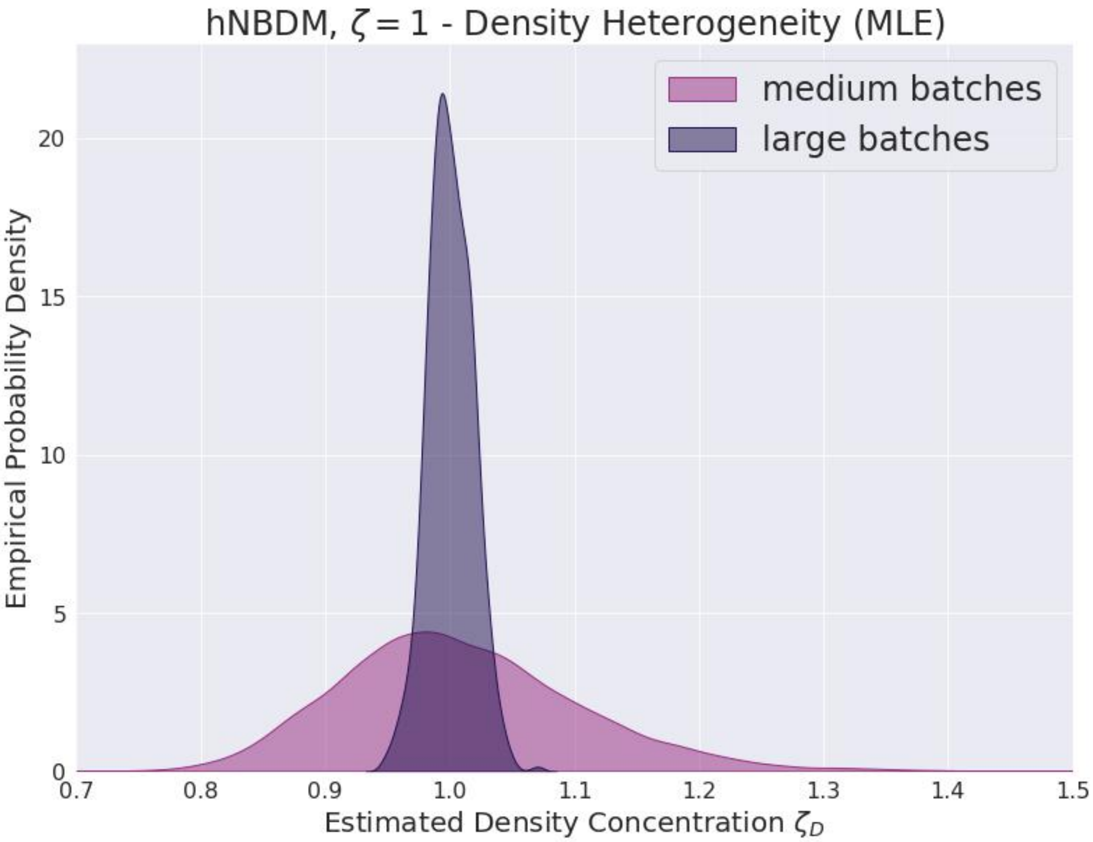

**Supplementary Figure S87**

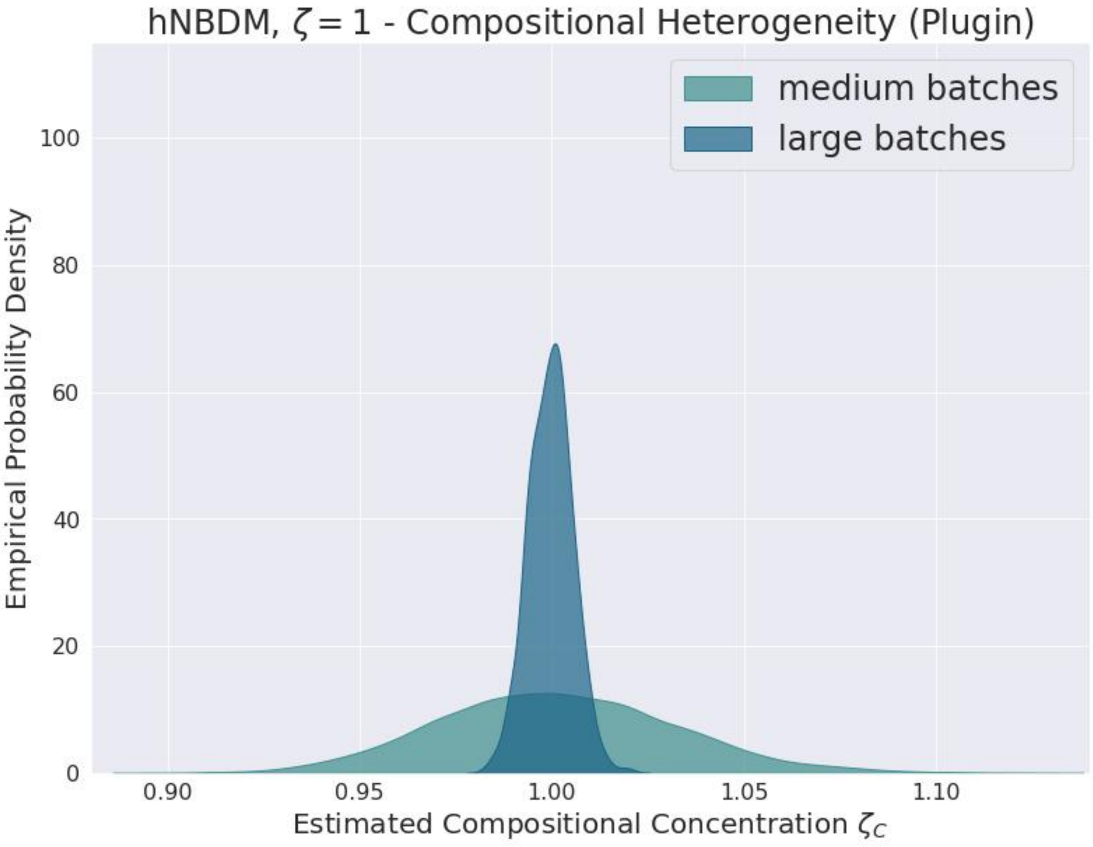

**Supplementary Figure S88**

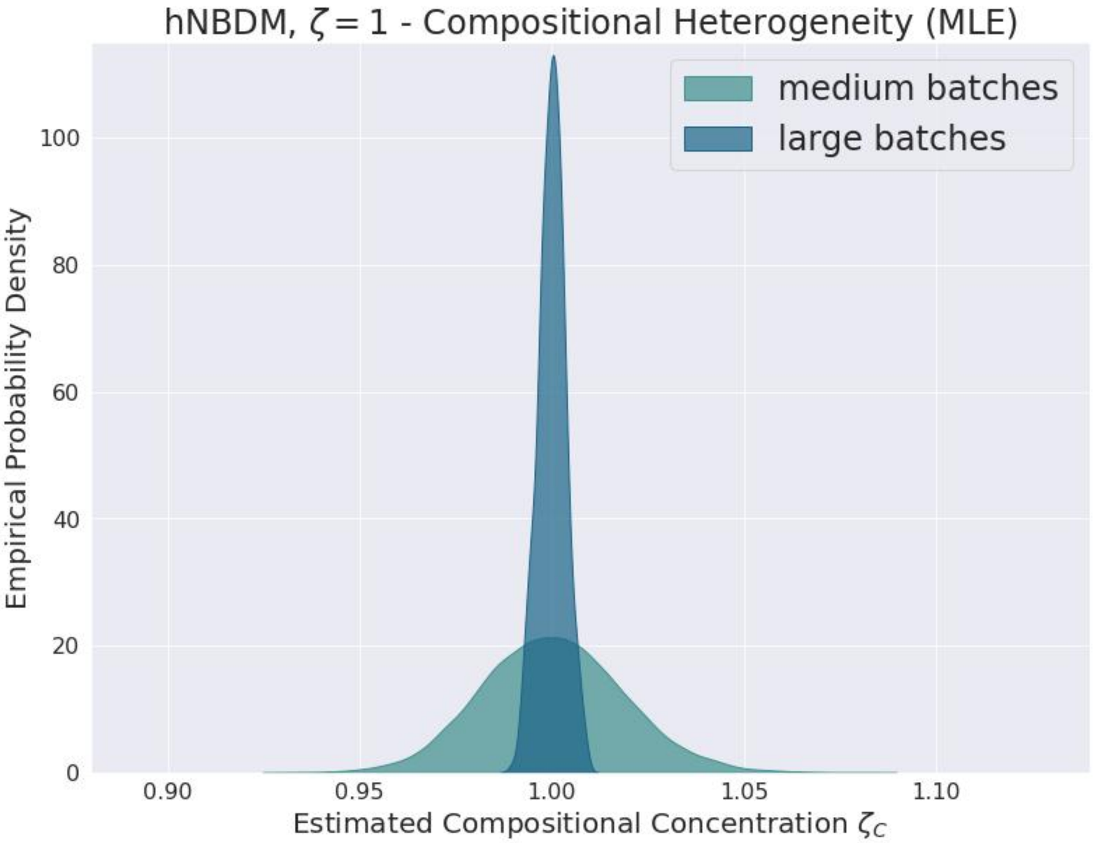

**Supplementary Figure S89**

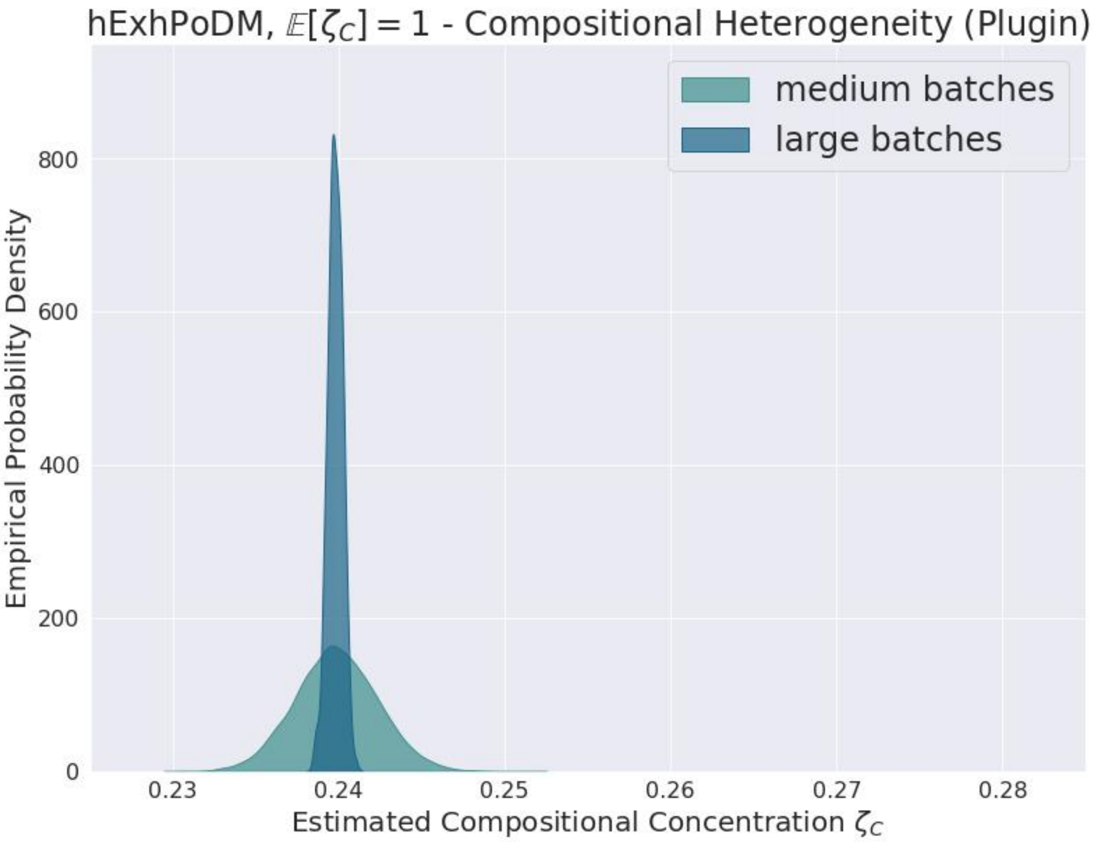

**Supplementary Figure S90**

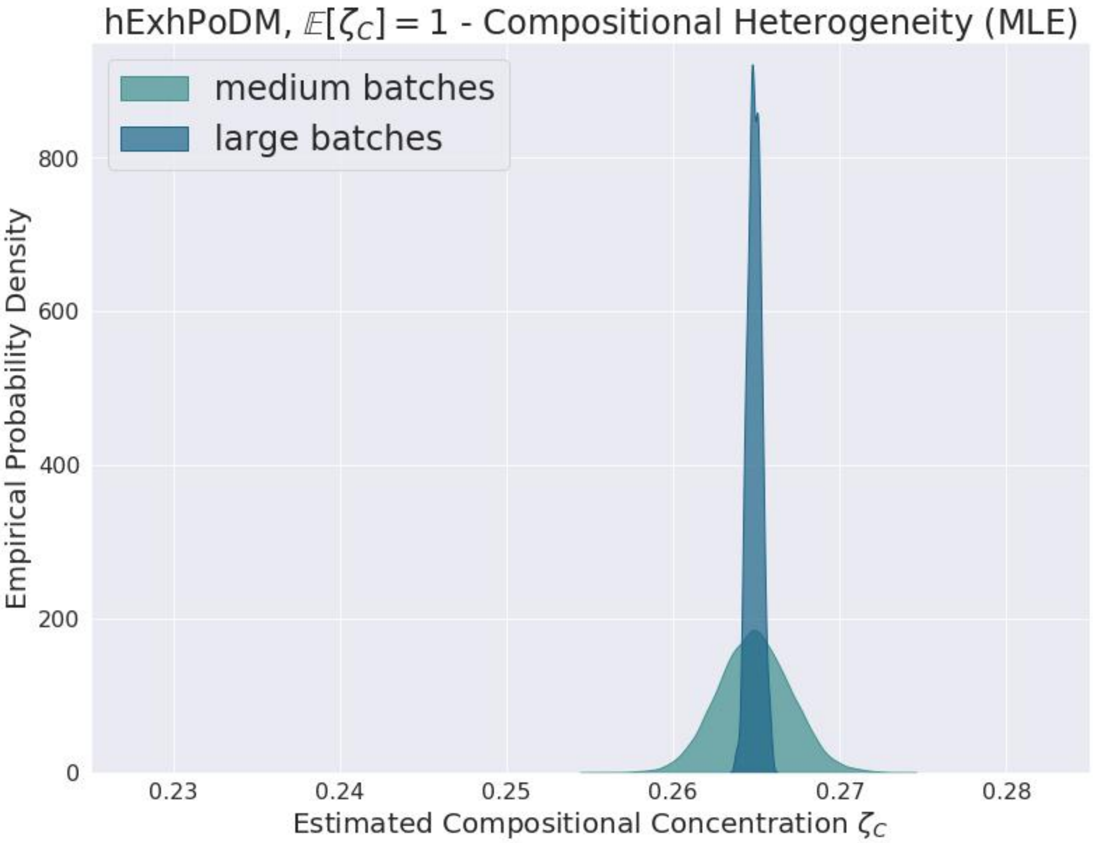

**Supplementary Figure S91**

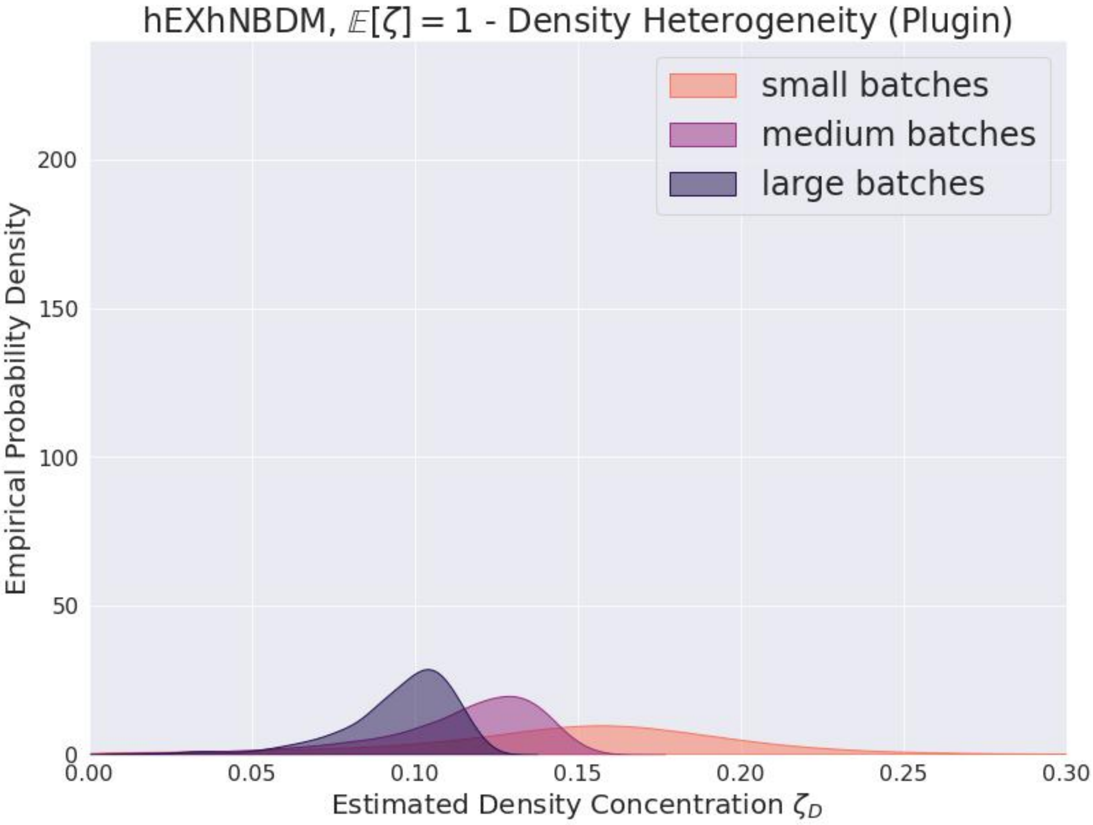

**Supplementary Figure S92**

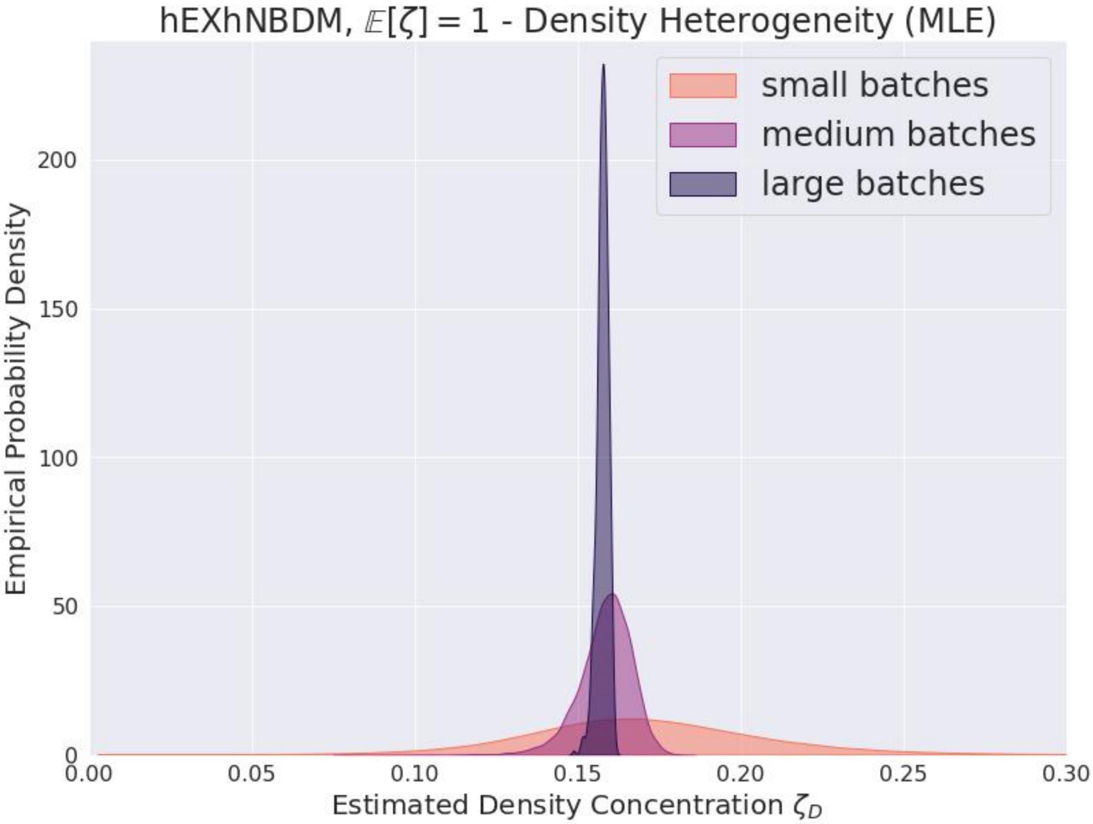

**Supplementary Figure S93**

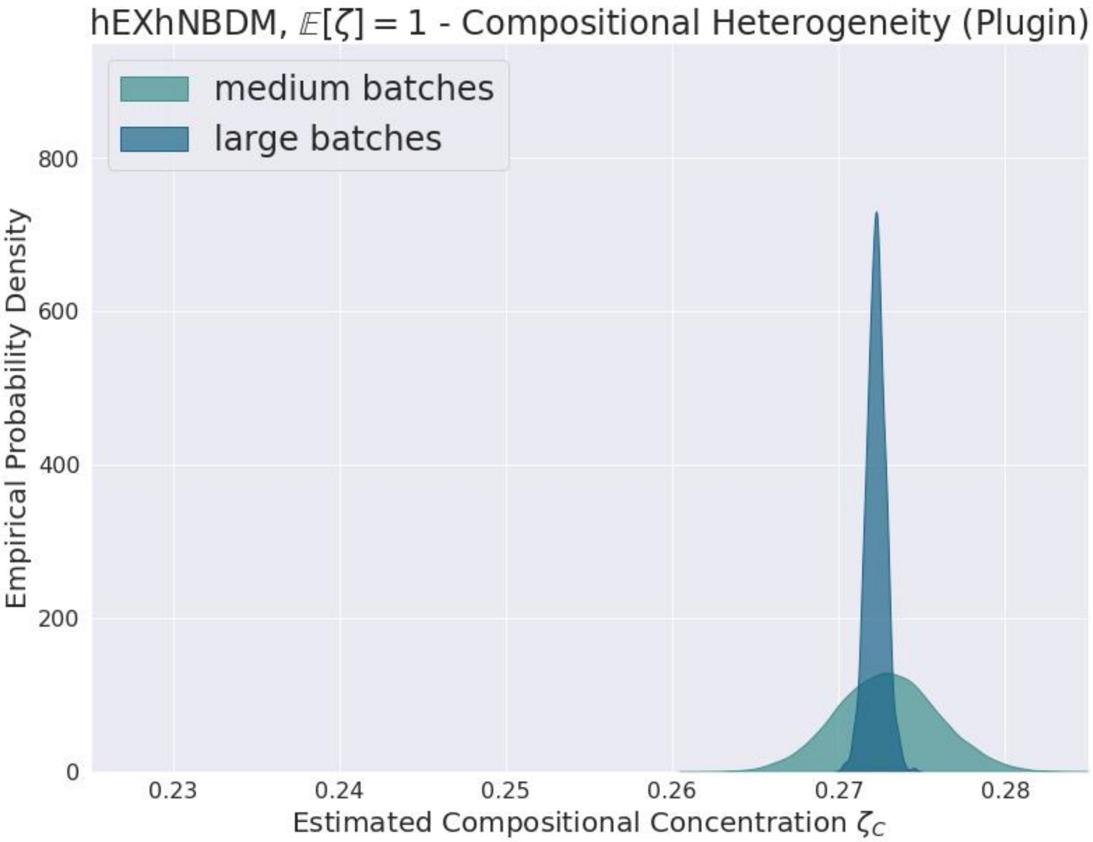

**Supplementary Figure S94**

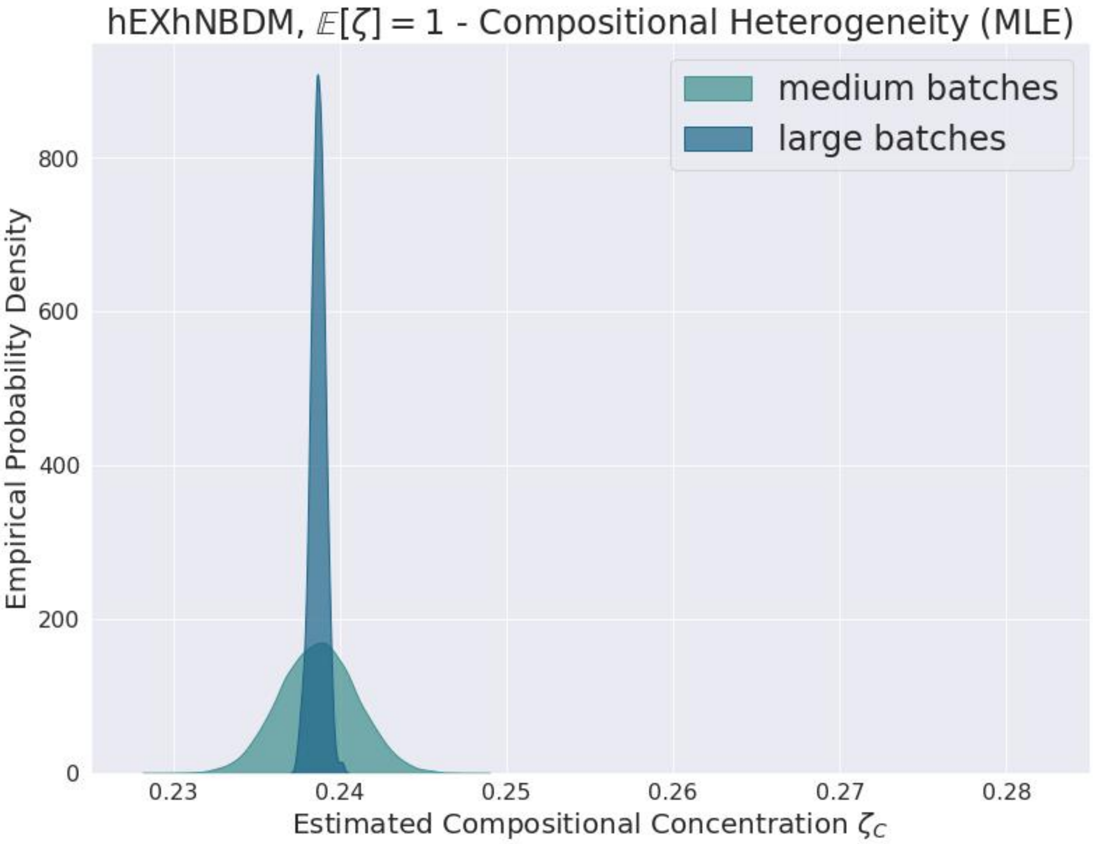

**Supplementary Figure S95**

(as computed in (Anscombe, 1950)) of the negative binomial ML and plugin estimators is negligible for the chosen sample sizes.

Bizarrely, hExhNBDM appeared to be the only distribution for which the ML density estimator was noticeably more efficient than the plugin density estimator, despite (or perhaps because?) of the model misspecification. Compare figures S92 and S93. I am unsure why this only occurred for hExhNBDM.

The bump in the density concentration estimates which consistently occurs around $10^4$ for the ML estimates appears to be an artifact of the difficulties encountered when relying on Powell's method (Powell, 1970) instead of Brent's method (Brent, 1973) to find the root of the score, cf. the discussion in section S5.3.2. See figures S53, S63, S55, S65, S75, S81, S57, S67, S59, and S69. These bumps probably would not occur with a better implementation of the MLE.

**Precision of ML Estimator vs. Plugin Estimator for Compositional concentration** Unlike the situation for the density estimators, the compositional ML estimator is noticeably more efficient than the compositional plugin estimator for all of the distributions, regardless of whether the model was correctly specified.

See figures S60 and S61 for survival curves when the true value of the estimand is infinite. See figures S72, S73, S76, and S77 for survival curves when the true value of the estimand is finite. Despite how difficult it usually is for most of the estimators to distinguish distributions with infinite concentration from those with concentration $\approx 100$, for the large batches the ML estimator remarkably produced almost no infinite estimates. The increased efficiency of the ML estimator is evident from the more rapid decrease compared to the plugin estimator of the survival curves when the true value of the estimand is finite.

See figures S70 and S71 for KDE plots when the true value of the estimand is infinite. The greater efficiency of the ML estimator is reflected by the distribution of finite estimates shifting further to the right more quickly. When the true value of the estimand is finite, as well as when the model is misspecified, the greater efficiency of the ML estimator compared to the plugin estimator is reflected by the narrower distribution of estimates. See figures S78, S79, S82, S83, S84, S85, S88, S89, S90, S91, S94, and S95.

**Behavior of ML Estimator Compared to Plugin Estimator Under Misspecification** Under the misspecified[38] models hExhPoDM and hExhNBDM, the effect on the limit of switching from plugin to ML was opposite for hExhPoDM compared to hExhNBDM. For the compositional concentration, switching from the plugin estimator to the ML estimator *increased* the limiting value of the estimates for hExhPoDM (cf. figures S90 and S91), while for hExhNBDM switching from the plugin estimator to the ML estimator *decreased* the limiting value for the estimates (cf. figures S94, and S95). Interestingly, for hExhNBDM the effect of switching from the plugin estimator to the ML estimator for the density concentration was opposite the effect for the compositional concentration; the limiting value of the estimates *increased* (cf. figures S92 and S93). The fact that the plugin and ML estimators converge to different values from each other under these distributions reflects the model misspecification.

#### S5.5 Discussion

Section S5.5.1 discusses why the results from section S5.4 are important for showing that estimates from the estimators will be useful in practice. Section S5.5.2 discusses how the results from section S5.4 suggest that future work might be able formulate density and compositional heterogeneities in a nonparametric context. Section S5.5.3 discusses how the results from section S5.4 bring us one step closer to the final goal of this thesis.

##### *S5.5.1 The Estimators are Good Enough for Use in Practice*

The above results show that the proposed estimators both (1) converge towards correct answers as the data size increases and (2) have decreasing variance as the data size increases. Therefore they satisfy the minimal requirements one generally expects of "decent" or "adequate" statistical estimators.

The estimators were already guaranteed to be consistent asymptotically when the statistical model was correctly specified. Nevertheless, that theoretical guarantee does not necessarily translate in practice into estimates usefully approximating the truth for realistic data sizes. (In this case realistic data sizes correspond to the medium or large batches.) Such a failure of "practical consistency" occurred, for example, for the version of the plugin compositional concentration estimator that equally weighted the estimates resulting from all pairs of strains. Consistency of an estimator need not imply efficiency of an

[38] Cf. footnote 36 above.

estimator. The above results show that the proposed estimators are in fact "practically consistent", not merely asymptotically consistent, and thus useful in practice. Not only that, but they often still managed to give useful results even for unrealistically small data sizes (the small batches).

###### *S5.5.2 Heterogeneity Might Correspond to a Nonparametric Definition*

The plugin estimators also gave sensible results in the cases of model misspecification. This suggests that functions of moments, which are at least similar to the "effective density concentration" (S108) and "effective compositional concentration" (S123) that these estimators converge to, may be useful for describing qualitative notions of "density heterogeneity" and "compositional heterogeneity" for general[39] count-categorical distributions. Exploring how, or whether, to define such notions precisely for more general distributions outside of the ghNBDM family would be a worthwhile goal for future work. The argument from section 4 of (Nakashima, 1997) that the plugin estimator (at least for the negative binomial) is only "semi-parametric" is similar.

It is worth noting however that, at least for hExhPoDM and hExhNBDM, the ML estimators were also "unreasonably effective" at giving sensible results even under model misspecification. Thus we might anticipate either that (1) a "nonparametric" or "semi-parametric" interpretation of their estimands also exists, or (2) that the ML estimators would not prove equally "unreasonably effective" for other misspecified models. (Nakashima, 1997) appears to argue the latter, at least with respect to the density concentration estimator. Unfortunately this work does not investigate the behavior of these estimators under misspecification enough to distinguish between these two possibilities, cf. footnote 26.

###### *S5.5.3 Conclusion*

This section shows that we have viable statistical estimators for density heterogeneity and compositional heterogeneity. Therefore, neither quantity needs to be treated as an "unfathomable unknown". Instead, we are able to actively account for them when simulating or analyzing the data from MOREI. Given real-world data, even of only moderate size, we can get useful estimates for concentration parameters describing realistic levels of heterogeneity. Using these estimates as the parameter values for our simulations, we can make simulations of MOREI more realistic. This in turn enhances our ability to find the methods that best recover true interaction networks from MOREI data.

##### S5.6 Plugin Estimators for Strain Population Frequencies

When $\mathbb{E}[\vec{\mathbf{N}}(0) \mid N(0)]$ is Multinomial or Dirichlet-Multinomial distributed, one has that for all $n \geq 1$ and all strains $s \in [S]$:

$$f^{(s)} = \frac{\mathbb{E}[N^{(s)}(0) \mid N(0) = n]}{n} . \tag{S135}$$

Averaging over all values of $n \geq 1$ leads to

$$f^{(s)} = \sum_{n \geq 1} \frac{\mathbb{E}[N^{(s)}(0) \mid N(0) = n]}{n} \mathbb{P}(N(0) = n \mid N(0) \geq 1) =: (f^{(s)})_{eff} . \tag{S136}$$

This leads to the following plugin estimators for the frequencies:

$$\hat{f}^{(s)} = \sum_{n \geq 1} \frac{\hat{\mu}[N^{(s)}(0) \mid N(0) = n]}{n} \cdot \frac{\sum_d \mathbb{I}\{N_d(0) = n\}}{\sum_d \mathbb{I}\{N_d(0) \geq 1\}} . \tag{S137}$$

###### *S5.6.1 Effective Frequencies*

Even when the left and right hand sides of (S136) are not equal, the plugin estimator (S137) is still consistent for the right hand side of (S136), the "effective frequency" $(f^{(s)})_{eff}$.

Note that for all of the distributions used in the simulations, the true frequencies *do* equal the effective frequencies. Even for hExhNBDM and hExhPoDM this is true, because

$$f^{(s)} = \int_{\xi > 0} f^{(s)} \mathbb{P}(\zeta_C = \xi) \mathrm{d}\xi = \int_{\xi > 0} (f^{(s)})_{eff} \mathbb{P}(\zeta_C = \xi) \mathrm{d}\xi = (f^{(s)})_{eff} . \tag{S138}$$

[39]It probably makes sense only to attempt to define such notions for distributions with finite second moments, or to at least automatically say that any distribution lacking finite second moments is "infinitely heterogeneous" without exploring further.

In other words, (S136) holds for any individual value of $\zeta_C$ and will therefore still hold after averaging over any choice of "prior" for $\zeta_C$.

For distributions such that $f^{(s)} \neq (f^{(s)})_{eff}$, the estimators (S137) will no longer be consistent, and a different choice of estimators $\hat{f}^{(s)}$ which actually are consistent for the true frequencies $f^{(s)}$ will need to be used inside of (S122) to guarantee that (S121) remains consistent for the effective compositional concentration (S123).

Alternatively, one could instead change the definition of the effective compositional concentration inside of (S124) to use the effective frequencies $(f^{(s)})_{eff}$ in place of the true frequencies $f^{(s)}$, in which case using (S137) inside of (S122) will continue to ensure that (S121) is consistent for (S123). Cf. section S5.2.3.

#### S5.7 Convergence Sometimes Disappointing for Small Batches

The variance of the distribution of estimates was sometimes much wider for the small batches than for the medium or large batches. This is unimportant in practice since the small batches represent an unrealistically small problem size.

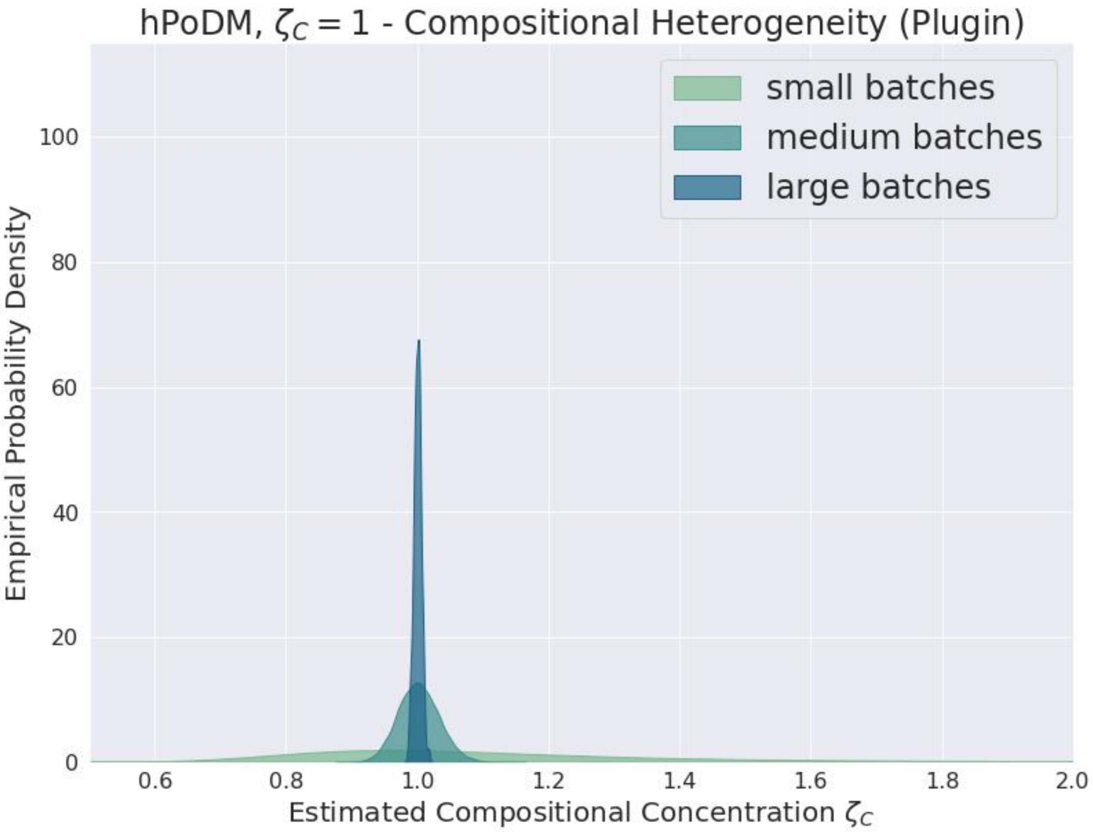

**Supplementary Figure S96**

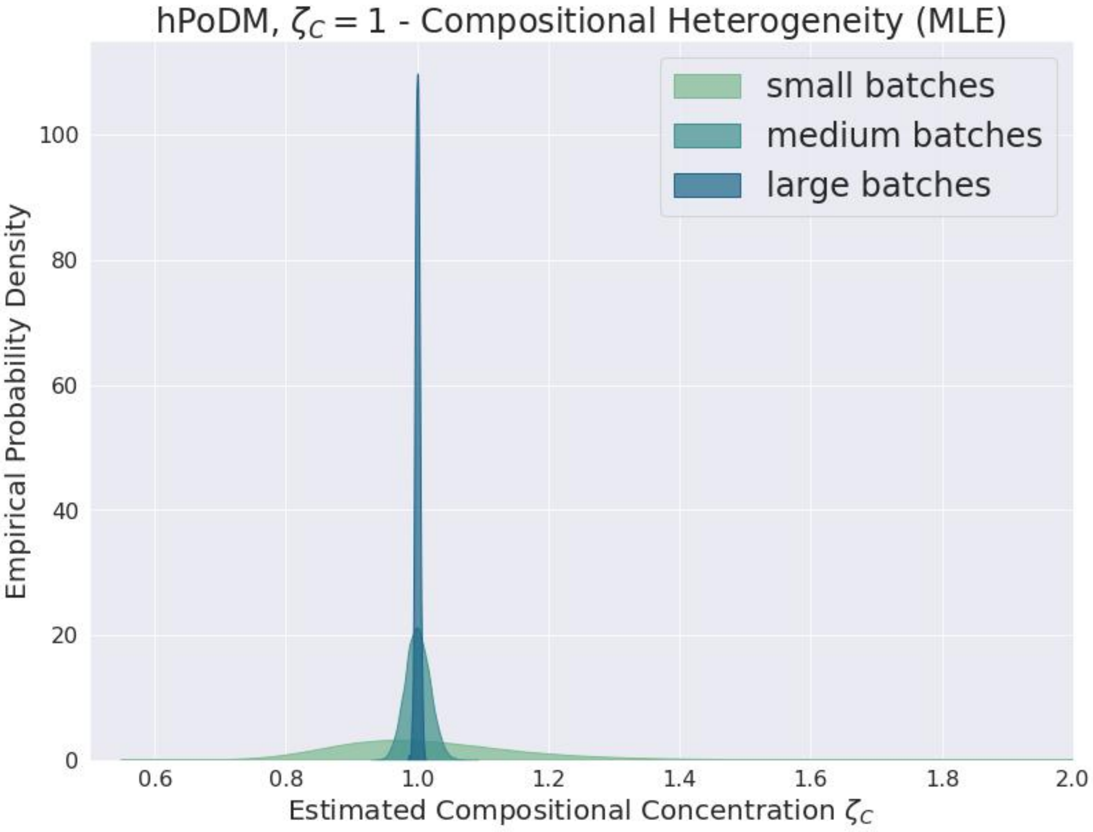

**Supplementary Figure S97**

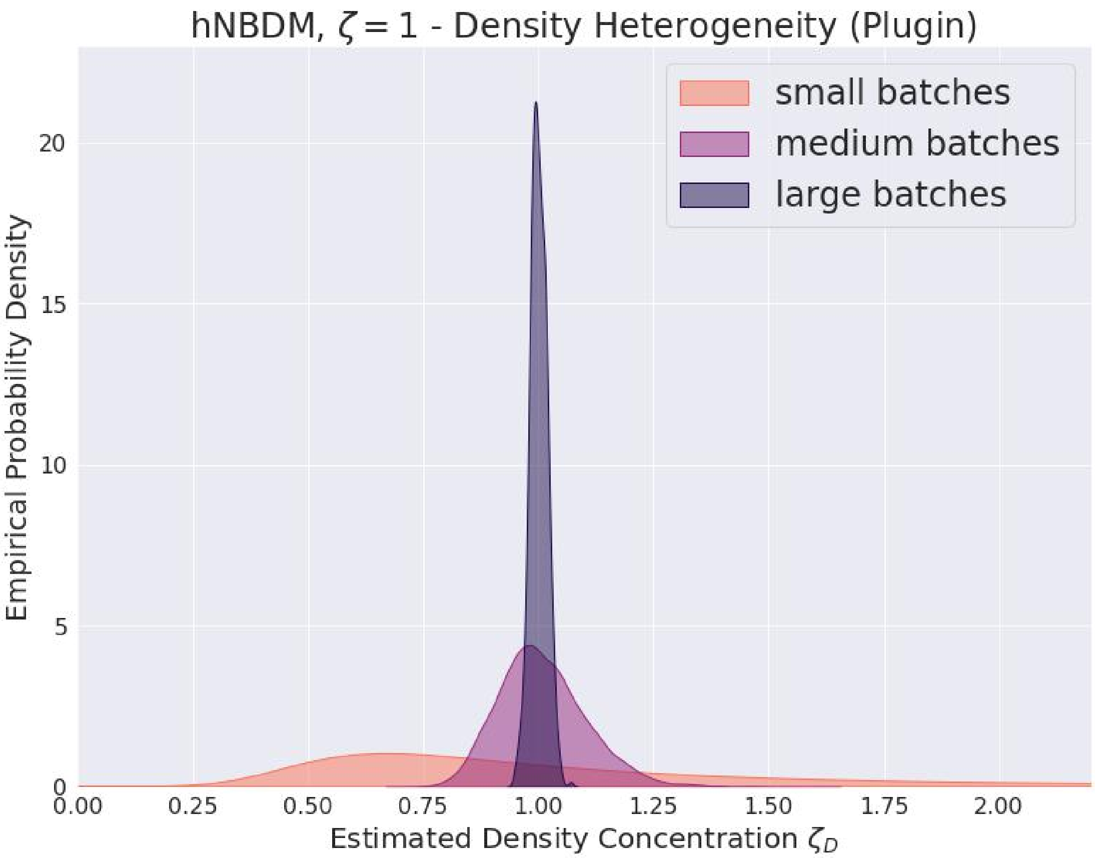

**Supplementary Figure S98**

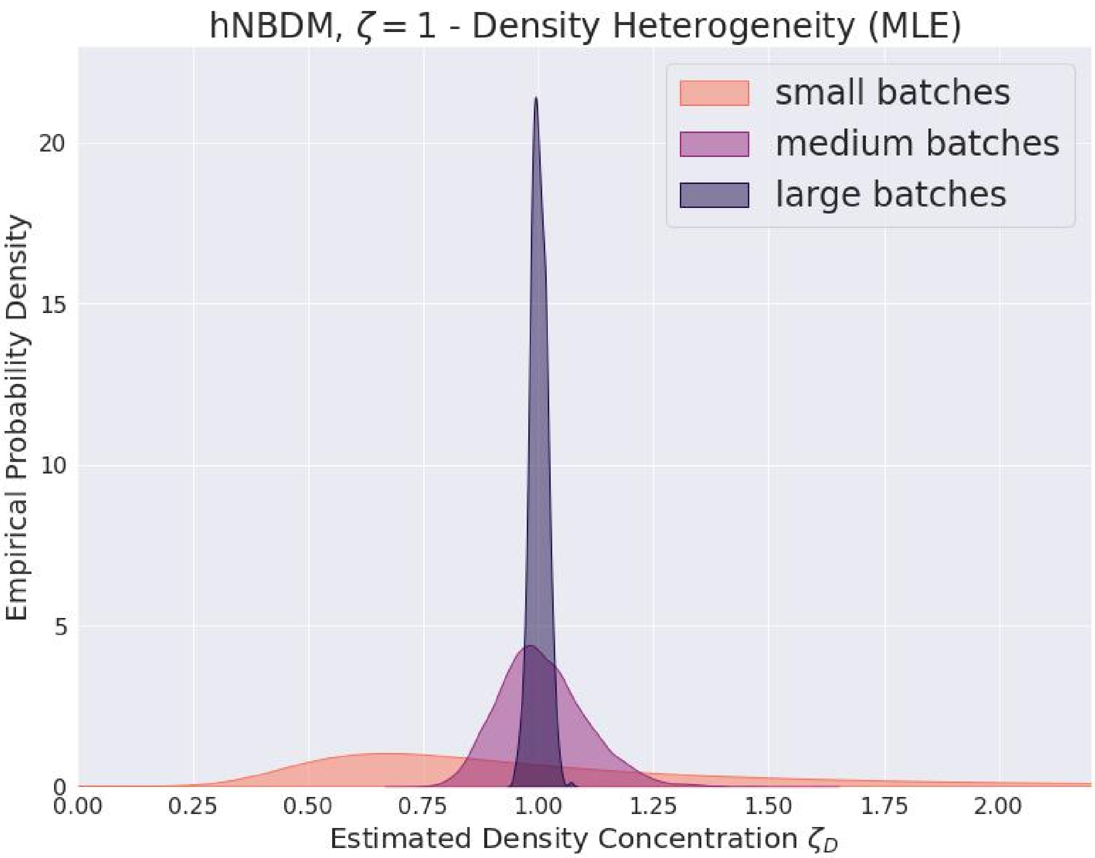

**Supplementary Figure S99**

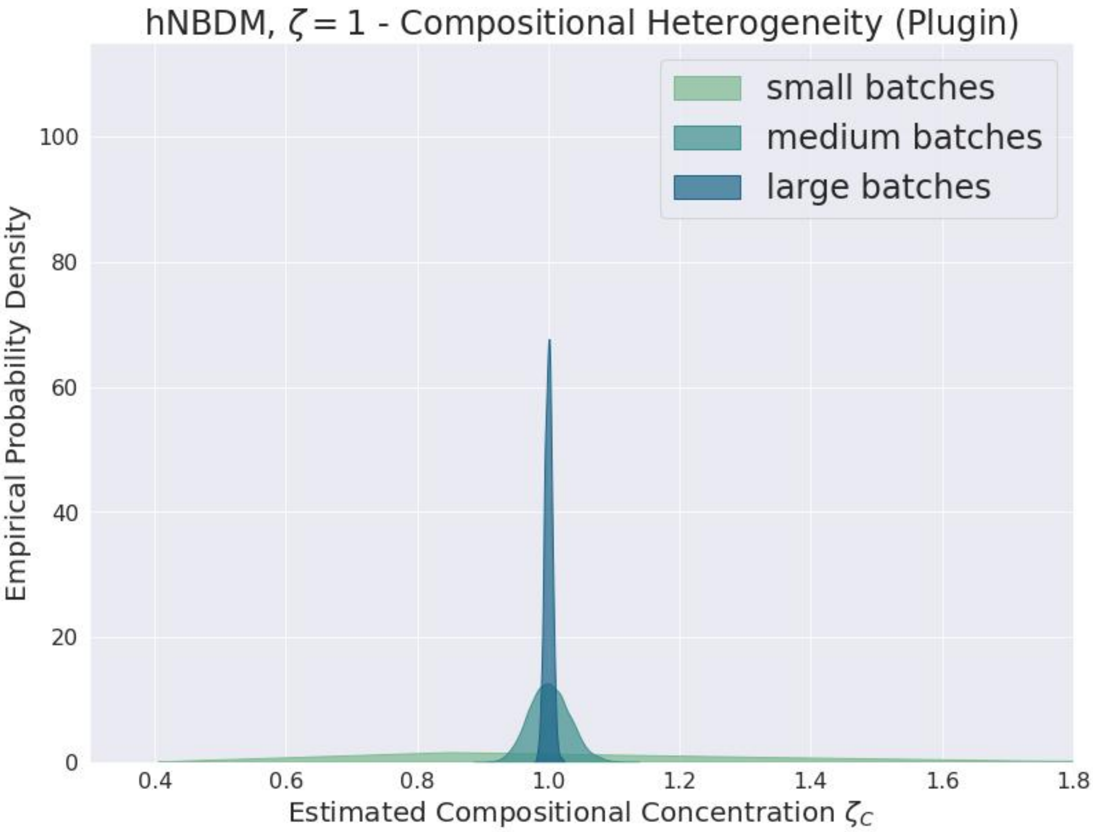

**Supplementary Figure S100**

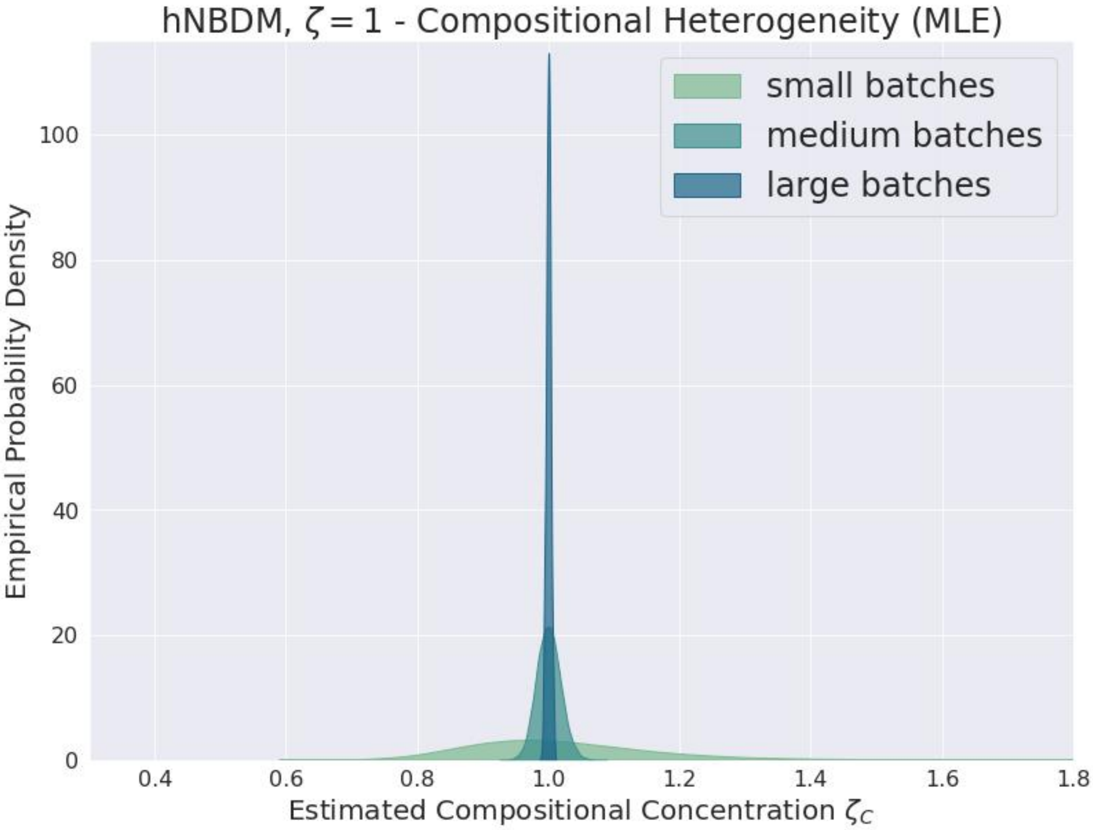

**Supplementary Figure S101**

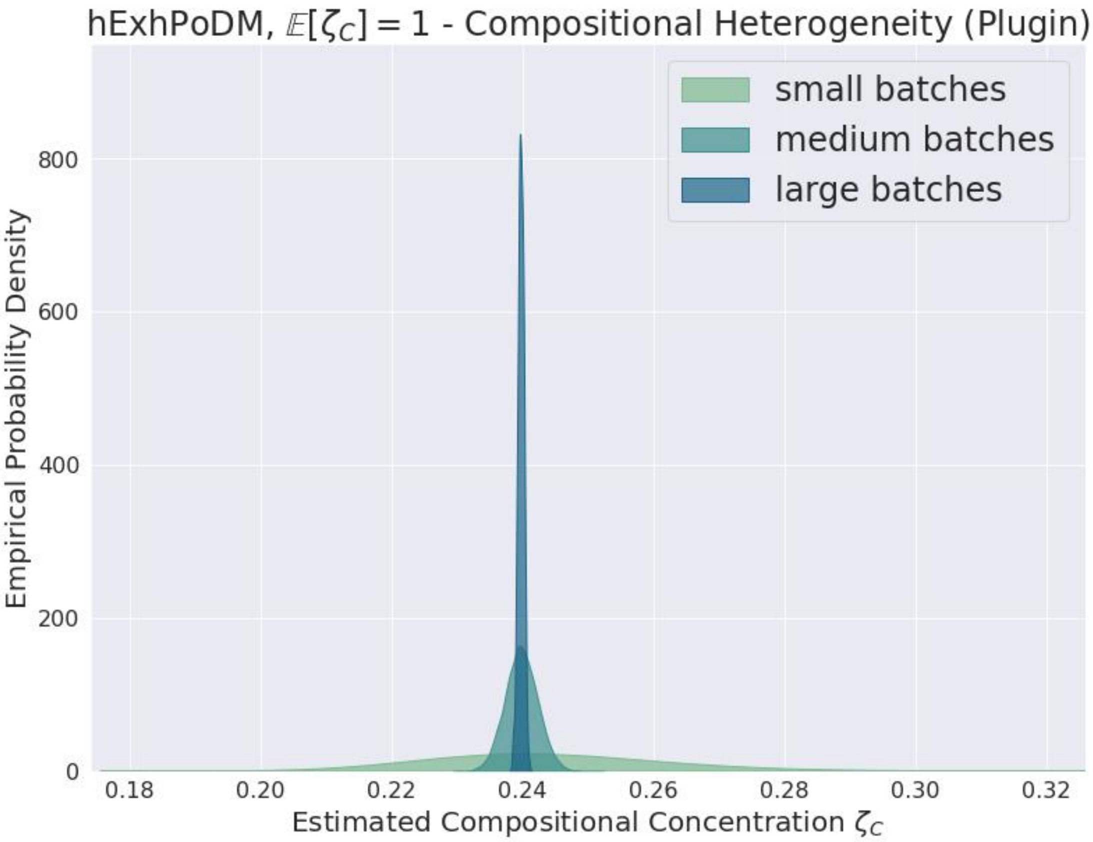

**Supplementary Figure S102**

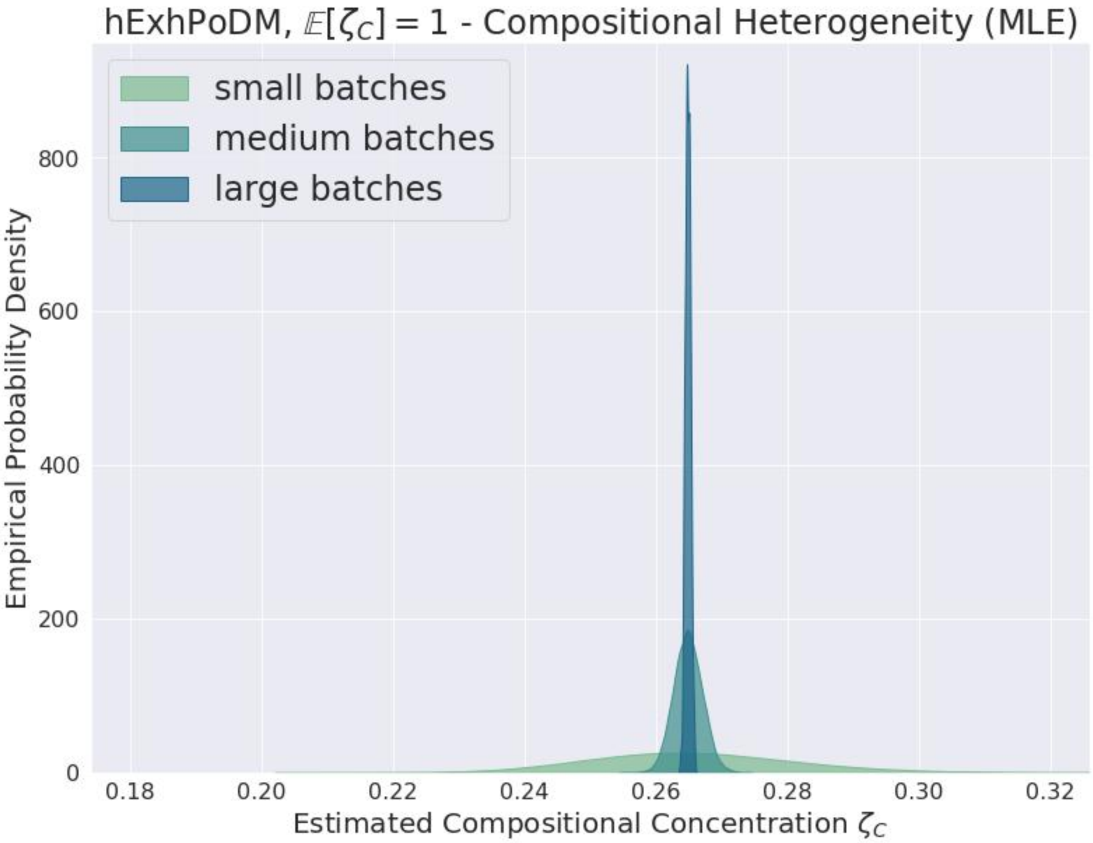

**Supplementary Figure S103**

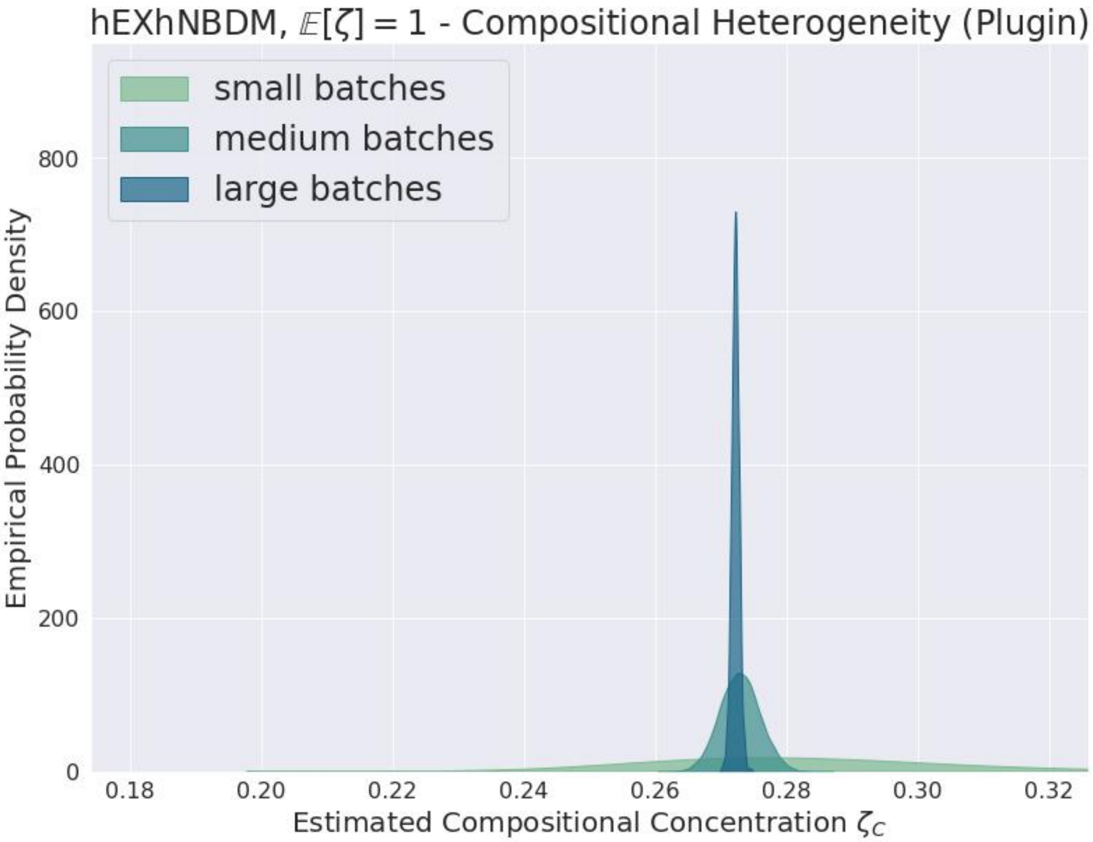

**Supplementary Figure S104**

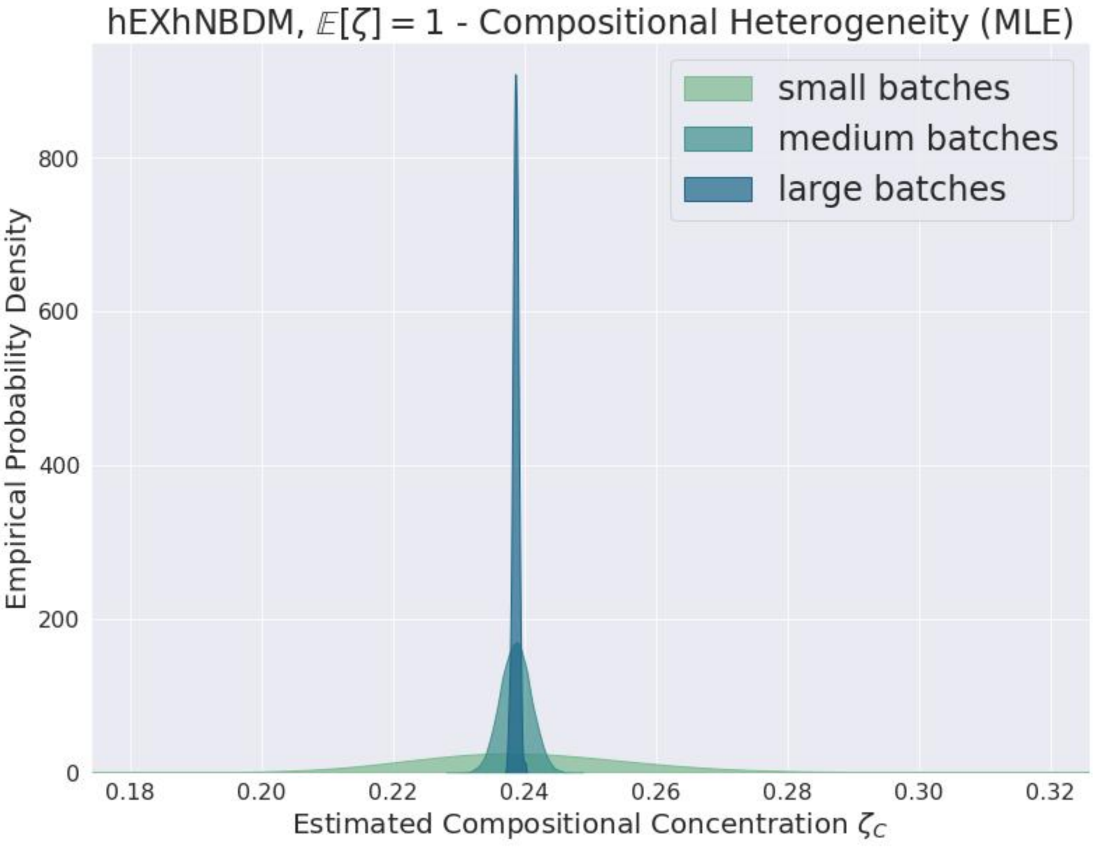

**Supplementary Figure S105**
